# Multiple nucleotide polymorphism DNA markers for the accurate evaluation of genetic variations

**DOI:** 10.1101/2021.03.09.434561

**Authors:** Zhiwei Fang, Lun Li, Junfei Zhou, Aiqing You, Lifen Gao, Tiantian Li, Hong Chen, Ruixi Han, Yehan Cui, Lihong Chen, Huafeng Xiao, Jing Zhang, Na Xu, Xiqin Fu, Jianan Zhang, Xiuting Li, Aijin Ma, Weixiong Zhang, Hai Peng

## Abstract

DNA markers are an essential tool for the detection and evaluation of genetic variations, a central theme in genetics and biology. Effective markers must be highly reproducible, polymorphic, accurate and efficient to profile. We developed *multiple dispersed nucleotide polymorphism* (MNP) DNA marker and an efficient MNP genotyping method called *MNP-Seq*. The MNP marker was 17.48% more polymorphic than the highly polymorphic marker of microsatellites on a collection of hybrid rice plants. When applied to genotype more than 80,000 individual MNP markers of diploid rice and polyploidy hybrid cotton varieties which were notoriously difficult to genotype accurately, MNP-Seq finished in two days and achieved accuracies of 99.999% and 99.988%, respectively. We adopted MNP-Seq to reveal the ubiquitous, albeit subtle and neglected, genetic heterogeneities in homonyms of Nipponbare rice, a popular model organism for plant biology. This result raised a question on the consistency of the published results using the model plant. We also used MNP-Seq to accurately and efficiently determine the identities of plant varieties, a key but difficult problem for the protection of plant intellectual property rights. While being applied to plants in the current study, the MNP marker and MNP-Seq are general and readily applicable to similar problems in animals and micro-organisms.

## Introduction

DNA markers, fragments designated to specific genomic loci, are an important tool for studying various biological problems such as genetic diversity, evolution and speciation, and genetic linkage, and can be used for, e.g., genetic fingerprinting, forensic analysis, and genomic breeding^1-5^. To support such broad applications, it is imperative to accurately evaluate genetic variations using DNA markers that are highly reproducible, polymorphic, accurate and efficient to genotype.

Among the early DNA markers are random amplified polymorphic DNA (RAPD), inter simple sequence repeat (ISSR), restriction fragment length polymorphism (RFLP), and amplified fragment length polymorphism (AFLP)^6-8^. The profiling of these markers is complicated. Their results are often determined by the presence or absence of specific DNA fragments that are derived from the restriction digestion or amplification of oligonucleotide primers. Technical expertise and strict experimental conditions, such as enzymic digestion and electrophoresis, are required. Therefore, it is difficult to ensure the genotyping reproducibility and accuracy of these markers.

One type of popular DNA markers is the simple sequence repeats (SSRs), also dubbed as short tandem repeats or microsatellites. An SSR is a fragment of repetitive DNA where a motif of one to six base pairs is repeated in tandem for 5 to 50 times^9^. The copy number variation of the motif gives rise to high polymorphism. However, repetitive SSRs can induce DNA polymerase slippage during polymerase chain reaction (PCR)^10^, which introduces erroneous SSR alleles to downstream analysis and consequently increases genotyping error as high as e.g., 5%^11,12^.

Moreover, the size of the amplicon of an SSR marker is typically determined by laborious electrophoresis^13-15^, resulting in low genotyping efficiency. The slippage error and low efficiency have been addressed in specific applications, e.g., forensic analysis, by designing species- and application-specific markers and optimizing experimental conditions^16,17^. Such a scheme is infeasible in general due to high cost.

Another common DNA marker is the single nucleotide polymorphisms (SNPs). As SNPs are mostly biallelic and highly abundant in the genome, a high-throughput genotyping method, e.g., microarray or whole-genome sequencing (WGS)^18-21^, is used to detect a large number of SNPs to guarantee the accuracy for the estimation of genetic variations. However, microarray has a typical error of 1%, which increases to as high as 6.1% on hybrid plants^20-22^. Besides, microarray has intrinsic technical issues of missing data, low sensitivity, and poor reproducibility^21,23^. A WGS method requires to sequence the genome, which limits the sequencing depth with acceptable cost, resulting in a genotyping error of 0.9% to 7.9%^18,19,24^. While a strategy of random DNA barcode has been introduced to increase genotyping accuracy^25,26^, the high cost and heavy workload restrict the strategy to a limited number of SNPs and introduce sampling biases to SNPs on the genome.

For polyploidy plants, the detection of multiple alleles on one locus, particularly minor alleles, is required but technically challenging using the existing markers. For example, for SSR electrophoresis and SNP microarray, 30 or more PCR amplification cycles are typically needed to produce enough signals for adequate visual inspection. Such a large amount of amplification may distort the actual proportions of marker alleles because of nonlinear PCR amplification, resulting in inaccurate genotyping. The situation is exacerbated by hybrid samples and samples from a population that has a complex genetic background and high genetic heterogeneity. It should be noted that about one-third of flowering plants^27^ and many crops such as cotton are hybrid and polyploidy, and the derived samples often form a population.

We developed a novel type of DNA marker named *multiple dispersed nucleotide polymorphisms* (MNP) and an efficient MNP-genotyping method, called *MNP-Seq*, to accurately detect and estimate genetic variations, particularly for polyploid hybrid plants. We analyzed the eminent features of the MNP marker and MNP-Seq on rice and cotton, including reproducibility, polymorphism, accuracy, and efficiency. We also applied them to two case studies, each of which addressed a challenging problem in plant biology and agriculture, to demonstrate their utility and power. While we only applied the MNP marker and MNP-Seq to plants, they are a generic tool that is readily applicable to animal species.

## Results

### Genotyping MNP markers

An MNP is a set of multiple dispersed SNPs within a genomic segment of less than 300 bp in length. A total of 930 and 943 MNP markers for rice and cotton, respectively, were designed and used in this study (Supplementary Table 1). The goal of development of MNPs marker is to discriminate varieties for the targeted species and the discriminative power was determined by marker’s polymorphism, number of markers and accuracy of genotyping. Thus a similar number of MNP were selected for these species despite the big difference in genome size, and same MNP locus could be used for other varieties. A method for genotyping MNP markers, named as MNP-Seq, was developed (Fig. 1). First, a multiplex PCR reaction was performed to enrich all the designed MNP markers from the genomic DNA of a sample to be genotyped. A unique DNA barcode was then added to the PCR products of the sample to construct a sequencing library. The libraries for all samples were pooled and sequenced by a high-throughput sequencing method. The sequencing reads were partitioned into the samples according to their unique DNA barcodes and mapped to all MNP loci for MNP genotyping (see Method).

**Table 1.** The reproducibility and accuracy of MNP-Seq.

| Species | Ploidy | Heterozygosity | Variety number | MNP markers | Compared genotype pairs |  | Reproducibility | Accuracy | Erroneous genotypes |  |
| --- | --- | --- | --- | --- | --- | --- | --- | --- | --- | --- |
|  |  |  |  |  | Total | Reproducible |  |  | Number per sample | Ratio |
| Rice | Diploid | Hybrid * | 47 | 930 | 127,875 | 127,872 | 99.998% | 99.999% | 0.009 | 0.001% |
| Cotton | Tetraploid | Hybrid | 43 | 943 | 116,700 | 116,671 | 99.975% | 99.988% | 0.113 | 0.012% |

\*for 46 of 47 rice varieties.

**Figure 1.**
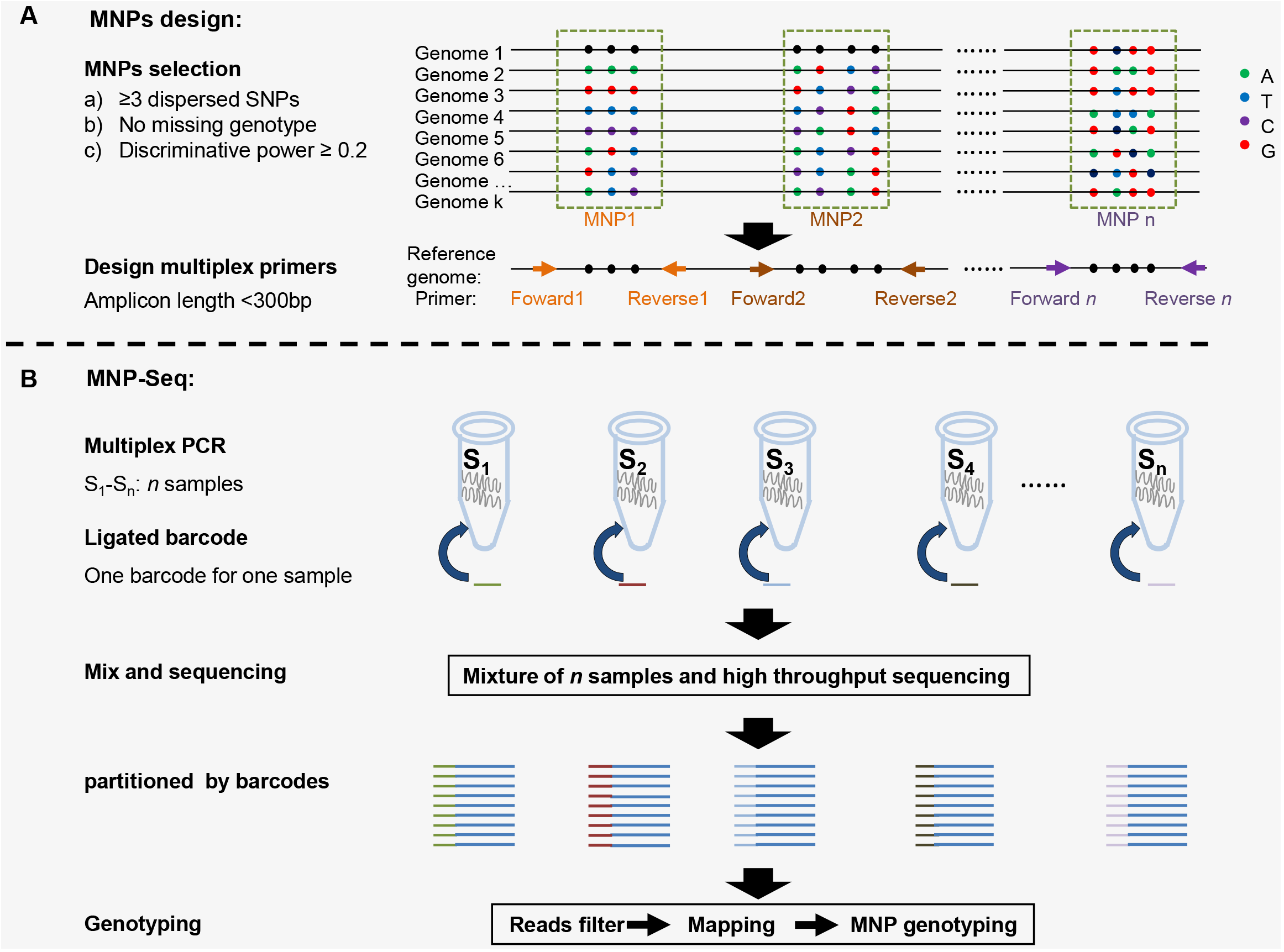
The procedure of MNPs design and MNP-Seq. **(A)** The development of MNP markers involves two steps, first to identify genomic loci of candidate MNPs that have diverse alleles across plant varieties and then to refine the candidate MNPs to accommodate effective PCR primers. Each colorful dot represents an SNP site on k genome sequences. **(B)** The major steps of MNP-Seq genotyping method, including library preparation, sequencing and bioinformatics analysis.

A total of 196,310,412 reads were produced from the technical triplets for the 47 rice and 43 cotton varieties (see Method), and each of the MNP loci for these varieties was, respectively, covered by 2,739 and 1,918 times in each replicate on average (Supplementary Table 2). Besides, an average of 924 (99.40%) of the 930 rice MNPs and 936 (99.21%) of the 943 cotton MNPs were genotyped for each variety in each replicate (Supplementary Table 2), showing a low missing rate of MNP-Seq. The MNP-Seq method was also efficient; we were able to genotype more than 80,000 MNP markers (47 rice X 930 markers plus 936 markers x 43 cottons) of all the varieties in one of the technical triplets within 48 hours, a significant improvement over the conventional methods. Due to the high reproducibility and accuracy of MNP-seq (see next), no replicate was required for the determination of MNP genotype in practice.

**Table 2.** Three major maker types and their genotyping features and performance.

| Marker type | MNPs | SSRs |  |  | SNPs |  |
| --- | --- | --- | --- | --- | --- | --- |
| Genotyping method | MNP-seq | SSR-seq | Sanger sequencing | Capillary electrophoresis | WGS | Chip array |
| Polymorphism | Highest | Higher | Higher | Higher | Lowest | Lowest |
| Accuracy | Highest | Higher | High | Middle | Low | General |
| Efficiency | High | High | Lowest | Low | Highest | Higher |
| Genetic discriminative power using the same number of markers | Highest | Higher | High | Middle | Low | Low |
| Forms of the genotype | Base | Base | Base | Amplicon length | Base | Base |
| DNA polymerase slippage | No | Yes | Yes | Yes | No | No |
| Genotyping cost per maker locus | Middle | Middle | Highest | High | Lowest | Low |
| The time required for genotyping | Short | Short | Short | Shortest | Long | Long |
| Parallel experiment | No need | No need | No need | Required | No need | No need |
| Amount of template DNA needed to genotype the same number of markers | Low | Low | Highest | Higher | Lowest | lower |

### High polymorphism and discriminative power of MNP markers

The SSR marker is well known for high polymorphism. We assessed the polymorphism of the MNP marker by genotyping 39 rice varieties and then comparing their 930 MNP markers and the 412 highly polymorphic SSR markers. On average, each MNP had 3.36 ± 1.13 alleles, and in comparison, each SSR had 2.86 ± 1.54 alleles (Supplementary Table 3), showing the former was 17.48% more polymorphic than the latter (p = 1.59 x 10^−10^, Student’s t-test). The higher polymorphism of the MNP markers provided a higher discriminative power than the SSR markers (Fig. 2A). Indeed, 53.66% ± 15.38% of the 930 MNP markers had different genotypes between varieties, and in contrast, 44.67% ± 10.91% of the 412 SSR markers had different genotypes. It is interesting to note that a strong correlation (r=0.94, Pearson correlation coefficient) could be observed between the distances calculated based on MNPs and SSRs for any two varieties [Fig. 2A].

**Figure 2.**
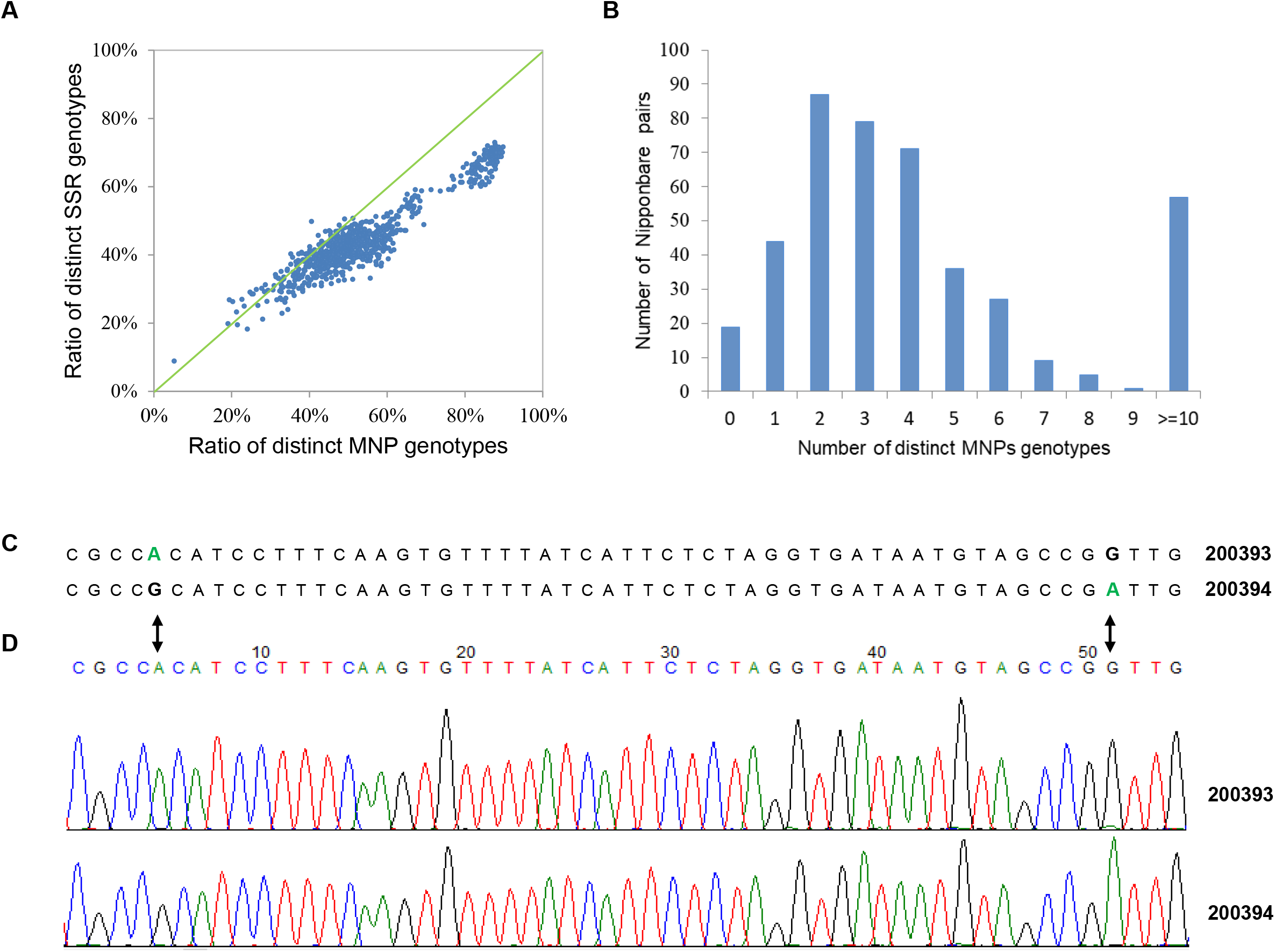
The discriminative power of rice MNP markers and MNP-Seq accuracy. **(A)** Comparison between the ratios of the distinct rice MNP and SSR genotypes. Each dot represented a pair from the 39 rice varieties (Supplementary Table 6). The majority of the dots are below the diagonal line, showing that the MNPs are more discriminative than the SSRs on average. **(B)** The distribution of distinct MNP genotypes among the 30 Nipponbare lines from different labs. **(C)** The distinct genotypes between Nipponbare line 200393 and 200394 on locus AMPL1563256 determined by MNP-seq. The variant bases were listed in bold. **(D)** Validation of the variants shown in (C) by Sanger sequencing. The corresponding variants are marked with arrows.

### High reproducibility and accuracy of MNP-Seq

Beyond polymorphism, reproducibility and accuracy are also essential for a genotyping method. Here the reproducibility was estimated by comparing the genotypes of each MNP locus between each two of the triplets separately. We used the data of the technical triplets to compare a total of 127,875 and 116,700 MNP pairs of genotypes of rice and cotton varieties, respectively. Among these MNPs, 99.998% (127,872) and 99.975% (116,671) were reproducible for rice and cotton varieties (Table 1), respectively. Under the assumption that genotyping errors ware independent and identically distributed, we estimated the accuracies of MNP-Seq to be 99.999% and 99.988% for hybrid rice and polyploidy hybrid cotton varieties, respectively (Table 1).Thus accurate genotypes could be got by only once experiment in practice and no cost need to afford for repeats. For comparison, these high accuracies improved upon the reported accuracies of SSR (94.47%)^12^ and SNP (94.3% - 99.1%) genotyping methods^19,20^. It should be highlighted that the population (e.g., plant varieties), polyploidy, and hybrid samples considered in this study were difficult for accurate genotyping.

### Applications of MNP marker and MNP-Seq

Taking advantage of the high polymorphism of MNP markers, high reproducibility, accuracy and efficiency of MNP-Seq, we applied them to two problems where it is critically important to accurately identify subtle genetic variations or to efficiently and conveniently compare DNA fingerprints.

#### Detection and analysis of genetic homonyms for ensuring scientific reproducibility

As the first sequenced model plant of *Gramineae*, Nipponbare has been widely used in plant biology and agriculture research. In this study, we collected 30 Nipponbare lines from different labs at geographically diverse locations in China (Supplementary Table 4), and constructed and compared their DNA fingerprints of the 930 rice MNP makers (see Method). We compared the fingerprints of each two of 30 Nipponbare lines, and a total of 435 pairs of fingerprints were compared. 416 (95.63%) of 435 pairs had differences of 0.11% to 12.76% (Fig. 2B and Supplementary Table 5). Further analysis suggested that a total of 138 MNP loci have different genotypes between at least one pair of lines, and many of the these MNP locus located at protein coding genes, such as disease resistance protein RGA4 gene, F-box protein At4g22030 gene and ethylene-responsive transcription factor ERF018 gene. This large range of genetic heterogeneity among the homonyms raised strong concern regarding the reproducibility of the research results using Nipponbare.

The more accurate a DNA marker method is, the more subtle genetic variations it can detect. Four pairs of Nipponbare line, which had the smallest genetic variation of only one different genotype, were chosen for validation of their distinct genotypes by Sanger sequencing. All (100%) of them were successfully validated (Fig. 2C, D and Supplementary Fig. 1), confirming the observed genetic heterogeneities in the 30 rice lines and further demonstrating the accuracy of MNP-Seq.

#### Accurate and efficient determination of plant variety identity

Registration and protection of the plant variety intellectual property rights required to accurately and efficiently determine the identities of plant varieties. Comparing SSR fingerprints from different experiments may be efficient but is not accurate to determine plant variety identities because of genotyping error. In this study, we constructed an MNP fingerprint database using the 47 rice varieties of the first technical triplets plus 150 other widely-used hybrid rice varieties (Supplementary Table 6). The second technical triplets of the 47 varieties were taken as blind samples and their identities were determined to be the ones in the database based on MNP fingerprint differences. As a result, the identities of all 47 blind varieties were correctly determined (Supplementary Table 7), indicating a 100% accuracy for the identity determination by MNP fingerprints across experiments.

In another related application, this scheme of variety-identity determination using MNP fingerprints was adopted to investigate the possible infringements of 29 rice varieties that were directly from the seed market but had different labeled names. Variety pairs that had MNP fingerprint differences less than 1% were deemed to include infringed varieties. As a result, 3 pairs were found to be suspected from the 406 pairs of the 29 varieties. These three pairs were all validated to include infringed varieties since they had greater than 1% different MNP fingerprints from their standard varieties (Supplementary Table 8), showing the ratio of infringement to be 3/29 = 10.34%.

Combined, these results suggested MNP-Seq be an effective and enabling technique for accurately detecting infringing plant varieties and for protecting the intellectual property rights of plant varieties. Indeed, the new MNP-Seq method has been adopted as the Chinese national technical standard for plant identity determination^28^.

## Discussion

The novel MNP markers and its genotyping method provide significant improvement over existing DNA marker technology. First, existing amplicon-based approaches (e.g., cancer gene, 16S rRNA, plant and animal sequencing) often focus on known housekeeping genes or disease-causing genes, and hardly used to identify the identity of plant variety due to low polymorphism or low discrimination of these targeted genes. In contrast, each MNP markers include several SNPs and thus have at most 2*^k^* distinguish alleles, which mean high polymorphism and high discriminative for each MNPs in the analysis of plant variety. The developments of MNP were based on the whole genomic SNPs of several variety or accessions, thus a large number of polymorphic loci can be obtained simply and quickly. To detect variants from amplicon sequencing data, most genotyping methods employ a probabilistic framework and genotyped SNPs one by one. Thus the confidence of genotyping was often confounded by numerous factors such as extreme read depth, homopolymer and population samples. In contrast, the genotyping of each MNP locus were conducted based on the whole reads pair (one pair by another), and errors randomly occurred could be excluded easily. Finally, the unique genotyping method of MNP-seq makes it more suitable for population (e.g., plant varieties), polyploidy, and hybrid samples compared with other approaches.

### The features of MNP marker and MNP-Seq

A high degree of polymorphism can reduce the number of markers needed and subsequently genotyping error. High genotyping efficiency can help rapidly genotype a large number of markers to decrease sampling bias. The MNP marker gained high polymorphisms by using multiple *k* SNPs within a short DNA segment to capture 2*^k^* possible alleles. In this study, the polymorphism of the MNP marker for rice was shown to be greater than that of the SSR marker, which has been widely applied due to its high polymorphisms.

The efficiency of MNP-Seq was attributed to multiplex amplification, high-throughput sequencing, and bioinformatics analysis. The multiplex amplification in the first step had a high efficiency to enrich thousands of marker loci by a single PCR reaction^12,29^. When combined with deep sequencing and bioinformatics analysis using the efficient MNP-Seq software, we were able to efficiently genotype more than 1,000 MNP markers for a collection of rice and cotton in about two days.

Several factors contributed to the accuracy of MNP-Seq (see Method). Marker design and selection avoided error-prone sequences, such as successive SNPs and simple repeats; the sequencing-based detection of MNP markers overcame the uncertainty inherent in the SSR amplicon length displayed on a gel and the microarray hybridization noise for SNP markers; the amplicons of MNP markers were often sequenced thousands of times (Supplementary Table 2) so that random experiment errors were minimized; the chance that two different MNP genotypes were mistaken as identical dramatically decreased to approximately the level that all of the SNPs within the MNP were incorrectly genotyped. As a result, the accuracy of MNP-Seq reached 99.988% for polyploidy hybrid varieties, which were popular in crops but rather difficult to accurately genotype. Table 2 summarizes the advantages of our novel MNP-Seq method over the major existing methods for genotyping DNA markers.

These imminent features made MNP-Seq the method of choice for fast, accurate and genome-scale genotyping, which was compared favorably over the other existing methods. Indeed, MNP-Seq has been adopted as the Chinese national technical standard of genotyping for plant identity determination^28^.

### The applications of the MNP marker and MNP-Seq method

Combined, the MNP marker and MNP-Seq genotyping method offered the *first* practical and genome-scale genotyping approach that is potentially *the method of choice* for many applications. We demonstrated the usage and power of MNP-Seq using two important but challenging problems as case studies in the current paper, which would be rather difficult or even impossible without the MNP marker and MNP-Seq.

Homonyms are biological materials for research that were labeled by the same name but genetically heterozygous. Homonyms may cause serious problems regarding the reproducibility and consistency of research results from multiple labs and the results overall a period. Homonyms and their problem of result reproducibility have been reported for Hela cells, which were widely used in biological and cancer research^30-32^. In the current study, we also revealed homonyms for Nipponbare, a widely adopted model organism in plant biology.

The genetic variations among biological homonyms are usually subtle due to low mutation rates, i.e., typically 10^−9^ per generation in human^33^ and thus were often shadowed by technical errors of the existing DNA marker profiling methods, e.g., 5% error rate for SSR^11,12^ and 1% to 6.1% error rates for SNP marker^20-22^. MNP-Seq greatly reduced the average number of erroneous genotypes to no more than 0.113 per sample (Table 1), suggesting that any distinct genotypes discovered by MNP-Seq were highly likely real genetic variations. Indeed, even the smallest genetic variation discovered by MNP-Seq, i.e. the only one different genotype between two Nipponbare homonyms, was validated to be true by the Sanger sequencing (Figure 2C, 2D and Supplementary Fig. 1).

The accuracy of the identity determination of a plant variety is a core problem for registration and protection of the plant variety intellectual property rights. The efficiency of the identity determination is also important to handle many varieties whose rights have to be protected. Although comparing SSR fingerprints is popular for plant identity determination, a substantial amount of parallel experiments is needed to compare the variety under the test with the varieties to be protected, e.g., over 6,000 registered maize varieties in China^34^, to determine if the former is an essentially derived variety (EDV) or an infringed variety of one of the latter. In this process, the heavy workload of parallel experiments defeats the SSR fingerprint method. Since the reproducibility of the MNP marker was over 99.974% (Table 1), comparing two MNP fingerprints from different experiments can have high accuracy, as shown to be 100% for the two cases of determining the blind sample identity and screening some rice varieties from the seed market. This strategy of direct comparison of MNP fingerprints is also highly efficient by avoiding laborious parallel experiments. For example, to determine the identities of the 47 blind varieties from the 197 to be protected, the workload using the MNP fingerprinting is only 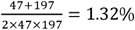of that using SSR fingerprinting.

As a final note, we like to highlight that while we focused on difficult problems in plant biology, the MNP marker and MNP-Seq can be directly applied to many similar problems in animals and micro-organisms.

## Materials and methods

### Plant materials

The study used plant materials of 43 tetraploid hybrid cotton varieties, 197 hybrid or selfing diploid rice varieties, 29 rice varieties collected from the market in China (Supplementary Table 6), and 30 ‘Nipponbare’ rice lines from different labs (Supplementary Table 4). All these materials were collected in this study. Genomic DNA of each variety or line was extracted from a mixture of equal-amount young leaves of 30 individual plants, following the protocol of E-Z 96 ® Mag-Bind® Plant DNA Kit (Cat. No. M1027, Omega Bio-tek, USA).

### Design of MNP marker and multiplex PCR primer

The public SNP data of 50 rice varieties from^35^ and 318 cotton varieties from^19^ were used to design MNP markers for rice and cotton, respectively. A sliding window of 125 base pairs (bp) was used to scan the genome with an increment of 5 bp. The discriminative power (DP) of a window was defined as *t* / *c*(*N*,2), where *c*(*N*,2) was the number of variety pairs among *N* varieties used and *t* the number of the pairs each of which had at least two dispersed SNPs within the window. The windows with DP ≥ 0.2 were chosen for multiplex PCR primer design and synthesis at Molbreeding Biotechnology Co., China. About 1,000 windows without simple sequence repeat and successive SNPs were randomly chosen from those with adequate multiplex PCR primers and the SNPs within these windows were used as MNP markers (Supplementary Table 9). The length of all primers ranged from 18 bp to 39 bp. No further optimization were conducted for PCR conditions after the primer designed and an MNP locus will be ignored if no products generated in the multiplex PCR amplification.

### Library construction and high-throughput sequencing

Sequencing libraries were constructed from 10 ng genomic DNA following the protocol of MGIEasy Library Prep Kit (Cat. No. 1000006985) with some modification to adopt BGI-seq high-throughput sequencing platform as done in^12^. The PCR cycle for library construction was set below that of the nonlinear amplification, i.e., less than 17 cycles. To allow for multiplexing during sequencing, ten □ nucleotide barcode sequences specific for each sample were appended to all libraries. The libraries were sequenced on MGISEQ-2000RS platform (BGI, Shenzhen, China) using 150 bp paired-end sequencing at our lab. To test the reproducibility, 47 rice and 43 cotton varieties were randomly selected and the library construction and sequencing were repeated for three times (technical triplets) with at least 30-day intervals between any two repeats.

### MNP Genotyping, fingerprinting and comparison

Sequencing reads without ambiguous bases were retained and mapped to the reference genome (IRGSP-1.0 for rice or *Gossypium hirsutum* acc. TM-1 for cotton) with Bowtie2 (version 2.1.0) using the default parameters. The unmapped reads were further mapped with BWA (version 0.7.9a) using the default parameters. An MNP allele was defined to be a haplotype of the SNPs in the MNP locus. The allele with at least 20 mapped reads and having the highest read count was defined as the major allele of the MNP locus. For clarity, define the α value of an MNP allele to be the ratio between its number of supporting reads and that of the major allele for the locus. For a locus having a major allele, the candidate alleles were the ones with over 2 supporting reads and their α values greater than 0.2. A candidate allele was called a true allele when its α value was greater than 0.5. The genotype of an MNP locus was the set of all its candidate alleles. The MNP fingerprint of a sample was the collection of all its MNP genotypes.

When comparing two genotypes of an MNP locus in two samples, if all the true alleles of either genotype were the same or a subset of the other one, the two genotypes were identical, or different, otherwise. When comparing the fingerprints of two samples, the difference of their fingerprints was defined as the ratio between the numbers of the different genotypes and all of the genotypes that the two samples shared in common.

### Estimation of the reproducibility and accuracy of MNP-Seq

The MNP profiling results from the technical triplets of the 47 rice and 43 cotton varieties were used to estimate the reproducibility of MNP-Seq. Here the reproducibility was estimated by comparing the genotypes of each MNP locus between each two replicates separately. A reproducible MNP genotype was observed if identical genotypes were observed between two replicate, or irreproducible, otherwise. The reproducibility of the MNP-Seq method was then estimated as *r* = *n*/*N*, where *n* and *N* were, respectively, the numbers of the reproducible and all genotype pairs compared. The chance that two reproducible genotypes were incorrect but identical should be rather small. Thus, both reproducible genotypes were taken as correct, or one of them as incorrect, otherwise. Therefore, the accuracy of MNP-Seq was estimated as 1-(1-*r*)/2 = 0.5 + 0.5*r*.

### SSR genotyping and validation of MNP genotypes

The genotypes of 3,105 SSRs in 8 rice samples have been reported in^12^. The polymorphism of each SSRs was calculated based on the reported genotyping results and the top 412 most polymorphic SSRs were used in this study (Supplementary Table 9). These SSR markers were genotyped for 39 of the 47 rice varieties (Supplementary Table 6) following the method in^12^.

The validation of MNP genotypes began with PCR amplification of the chosen MNP loci using the same primers used before (Supplementary Table 1) and following the protocol of AmpliTaq Gold® 360 Master Mix (4398876, Applied Biosystems™, USA). The PCR products were then sequenced by Sanger sequencing (TsingKe Company, China).

## Data Availability

All the genomic sequence data sets are available in the NCBI Sequence Read Archive under accessions PRJNA622759.

## Acknowledgements

This work was financially supported by Application foundation Project of Wuhan Science and Technology Bureau (2018020401011298), National Key R&D Program of China (2016YFF0202303,2018YFD1000500), National Natural Science Foundation of China (31800306, 31830069), Project of Hubei Intellectual Property Bureau (2019-23), Open Research Fund of State Key Laboratory of Hybrid Rice (Hunan Hybrid Rice Research Center)(2018KF02).

## Author Contributions

H.P. initiated the research and coordinated the project. W.Z. supervised the study. H.P., W.Z., Z.F. and A.Y. wrote the manuscript. T.L., A.Y., H.C, N.X., and R.H. collected plant materials and extracted high-quality DNA. J.Z., J.Z., and L.G. constructed DNA-sequencing libraries and carried out high throughput sequencing. Z.F. and L.L. performed bioinformatics analyses. X.F., A.M., J.Z. and L.C. evaluated some of the results. Y.C. and H.X. participated in the analysis.

## Figure Legends

**Supplementary Figure 1. Validation of distinct MNP genotypes between Nipponbare lines.** The numbers on right are the codes of the Nipponbare lines. The upper and the lower parts for each MNP locus are the genotypes from MNP-Seq and Sanger sequencing, respectively. Variants bases are indicated by arrows. **(A)** Locus AMPL1563256. **(B)** Locus AMPL1563256. **(C)** Locus AMPL1563757.

**Supplementary Table 1. Supplementary Table 1. Multiplex PCR primers used for MNP genotyping rice and cotton varieties.**

**Supplementary Table 2. Statistics of sequencing data for rice and contton varieties.**

**Supplementary Table 3. The allele number for each of SSR and MNP loci.**

**Supplementary Table 4. Nipponbare rice lines used in this study.**

**Supplementary Table 5. Distinct MNP genotypes for each pair of Nipponbare lines.**

**Supplementary Table 6. Rice and cotton varieties used in this study.**

**Supplementary Table 7. Identity determination of blind samples.**

**Supplementary Table 8. Infringed varieties.**

**Supplementary Table 9. Multiplex PCR primers used for SSR genotyping in rice varieties.**

## Reference

1. Wang, W. et al. Genomic variation in 3,010 diverse accessions of Asian cultivated rice. Nature 557, 43–49 (2018).

2. Zhou, Z. et al. Resequencing 302 wild and cultivated accessions identifies genes related to domestication and improvement in soybean. Nat Biotechnol 33, 408–14 (2015).

3. Xie, W. et al. Breeding signatures of rice improvement revealed by a genomic variation map from a large germplasm collection. Proc Natl Acad Sci U S A 112, E5411–9 (2015).

4. Chen, H. et al. A high-density SNP genotyping array for rice biology and molecular breeding. Mol Plant 7, 541–53 (2014).

5. Stoffel, M.A. et al. Demographic histories and genetic diversity across pinnipeds are shaped by human exploitation, ecology and life-history. Nat Commun 9, 4836 (2018).

6. Cunha, J.T. et al. RAPD and SCAR markers as potential tools for detection of milk origin in dairy products: Adulterant sheep breeds in Serra da Estrela cheese production. Food Chem 211, 631–6 (2016).

7. Alqahtani, A. et al. Differentiation of Three Centella Species in Australia as Inferred from Morphological Characteristics, ISSR Molecular Fingerprinting and Phytochemical Composition. Front Plant Sci 8, 1980 (2017).

8. Asadzadeh, M. et al. Simple, Low-Cost Detection of Candida parapsilosis Complex Isolates and Molecular Fingerprinting of Candida orthopsilosis Strains in Kuwait by ITS Region Sequencing and Amplified Fragment Length Polymorphism Analysis. Plos One 10, e0142880 (2015).

9. Ellegren, H. Microsatellites: simple sequences with complex evolution. Nat Rev Genet 5, 435–45 (2004).

10. Hosseinzadeh-Colagar, A., Haghighatnia, M.J., Amiri, Z., Mohadjerani, M. & Tafrihi, M. Microsatellite (SSR) amplification by PCR usually led to polymorphic bands: Evidence which shows replication slippage occurs in extend or nascent DNA strands. Mol Biol Res Commun 5, 167–174 (2016).

11. Treangen, T.J. & Salzberg, S.L. Repetitive DNA and next-generation sequencing: computational challenges and solutions. Nat Rev Genet 13, 36–46 (2011).

12. Li, L. et al. An accurate and efficient method for large-scale SSR genotyping and applications. Nucleic Acids Research 45, e88–e88 (2017).

13. Pourabed, E. et al. Identification and DUS Testing of Rice Varieties through Microsatellite Markers. Int J Plant Genomics 2015, 965073 (2015).

14. Choi, S.P. et al. Genetic characterisation of commercial Chinese cabbage varieties using SSR markers. Seed Science and Technology 44, 595–608 (2016).

15. Delfini, J. et al. Distinctness of Brazilian common bean cultivars with carioca and black grain by means of morphoagronomic and molecular descriptors. Plos One 12, e0188798 (2017).

16. Gettings, K.B., Aponte, R.A., Vallone, P.M. & Butler, J.M. STR allele sequence variation: Current knowledge and future issues. Forensic Sci Int Genet 18, 118–30 (2015).

17. Daunay, A. et al. Low temperature isothermal amplification of microsatellites drastically reduces stutter artifact formation and improves microsatellite instability detection in cancer. Nucleic Acids Res 47, e141 (2019).

18. Huang, X. et al. Genome-wide association studies of 14 agronomic traits in rice landraces. Nat Genet 42, 961–7 (2010).

19. Fang, L. et al. Genomic analyses in cotton identify signatures of selection and loci associated with fiber quality and yield traits. Nat Genet 49, 1089–1098 (2017).

20. Li, G. A new model calling procedure for Illumina BeadArray data. BMC Genet 17, 90 (2016).

21. Bayer, M.M. et al. Development and Evaluation of a Barley 50k iSelect SNP Array. Front Plant Sci 8, 1792 (2017).

22. Sasaki, S., Yoshinari, K., Uchiyama, K., Takeda, M. & Kojima, T. Relationship between call rate per individual and genotyping accuracy of bovine single-nucleotide polymorphism array using deoxyribonucleic acid of various qualities. Animal Science Journal 89, 1533–1539 (2018).

23. Allen, A.M. et al. Characterization of a Wheat Breeders’ Array suitable for high-throughput SNP genotyping of global accessions of hexaploid bread wheat (Triticum aestivum). Plant Biotechnology Journal 15, 390–401 (2017).

24. Malmberg, M.M. et al. Evaluation and Recommendations for Routine Genotyping Using Skim Whole Genome Re-sequencing in Canola. Front Plant Sci 9, 1809 (2018).

25. Stahlberg, A. et al. Simple multiplexed PCR-based barcoding of DNA for ultrasensitive mutation detection by next-generation sequencing. Nat Protoc 12, 664–682 (2017).

26. Jee, J. et al. Rates and mechanisms of bacterial mutagenesis from maximum-depth sequencing. Nature 534, 693–6 (2016).

27. Moghe, G.D. & Shiu, S.H. The causes and molecular consequences of polyploidy in flowering plants. Ann N Y Acad Sci 1320, 16–34 (2014).

28. Identification of plant varieties - MNP marker method. (state administration for market regulation, China, 2020).

29. Pater, J.A. et al. A common variant in CLDN14 causes precipitous, prelingual sensorineural hearing loss in multiple families due to founder effect. Hum Genet 136, 107–118 (2017).

30. Liu, Y. et al. Multi-omic measurements of heterogeneity in HeLa cells across laboratories. Nat Biotechnol 37, 314–322 (2019).

31. Yu, M. et al. A resource for cell line authentication, annotation and quality control. Nature 520, 307–11 (2015).

32. Ben-David, U. et al. Genetic and transcriptional evolution alters cancer cell line drug response. Nature 560, 325–330 (2018).

33. Willems, T., Gymrek, M., Poznik, G.D., Tyler-Smith, C. & Erlich, Y. Population-Scale Sequencing Data Enable Precise Estimates of Y-STR Mutation Rates. Am J Hum Genet 98, 919–933 (2016).

34. Tian, H.L. et al. Development of maizeSNP3072, a high-throughput compatible SNP array, for DNA fingerprinting identification of Chinese maize varieties. Mol Breed 35, 136 (2015).

35. Xu, X. et al. Resequencing 50 accessions of cultivated and wild rice yields markers for identifying agronomically important genes. Nat Biotechnol 30, 105–11 (2011).

