## Supplementary table 1 for "Multiple nucleotide polymorphism DNA markers for the accurate evaluation of genetic variations"

**Supplementary Table 1. Multiplex PCR primers used for MNP genotyping rice and cotton varieties**

| Locus code | Forward primer | Reverse primer | Species |
|---|---|---|---|
| AMPL1563062 | CCAGATTCACCTCCTCTCCTCAA | GATGGACGTCAAGATTCGTGCTAT | *Oryza sativa L.* |
| AMPL1563095 | GGATCCCTTAACTTACAATACCGGTT | GTCGCTCGTTTGCACTTGAG | *Oryza sativa L.* |
| AMPL1563096 | ATGAGGTTAATAATCTGGACATGGCAA | GGAGGTAGTAGTAAGCAATCCCATTG | *Oryza sativa L.* |
| AMPL1563097 | TTCTTATGGCATCACGATTTAGAGGTAAC | ACAACTGAATGAGACTTATGCTACCAAG | *Oryza sativa L.* |
| AMPL1563100 | TCATCTAGTCTGAATCCAGTTCCACT | AACTTCTTGTTACTGAGGCTTCTAGATCTA | *Oryza sativa L.* |
| AMPL1563102 | AATTTTAGATTTAACTGCGATGCATTCGA | CCCGTTTCGATAATCGAATTTCATTACG | *Oryza sativa L.* |
| AMPL1563103 | TGTGAGATGTACTCCGTATTACTAATTACTT | AATGGATAGGTCATTAACCTAGTATTTGATTCA | *Oryza sativa L.* |
| AMPL1563116 | CAAGTATTTCTGGTATGTATCTGGACAAGA | CAACAACCATGCATGCAATGGAT | *Oryza sativa L.* |
| AMPL1563121 | CATCGGAGTTTGTGAAAATAGAGAATAACC | GAGTATCAATTTTGAGCATGTAGCTAGACT | *Oryza sativa L.* |
| AMPL1563123 | GACAAGGTGGCCTATAAATAAGCAGA | CCTCTCACAAACCATGGTATGCAT | *Oryza sativa L.* |
| AMPL1563126 | CGCCACTACCTCGCTTGT | GACTATTCATCTTTCTGAAAAATGTTTTCACAA | *Oryza sativa L.* |
| AMPL1563133 | TATAAGTTGAGGTGTTGTTGAGTGCA | ACTAAACGGAATCTTTTTAATAGCTCACCA | *Oryza sativa L.* |
| AMPL1563134 | GGTCACGCCTTCTCCACTTG | CGAAGACTCACCACCATGAGT | *Oryza sativa L.* |
| AMPL1563137 | GCTTGCTGAGCTGTTGTTTCAT | GCGTTCCAGGAGCACTAAGATG | *Oryza sativa L.* |
| AMPL1563138 | CCGTGGAACTGGGACAACAATT | GCCAGACACGTCAGAGATTCAAC | *Oryza sativa L.* |
| AMPL1563143 | TTTTTGCCCATCTGAAACAAAATGTATTTT | GGGCTAGGGTTTGGGAATAAATCAG | *Oryza sativa L.* |
| AMPL1563147 | GATTATTCGGTCACTTGTTAAGGGAGT | GTTTTTCTTCTACTGCCACATAAGCATC | *Oryza sativa L.* |
| AMPL1563153 | TCTAGAAAACTCAATACGGAGTATTGCC | AAAGGTGTTCTAAATTTTGTTTAAAGAGCTACA | *Oryza sativa L.* |
| AMPL1563154 | AGATTGATCTGGACAAAATGGCTAATTAGT | GCTATTTACGCCAAGTTTAGTTCCAAAA | *Oryza sativa L.* |
| AMPL1563157 | GCGCTTGAGACAGTGGAGATTC | GTGGTTTCGTTGATCACCAAAGTG | *Oryza sativa L.* |
| AMPL1563158 | CTTAATTTCATCCTTGGGCTTATACTCAGA | GCACATACACATCACTGTACACGT | *Oryza sativa L.* |
| AMPL1563159 | GCTCGGCAACGACTACGT | ACTTATGTTGCGTGCGAAGACA | *Oryza sativa L.* |
| AMPL1563161 | ATATGCCGCTTGCTTGACTCT | GCATGATGCTCAGCATCCCTT | *Oryza sativa L.* |
| AMPL1563166 | GTATACGCTACGTAGAACAAAACCG | TCGATTCAAAGTACCCTGCCAAAA | *Oryza sativa L.* |
| AMPL1563168 | GTCTTAAATTAGGCAGCCGTACGAT | CTGGTGTTTAGTGTGGCTTCAAG | *Oryza sativa L.* |
| AMPL1563169 | CAGTGGGAAGCCACATAAACCA | GGGACATGTTCTACCTCATGATCATG | *Oryza sativa L.* |
| AMPL1563172 | TTCATGCTAACCACGTGCACTA | TTTTCCCGGTTTTATTTTTCTATAGGGACA | *Oryza sativa L.* |
| AMPL1563173 | TGGAGCAAATAATCAATTTCAAAGTTCAGC | TGTTAGGCTGTAGTAATTCACATGATGTG | *Oryza sativa L.* |
| AMPL1563175 | AGTGCCTTTCGTATTTCTCAATGTGT | CTTGATTTGGCTTTTGCCTTCTCT | *Oryza sativa L.* |
| AMPL1563176 | CCCACACAACTACGACTGTGTT | ACAGTTTATGCGTCCTTGCACATA | *Oryza sativa L.* |
| AMPL1563178 | ATGATACGAGTAAATATTATTAGGCCGTGTTTT | GACAACAGAAATTGCCGGTGT | *Oryza sativa L.* |
| AMPL1563182 | AACGAAAATGTGCATCTGTTTAGTTTCTT | GCTAAAATGAGGAATGAGGCCAAGA | *Oryza sativa L.* |
| AMPL1563184 | GAATGGGTAAATTGGAATCATGCCA | GACATTACGACGAGTAATGAACAGGTAAA | *Oryza sativa L.* |
| AMPL1563186 | GATGGCTAGTGTTTGGTGCAAAG | GCGGACAACTTGTGTAACCCTT | *Oryza sativa L.* |
| AMPL1563187 | GGCTCCTATTTGCCGCTGATTA | TCTAGCTCGGCGAATTAAGAATACAAG | *Oryza sativa L.* |
| AMPL1563189 | TTCGGTTTATAACATACTAAGTCGTGCAT | CGTGTAGTCCCTCTCTGAAACG | *Oryza sativa L.* |
| AMPL1563193 | CACTCCAAACAGTAGTGAGTAGTGT | GGGCCTGTTTATCCTAATTTTGTTTTCTT | *Oryza sativa L.* |
| AMPL1563194 | CGAACCCGTTCTCTCAATTTATATAGGA | GCAAGTCCTATTTGGAGAATTAATTCGAGA | *Oryza sativa L.* |
| AMPL1563196 | TTTTTGGGTTAATTTTAGCATCACCAGA | TACCGTGTTAAAATATTGGCAAGGTCTAAA | *Oryza sativa L.* |
| AMPL1563200 | CCAATTTATACACACACTACGTCTTGGA | AAATTCTAGATGCACTCACAGCATTACT | *Oryza sativa L.* |
| AMPL1563203 | GGTCATGTGAATATAGCCGGTGAT | CTGCTAATCACATAGAAAACATACCAAACC | *Oryza sativa L.* |
| AMPL1563205 | AAGTCGTCCCTCCTGAATTGC | GGCTAACTCCAAGGCCCATA | *Oryza sativa L.* |
| AMPL1563212 | TGGTTTTCACGGGCTTTTCTAGA | CGATCCTACACACATAGCACAGAT | *Oryza sativa L.* |
| AMPL1563213 | TGTGTCGGTTAGACCGTCTAGT | TGGCCCAATCTTTTAGTGAGTTAAGATAAC | *Oryza sativa L.* |
| AMPL1563214 | GTTTTTGTTCTTTCAGTTCCAACCCAA | GTCTCATCCATCTCCATGAAACCTC | *Oryza sativa L.* |
| AMPL1563215 | TCTTAATCTTTTGAAAATTTGATGCCGCT | ATCCATGTACGTAACTCCAAGCC | *Oryza sativa L.* |
| AMPL1563218 | CCCACTTGACAAAATTGAGCGA | TGTTACTAGCATGTTTAGGGAAAGTGTC | *Oryza sativa L.* |
| AMPL1563222 | GGATTAAGCGCGAAGATCCGAA | TGCACTATCACTTTGGTAAAAAGGCA | *Oryza sativa L.* |
| AMPL1563223 | CGAGGCAATATCGCCATCTCTT | AAAGACAACTATGTGCCACCAAGA | *Oryza sativa L.* |
| AMPL1563234 | TTGCATGTTATATTTAAGGCATCCAGGA | GATACTGTTCCGGCTGGCA | *Oryza sativa L.* |
| AMPL1563238 | GCCCATAATGGGTTACCTGTATCC | GGCTAGGGTCAATCCACTTATAAATTGAAT | *Oryza sativa L.* |
| AMPL1563244 | GTCCATGTGGTGTGCATGTAG | AGAGGTGGCATTAGAAATCCATAGCTATA | *Oryza sativa L.* |
| AMPL1563256 | CAGACTTTTCCTCCTGATTGGATGA | GATTCAGCAGCAATTTTGAAGAAATGTG | *Oryza sativa L.* |
| AMPL1563261 | AGAGGAAGGAAAAATCACACGGTT | ACCCATCCTCCGTCCATATGA | *Oryza sativa L.* |
| AMPL1563263 | CCTCAGTACGTCTATCACTCCCTT | TGCTCTTAGGATCTACTTATTGTCTCCTAC | *Oryza sativa L.* |
| AMPL1563267 | TAACGTTGTTGCGATAACAACTATTGC | CAAGAGCACACACTACAAGGAAATC | *Oryza sativa L.* |
| AMPL1563268 | GATTGGTTACTCGTAGACTAGAGATAGGAG | CGGGTGTATTATTGATCTCACAATCTTAGT | *Oryza sativa L.* |
| AMPL1563272 | AAGGAAAGGATCCTTTTGAGTTAAGGAAAT | TCGATGAGATCTATGGGCGAATTG | *Oryza sativa L.* |
| AMPL1563275 | CTATAAAATCATGCTACACCATTAATATCCGG | TTTTACGTTAAATAAGCTTAAAGCCCAAAC | *Oryza sativa L.* |
| AMPL1563276 | TGGGCCAAAATCGACTAATACATATGTT | GATGTATGTACCACGATCCATTTCTTACT | *Oryza sativa L.* |
| AMPL1563279 | TCAATCTCAATCAATCACCATTTACCAAGT | GTGATGCTTGAAGTCATTAGGAGTCA | *Oryza sativa L.* |
| AMPL1563282 | TACACCAACATTCTCTTCTCAACTAATCAG | CTGTGCGGTGCCAAAACAATAA | *Oryza sativa L.* |
| AMPL1563285 | GCGTGGTGTTCGACGAAAC | CTTCCTTGCCTTCGCTGTCTTA | *Oryza sativa L.* |
| AMPL1563286 | ATGAGCAAGAGCTCATTGTGGT | AACGTAGATGAGGACCTGAGGAA | *Oryza sativa L.* |
| AMPL1563289 | TCATGACAACCACATATCTCCAACTTC | TCTAGGATGAATCTGTTTCCTCGTCT | *Oryza sativa L.* |
| AMPL1563296 | TTAATTACATCGATCAATACGTGTTGCCTA | TATGCTCTGTCACTTCGTATTATGCG | *Oryza sativa L.* |
| AMPL1563298 | GATAGGGTTCGGAAACTAATTCTCCA | CCATAAGTCTCGTGGTGTTTGCT | *Oryza sativa L.* |
| AMPL1563299 | CATCAGTTTCTGCTGTCCTGGTA | TGGTTGTAGGACATCCGGGTATT | *Oryza sativa L.* |
| AMPL1563304 | CCAGGGAATGTTTGGACATGGT | TATGTGCAGGAGGCCAACAAGGT | *Oryza sativa L.* |
| AMPL1563308 | TCAGGGATTTCTTTGTTGGAAAGGA | TGTATGGTAAGCTCAATATCTCTCGATTCT | *Oryza sativa L.* |
| AMPL1563310 | GAAAATCCAAAACTTCATCCGGTATTTCT | AAATATGGCTATCTTTAGTCCCACGTAAAA | *Oryza sativa L.* |
| AMPL1563311 | AGTTGGATTTTTATAGCTCTCGTAAACAGG | CGTTAGGTTGAAGTGTTCATTCAATGAA | *Oryza sativa L.* |

| | | | |
|---|---|---|---|
| AMPL1563315 | AGTGTTCCAACCCTGTCTCCTTA | CATTGCAGTCATGCAAATTTCCCA | *Oryza sativa L.* |
| AMPL1563317 | GTATAATTCATGGCTCGATCGAATCGTA | GTCTTCGGATGTTTGGGAGGTT | *Oryza sativa L.* |
| AMPL1563318 | GTGCCCAGCAACCCAGTTAAAA | CAACGAAGTGCCTCGAGAGAAA | *Oryza sativa L.* |
| AMPL1563326 | GCCAGCAAGTAATAACTCTGAGCT | CAACAACAAGCTCAATTCCTTGCTT | *Oryza sativa L.* |
| AMPL1563332 | AGAACTGGGCAAAGTCGATTTGT | CTTGATGAGATGCTTTCCATGCTTG | *Oryza sativa L.* |
| AMPL1563333 | TGTTCTATCCCTATAGCATAGGGAACTTG | TAAGAGCAGCTAGCGATAGATCTGTAG | *Oryza sativa L.* |
| AMPL1563334 | ATTTTGTGGCACCATCGAATATGC | CTGGGAGGACCAAAAGATACAACTAG | *Oryza sativa L.* |
| AMPL1563337 | GTACGAGGAACATATTTTGTTCAAACAGTT | CTACCTGAGCGACAGAAAGAACA | *Oryza sativa L.* |
| AMPL1563345 | GGCGTTTAAATTTATTGAGCATGTAGGT | GTATTGTGATTTTCAGTGCTAGTTTGTGAT | *Oryza sativa L.* |
| AMPL1563353 | TTTTAGCAGAACCGAGCTGTCAA | TTCCAATGTTGACATCCTCTTGCT | *Oryza sativa L.* |
| AMPL1563357 | GGATTTTGCCGGAAAACTCTTCTAAAAA | CAATGGAATCCCTAATCTATATGCACATCT | *Oryza sativa L.* |
| AMPL1563358 | ATGTATAATGTCCTGTGATTATATGTGGCA | GGTCGCAAAATGCGAAGAACAA | *Oryza sativa L.* |
| AMPL1563362 | AATTCTGCAGCTTTAATGCTACTAGCTTA | GTCAAACAGTTTGAACTCAGAAATACCAG | *Oryza sativa L.* |
| AMPL1563365 | TGATAATATGCATATGGCATACTGTGGA | CCACTATCGCCAATACACTTCTCAAC | *Oryza sativa L.* |
| AMPL1563366 | GCTACCAGCATTCAGCCATTCA | GATCCAAACAAGGCCATTGACAG | *Oryza sativa L.* |
| AMPL1563369 | TAAATCAAGCTTTGCAGCAGCTAAAG | CCAATCTCACTTGCACGAGGTT | *Oryza sativa L.* |
| AMPL1563373 | TGCTGCAAGGTTTGTTTGATTCTG | TCCGCTACAATTACAGGATCCAATG | *Oryza sativa L.* |
| AMPL1563375 | GTGCTGTGTGATTCATTATGATTCTCC | CTTGCGCACGTACTACTCATTAGA | *Oryza sativa L.* |
| AMPL1563376 | GACTTGGTGGCCACAACATCTA | GACTTCTGCTTCCTTTTGGATTCG | *Oryza sativa L.* |
| AMPL1563377 | GGGATCTCAATTGCTTTTGCAGA | GCTGCCTTTTGACAGCAAGATC | *Oryza sativa L.* |
| AMPL1563381 | CCCAAGGCTAAAGCTCGTTGTA | GCACTCGACGAGGATAAGGTAG | *Oryza sativa L.* |
| AMPL1563385 | CAACACATAAGCTGTGTGCACTT | CCCTACATTCATCCATGTATTAAAACATCC | *Oryza sativa L.* |
| AMPL1563387 | CCACATTTGCTCGGTTTTCGTAAG | CGAAATCTTGTTTTGACAGTGGACAAG | *Oryza sativa L.* |
| AMPL1563389 | TCAAGAGGAAAATAAGCATGAGGCAA | TCAGGTCAGTAGCTATAGTATTCAAATCGT | *Oryza sativa L.* |
| AMPL1563390 | GGAGCATAGAAGCACTTCTCACAAA | AAGACACATTTTGGTATCAATCCAAATTCC | *Oryza sativa L.* |
| AMPL1563392 | AAGTTCGTTCGCTTTTGAAATTGAAGA | GTCTGACTTTCATCTCAAGTTAACATTCG | *Oryza sativa L.* |
| AMPL1563396 | GACCTTCTGTTTCTACCAGAAAAACAC | GCTCTCTTCCTTCTGTTCAATGGATG | *Oryza sativa L.* |
| AMPL1563397 | CGATTGTTTTCGGTATGTCAGTTGTG | TGATGCCAGTGAACATTTGGGT | *Oryza sativa L.* |
| AMPL1563398 | GAAATCAACCTGATTTCCAAATAGTTCAGT | CGCACGAAAATAATATAGCTCATTAACACA | *Oryza sativa L.* |
| AMPL1563399 | TTTCCATGTTAAGGGATCAAATGACCA | CCATCCTACTCATTCGAACACCA | *Oryza sativa L.* |
| AMPL1563409 | GTAAGTGACTTATCTACACATACGTGCA | TATTCCACCCAAGGAGTGCTAATTG | *Oryza sativa L.* |
| AMPL1563410 | TGTGATGATCAGTCCTAGAGGTGA | ATTTAGGAGGAAACTGCATCTATTCATGTT | *Oryza sativa L.* |
| AMPL1563412 | TGTTTGGACTTATATGACACCCGAAA | TGATGGCTTGAAGAATATAAAGCACTAGT | *Oryza sativa L.* |
| AMPL1563418 | CCGGTGGGCCTTTTGTCTTTAT | CATTATACTAAAACAGAGTGCCTCAACAGT | *Oryza sativa L.* |
| AMPL1563419 | CAAAGAGGGTAAGCTGAGGCTT | CCTAAGAGCGGTACAATAAAAGGCTATAAA | *Oryza sativa L.* |
| AMPL1563422 | CTCTCTGCTGCAAAGAGCTTCT | CAGCTAGCCACGATACCTGATAC | *Oryza sativa L.* |
| AMPL1563424 | ATCTTAATACAATCCGATGGCCACAA | CCTCATTTGATGGGACATTAGTCGAC | *Oryza sativa L.* |
| AMPL1563425 | CTTGTATCCCGTACGCATCGTA | ATCAACGGTTCTAGAAAGCCTATGATAAAA | *Oryza sativa L.* |
| AMPL1563427 | CACAATTTTGGAACGGATGCAGT | AAAATTTTCGGAAATTCCGAGGAAATTTCT | *Oryza sativa L.* |
| AMPL1563429 | AGAGAGCCAGCGTCTCTCATAA | GAAGTCGGCCTCGAATCTATCC | *Oryza sativa L.* |
| AMPL1563435 | TTGTCACGTAACGCCAACTTTTG | GCTAGGCTTGATTTCAATCCAACAAATATG | *Oryza sativa L.* |
| AMPL1563436 | AAAATGTAACGGAGGGAGTAATAAAGGATC | CACTTCTCCCTTATCTACGTACTCCA | *Oryza sativa L.* |
| AMPL1563437 | GGGTTCTCCATCGTCGGATTTG | CGAGTTGCGAGCTTCATTTGTTC | *Oryza sativa L.* |
| AMPL1563438 | GGGTTTGCAGCAACAGTGGATA | GGAATGCCATCGTGTCGTCAT | *Oryza sativa L.* |
| AMPL1563443 | TGCATTCGTGCGAGAAAAATCC | TGCATGCAGACGAGCAGATAAT | *Oryza sativa L.* |
| AMPL1563445 | TCAGACATTTCTTCTCATGGTCCATTG | GTGAGTTGTGACAGGATAGTACTACTAGT | *Oryza sativa L.* |
| AMPL1563447 | TAGCTGCAGCTATAATTCCATGTATAACG | CCCAACTCCTGACATTGAAGTATAAAAAC | *Oryza sativa L.* |
| AMPL1563449 | GGTAGCGACGGGCTTTAGATTT | ACTAGGAGATACACAAGGTTTGAGAAAGAT | *Oryza sativa L.* |
| AMPL1563451 | CTGATTGAAGCTTTGGTTCTTAACTATTCC | GGGAGTAAAAATGACCCAAATAACCC | *Oryza sativa L.* |
| AMPL1563454 | TGGGTTTAAGTTGTACAAAACCATACGA | AGTAGATCAAAGAAAAATTAGTTAATGGTGTGA | *Oryza sativa L.* |
| AMPL1563456 | GTGTGTTTAGTTCATGCTAAAATTGGTTGA | GAGAGAGATATATAGATCGATCGATCGCTT | *Oryza sativa L.* |
| AMPL1563457 | GTAATGGTAAAAACATCTATAGCTGACCTTAGT | GGACTGGCTAGCATCTGACC | *Oryza sativa L.* |
| AMPL1563460 | TAACGTATGTTCACAACTAAATTTGTCCTTTTC | CCCTCGTTATTTGTATACCACTCAAATAGTTAT | *Oryza sativa L.* |
| AMPL1563462 | GTGGATATGCAACGATATCATTTTTACTCC | CCCGTCCTAAAGTGCTTTTTGAA | *Oryza sativa L.* |
| AMPL1563464 | TGAAAGATATATTTTGGAGACAGCGTCAT | GTGATTCAACATATAATGGCTGATAATCCC | *Oryza sativa L.* |
| AMPL1563469 | AATTAATCAGCTCATTAACTGGTGTGTTAG | AGTGTATCATGCATCATTTGCTTCTTTG | *Oryza sativa L.* |
| AMPL1563472 | GACGCAAAACGGGATCAGAATC | CCGTAGTTAAGCTGAAACAGGCTA | *Oryza sativa L.* |
| AMPL1563474 | CCATCCCTTGCTGGTATATGGTG | CGTAGTATCCTATTCGGGAGAGAACTG | *Oryza sativa L.* |
| AMPL1563482 | GGGTGCCCAAAAATTTGGTTAAGA | AAGTGGTATCTCATGATACCTTCTTAAGGA | *Oryza sativa L.* |
| AMPL1563488 | GCTAATGAGAGGGCTGTGTTCT | CCCACAAGTAAATCGTTAATTTGATGTGAA | *Oryza sativa L.* |
| AMPL1563489 | CACTTTCAGTTTACCTCTTTTCAAGTTACC | TCTCAACTATCTCTACTCAGGGTTTACG | *Oryza sativa L.* |
| AMPL1563491 | TGATTTTCCGTACACAACTTTGACCA | GAACAAAATGGCTGGCCTTACT | *Oryza sativa L.* |
| AMPL1563493 | AGTAGTAATAAGTCATCGGAGCGAGTA | CTGGCAGGTAGGGCAAGGAAAAA | *Oryza sativa L.* |
| AMPL1563495 | TACGTGATACGTCTATGCGTAGGA | TCGAGGAGATCTCTCTAGTACTCCTATC | *Oryza sativa L.* |
| AMPL1563497 | GGAGCGTGACTCGAGCAA | GCGAGCTAATTAGCACAAATTTTACACTAC | *Oryza sativa L.* |
| AMPL1563501 | ACGCTTAAAGGGAAATGATCAATGAGT | TTGGGTATCTAATCGCATACAAAGGAATT | *Oryza sativa L.* |
| AMPL1563509 | TCCCTAAACAGAGGTCGGGAAT | GAGGAATAGCCAAATAGGTGGAATACC | *Oryza sativa L.* |
| AMPL1563511 | GGCAACCAAATTGTAGCTAACATTCT | ACACCAAGTCTTGCTGAATTGTATTAACTA | *Oryza sativa L.* |
| AMPL1563512 | CCTTTCGATGGGTTCTTGTTTTCG | CTGACTTAGTCAGAATATCAAACGTAACCA | *Oryza sativa L.* |
| AMPL1563513 | GGATATCCCAACCATAGTATTTGTAGAGC | CTACACTAGTAGTCGATTGCGTGTT | *Oryza sativa L.* |
| AMPL1563520 | CGGTATTGTATCATCTTTCCCATTGGA | CTCATCTCCTGTTCTTGCCCAT | *Oryza sativa L.* |
| AMPL1563524 | CCTTACTTGGAGGCCAAGATGA | TTGCTGACTTTATCAACGAGGGATAC | *Oryza sativa L.* |
| AMPL1563525 | GCCCATGTTTGGAGATCAATCTC | TTTGCCCTGTACTCAAAACATAAAAGTTG | *Oryza sativa L.* |
| AMPL1563529 | GGGAGAGAGAGAGGTGAAGAGATT | GCAGAATCGGACGGTAGTACAGTAT | *Oryza sativa L.* |
| AMPL1563533 | GATATTCCCGTGGGTTGCTTTG | ATAAACCAAGATAAGGACAACACAACAGCT | *Oryza sativa L.* |
| AMPL1563534 | TCTATTTAGTGCCAACTTCGATACCATG | TCTCTACTGTTTTATCTCCTCGACATATGT | *Oryza sativa L.* |
| AMPL1563535 | CAATTCTGTTCAGACCATCAGCCTA | GTTACTAAATGGGATGTTGTACCTACTGTT | *Oryza sativa L.* |
| AMPL1563536 | AATTTCAAGTCTTGGAATGTTTTGTGCT | GGGTTAACTTCTTTTGGAGTCCCTAAG | *Oryza sativa L.* |
| AMPL1563539 | TTCGTTTTCTATGCACACAACTTGTG | GTTCTTTGGAGTGGAAACCGTTTG | *Oryza sativa L.* |
| AMPL1563543 | GGTTTTTGGATAGTGGTTGGGAGTT | CCTGTCGTCCCTGTCTCTTGTA | *Oryza sativa L.* |
| AMPL1563545 | TATGGAAGTTGAAGTCCTGTAACGAAAATT | GAATATTTCAGGCTCTGGCATATCCAA | *Oryza sativa L.* |

| | | | |
|---|---|---|---|
| AMPL1563548 | AGTACTAGAACTTTTATGCCTTGCAACT | GATGCTTCATTTCGTACGAGAAAAAGG | *Oryza sativa L.* |
| AMPL1563549 | AGGTATTCCTTTTTGCATCTGTAATTCTGT | GCAAAGTGTGCAGGAGATGGTAA | *Oryza sativa L.* |
| AMPL1563550 | CCCTTGTGTAAAATGGTTTCAGTCA | TTGAAAATGCAATTAGGTTTTTGGGACA | *Oryza sativa L.* |
| AMPL1563552 | CATATGAACTATTATAGCCATCGTTTCGC | GGTATGTGTGTATGCATGCATGC | *Oryza sativa L.* |
| AMPL1563555 | GCGAAACGGTATATTTAGAGCAGGT | TACAAATTATTTTCCGTTTGTAAGAGCAAGTTT | *Oryza sativa L.* |
| AMPL1563557 | TGCATTTCTTTTTCGTATTGCGCTT | CCTCCTTTGTAAAGAATCTCATTGGTTTG | *Oryza sativa L.* |
| AMPL1563559 | CCAACTGAAGGTTTGTTGTTAACTCC | TGAAAAGATAGTTCATCACATTGTTCAGGT | *Oryza sativa L.* |
| AMPL1563561 | AATGGTACTACTCATATGAGGCTAGGTT | GCTTCATGATTTGACAATATAATGCTACAT | *Oryza sativa L.* |
| AMPL1563563 | CCCACTATTGTTGGAGCCAACA | CACAGACCCTTTGTTATAGAGACAACATAT | *Oryza sativa L.* |
| AMPL1563567 | TCACTGTATCGTCATACAAATCATAGTCCT | GTGGCTTTTAGCTTTAGCATTTTAAGTACA | *Oryza sativa L.* |
| AMPL1563568 | GTTTTTAGCCAGAAAGAATTGAATGTGC | TCTTCAAGTAACACAAGGAGAAAACGT | *Oryza sativa L.* |
| AMPL1563569 | TTTCTTAGCGTTGGTGCAAAGATAAAC | GACAAGAGGAGGCTAAAATGATATTGAACT | *Oryza sativa L.* |
| AMPL1563576 | CAGTCAATCTAGTTAACTGAGGTTGGT | CTCTCTCTCGATCACTTGTGGAAAAG | *Oryza sativa L.* |
| AMPL1563577 | GCGTGCATGTTTGTTCGTACTA | GCCGTTAAGCCGTCACATGTAT | *Oryza sativa L.* |
| AMPL1563587 | CTGCTCAACAAAGAACTGCTCAA | CCTAGGCCTCAGAGCCTCTTAG | *Oryza sativa L.* |
| AMPL1563589 | TATGTGCTACCAGAGTATATCACAGCA | GAAACATGCCCAAAGAGCACATC | *Oryza sativa L.* |
| AMPL1563591 | ATTCATAATGTCGGCATGAACAAATACG | TTACGTGTTTAGCCAAGCCAATAAGTA | *Oryza sativa L.* |
| AMPL1563595 | TGTGGCACATATGCACACCATA | GGTTCAAAAATAAACCCTACCCTATTGGA | *Oryza sativa L.* |
| AMPL1563598 | ACATTAACATCACGCGGATAACCA | GGCATGATGGGAAATTGGGAAGA | *Oryza sativa L.* |
| AMPL1563603 | TATGTATCAAGGTGATCCAACCAAAAAGT | CCTAGCTGGGATCGAAGCAATG | *Oryza sativa L.* |
| AMPL1563605 | GGCCGAATGAACTATAGATGCCT | ACCAAGATTTATAAGACCGTGTGGTTATC | *Oryza sativa L.* |
| AMPL1563608 | AGCGAAATGTACACCCAAACCA | GGGCACAGGTATATAACCCTCTTGA | *Oryza sativa L.* |
| AMPL1563609 | CTCTGCGATAGATTGTGGACGT | GGTAGCTGGAGCAAGAGAAGG | *Oryza sativa L.* |
| AMPL1563611 | CAAAACGGTGGTGGTAAATACGG | GTGGTATTTGGATCTTCGAAACATGT | *Oryza sativa L.* |
| AMPL1563614 | CACATCCTCCTGCAAAATGACTC | CCATCCACTTTTGCAACCATTTCC | *Oryza sativa L.* |
| AMPL1563616 | TCCTGTATGTAATGCTAGTATTTGTTAGCG | CTCACAATGAATTTTGCCTTAACTAAACCT | *Oryza sativa L.* |
| AMPL1563620 | TTTTTGACGGAGGGAGTAGTAATCAA | GACGGAGGGAGTACTAGTAATTGC | *Oryza sativa L.* |
| AMPL1563623 | GGTGGTGTAGTGGGCTAGATCT | CTTGTGTAGCTATGATTCTTCACATCACT | *Oryza sativa L.* |
| AMPL1563626 | CCTATAGGAGGAAAGTATAGGTGCTGA | ATACCTATCGAAAGATTAGGACCTAACGT | *Oryza sativa L.* |
| AMPL1563633 | AATTAAGTCTGTCTCTGTGGCGA | CGATTCTATTGAGCCTAATTAACCCATGATTAT | *Oryza sativa L.* |
| AMPL1563635 | AACAGAGGATTACCATTTTACCAGCA | TCCATGTGCTTCTGAAACTTGATTTTG | *Oryza sativa L.* |
| AMPL1563636 | ACTAACCTCACGCTGCATATGC | GTAGAGCAAACAATGGAAGCACATAG | *Oryza sativa L.* |
| AMPL1563637 | TTCTCGTCTCAATCAATCACAACTATTCTC | TTCCAAACAGAAGTATACATAACATGCACT | *Oryza sativa L.* |
| AMPL1563639 | GCATCTGCATCTGCAAATGGTG | GCACTGTGCGAAAATCATTGGAG | *Oryza sativa L.* |
| AMPL1563641 | TCTAGGTCACCGTCAATTGCAAG | CAGGAGGGAAAGGCTCTACATG | *Oryza sativa L.* |
| AMPL1563643 | TTGTTGTAACCTTGATGTTACAAGGTAGT | GGGTTTCCCAGCTTCTAGCTTTTTATT | *Oryza sativa L.* |
| AMPL1563648 | ACTGGACTTGTCTGGTAGTGATACA | TGATGAATGCATCCTCAGATAACATCG | *Oryza sativa L.* |
| AMPL1563651 | TGACACTGGAGTTGATTACAAAGACAA | ATCATCATGGATTCCTTCGAAGAAAGTTAT | *Oryza sativa L.* |
| AMPL1563653 | GCAGTATATGGATGAGAGACGAATGAAC | GTTTGACTTGGCATAAAACAAAAATGGC | *Oryza sativa L.* |
| AMPL1563656 | GCGTCAGAACTTCAACACCAAG | CCTTAGCCTTGCTCAGCTATTTAATTG | *Oryza sativa L.* |
| AMPL1563657 | CTCCTCCCTCTACCGCCATA | TATACCTCAGTAACCGCACGACTAT | *Oryza sativa L.* |
| AMPL1563658 | CATACAGATTACACTTGGATCTCAGGA | GCACTGATCCAACCATATATCCAAGG | *Oryza sativa L.* |
| AMPL1563663 | TGTGCATGACCTTTGCTCAGAA | AGTAACTGGATTGATCGATGCAGATTT | *Oryza sativa L.* |
| AMPL1563664 | GCTGTTGTTTCTCTTGCCTTCA | GTTTGGACTGAAAAACAACAACCGT | *Oryza sativa L.* |
| AMPL1563669 | CGATCGACTTCAGGCCTTGTTT | ATGCTGGCTACAGATTTATAGCTCG | *Oryza sativa L.* |
| AMPL1563670 | CGTGTCAATGGAAAACAAGGCA | GCTAGCCTGTGAAAAATAATTATTTTCGCA | *Oryza sativa L.* |
| AMPL1563672 | TCCTCTCCAAAGATTCCGACAAC | GCTCCCTTCCTTCTCTGCAAAC | *Oryza sativa L.* |
| AMPL1563673 | CATCCGAACGTAAAGGAGGGAT | AGTTTAAAATTGCAAATGACACGGCTTA | *Oryza sativa L.* |
| AMPL1563675 | ACGCCAAAACTTTACACTAGGCA | ACGGGCAGGAAGATTTGGAATT | *Oryza sativa L.* |
| AMPL1563684 | CCCATCTTGCACATCCTCTTGA | GGCTAGAGGGATCTGCTCTGAA | *Oryza sativa L.* |
| AMPL1563685 | GACAGAGAGAGTACATTATTTGTGTTATTTGC | ATCTATTTTTAGTCCCGAGTATTTGAACAGG | *Oryza sativa L.* |
| AMPL1563686 | TATTATGCAGACACGGAAGGACACCAT | AATTCCCAGTTTTGAGCTCTGCTAT | *Oryza sativa L.* |
| AMPL1563687 | GGCTGCACCGATTAGTCACTTT | AACATGAATTCTAGTTTGCAAAATGCCA | *Oryza sativa L.* |
| AMPL1563689 | CCTGTGGAGTTATCGAGAAAGCT | ATCTCCAATGAAATGTATTTACATTGTCTCCT | *Oryza sativa L.* |
| AMPL1563690 | ATGTCAAATTTCAGAAACAATGACCTCTTG | CTTCTTGGTCATTTCTGGTGGTAGTTTATA | *Oryza sativa L.* |
| AMPL1563695 | CCAACAGCAAAAGGTAAACCACATC | CAGTTCAGATTAATTATGGGTCTGGGTTT | *Oryza sativa L.* |
| AMPL1563698 | CCACGAGGTATACCGATGTTTCA | CAAACCCAACATATTAATTTATCCTGCACA | *Oryza sativa L.* |
| AMPL1563701 | ACTTCTATCATCAACCAGAGGTACATAGAA | CACTGCAGAGTAACAGCACACT | *Oryza sativa L.* |
| AMPL1563704 | CCATCCATTTCACACTCCTCACT | GCTTCCCTTCAAAATTGATTGAAATCCATT | *Oryza sativa L.* |
| AMPL1563705 | ACTGGGAGGGAGTATATATAAGCCAAA | CAATAATAAGTATAGCCACTGTTGGTATTTCT | *Oryza sativa L.* |
| AMPL1563712 | GAAGTGCTAAGTATATACTTCTTCCGTCA | TTTAGATATTAAGCCCTGTTGTGCCTT | *Oryza sativa L.* |
| AMPL1563713 | CCTCCGTCTCCGAGTGATATCA | CGAATTGGTGGAGAGAGGTGAGA | *Oryza sativa L.* |
| AMPL1563716 | CATTAAGATAATAAACCAAGCTTGGCACAT | GGCATCACTCATATTCCATGAAAAATATCA | *Oryza sativa L.* |
| AMPL1563718 | GAAACATCACGAGTATAATTGCTGAAACA | TTTGCAATAAATAACGGAAAATGTTAGCGA | *Oryza sativa L.* |
| AMPL1563721 | GTGAACAGAGCAACCACGCAACATT | CATTGTAAAGTTGATACCTCCTGGTATCT | *Oryza sativa L.* |
| AMPL1563722 | CCAGCAAGAATGCTAGAGTGCT | TCCCTTTCCAACATCTCTCCATTG | *Oryza sativa L.* |
| AMPL1563723 | CACAGCCACAACGAGAATGTT | GCATTTGCTCTCGGTCTGGA | *Oryza sativa L.* |
| AMPL1563725 | ATTCGGATTTGTTTTTCGGAAACAGTT | TGACGGTGCAATACAGAGGAAAAA | *Oryza sativa L.* |
| AMPL1563728 | CGCTCGGGCTTTGATGTCAA | GCCTCACCTCCACGAACATC | *Oryza sativa L.* |
| AMPL1563730 | GGCAGCTCTGATAACAACGGT | TAACCTTGGCCGTGAACCAAA | *Oryza sativa L.* |
| AMPL1563733 | GGTGAGAGCGATCGAAATGAGG | CTTTGCAATGTGCATGCAGTTTG | *Oryza sativa L.* |
| AMPL1563737 | CTGGCTCTCACCATCTTCTCAG | GCTTCAGCGGGTATCCGAATAAC | *Oryza sativa L.* |
| AMPL1563738 | CATCTAGCTCATACTTGCTTAGCTGT | GACATGCGGATAGTATGCGGAT | *Oryza sativa L.* |
| AMPL1563741 | GCACCCACTGACCCACATAATG | GCCTGTAGATCGTGTTGCTACTATTAC | *Oryza sativa L.* |
| AMPL1563742 | GCCGTAGTATACACCTCCACCTT | CCTCCTCCTCCAAGATTCCCAA | *Oryza sativa L.* |
| AMPL1563745 | GTCACTGTTAATGCATTAGTACATCAGGA | GCTATAGAAACTTTTCATCAGCTCAGAACT | *Oryza sativa L.* |
| AMPL1563748 | GCTTAATTGTACTACCATGGCCACAT | AGGCTCTACCAGGGTTACTGAC | *Oryza sativa L.* |
| AMPL1563751 | ATGCCTGCATTAGGTGGGATAAAA | AATTCGTAATGTGGCAAGATGACAGTA | *Oryza sativa L.* |
| AMPL1563752 | GCATTGCGACGAGCTCTGTA | CCTTGACATTCAGCCATGAATAGC | *Oryza sativa L.* |
| AMPL1563755 | AGGAGATGTGAAAATGTGAATAGTGAACA | GTCGGTGCTGCTATCTTAGTTTCT | *Oryza sativa L.* |
| AMPL1563756 | TTGCCAACACCTTCCTCACTAG | GTCTAAAAGAGGTTCAGTGCTGTCAA | *Oryza sativa L.* |

| | | | |
|---|---|---|---|
| AMPL1563757 | AGATCTTCATAAACGTCCAGGGATCT | AATCAATTAATCGTGCTTAATCCTGTCTTG | *Oryza sativa L.* |
| AMPL1563759 | GATTGAAATCGATGCAGTCAACTCG | CTCCAGAGCAAAGCTACCAGCT | *Oryza sativa L.* |
| AMPL1563764 | GCAAGATCTGCATATGTATAACTGATGGT | GCTTTTCGAGTTCGGTAGCTATATTTTAGA | *Oryza sativa L.* |
| AMPL1563767 | CCCAATCACTGTGTCTCAGATCAT | GGTTATTGGTGCACAAGGCATAATAAAC | *Oryza sativa L.* |
| AMPL1563773 | CACGTCAGATCCTCCACATGT | TGACTTCTCTCACCTAGCAAAGCTA | *Oryza sativa L.* |
| AMPL1563776 | CAACTTCTTCTGTGTTTCTAAAAGCGAA | ATGTGGATCGAATTTAAACTTGCATGTTT | *Oryza sativa L.* |
| AMPL1563779 | TTTTGTGGAATGTCGTAAACATGAGC | GCTAGCTGCCAGCGAAAAGTTAT | *Oryza sativa L.* |
| AMPL1563785 | CGCTCTTGATCATCGCGTTTTT | CTCCAGGCCAGGAAACGACAAGATA | *Oryza sativa L.* |
| AMPL1563788 | CACATTTAGTGCGGACGGAGA | AAAATAGACTTTTCAAGAGGATTTGATTTGGG | *Oryza sativa L.* |
| AMPL1563790 | AAAAAGTCACTGTGTGCAAACATT | TGCTTCGTCGAGGTAAAATCCTTTT | *Oryza sativa L.* |
| AMPL1563792 | CATGAAGGAACCTGGGCATAGAT | AGACGAGAAAGTTGTAGTTTAGAATGGG | *Oryza sativa L.* |
| AMPL1563797 | TTGCTGACTCAAAGTCAAGAGCT | GTTGTGGTTGCCAAGTGGAA | *Oryza sativa L.* |
| AMPL1563798 | TGCCAAGAAATTCGAGTCACCAA | AGGAGAAATTAAATCACCTCGGATGTTG | *Oryza sativa L.* |
| AMPL1563800 | ATCCTTCTTCATGGATTTCTTCTTTCGT | TGGGTATACTATACCAATACACGACGTAT | *Oryza sativa L.* |
| AMPL1563805 | GTTTACAGGTTTCATCATTTCTCCACATC | CCCATGGGTGTGTGTAGTTCAC | *Oryza sativa L.* |
| AMPL1563808 | TCTTCACATGCACTCACATGCA | ACAAAATTTCCGAAATTTTGGTCCTATTGG | *Oryza sativa L.* |
| AMPL1563811 | CCATCGCACATATAGGGTCTGTT | ACATTTTCCTCACCATCCCAACT | *Oryza sativa L.* |
| AMPL1563814 | GTTTTTAGAGTAAATTGCATCAACGGTACA | TGATGGCAGCACCGGTTC | *Oryza sativa L.* |
| AMPL1563817 | CGAATAGCCTCCCTGTGACTGA | GTTCTTCCTCTGCTTCCTCTGTTC | *Oryza sativa L.* |
| AMPL1563818 | TTTGCTAATTCGTGTGTAATTTTAGCGT | AAATAGTTGTTCATTCCTGTTTTAGCGAAG | *Oryza sativa L.* |
| AMPL1563819 | AGCTACAATGTCTATTTGCCCTACTATAG | TGGTTTAAAACCAAATTTTGCTAAGCCT | *Oryza sativa L.* |
| AMPL1563831 | GAGATAAATCGGAGAAGCGACAAGT | CTCCCTCCATCCCAAAGTAACTTTATAAAG | *Oryza sativa L.* |
| AMPL1563832 | GTACGCTAGAATTTGTCTTGATTGTCAC | AGTCTTTTTGGTCTATTTGGTTAACCGA | *Oryza sativa L.* |
| AMPL1563833 | TGTCGTCTCAAGTTCTAGCTAGGT | AACAACGATATCAACGGTAGCGAT | *Oryza sativa L.* |
| AMPL1563835 | ATGTTGATAATAAAATTGAAGCCATACTTCAGC | GGAAATAAAAGATCTTTTCACTTAAACATTGGC | *Oryza sativa L.* |
| AMPL1563843 | ACGGCACAACAATAACAACACATG | GGATCTGAGCGGGATCAAGAAC | *Oryza sativa L.* |
| AMPL1563845 | AGCAGATACACGTCGATCGAGTA | GATCAGGACGAGAATGTAGTATTACAGAG | *Oryza sativa L.* |
| AMPL1563851 | CGGGTTAGCCATGGACAAGAAA | CCCTACTGTACACCACCAAAAGTG | *Oryza sativa L.* |
| AMPL1563855 | CCGGGTAAAAACACCATTAGAGCA | CACATTGAACGTTTGAAACTTGCATAAAAT | *Oryza sativa L.* |
| AMPL1563865 | TAGGTTTATGAACATGATGGCTTCAGTT | TTGACTTGATGCGTGTAGATAATAACATGA | *Oryza sativa L.* |
| AMPL1563870 | ACAAGTATTGGTAGCAAACTAAACATTGC | CCCTAGAGGTACAAAGTTAGGCCAT | *Oryza sativa L.* |
| AMPL1563871 | AGCACATTGTATTTTAGACTTAGTGGTGT | TGAGTAAGGGCAAAAAGGACATTTTATACT | *Oryza sativa L.* |
| AMPL1563874 | ACTGTCCAGTATTGTGGTAGGTCT | GCTACTGAAGGAGCCTTCTTCTCA | *Oryza sativa L.* |
| AMPL1563878 | GCTAATAAGGTTTATCGTTTTGTTAGCAA | GGAACTAAACAAGGCCATTGTTACC | *Oryza sativa L.* |
| AMPL1563882 | CACAACTATCGCTTCATTACTCCATCA | GACTTGGTATAAGTTTCTGGTCGGAAT | *Oryza sativa L.* |
| AMPL1563886 | TCTCAATAGATCAGATTGTCCAAGCTACT | AGAAAATAAAGTCCAGCAATCATCTTCAGA | *Oryza sativa L.* |
| AMPL1563888 | CTTGAACATTTCACATGCTCACCTT | CAACAGTGTTAAACTGTCCGTTTGTATAC | *Oryza sativa L.* |
| AMPL1563889 | GCCTCTTCAGTTTGGTCCATTTTG | GGCATAATCCTGTTGCACAGATG | *Oryza sativa L.* |
| AMPL1563894 | CTAGCTAGGGCCTTTCATATGTCG | GATCAGTCAGGCGTTGTTAGCT | *Oryza sativa L.* |
| AMPL1563895 | AGTCCCTTAAGAAAATGCAACGAGT | AATGCTTCGCTTGTAAATTAAGCGTAAT | *Oryza sativa L.* |
| AMPL1563898 | GTTTTTACATGCCTCAAGGCAAGT | GACACAATCACAATGTTTCCAAAAGGA | *Oryza sativa L.* |
| AMPL1563899 | GCACAAATTGGTACGTGCACTT | TGTAACTAGGCCTGTGGTGTACATA | *Oryza sativa L.* |
| AMPL1563900 | GAGATTCGATTGAGACCCATGCAT | AGCAGTTTGAAGAATGTGCTAATGAAA | *Oryza sativa L.* |
| AMPL1563902 | TAGATTTTCACCAATAAGCTAGCTTCCTTT | CCCTTCATATTTCTACTACCGCTAACTG | *Oryza sativa L.* |
| AMPL1563907 | ACCCGCTTGCACCCTTAAAAATA | GTTCACGTGGCTTGACACATTG | *Oryza sativa L.* |
| AMPL1563909 | GTTTCTCTTGAGCAAGGAATCCGA | TTCTATCTGCTTAACGCTCCACATTATC | *Oryza sativa L.* |
| AMPL1563910 | GTCACAAACTGTTCGAATTTTACATACTCA | GTTATGAACGTGCACAAGAGGGTA | *Oryza sativa L.* |
| AMPL2000000 | GAGGGAGTAGTATGGTTGTTTTGTGTTAG | CTCTTGCTAGGAGTACGTGTGATGTC | *Oryza sativa L.* |
| AMPL2000001 | CGATCGATGAGCACTTGCAAAGT | TGGAGTTGGAGAGACGTATATATGAGAAGTA | *Oryza sativa L.* |
| AMPL2000002 | TTCCAGTTTCTCCCTTTTTAAACTCA | TGAAGTGTGTTGTTTCTGTGTGAAACTA | *Oryza sativa L.* |
| AMPL2000003 | GATGTATGGTCAAACGTATCTCAGAAAA | CAAACTGTGTGCAGTCGCAATAG | *Oryza sativa L.* |
| AMPL2000004 | TTTCAAGGTATCATGGTACTACGATGTAGT | GGCTAAAGGTCAACTGTGATGTGTAG | *Oryza sativa L.* |
| AMPL2000005 | AATTGCCATTGCTCTTCGCTT | GCTAGGTTTTCGACGGTGAGGTTA | *Oryza sativa L.* |
| AMPL2000006 | TGAGGATGAACCCAATACTGAAAAA | TGAAGAATTTTAATTGGCTTTTGGTC | *Oryza sativa L.* |
| AMPL2000007 | CTCCCTCTTTCTGCCTTCCTATG | TTGCGGTGCAATTAAATAAGTGC | *Oryza sativa L.* |
| AMPL2000008 | AGCTTCGTGCTCCAGCTCAT | TGCAAAACGAAGAAAATTCTCTCAC | *Oryza sativa L.* |
| AMPL2000009 | TCATGCTCATGCATGTGCACT | ATCCTCAATGACCACTAGTCAAGACTC | *Oryza sativa L.* |
| AMPL2000010 | AATCCATTAGCTGGGCTCTAGGTT | TCTCGTAGTTTGTACAGGCGGAAT | *Oryza sativa L.* |
| AMPL2000011 | TTATTCCTGGTCTCATGTGTTCTTGA | CACTGAGTAGAAGGCAACTACAAGAGGTAG | *Oryza sativa L.* |
| AMPL2000012 | CCACGAGCTCCATATCCAGTTG | CAGCCTCTCCATCCACTCTCTCTAC | *Oryza sativa L.* |
| AMPL2000013 | GGATTCAAGTCATGGCAGATTGA | CAAAACTCAAGACTGAGCAATTCG | *Oryza sativa L.* |
| AMPL2000014 | ACAGGAAATATGTGTTCAATGATTGC | TTAGCAAAATAGTAAGGACACTACTGGTCA | *Oryza sativa L.* |
| AMPL2000015 | CGTAAGCCCGTAACAACCGTAA | GACTTTTGATCTCTTCAGGCATTCA | *Oryza sativa L.* |
| AMPL2000016 | ATACCTGGGATTAAGTGATCAAGCC | CTAATCCCCTCACCCTGATTATCAC | *Oryza sativa L.* |
| AMPL2000017 | GCAGATTTGCTGCTGCTACAGTT | GGATGGAGAGAGCATGATGAAAAA | *Oryza sativa L.* |
| AMPL2000018 | TTTTTGAAGATACACGAGCCATAGAA | TTAGCAGTGACTAAACATTTTTCCACA | *Oryza sativa L.* |
| AMPL2000019 | TGCAACGTAGTCCCACATACCAA | GCACTTGGAGAACGGTAGTGGTATT | *Oryza sativa L.* |
| AMPL2000020 | ACTAAACAAGCCCACAGTCGTAATG | CAATGAGTACGATGGTGTAATTAGATTCA | *Oryza sativa L.* |
| AMPL2000021 | ACCGAGGGTTTTCTGTTAAGCAC | TTGACGCATCTTCTTCCCTTCT | *Oryza sativa L.* |
| AMPL2000022 | GCCTTCAAGGGAGGGAAAGCTAC | GTACTGATTTTCATAGAGCACACTTTTGTG | *Oryza sativa L.* |
| AMPL2000023 | TGGACCACGATATGTGTAAATCATG | TCACCATGTTGGCCAAAACTG | *Oryza sativa L.* |
| AMPL2000024 | CGTCTTTTCGTTGTCGTATTGAGG | GCTAGGCACAATAATTCCCTCGAT | *Oryza sativa L.* |
| AMPL2000026 | ACACGGGGATGATAAGGTTTAATCT | CCACATCCTCTCAAACAATCACAAA | *Oryza sativa L.* |
| AMPL2000027 | GCAAGTTGTCCAGGGCAAAATATATA | TGGAAGTGAAGGAAAAACAGGAGATA | *Oryza sativa L.* |
| AMPL2000028 | CCCACAAACCCTAGGAATTTTGTTT | CCCAAACTCCACCTCCTCTGAGT | *Oryza sativa L.* |
| AMPL2000029 | ACGCCATCCATCCGATCAT | AATCCTTCTTTATCTCCTCCTCGTGTA | *Oryza sativa L.* |
| AMPL2000030 | CCATTCATATTGGTTCTACATGGTACATG | TTAAGTTACCTGACGGTCATTTCATTAAG | *Oryza sativa L.* |
| AMPL2000031 | TCAGGTATGCACCCATTACGAAG | GCAAGCATATCACATTTGGAACAAT | *Oryza sativa L.* |
| AMPL2000032 | CACTACTGCGGTTCTAAAGGATATTCG | GAACGAATTAGGTATATTATTGTGGCCTT | *Oryza sativa L.* |
| AMPL2000033 | CACATGGATTCACCTCTGTTATGATTT | GATCGGTGTAGCCCAGAAAATG | *Oryza sativa L.* |
| AMPL2000035 | GGGACTAAACTGGACTTAGGCGAA | ATAGCCTATTCGGTTTGATCAATTGT | *Oryza sativa L.* |

| | | | |
|---|---|---|---|
| AMPL2000036 | GGTTTAAAAGATTTGTCGCGCAATT | ACTCCAAACTGTACTGTATAAGCCT | *Oryza sativa L.* |
| AMPL2000037 | AACTCAACCCTTTGATGCAATCTG | TGATTAAAATAGGAATTCAAATTGGTGCA | *Oryza sativa L.* |
| AMPL2000038 | GCAAGCAATCAAACACCTGCAT | CGTTTGGAGGTGCCATTTCCT | *Oryza sativa L.* |
| AMPL2000039 | CGCCTCCTCCTACACCTACTGTT | CGTCGTCGTAGAGGAGATCGAA | *Oryza sativa L.* |
| AMPL2000040 | AGGTGTACAATGGGACGAGATTAAAG | GATTTTATTTAGGACCACCAACTGCTATT | *Oryza sativa L.* |
| AMPL2000041 | GGGACTTTGGGAGTTTTGGGAT | CATCCTGGTTCTCTCGACCAA | *Oryza sativa L.* |
| AMPL2000042 | TCGAGGAAGGAGGGAGGAGTT | TGCAATATATAGTTGCACTCACTCCTCA | *Oryza sativa L.* |
| AMPL2000043 | TGAAAAGTGCACACTTCCACATG | AGGCTCCACTTCATGTTGTTTAAATC | *Oryza sativa L.* |
| AMPL2000044 | CATAATCACGCCGTGAGAGAGA | AATTTGGTTGTACTGCGTTCTGCTA | *Oryza sativa L.* |
| AMPL2000045 | CCAACTCCAACTCCACCCTTT | CGGAGTTGGCGGTGACAAAGT | *Oryza sativa L.* |
| AMPL2000046 | GGACAAAACCGCTCTGGATTAGA | AAAATTGACCCTAGCTACTCCATTCAT | *Oryza sativa L.* |
| AMPL2000047 | CTCATACACCCTCCTGTTCGATCT | GGTTTACCTCACATATACTCTGAACAATGA | *Oryza sativa L.* |
| AMPL2000048 | TCCGATGGCGACACATATTG | GAAAAGAGAAGCTAGCTCCAAGCAT | *Oryza sativa L.* |
| AMPL2000049 | TCCGGCTCAAATATCCGAATC | TTGTTGTAGAACACCGAGTTGGAGT | *Oryza sativa L.* |
| AMPL2000050 | AATTGCGCTATTTTGTACTAGCTTGTG | AGGTTGTCTCTTAGGTAGACCCAGATAATAC | *Oryza sativa L.* |
| AMPL2000051 | CGAATCGCTTCTAGACCTGCAA | GCAATAATTCGGGAGTAATTCGTACTT | *Oryza sativa L.* |
| AMPL2000052 | ACAAACTCCACTCTAGCCACCTCTT | GTGAAAATCCAGTCTCAGCTCTCTTT | *Oryza sativa L.* |
| AMPL2000053 | ACAAAAATCGATCCATGCCACTA | CAGACGTAGCTGCTATTGGGAAG | *Oryza sativa L.* |
| AMPL2000054 | ACGTACGTGTCCAGCGTTTACAT | GAATCATGTATTTTGTCGATCGGTT | *Oryza sativa L.* |
| AMPL2000055 | TTGTGTAATGCTCCAAAAGGCTAGTA | CATGTGTGAATTAAGGAAGCAATCTTAGA | *Oryza sativa L.* |
| AMPL2000056 | CAATAAATATCGTGCATACATGCATG | CATCAAGCATTGTGTGATATATTGTCCT | *Oryza sativa L.* |
| AMPL2000057 | AAGCGATGCCTATAGGGATGAAC | CCTTCCTTCTCTTCTAGCATATTGTGC | *Oryza sativa L.* |
| AMPL2000058 | GGCCATAAGAAAACCCTCGTGTT | CCGGAACCTTTTGGCAAATTAAA | *Oryza sativa L.* |
| AMPL2000059 | GACCCTGTAACCTCAGCTACAATACC | CACCACCAAGGACATAAGCGA | *Oryza sativa L.* |
| AMPL2000061 | GCTACTCGCAAGGCTCAACTGAT | CGGTGCTGATGCAGCATTAATC | *Oryza sativa L.* |
| AMPL2000062 | CGTAGTTGCTGACTGTCGTGTGAATT | TGCAGCAGGTGCGAATGTATTA | *Oryza sativa L.* |
| AMPL2000063 | GGCTCATCAGTCCTTAGGGTGA | ACCATGCATCGAGAAGAAAAGG | *Oryza sativa L.* |
| AMPL2000064 | CCGGACTCTTCCCATACCAAGA | GACATTGGCATATGCTCAACTAAATTTAT | *Oryza sativa L.* |
| AMPL2000065 | TCCCACCGATGCGCTATACTA | GCATTGATGATGAATGAATAACGG | *Oryza sativa L.* |
| AMPL2000066 | AGCACATCGTTTTAGTGGATTTCTG | TTCTTGTCACATCAATATTGCTTGC | *Oryza sativa L.* |
| AMPL2000067 | TGGGACGGAGGGAGTATGTTATTAC | CAAGAGCACCAAATTCTCAATTTTG | *Oryza sativa L.* |
| AMPL2000068 | CACTGTGTTAAACAGTGCTACACAAACA | CCATACCGAGTGATCACATACATCC | *Oryza sativa L.* |
| AMPL2000069 | CAAGAGTAAACAAAGAGCAGTTTCTTACAGT | TCCCTGGAAATTGTGAAAGCA | *Oryza sativa L.* |
| AMPL2000070 | AAAAGAGGCAGTTTTGACCAAGATAAT | TGTTGTTTCGGTTACTGTTGTATTGG | *Oryza sativa L.* |
| AMPL2000071 | AGACCCTGCAGCAACAATTTCT | AGAAATGAACGACCACTTTGGAAC | *Oryza sativa L.* |
| AMPL2000072 | TCTCTCTCACTCCTCCACCTCAA | TCCCTCATCAACTATGATTATAGAGCATC | *Oryza sativa L.* |
| AMPL2000073 | CAAACCGTTCCATATTAACTTGAAGG | GCTGTGTTGACAGCTGTCATAGTACTTCTAC | *Oryza sativa L.* |
| AMPL2000074 | GACACAGTGATGCAATTTGTGGA | CATTGGCTGCTTTCCTGCATAG | *Oryza sativa L.* |
| AMPL2000075 | CACCCGAGTGTGTCTAATTGCAG | TGCGTGTAGCAGTTAAGTGTAAACAATATAC | *Oryza sativa L.* |
| AMPL2000076 | CAAACATATGTCCACCAAAAAGAACA | GGAAAAGAGGCATTGCTTATGGAG | *Oryza sativa L.* |
| AMPL2000077 | GGACTACAAATTTGACCATCAGGACA | TTTGCATCTTGCATTTGTAATGTGT | *Oryza sativa L.* |
| AMPL2000078 | CTGCTGCTGATTCAGTTGATCG | TGCCATTTCATTTTTGGTGTTGT | *Oryza sativa L.* |
| AMPL2000079 | TGCAGTATGTATCGGATGATCCA | GCACCAACTAGCAACCTTGCTGTAC | *Oryza sativa L.* |
| AMPL2000080 | CCGGTCGGTTTCCTAACATACA | CTATAAAAAAGTAGTAGCGACTGCCACAT | *Oryza sativa L.* |
| AMPL2000081 | GGTGTTGGTTCTTTAATTAGCACCA | CATGCCATTTACAATCTTCGTCTTCT | *Oryza sativa L.* |
| AMPL2000083 | CTGAATTCTGAAATACTTAGCACGGTT | TGCAGATGAAGCAAGAACTTATAAAACTG | *Oryza sativa L.* |
| AMPL2000085 | CACCCGAAAACTAAGTGGCTGATA | CATCAATTAAAACCTCGTTGCGAT | *Oryza sativa L.* |
| AMPL2000086 | AACTCCCTTACCTAGTTGTTCTCTTCG | TGTTTCTCTTTCTCTTTCAGCTCAAAAC | *Oryza sativa L.* |
| AMPL2000087 | TCGGCAGGCGTATTATTTTCA | AATGGGTCCACTAAAACCAATGG | *Oryza sativa L.* |
| AMPL2000088 | CCGCCTGTTGTTGATGTTGAAT | CAAATCTGAAAAGAGCAAGTGTGGTAC | *Oryza sativa L.* |
| AMPL2000089 | GTCCAGTCCATTACTCCCTCCAT | AATCTAAACACGGCCTCTATATTGCA | *Oryza sativa L.* |
| AMPL2000090 | ATCACGATCAGGATCCTCAAGTTT | GCACGCAAGAAGTATAAAACCACAG | *Oryza sativa L.* |
| AMPL2000091 | GTTGCTATTTGTTGAAAGCTTTAGTCG | TTACCTATAGCGGTTCTCAGAGAAGATT | *Oryza sativa L.* |
| AMPL2000093 | GCTAAGCGAGCGAGGTGATATG | CATAGCGTCTTTACGTACCTCAGAGATAT | *Oryza sativa L.* |
| AMPL2000094 | GGGTTATACTGCTAGGTTGCTACAA | TAGAGAGACTTACTACATGGCAACC | *Oryza sativa L.* |
| AMPL2000095 | GTGTGAGCAGCGGAAGAACTAAC | GCACTACTCAATGATCCAGCATATTACAC | *Oryza sativa L.* |
| AMPL2000096 | GCTTACTTTGGATGAAATTGTGTGTG | TGGCCAAAATGCAGATTCACTAAT | *Oryza sativa L.* |
| AMPL2000097 | CAACAATTAAAGGCTCCTGTGTACGT | AAGAACCAGAGAAAGTAGACATGTAACTGAT | *Oryza sativa L.* |
| AMPL2000098 | CCACCGAGACAGGGAAGATG | AGATTCGTGTAAATCCTTAGAACCGTAA | *Oryza sativa L.* |
| AMPL2000099 | GGAAGATTGTTGGGTTTAGTCCCA | CGTGTTTCCTGCTCCACTTGTCT | *Oryza sativa L.* |
| AMPL2000100 | TGCTGAATCTATACTGTTGAGTTTGCA | GCAAGTCTTTCCTTCTTCACATGCT | *Oryza sativa L.* |
| AMPL2000101 | CAACAACAACAACAACGATGATGA | GGTGATCGAGCTGCACAACTACT | *Oryza sativa L.* |
| AMPL2000103 | TTCATTCATGCAGACTCGCAA | TGGAATGCATTGTCGTTCTGA | *Oryza sativa L.* |
| AMPL2000104 | CATCGTTCGTCAGACTATGCAGA | AGAGATCCAGGAAGAACTACCGTGT | *Oryza sativa L.* |
| AMPL2000105 | TTCACGGGAAATTTCAGAGAGTACA | AGTTGACTTGAAGAACTCAAAGCGA | *Oryza sativa L.* |
| AMPL2000106 | CGGTTGGTTAATCTCTCGGTAGTGT | AAATGAGCCTTTTATGTCCAGAAAATAG | *Oryza sativa L.* |
| AMPL2000107 | CGTATTCTGTCCTGTGTGAATTTCG | GCAAATAAAACAAGTGCCTGAATTACA | *Oryza sativa L.* |
| AMPL2000108 | CACCTGCAAACACTGGATGTTC | CTAGCAAAGCCTTCTTCATGCTCT | *Oryza sativa L.* |
| AMPL2000109 | GGTTTTTCCAATCATCTCTGTTTCA | AAGAGGAACATGCTGAGACTCTTACC | *Oryza sativa L.* |
| AMPL2000110 | CCATGTACAGTATTATTACATGGGAGCTT | CTAATTTAGCACAACAGCAACATAATGAC | *Oryza sativa L.* |
| AMPL2000112 | CATCTCCTCCTGAACACAACTTTGAC | GCATGAGTGAAGATGTCGATGAGT | *Oryza sativa L.* |
| AMPL2000113 | CCACAAAACTGTCGGTAATAATCAGG | GCTAGCTAACCAGATTAGTTCCTTGCTATAA | *Oryza sativa L.* |
| AMPL2000114 | CAGCGATGGAGAGTGAGGCA | GGGAAAAAGTTTATATTACCCTTTTGAACTT | *Oryza sativa L.* |
| AMPL2000115 | CGTCAACAACGGCAAGGACAT | CAACCTGCCTACCATGGTAAAATAA | *Oryza sativa L.* |
| AMPL2000117 | TGTTTGTCACCAATCTATCCTGTAAAAGA | GCCATGCTGAACTGTTCGAGAT | *Oryza sativa L.* |
| AMPL2000118 | GCAACAGATGAACCGGACAATG | GTGAGGTGTCTCCATGAACTGGT | *Oryza sativa L.* |
| AMPL2000119 | GGCCCATAGGGAAGCATTCTA | GCTTACTGGCTTGTTTGCAAAAG | *Oryza sativa L.* |
| AMPL2000120 | TCTTTTTGTGTGTAGGACAAAGGTG | TGGCTCGAAATGTCAAACCAAATTA | *Oryza sativa L.* |
| AMPL2000121 | AGGTGACTATGTCTACAAACACCTATGAAC | CATATGGTGGCCATGGATTTAATC | *Oryza sativa L.* |
| AMPL2000122 | TTCTCTGAGTGCAATCAACTCCTCT | AGTTCCTGCCTGACAGAAAACCT | *Oryza sativa L.* |
| AMPL2000123 | ATGATCAAACGTCACATCAAAAGC | CGAGTACACCGTCGTCCATGT | *Oryza sativa L.* |

| | | | |
|---|---|---|---|
| AMPL2000124 | CGAATTCAGCTCGATTCGAACTTC | CCTTGGGTTTCACTGATCGATC | *Oryza sativa L.* |
| AMPL2000125 | ACACACCCACTAAGACAAATCCTTA | TATGTCCTGCCATGCTCCTTCATC | *Oryza sativa L.* |
| AMPL2000126 | CACATGTGATCGACGTCTTAATAGAAG | TCACCTCTAAATTCACCACTCTTCAAA | *Oryza sativa L.* |
| AMPL2000127 | TGTTAGTTGTTGTTATGGGTTGTGTATTTG | GGGTTCTATCTTGTCTGCTCAAATTACA | *Oryza sativa L.* |
| AMPL2000128 | GCGTATTTGTTTTTGTTTCATTCGATAC | CATACACCAAAAATGATGACTATGTGCT | *Oryza sativa L.* |
| AMPL2000129 | CCTCTTCCCTACCGGTTATGCT | CGACACCCGATGGGTTTTTACTC | *Oryza sativa L.* |
| AMPL2000130 | GCAAGAGCTAGCCACGCAAA | CAGCTCAATGCAACAACTATATATGGA | *Oryza sativa L.* |
| AMPL2000131 | CTATCTCCTAGCGAACGTTAGCTACCT | ACCGGATCTGGTTTGTAGATAGTAAAAA | *Oryza sativa L.* |
| AMPL2000132 | CAGTATGATGTGTATTGCAAGTATGGCTA | GAGGACTAATTTGTCCATGTGCATTT | *Oryza sativa L.* |
| AMPL2000133 | TGCCTCGAGTTTAGTGATGTAGGGT | TTTACGTCAAAGGAGTTGCTTTCC | *Oryza sativa L.* |
| AMPL2000134 | TGCAGGATAGTTAGGAGGAAAATTAGATG | AAAACTTCAAGTGGGTGTTCAACAA | *Oryza sativa L.* |
| AMPL2000135 | CAGGCTCACTGTAAACAAGAGACTCTACT | AATTCAATTTTGCTTGCTGATAAGATG | *Oryza sativa L.* |
| AMPL2000136 | CCCAGCAATGATAGTTTGTAAACTAAATCA | CATTCAAAAGAAGAAGAGGACGAGGT | *Oryza sativa L.* |
| AMPL2000137 | TCTATGGCTCGTGGGCCTATAA | GACCTAGCTTGTTTCCATGGAAAC | *Oryza sativa L.* |
| AMPL2000138 | GATAATCCTGGCCTTCTACTTCTCCT | CGCATGTGCATTTTGCTATACCTC | *Oryza sativa L.* |
| AMPL2000139 | TACGACAATCTAACCTGCCCAAATA | TCCCCTAAATATGGGCATTGTAACA | *Oryza sativa L.* |
| AMPL2000141 | CTATTGGCTGGTTCTGTAAATTTGGT | CATTATCAAGATACAGGACATTGTCTTGC | *Oryza sativa L.* |
| AMPL2000142 | TCTACTCTGCCTAAACAAATCTCAACC | CCTCTAATGTTCTTTCAGTCACAAGATCAC | *Oryza sativa L.* |
| AMPL2000143 | TGGATGACGCATCAGAATGTCA | GCTGCTGATGTTATATGGAGGAAACTATC | *Oryza sativa L.* |
| AMPL2000144 | CCAGGGAGGTATATTTTGGATGGA | CCCTGTCTCAGTCCTCTGAAGGTT | *Oryza sativa L.* |
| AMPL2000145 | CCCTTCATAAGAGAGTAATTCTGGTTGA | CGATGTAGACATTACATGCTTGACCTTAC | *Oryza sativa L.* |
| AMPL2000146 | AGACTTTAACATGCACCTCATGCA | AAAGCCTTTAGGAGCATAATATAAAGCC | *Oryza sativa L.* |
| AMPL2000147 | ACATCGAATTAAGGATGCACACAA | GCCAAAATTTGAATTTTTAACGCTAA | *Oryza sativa L.* |
| AMPL2000148 | TGCTTGATGTAAGTGGCAACCAT | GTTCAGATAGTTTGATCAATTGCATGA | *Oryza sativa L.* |
| AMPL2000149 | GAGAATGCGGGAATGACTTCAC | CACAACCTTTGCATCCTTGTCA | *Oryza sativa L.* |
| AMPL2000150 | CAACGATGGCATTTGTGGGA | GCCACGCCCATTTTTAATTTTAC | *Oryza sativa L.* |
| AMPL2000151 | ACAAAAATTGGAGATGATGCGAAA | CTTTATTACCGTGATCGAGTCCAAGA | *Oryza sativa L.* |
| AMPL2000152 | AAAGCAAGCTTGTTCAGGACAGTTA | CCAAAGTAATGGTAAACGGAAGCTT | *Oryza sativa L.* |
| AMPL2000153 | CGCAGCAAAGATTCTTCAGGTT | TTCCACAGGTGAAACTCGTTGA | *Oryza sativa L.* |
| AMPL2000154 | CACTGTTTGTGCATGGCTGATT | CACACCAGAACCAGATCCCTCTT | *Oryza sativa L.* |
| AMPL2000155 | CCTTTCGAGAAAAAGGAAAATGTCA | TTTCAAGATTTGCTATTTGACATTCG | *Oryza sativa L.* |
| AMPL2000156 | ATGAAAGCGATGGAAAGAATGTG | TCGTGAGCATCCTAATCTCTCTTTC | *Oryza sativa L.* |
| AMPL2000157 | AGAACTGCTAGAGATGCACACCAA | TGGAGTGCTGAAACCTGATGTG | *Oryza sativa L.* |
| AMPL2000158 | TGGATGTAGAGGCCTGTTTAATTCA | CTTGGTTTAAGTCGTCTGAAGACCA | *Oryza sativa L.* |
| AMPL2000159 | GCACATACCGATGATGCAGAAGTAGA | GGATGGATGGGATACCTTCCCTT | *Oryza sativa L.* |
| AMPL2000160 | CCACATTAATGTTGTGATTACCAACG | CACTTTCAAACTAAACCTACCTAGCTAGCTT | *Oryza sativa L.* |
| AMPL2000161 | GGATTCAGTGGCAGCATGGTTA | AACCTCTGAGAGACATGACACTAAACC | *Oryza sativa L.* |
| AMPL2000162 | GGACCAAGCAACTTTTGCTTCA | ACATGACCATCTTCTGAAACTACCCT | *Oryza sativa L.* |
| AMPL2000163 | TTATACTTTACAGCTGTCATGTGAACAAGAA | GGCCTTGATGACCATAACCTTTG | *Oryza sativa L.* |
| AMPL2000165 | GACCAAATCGGGATTAGTCATGAAC | GACACATTTCCGTCTTGTTTCATCT | *Oryza sativa L.* |
| AMPL2000166 | TCAGGGACAAACTTGATGAGACCT | TGCTTCAGGAGAGAAGTTGATGATG | *Oryza sativa L.* |
| AMPL2000168 | GATATCCACTGCATTGTAAGGGAGA | GTCAGCTCTTTTAATACCTAAATGGGTTT | *Oryza sativa L.* |
| AMPL2000169 | GCTTTGACGATGCAAAAACGA | GGCCCTAATTATACGTTATACGTTAATTGG | *Oryza sativa L.* |
| AMPL2000170 | TGGCACAATTATATTTCCATTGCATT | GACCACCTCATTACACTAATCCCATT | *Oryza sativa L.* |
| AMPL2000171 | TGATCAATCAACTTGCTGTATACCACA | GGGAAAAATCCAAATAAGAACCAAA | *Oryza sativa L.* |
| AMPL2000172 | ACAGTATATGTGCTCTAGAAACATCGAAAC | AACAGAAAGCAACTTCAGTTAAAAGACAC | *Oryza sativa L.* |
| AMPL2000173 | CACTGTCATCTGCTTATCCAGCTGT | ACATAGGATCTGAGCAGAAAAAGAAGG | *Oryza sativa L.* |
| AMPL2000174 | CCCTATTCCGGTCGATCGAT | CATCATCGTGAAAGGTGTAGTAGCTCT | *Oryza sativa L.* |
| AMPL2000175 | GGATGTACTTTCAAATTAGGTCGAGTGT | GGTCGCCCTCATGGTAGAACTG | *Oryza sativa L.* |
| AMPL2000176 | CCCGCCTCAAGATGCATGG | CAAATTCAAGTATAGCCTAGTCCCGA | *Oryza sativa L.* |
| AMPL2000177 | CGAAGAACTTATTTGGACAAATATGACCA | GGAACATTTGGAAAACCATGAGAAA | *Oryza sativa L.* |
| AMPL2000178 | TCTGTATAGCCTCAGACTGCCTTGA | AGCGGACAATTTATTTATTACTGTCCA | *Oryza sativa L.* |
| AMPL2000179 | TCAAAATGGTAGAGAAGACTAGCTAATGCT | ACCCTCGTAAATGTCGTGAGACA | *Oryza sativa L.* |
| AMPL2000180 | TTTTTCCCTAGTAGGTGTAGATATCTAGTAGCTC | CATAGATATCCCATTGTATGTCAACAAGA | *Oryza sativa L.* |
| AMPL2000181 | CGGCATCACGGACATCTACAACT | CGAAAGTTGACACCGTAGGAGAA | *Oryza sativa L.* |
| AMPL2000182 | TGGAATTAAGAGGAAAGAGAGGAATCAC | TTCCACGCGGATTGCTTTT | *Oryza sativa L.* |
| AMPL2000183 | AAGAGGGATCGATCTACCCAGCTA | TGGTGATCAGGTAGTTGTTGTTGC | *Oryza sativa L.* |
| AMPL2000184 | CCCACGTTGATTGAGTAGATGGA | CGTAACATGCTATTTCTTTGGAATAGAGGT | *Oryza sativa L.* |
| AMPL2000185 | AAAAACTTGCACGCTCAATATCGT | TCATAGTTGTGCTTAGTGGACCTACAACT | *Oryza sativa L.* |
| AMPL2000186 | TGCATCCGCTGATTGTATCTCC | CAAGTAATCGCTCGACAGCTTCATC | *Oryza sativa L.* |
| AMPL2000187 | TGGAGGAAGTATCTCATGGGTATTG | CCACATCATCAAACAAATATTAACCCTTC | *Oryza sativa L.* |
| AMPL2000188 | CAAATATATCTCTGGCGCTTGGAA | CCTGCTGTTCTTTCTTCAGTTCGT | *Oryza sativa L.* |
| AMPL2000189 | AATAACGACATAGACTTCTGAGTGACATTTC | TGGTCACAAATACTAGTTGCTTTATGCATAC | *Oryza sativa L.* |
| AMPL2000190 | GGTAAAGAACATACCCATATCCACGT | TGCACCAAGAGATGGCAATG | *Oryza sativa L.* |
| AMPL2000191 | CGACGGCATTGCACACGAA | TGGCGTAGCAAAAACAATGTCAC | *Oryza sativa L.* |
| AMPL2000192 | AATAGACCAGCTAGAAATCACAACTATGC | GTCTTGCATGAAAATAGTTGCAACTC | *Oryza sativa L.* |
| AMPL2000193 | GCTCATCTGATCCGTATACTCCGA | GAAAGACCGAAAGAAGAGGAGGAT | *Oryza sativa L.* |
| AMPL2000194 | GTCCAGTACTAACCACTCAGATTTCAGA | GGCCGGTTCATACACGAGG | *Oryza sativa L.* |
| AMPL2000195 | TTCTGATTATGACAAAGGTGGCTATG | GGAGCTGGTTTGAAGTGGATTAATTAT | *Oryza sativa L.* |
| AMPL2000196 | CGTGGAGATAAAATACAATGGCCC | CACCAATAGTCGGATTCCTGCTATA | *Oryza sativa L.* |
| AMPL2000198 | ATAGTCTGCTATTGTACCTGCTCTTAGAAAG | CAGTTTGCCCAATTTTCAAATTCTTA | *Oryza sativa L.* |
| AMPL2000199 | CATGTCAGCAACACCAGACTTCA | CACTTACTGGTGCATGTGTCGTTG | *Oryza sativa L.* |
| AMPL2000200 | GCAGAAGTTCTTCGAGCATAACG | GAGGATGAACATTTCTCTCGCAA | *Oryza sativa L.* |
| AMPL2000201 | TCATCTCTCTCTCCAACTGATGTCT | TGACAGACGAGGTTGGCAACT | *Oryza sativa L.* |
| AMPL2000202 | TCCTTCTAGATGAAGGACCTGAAGATT | GCACGGGTATGGAATCTATACATGG | *Oryza sativa L.* |
| AMPL2000203 | GAGTGTGGTCAAACCTTGAATTGA | AAAAAAACAGGTACTAGTAGTATGTCATGTCCT | *Oryza sativa L.* |
| AMPL2000204 | CCTTCCTCGTCCTCTTCCTCTTC | GGGTGGACGAATCTGGAATGTC | *Oryza sativa L.* |
| AMPL2000205 | GTGTAAATGACTCACAGATGGTTTGG | CAACACGTGACACACACACATACATT | *Oryza sativa L.* |
| AMPL2000206 | CATTCAAACCGTACAATTGAAGAGG | TGACCTGTGTTTGTCGAAAATGTT | *Oryza sativa L.* |
| AMPL2000207 | CCCACAAGATTCAGTACTGTGCAGA | ACTGTTCTCACCTGGCTGATATCC | *Oryza sativa L.* |
| AMPL2000208 | GCTTACACAAAATATGAACTCTGTGGTGA | TTGGCTAATATTGCACGATTTTAGTG | *Oryza sativa L.* |

| | | | |
|---|---|---|---|
| AMPL2000209 | CACCTGGGTGAGGATCCTCTAA | GAACGATTACCTTGTTTTACAGTTTGG | *Oryza sativa L.* |
| AMPL2000210 | CAACTACTTGTTGGTCATTGTTTGTCA | AGAAGGCATCCAGGAAACTTATCAT | *Oryza sativa L.* |
| AMPL2000211 | CATGACCATATTTAAGGGCCTTTTG | CTCTTGCACTAGAGATGAGGCTTAGAG | *Oryza sativa L.* |
| AMPL2000212 | TTGCTAGGGTCGTTTTCGAGAAC | TTTATTAGAAGTGTATTCACGGTCCTAACAA | *Oryza sativa L.* |
| AMPL2000213 | TGTACATGCGGGACGATGAAG | CACAATCCAATTTTGCTAGCTAAGATG | *Oryza sativa L.* |
| AMPL2000214 | GGGAGTGAAATGCCCTGAAAA | CAAAGATTGTTAATGTTGAGTACCTGTGAT | *Oryza sativa L.* |
| AMPL2000215 | GACCTTGCATATTTACGAAGTGATCA | GGCTACTTTCCTTGCTTTGGTCA | *Oryza sativa L.* |
| AMPL2000216 | TTAGGCATGGGATGCAAGGAT | CTGAATCAAATTGATGTCGCTGAA | *Oryza sativa L.* |
| AMPL2000217 | CGAAAAATATTTACCAATGAGTAGTGCAGT | CACTAGTACTGTGTTCTTCGCCTACGTT | *Oryza sativa L.* |
| AMPL2000218 | CACTGCACTGGAGTAAGCCAAAC | CCAGATAAATTTCTTCTCCCACAAAGAC | *Oryza sativa L.* |
| AMPL2000219 | CGAAGGAGTGGTTCGAATCTGA | CACTAGAATTGTGTCTGACAACTAACAACC | *Oryza sativa L.* |
| AMPL2000220 | AAATTTTATGTCCTAAGTGCAGAAGAGTG | CCTTGTTGCAAAATTGTTCTCCA | *Oryza sativa L.* |
| AMPL2000222 | TTTACATCTCTAGTCTCTTATGTTGGACACA | CCAACTTGTCATTATAGTCTTGAGTGCAT | *Oryza sativa L.* |
| AMPL2000223 | ACGCTCTGGTCAGATTCTGGTTAGT | CCACTATAGGTAGCACTTGTTGCTTCAG | *Oryza sativa L.* |
| AMPL2000224 | TTCAAGACTGAGAACATTTCAACTGAA | TGGGTATGTACAGATTCAGATTCAATTC | *Oryza sativa L.* |
| AMPL2000225 | GACCTTAAACCTTAACTCCACTGTCTTCA | TCTGAAATATCGATGCTTTTCACCA | *Oryza sativa L.* |
| AMPL2000226 | TTGTATGACATAGCGCATTCCTCTA | AACCCCAGATCGGTCATATGAATAG | *Oryza sativa L.* |
| AMPL2000228 | CGGATCGGCAGTCGTGTCTAT | GGCCGTGATCTCCTCACACTCT | *Oryza sativa L.* |
| AMPL2000229 | TGCTAGATTCGATGAAACTACTCTGGT | TGGTCAATATCGTTGCCACGTT | *Oryza sativa L.* |
| AMPL2000230 | ATGAGCCAATCCTTTGATTACCC | GCGAAACTTCCTCCATACTCCATA | *Oryza sativa L.* |
| AMPL2000231 | ACTTAACTTAGTGGGTGTTGATCGA | TAACAACATGTGGAGTGTTTTCTGC | *Oryza sativa L.* |
| AMPL2000232 | CTATTTACTAAGAGAAGATGCCTCTACAATCTGT | AATGCTTTTCCCTTTACAATGAAGTACC | *Oryza sativa L.* |
| AMPL2000233 | CTATTTTGGTGAGCAGCGACAA | CTACCATGTTTTGTTTGGGAATTTTT | *Oryza sativa L.* |
| AMPL2000234 | TTTTTTACGGAGGGTGTACTCTTTAAGA | CGGTATTATCACCAACTCATTACCTACC | *Oryza sativa L.* |
| AMPL2000235 | GCATGCAATGTAGGATATAATGACCA | CGAGCTTTTAAATATCTCGTTGCAA | *Oryza sativa L.* |
| AMPL2000236 | GACAGTTGGGACCATGTATTAATAGTGTAC | TTGTTTTTAAAAGAACCGGCACATAT | *Oryza sativa L.* |
| AMPL2000237 | TTTGCATCATGTGGTGACAAATTAA | CTTTGTGCTACATGAAAGAGGATAATACAC | *Oryza sativa L.* |
| AMPL2000238 | GAATGAAGACGTGTGGTGTCTAGGA | CACAAGTGCATTACCCATAGAAATTTATT | *Oryza sativa L.* |
| AMPL2000239 | AGGACCGGCTCAACATGGA | CATGGAGTCCACATGTAAGGGAGA | *Oryza sativa L.* |
| AMPL2000240 | GTATTCTTGGAGAGCAGCACCATT | ACACTCGTCATATATTGCAGAATTCTACA | *Oryza sativa L.* |
| AMPL2000241 | GCAAGCCTGCAACCTGTAAACAG | TTTCCAACTCTACATGCTCTTCCATA | *Oryza sativa L.* |
| AMPL2000242 | GTACGGATCTGCAGCTGTTGTTT | AAAACGAGGTGTGTTTAGTTCCAAAG | *Oryza sativa L.* |
| AMPL2000243 | CGGAAGAAAACGCCAGTAAAAATA | TTGTGCGAACTTTGCATCACTC | *Oryza sativa L.* |
| AMPL2000244 | CGTATACTCAATTGCGTTGAGCTATG | CGGATGGTAGGGATTAGCAAATCC | *Oryza sativa L.* |
| AMPL2000245 | GGATTAGGTAACCCAAGACAGGGTAGT | TGTTCACATGGCAAACAGCATC | *Oryza sativa L.* |
| AMPL2000246 | GGATCATGTGAACTCTGGGAGGTATC | CAAGTCAAGTTTCAGAATTGTGTATGTAGCT | *Oryza sativa L.* |
| AMPL2000247 | AAAGTCTTACGTTGTAAAACGGATGAA | TCAAATTTGACGGTAGAGCTAACACA | *Oryza sativa L.* |
| AMPL2000248 | TCATAGCTAGGTGTCCTATTTGATTGGT | CTTGATTTTGCAGAGGAATTTGAAAT | *Oryza sativa L.* |
| AMPL2000249 | CATTGACGCCATCCCATCAT | GAAGGGTCAATTTGTCAAAATTATGG | *Oryza sativa L.* |
| AMPL2000250 | GGGTCAACCAGAGTTGTTGGTCA | CTTTGGTACGTATTCGTTCCGTATTT | *Oryza sativa L.* |
| AMPL2000251 | GGTGAGGTGAGATCGAGGAAGA | GGAGTTGAAGGCATTTTTAAAGTCCTAA | *Oryza sativa L.* |
| AMPL2000252 | AAACAACAGAAAGAACCAAGTCTGTCA | GCTATAGAACCATCCTTTACTCCTCTTTTGT | *Oryza sativa L.* |
| AMPL2000253 | TGAGTAGATCTAATGGGCGAGTGTT | GGATTGTTATATGGCATTCGAATGA | *Oryza sativa L.* |
| AMPL2000254 | TGGGATAGAGGAATCACCGAACT | TGTGCATACGCACATTGTTGTC | *Oryza sativa L.* |
| AMPL2000255 | GTGAATCTAAACACCAACTAGCAAA | TTATATAGCCACCTGTAGCATGACC | *Oryza sativa L.* |
| AMPL2000256 | GCGGTTTTGGAAACCCTGACTA | GGTGCAATAATTGAGGCTTCAACT | *Oryza sativa L.* |
| AMPL2000257 | CACGTCAGCCAAAACCATTTCT | CTCTCTCGACGGATTTTTGCA | *Oryza sativa L.* |
| AMPL2000258 | GACTTCATCATCAGTGAGACCATCA | TCATCAACCCAAATTGTGCAAAT | *Oryza sativa L.* |
| AMPL2000259 | AGCTATCCCATTTCTTTCAAAATGC | CAAAGCGGCAAAAATATACCGA | *Oryza sativa L.* |
| AMPL2000260 | TGCCGAATCCTGAACCTAACC | GGACACTATGGTACATCTGTGTCAAAAA | *Oryza sativa L.* |
| AMPL2000261 | AGTCTTCCGCGACTCAGATCATA | GGTTGCTGGTGAAGTGCTGATTA | *Oryza sativa L.* |
| AMPL2000262 | ACGGTTCCAACATTATTTGGTAATACA | TCTAATACTCCCGTCATCCCAAAATA | *Oryza sativa L.* |
| AMPL2000263 | AAACAGTTGCAACGTCACCAAAC | TGTTTTTGCCTTGCGTGCTAAG | *Oryza sativa L.* |
| AMPL2000264 | GCCAGTAACTTTTGTGACATCCATGTA | AATGTTAAGGCGTGACACCTGAA | *Oryza sativa L.* |
| AMPL2000265 | CGCCAGCCGCTGCTATATTT | GCCGCTCAGTTTGACTTTACAGA | *Oryza sativa L.* |
| AMPL2000266 | CCGGTTACCCGATGATGAACAT | GGAGAAGCACTGGCTCTTGATG | *Oryza sativa L.* |
| AMPL2000267 | GTGGCCCACCTAACCTGTTTTTA | TGGAGAGAGAAAGGAGTCACAAATAAGTTAA | *Oryza sativa L.* |
| AMPL2000268 | GGAATTTATAAAATCTCCCTGAAGCATG | AACTTTCATAGAGAAGGCTGGTTTATGTT | *Oryza sativa L.* |
| AMPL2000269 | CGGCGAATGAGGATGGTGAT | GCAACCGCAACGTACAACCTCT | *Oryza sativa L.* |
| AMPL2000272 | CATGCGTCACCATCACCGTT | AATATATATAGTGTTTTTCAATATTGGCACCTAGTATAT | *Oryza sativa L.* |
| AMPL2000273 | TGGCGCATCACCAGATGATAT | CAAAATGCTTATCCAGTGTCAATTTCTG | *Oryza sativa L.* |
| AMPL2000274 | CCTGAGCTTGCATACTCCCAAT | GCTCTGTCACTGTGTGTGACGAT | *Oryza sativa L.* |
| AMPL2000275 | GGCCTTAGAGGAAGAGATATCGCT | GATGGTCCCATCCCATCCATA | *Oryza sativa L.* |
| AMPL2000277 | CCGGATACACACGAGACGACTACT | GCCAAGATAACTGTGCACGACAT | *Oryza sativa L.* |
| AMPL2000278 | CACGCTCGCATCATTTTAAAAAG | CAAGGTAGCTTTTCTCTCAGTCCGTT | *Oryza sativa L.* |
| AMPL2000279 | TGGCAGAGGAAGACTGGCAA | GCATGCACCAGCCAATCAG | *Oryza sativa L.* |
| AMPL2000280 | GCAGATGCATGCATCTTTACTCC | CCAAATTTTGGCCTATATGGAAATG | *Oryza sativa L.* |
| AMPL2000281 | TGAAAACCAAGGGCTAGGGCTAA | GATCTGGATGAATTTTTGAACAAGGTT | *Oryza sativa L.* |
| AMPL2000282 | CATCGTTGTCAAAACCAAGACGA | GATCTTCTCGGGATTTGCCTCTATC | *Oryza sativa L.* |
| AMPL2000283 | CTAAACAAGGGCTTTATTAGTTGAAATTAAAA | TGTCTACGTTTATCAACTCTGAGCTTCAT | *Oryza sativa L.* |
| AMPL2000284 | CGAAACACGAGGCGTGAGAGTAT | GTGTTCGATGGCCTCATTGAA | *Oryza sativa L.* |
| AMPL2000285 | GCGCTGTCCAAGAGCTTCTC | TTTATGAAGCAATCGATTGCCA | *Oryza sativa L.* |
| AMPL2000286 | TGTAGCAACTTCTATTGTGTTTTTCAGG | CGAAGTGTTTTTGAGAGGCTAAGTTT | *Oryza sativa L.* |
| AMPL2000287 | TTTCGCTTTTACTTAGGCCTTGTTT | AGCCTAAGGTTTCGAACATTCTAGA | *Oryza sativa L.* |
| AMPL2000288 | CCCTCTGTTGCCAAAAAGATCAG | TCACCTTTCTGTAGAGATGCCCAT | *Oryza sativa L.* |
| AMPL2000289 | GTGGTTAAGGTCTCTGGCGAAA | CCACTTCGGCAGCAGATAACATT | *Oryza sativa L.* |
| AMPL2000290 | GGAATCCATTCATCAATCATGCAG | CTCCACAATGATAGTCTGATCTCTGATG | *Oryza sativa L.* |
| AMPL2000291 | GGAAATACATATCCGTGTTTGAGTCAGA | TGAGAGAGAGAGAGATAACAATTGGTGTAA | *Oryza sativa L.* |
| AMPL2000292 | AGAAGGTGCTTCAAGTGCATGAC | CCAAGATTACCAGTCAAATATCATCCTG | *Oryza sativa L.* |
| AMPL2000293 | GCACCTGATATGTAAAGTTATTTTCTTGAGAC | GCATTGTCTAGAGACATTAGATTGTACCACA | *Oryza sativa L.* |
| AMPL2000294 | CAAGACCAGCGTGTTCCTTTTT | TGTCCTTACAATCATATCACTTTTGGGT | *Oryza sativa L.* |

| | | | |
|---|---|---|---|
| AMPL2000295 | GCAGTACATGCCGAGCTTACCAT | TCTGATTCCTGGCGATAGGAATAC | *Oryza sativa L.* |
| AMPL2000296 | ACATAAGCAAAGCTTGGCATCACT | CATCTATAGTCATTGTTGCTTGCCTGTT | *Oryza sativa L.* |
| AMPL2000297 | CACACACCCGGTCTCCAATT | CACGTCCACGTACGTTGTTGAG | *Oryza sativa L.* |
| AMPL2000298 | CACCTCTACCATGAATCATCCTCTGT | GATATGTCACCACAAGTAACAAGTAATGAGC | *Oryza sativa L.* |
| AMPL2000300 | TCCAAGTTTAGCAAACCTTCTGACA | CATTCAGTACGACCATGCTAATTATGATT | *Oryza sativa L.* |
| AMPL2000302 | GCAGCACGATCAGCATACTCAG | TTCACAGGTCCTCACCAAATCC | *Oryza sativa L.* |
| AMPL2000303 | CGCTCTGTTTCCGGAACAATT | GGCAAATATGCCGAATTATTGG | *Oryza sativa L.* |
| AMPL2000304 | GCCCAGAAAGAGAAACGTGTGT | CCACATCCTACATTGACTGTTAAACCA | *Oryza sativa L.* |
| AMPL2000305 | AGTCGACGGAGAGGTGCGA | AAATTATTCCGCCAAAAATCCG | *Oryza sativa L.* |
| AMPL2000306 | GGGCTGTCTGCAATCTAAAGAGG | GGGCCTATTAGTTTATTAGAGAGCCACAT | *Oryza sativa L.* |
| AMPL2000307 | TGGTTGTTGTTGTAGCAAAGAGGA | TCCTAGATAAGCAGCGAAGGAAATT | *Oryza sativa L.* |
| AMPL2000308 | AGAAAACTTCGGTAATAGCCACCTTT | CCAATGCTCACTGAGACTGACTATTTG | *Oryza sativa L.* |
| AMPL2000309 | TTCAGAAACATTCCCTGACGCTAT | TGCTGATGATAAAGGAGAAGATTTCAA | *Oryza sativa L.* |
| AMPL2000311 | TGAACACGGAAATACAAAACTGCAC | AACAAAAGCACCTAGCTTTATCCTATCC | *Oryza sativa L.* |
| AMPL2000312 | TGCAACCTCATTTCTACTTCTTTCATG | CGAAGAGTGATGTCTATAGTTTTGGAGTTA | *Oryza sativa L.* |
| AMPL2000313 | CCATTTTCTAGGTAGAAGATGTTACATGC | AGGCCTCCAATAGAGTTTTCCCTA | *Oryza sativa L.* |
| AMPL2000314 | GGGTCTTCTTTACCTTCCCAGTT | GTAATGTTCCGGTCATTCTCTTGTC | *Oryza sativa L.* |
| AMPL2000315 | AGCTAGAGCTGTGCCAAACAGACT | GTCTCGGTTGGTGTTGGTTAGACT | *Oryza sativa L.* |
| AMPL2000316 | GGTTGAGGACATTGCACAAATCTTA | TATTGCCCAAGTCACATTCAAACTC | *Oryza sativa L.* |
| AMPL2000319 | TGAAAACCCGCTTAATTTTGTCA | GGGCTCTAACTGGTGTGAATTACTACTAGTAC | *Oryza sativa L.* |
| AMPL2000320 | GGATTGCAAACCAGCCCAAA | AGGCACTTAAGATTTGTCCCAAAATA | *Oryza sativa L.* |
| AMPL2000321 | TGCTATATGCTTAACACCAACAACC | GAAGAAACGTGCATTATTACATCGC | *Oryza sativa L.* |
| AMPL2000323 | CATTTGCAATGATGCAAACTAAAACA | CTCGACAATATCAAAGGTACCGAGAT | *Oryza sativa L.* |
| AMPL2000324 | TGAGCATCTGATTTTCGCAGAA | CGCATATGGTCAAATAGCATTGC | *Oryza sativa L.* |
| AMPL2000325 | ACACATTGGACTAGGGACAAAGTAA | AATTGAGCCTTCGGTAAAACTTGAG | *Oryza sativa L.* |
| AMPL2000326 | CCCCTCTATTTTTGTAGGTGAACTAAGAGT | GGTGATGGATCCACACCCTTAC | *Oryza sativa L.* |
| AMPL2000327 | ATGAGAGAACTACATACGCAGGATGA | TTGTGACTGACGAGGATGATATGG | *Oryza sativa L.* |
| AMPL2000328 | AATTGACAGCTGGGCCTTATTACA | CATTTGTGTTGATTTGACATTGTATAGCT | *Oryza sativa L.* |
| AMPL2000329 | CCATTCCTTTCATTATTTAGCAACCA | CAGTGTGGCTTATTACTTATTAGCTACCAA | *Oryza sativa L.* |
| AMPL2000330 | GACGGCGAGAGAGAGATACAATG | GAGCCCAACCTTGAAATCTTAACAA | *Oryza sativa L.* |
| AMPL2000331 | GACGGAGGGTGTAGTTTTCGATC | GCTAGGTAGCCAGATCAGTTTCGTTC | *Oryza sativa L.* |
| AMPL2000332 | GAAACCTGTGGAACAAAGCAACA | GCTGGTTGAACCTGTGAGAATCC | *Oryza sativa L.* |
| AMPL2000333 | CTTAAGAGAAGCCATCCCCATCAC | TTCACGCAACCCAAATACCAAATTA | *Oryza sativa L.* |
| AMPL2000335 | CAGATGATATATTCCCTTTGACAATGG | TGAATGATCCAATGGCTCTAATTAGCT | *Oryza sativa L.* |
| AMPL2000336 | CCATGTCCTCCAACCAAACAA | AACTATGTGCAAATGTTGTAAATGCAA | *Oryza sativa L.* |
| AMPL2000337 | AAATTCTAGATACGCCACTAGGAGATCA | ACCTGCCATGGAAGCTCGT | *Oryza sativa L.* |
| AMPL2000338 | CAAATCTAAGTCATACCAAATAGCAAGCA | AAAAAGCTCTTTCCAGGCGATT | *Oryza sativa L.* |
| AMPL2000339 | CCGATGAAGCGGAAGAACTG | AAACCGACATTTCAGACAAATGGTA | *Oryza sativa L.* |
| AMPL2000340 | GAGGTCATCGAAGCTGTACGTC | TTATCTTGATGTGATTTCGGTCGTG | *Oryza sativa L.* |
| AMPL2000341 | CGCTGAATCTTCTTCCAGGCTT | TGAAATACAGGGTAGGGCAGATTATT | *Oryza sativa L.* |
| AMPL2000342 | CAAAAGCTTCTCTCCTCTCTCTTTTCTCT | CGTTACTGTGAGTGAGTCAACCGTTT | *Oryza sativa L.* |
| AMPL2000343 | AAGAGCCAAGATCACCACATAACC | TCAAAATTCCCAAACTCATTTTCAG | *Oryza sativa L.* |
| AMPL2000344 | GGATGACCTGCATATGCCACAT | GCATGCGCAAGCATATAAACTTC | *Oryza sativa L.* |
| AMPL2000345 | TTGGGATTCACACGAGTGGAT | TTTTAAACCACGGTATACCTTATGTGC | *Oryza sativa L.* |
| AMPL2000346 | AGTTGTCTGCACATTTCTTCCCA | TGATTGTAACAACCAGATTAGATGGAGTA | *Oryza sativa L.* |
| AMPL2000347 | TCAAGTTAAATGAGAAGCGGGAAA | TGTGAGCATTGAGAATAGTTGAAGTTTG | *Oryza sativa L.* |
| AMPL2000349 | TTTTACTCTGCTATGTATGGCTCCA | CCATCAAATCGACTTTACTCGTGAG | *Oryza sativa L.* |
| AMPL2000350 | TCGTCAGCGCTAGAGGGTTTT | TGTGAGGGACCTTCCCTAATGC | *Oryza sativa L.* |
| AMPL2000351 | TTTCATTGTGTCCCTTCTCCAGA | CATCCTATATGTCAAAAACCACGGTAAA | *Oryza sativa L.* |
| AMPL2000352 | TGAGAAAGAGGAAAGGGAGAGGATA | ACAATTGCTTCTTTGACCATGTTCT | *Oryza sativa L.* |
| AMPL2000353 | GTTGTTCACAACTGACCCACTTACA | CCTTGTCATCGGTGACAATGTAGC | *Oryza sativa L.* |
| AMPL2000354 | CGACGTGCAATTAGGAAGCTTCAT | TCTATGCATAGGAGTGACTGAATCAGAG | *Oryza sativa L.* |
| AMPL2000355 | TGTTGCATGCATGTTTAAATTCTCC | GGTCCATCTACCAAAAGAACATATTTCAT | *Oryza sativa L.* |
| AMPL2000356 | CGTCTTTCGGCCTTTCTCACTT | GGGAGGGCAAAAATTACATTTAGC | *Oryza sativa L.* |
| AMPL2000357 | CGGCAAAACAAATGCAACTG | GAAAACCTATGCTATGTGAGCTCAGATT | *Oryza sativa L.* |
| AMPL2000358 | GATGGAGGTAGTAGCAAATGCAAAG | CCCGTTAGACCTAATGGAAGTAATCG | *Oryza sativa L.* |
| AMPL2000359 | TGCCACCTCTCCCTCTTGTATACA | TGGGCAACTGTACTCAATTTATTTCAC | *Oryza sativa L.* |
| AMPL2000360 | CGTAACTTGTAGCAAATTGCTCCA | TGCATTTTCCACGCTGGATTT | *Oryza sativa L.* |
| AMPL2000362 | GATTTGGAGCAAACGAAAAGAGGA | CCATTTAGCAGCGTCTTAAACCTTC | *Oryza sativa L.* |
| AMPL2000363 | TGTGTAACTTTTATAGTGTGCCTGCAAT | CCGACAGTTGGGTAAAATAGACTTATTTT | *Oryza sativa L.* |
| AMPL2000365 | GCATTCTAGCACAATAACTTGGCA | TTCTACATCCAGTTTAGTTGACCTTCAG | *Oryza sativa L.* |
| AMPL2000366 | CATTTCCAGCAGAAGGGCAGTA | TGATGGTTTGGTTAATCTTAGCCATT | *Oryza sativa L.* |
| AMPL2000367 | CTCAATGGGAGGAGCGTAGAAA | TGATGTTATATTTAGACGAGCTTCTCGAA | *Oryza sativa L.* |
| AMPL2000368 | TTGCCTCTTTCTCGTTAGTAAATGGT | TTATTTTAGAAGCCATAGCCACCAATA | *Oryza sativa L.* |
| AMPL2000369 | ACTACTACTCAACAACCAATGAACCAAA | GAGAGTAGTACTACGGCAGTGTTTAGTTCC | *Oryza sativa L.* |
| AMPL2000370 | GAGGATACAATTCAGGAAGCACAAG | TTGGTCTTGGTGTCAAAAGTAAAAGA | *Oryza sativa L.* |
| AMPL2000371 | TGGCTACTTTAGTTTCCTATCTGTCGA | CTAGCGGTTTGCACATAATGACC | *Oryza sativa L.* |
| AMPL2000372 | TTCTGAGAAGGGTATAACTCAGAAAGTACAC | TCGATCTCTGGGATCGCAA | *Oryza sativa L.* |
| AMPL2000373 | CGGTTTGCGACGATATGGATA | CATTGATAAACCAAAAGAAAACCGA | *Oryza sativa L.* |
| AMPL2000374 | TGTCCACCAGCTACGTAGCTAAGTATC | TGAAGCACGTCATGGAGAAAAATATAT | *Oryza sativa L.* |
| AMPL2000375 | GGATGCTCCAGAGCTGAAGGTAC | CAACATCCCGGTTTCCTTGAAC | *Oryza sativa L.* |
| AMPL2000376 | CAAGTCACCTTTCTCCATTGCA | CATTCTCCTTGGTGTTAACTTAGTCAGTAA | *Oryza sativa L.* |
| AMPL2000378 | CCTGGGAAGGAAGATGCAAGTG | GAAATCCTACAGCACCACACAGAAA | *Oryza sativa L.* |
| AMPL2000379 | TCGGCTTACATCTGGTCAGCTAGT | TCAACAAGGAGAACACATAACAAGAAA | *Oryza sativa L.* |
| AMPL2000380 | CAGCTCCCGGACTTATTTTTAATAATTCT | TGTTATTCTTTGCTTGGAAGACAATTGTA | *Oryza sativa L.* |
| AMPL2000381 | GTTGCACTGATGATCCCAGCTA | GGATTTCCCAGTTTCCAACTGAA | *Oryza sativa L.* |
| AMPL2000382 | GGGCTACTTCCTCTCTAAATGCG | AAAGTGATTGAGACGAGGGAGTTTT | *Oryza sativa L.* |
| AMPL2000383 | ATCAATGCAGCTCCTAATAAGACCT | AGATCCTAACACAATGAGGGGAAAA | *Oryza sativa L.* |
| AMPL2000384 | AATTAACATCAGTGGTGTATGTTTGCTC | GCAACGAATACTGATATGATATGGCTAC | *Oryza sativa L.* |
| AMPL2000385 | GGTTTTGTTTTGCAACCATCCA | GCTAACGGTGGCAAATTGTTTG | *Oryza sativa L.* |
| AMPL2000386 | GACCATGTAATGGCTGCAACAA | TCATGCATCATCATCTTGCTAACAA | *Oryza sativa L.* |

| | | | |
|---|---|---|---|
| AMPL2000388 | AAAAGGAGATGTGCCTATAAGTATGTCCT | AGCTACTGGCCAATTTTGTTTCAC | *Oryza sativa L.* |
| AMPL2000389 | TGTCATTGCTTTTTGTGCACCTA | CGTCCCATTTAATGATACAGGTACATTG | *Oryza sativa L.* |
| AMPL2000390 | TTTGGAATGGAACTTGCAGTAAGAC | TAAAATCAGCCACAAAAGGGTTCTG | *Oryza sativa L.* |
| AMPL2000391 | TCGCTTCTCTCCTAGAACAACTACTTTG | GGTAAACCCTGGTACTATCTCCGTTCT | *Oryza sativa L.* |
| AMPL2000392 | TTATTCCATCCTCCTAGTACTATGTCTGTAGA | GAAAGTTGAGGAAAGAGCTCAGTCTTAA | *Oryza sativa L.* |
| AMPL2000393 | GGCAGTCGGGCTATCTATGAGCT | GGAGAAGGAGAAAAGGAAATTGACAT | *Oryza sativa L.* |
| AMPL2000394 | CCAGCAGCCCAAGAACTTGT | CCCACGTTTTGCTGACTCTTTC | *Oryza sativa L.* |
| AMPL2000396 | GCTGTGTGCACATGTGTAGTACCA | CCTCATGTGCCATGCACCATAT | *Oryza sativa L.* |
| AMPL2000399 | TGCCAACCTTTTCGCCATTT | CGCTTTCCGATTATTTGTTGTTTCC | *Oryza sativa L.* |
| AMPL2000400 | TCACTTCACATAGCACATTGCACATA | GTGGCTTCATTGATGAAGCTACCT | *Oryza sativa L.* |
| AMPL2000401 | TTAGAGTTCAGTCTCACATTGGTGGT | AGGCACCACAAGCACTAGAACTATATTT | *Oryza sativa L.* |
| AMPL2000403 | GGTTGGTCATAAGACTGTCACGGTA | GGGTATATTCACTGGAATTGATAGCATG | *Oryza sativa L.* |
| AMPL2000404 | CATGTTCCCGACTGCAAAGTTC | GCTCACAGTTACAATGATCTGCAAT | *Oryza sativa L.* |
| AMPL2000406 | TGATTAGCTTATAGTTGCTTTTGTTCAGC | CACAGTCGTGGGAATATACGAACATT | *Oryza sativa L.* |
| AMPL2000407 | GGACAGCTGCTGCTATGAGAAAGA | CGTTGGATCGAGGATTTGTCGT | *Oryza sativa L.* |
| AMPL2000408 | GGCGAAAATAAATCTTTTGGCTGT | CCTTCAACTGAATTGATTGGCAAATAG | *Oryza sativa L.* |
| AMPL2000409 | TCACTTACCCAGAATCCAGCAGA | GGCAAACTTATTAGATTCATCAGGTTG | *Oryza sativa L.* |
| AMPL2000410 | CACTAAAAATGGAGGTAGTATCTCTGACCT | CGTACTAGCTTGGTTTTTACGGAATAAA | *Oryza sativa L.* |
| AMPL2000411 | CAGCTCAGAGCTATAAACTGTAGTACAGTACG | TGCCATTGTATTCCTCTAGTATTCCTATGT | *Oryza sativa L.* |
| AMPL2000412 | CACTGGGATTGTGGACTATAAAGTTACCT | TTTAGTAGGAGGGACCTGCCAAA | *Oryza sativa L.* |
| AMPL2000414 | CAGCCACAAGACTTAGTAGATGCAGA | CAGTGACCGACGAGGTGCTCTA | *Oryza sativa L.* |
| AMPL2000415 | AGTATAAAGCCACATCATCGCCTT | AATACACAGACCCAAAGAAACTAGACATC | *Oryza sativa L.* |
| AMPL2000416 | CCTCCCGACCAAGAAGAACCTAA | TGCTCAATAGGAACATGGTCCAT | *Oryza sativa L.* |
| AMPL2000417 | GGACCCGAACTAGCATGTCTCTCT | TGGTGGCTTCAGGAATTTTAGTTG | *Oryza sativa L.* |
| AMPL2000420 | GGGAAAAGTTAGCAAGAGCTCTAACCT | GACAGCATTCAGATGGAGTGGAA | *Oryza sativa L.* |
| AMPL2000421 | TGGAAGTGAAGAGGCTTCTGGAT | GATGAGCCTGTACTAACGTCAGAAAAATAG | *Oryza sativa L.* |
| AMPL2000422 | TCTCTAAAGAACTCTGATGTTAAGCGTGT | TTCACGTCCCAAAATCACATCA | *Oryza sativa L.* |
| AMPL2000423 | GGGAGTCGAACCCAAATATCAGG | CGCACCTCTGCTTCACTTCATTA | *Oryza sativa L.* |
| AMPL2000425 | TTAAGTTCCCTCGATGGTTTTACCT | ACCTCCTTTCCGTTTTCATGC | *Oryza sativa L.* |
| AMPL2000426 | CATCCTCGGTTTTCTTCGGTAAG | GAGATTCTTCCATACAACAATAGGCATC | *Oryza sativa L.* |
| AMPL2000427 | CTCGTTGTGTTTCTGTAGAAAACCA | AATTGGAAGCAATTCAACATGCATG | *Oryza sativa L.* |
| AMPL2000428 | CGAAGATGAAGGTTCCAAAGACCT | TTGTAGGCGAAATACTTACGTAGATATGC | *Oryza sativa L.* |
| AMPL2000429 | TAAATACACAACATGCATGGAGAGC | GTTTGGTTTCGCCCTAAAGTTTCTA | *Oryza sativa L.* |
| AMPL2000430 | CGGTGATGGACCATCATCCAAC | GGTTTATCAGCCGTTGCTCAAA | *Oryza sativa L.* |
| AMPL2000431 | CCTAAAACCCACGTGTCAGCTCTA | CCAAACCCAAAAATAATTTAAATCACA | *Oryza sativa L.* |
| AMPL2000433 | AATAACTAGGTCATATAAGGCTTGTGTGGTT | CCAAGCTGATGATGTCCTTATCAATACT | *Oryza sativa L.* |
| AMPL2000434 | ATCCATGATTGCCAATCTGTTTTGT | TCACTTCAGCTCAGATCTCTTTTCA | *Oryza sativa L.* |
| AMPL2000435 | CTTGTAAACTTCACTGAGATCTGCTCA | CATTGCGCATGCTTCCTTTAG | *Oryza sativa L.* |
| AMPL2000436 | TCCGTTTGCTGAGGTCAACTCTA | TGTGATGTCTGATTTCCTGAAATCAAT | *Oryza sativa L.* |
| AMPL2000437 | CCCACTAAAATGGAGTTTGGTGTTG | GAGGATACGGTTTGATGCAGGA | *Oryza sativa L.* |
| AMPL2000438 | GGCATCTCTTAAAAACCGATCTTATCAA | TGCGATAATTTTTCAGATCCACAATT | *Oryza sativa L.* |
| AMPL2000439 | TGGGCAAGGGAAGGAATGTA | GGAAGTGCTAGTTCTGGGCTATGACT | *Oryza sativa L.* |
| AMPL2000441 | CACCCTGAAAACCTCAGCTGTTT | CAGAGTTGAAACTCTGACGTGATTAGTTA | *Oryza sativa L.* |
| AMPL2000442 | GGAAAGACCGTACAAATTCAAGACAAG | GCAGAAAAACCCTAGCTTGCTCAT | *Oryza sativa L.* |
| AMPL2000443 | AAGTGCTGGTGTAATTAGCTAATTAACCA | GCGTTTCCCGTCCTGTTTGT | *Oryza sativa L.* |
| AMPL2000444 | AAGATGTTCAGGGTTGTGACAAAGT | GGAATAAAGAAAAGGAAAAGGATGAGAA | *Oryza sativa L.* |
| AMPL2000445 | TCGCGACCTTCAAATGCAA | TGTCTCTGGCCAAACATACAGAGAG | *Oryza sativa L.* |
| AMPL2000447 | GAAAACCCCGGCATTGATAAACTAA | TTGAGGACACTATGGTTGAAGGG | *Oryza sativa L.* |
| AMPL2000448 | CCTCGAATCCAAGCCTAGTGATTAGT | CAACTGCCAATGTACTGCTAAGGA | *Oryza sativa L.* |
| AMPL2000450 | GCTCAAGTTTGATGATCTTCATTTGCT | CCACAGGGAAGAGAATAATCCACAG | *Oryza sativa L.* |
| AMPL2000451 | CGCACCGCTGTAGCATAGAAAT | CGATGCTCTCGAGGGATTAATG | *Oryza sativa L.* |
| AMPL2000452 | GAGGCGCAGAGGTACCAACTC | CCTTTCCTCTACAGTAACCTCTCTGATTTC | *Oryza sativa L.* |
| AMPL2000453 | TGATCATCCATGCATGCATTC | CTGAGTCAAAAGCTTAAACCTTAGTGGT | *Oryza sativa L.* |
| AMPL2000454 | CCCCATTCCCTCTCTCTTTCTCTC | AGAGATTCACAAGCATGCACATCT | *Oryza sativa L.* |
| AMPL2000455 | GCGAGGCCAAAACTGGAATAGA | CTCGTCTCGGTGTGAGTTTGAA | *Oryza sativa L.* |
| AMPL2000456 | GGTGAATAGAGAAACCCAAAAATTCC | GCAAGACTTTTGAGGTGTTCCTAAATC | *Oryza sativa L.* |
| AMPL2000457 | TCTTCTCTGCTAAGCTTGCTAGCTTC | GCCACAGATTTATCCAGTCACATG | *Oryza sativa L.* |
| AMPL2000458 | GCGATGATGCGATGAGAATGA | AGCCCATGACTGGTGATGATAAG | *Oryza sativa L.* |
| AMPL2000459 | TGTGCTATCATCTGTGTTGCCAT | AATTCGTCAGCTTTATTTTCAATCTCC | *Oryza sativa L.* |
| AMPL2000460 | CCTCAGCTGGAAGGCAACTAGAT | GCACTAGAATGCACTTCAAGTAAGCAAC | *Oryza sativa L.* |
| AMPL2000461 | CATAGATAGTGTAGTGCCTGATGACCA | CCCTAATGGTGTAACCATATTGCTTC | *Oryza sativa L.* |
| AMPL2000462 | CCAGATACAGAACTGCGAATGATTG | CAGAAAACATGCATTATGTTTGTAGCTAGA | *Oryza sativa L.* |
| AMPL2000463 | TGAAATGCCTCATTTCAGCCA | CAATTTGGTCTTGGGTTCTAAATTCAG | *Oryza sativa L.* |
| AMPL2000464 | TCTACAAAGACATTTGAAAAACACGTG | CCCGTGTGTGCGTTCATTTATA | *Oryza sativa L.* |
| AMPL2000465 | CTTCTAGAAGCTTTTGCTTCTCTGGA | CGTTTGGCTTGGAGCCATTTATATATA | *Oryza sativa L.* |
| AMPL2000466 | CCGTTTGATTTTGTGAGACCACT | GGCCATCACAATTAATTTTTCTGAAGTT | *Oryza sativa L.* |
| AMPL2000467 | GTGACCTACAAGGCTACAAGGAATAACT | CAACCATGGATGCTAATGGGAT | *Oryza sativa L.* |
| AMPL2000468 | AACACAACGAAAGCTTTTGACGA | CTCCATCTTGATAGGCTGATAGTATCAAT | *Oryza sativa L.* |
| AMPL2000469 | TTATTTGCTATGCTAAGGTCATGTTTGT | CTTGAAACATCTTGTGACAAATCCAC | *Oryza sativa L.* |
| AMPL2000470 | TTGTCGACAAATTCAGTCCCAAGT | GCTGATGTGATTGTCTGATGAATCC | *Oryza sativa L.* |
| AMPL2000471 | CTGTATAAATCCTTGTGTCCGCATC | GCAAACTTTGGGTGCTTATTTGTTT | *Oryza sativa L.* |
| AMPL2000472 | CCAAGTCATGCATAGAAATCACATGA | TGAGTTCAGATTACAGCCTATACTGCA | *Oryza sativa L.* |
| AMPL2000473 | AGCCATATTCTAGCTCTTCCCTTAT | ACACATGGAAAAATGTTCACTCGAA | *Oryza sativa L.* |
| AMPL2000475 | GCATCATTTTTGGGATTTGGGT | CAGTGATGACCTTCTCAGTGTCATTT | *Oryza sativa L.* |
| AMPL2000476 | CATACAAGAACTTCAGCCTGACCAA | GCACACAATAGGTTTCCTAAGTATCGAC | *Oryza sativa L.* |
| AMPL2000477 | TGATACCTCGGCTAAAAACTGATCAA | ACCAGTTCCTACGTATGTTCCTCCT | *Oryza sativa L.* |
| AMPL2000478 | TTTCTTGTTGCAGTCGACTGATACA | CCAGAAAAACAAAAACTTGGACAATTC | *Oryza sativa L.* |
| AMPL2000479 | CATGTGTCATAGCAGTAAATTTGCAAG | CCAAACAGTTTTGCAGAAATAGCAA | *Oryza sativa L.* |
| AMPL2000480 | CACGTATCAAATATCCAAAGAAGTCATG | GGTGAAAATCAATTGGTCGATTTTG | *Oryza sativa L.* |
| AMPL2000481 | TCAATAAAATCAGCTGGGAGGATCA | CTCCTCTACTTCCGATGGGTAGATC | *Oryza sativa L.* |
| AMPL2000483 | TGGCAGAACAACGGACATTGT | TCCTTCGATTCCTCTCTTTCCA | *Oryza sativa L.* |

| | | | |
|---|---|---|---|
| AMPL2000484 | AACGGGACTCAAAACGAATTTCT | CAGGTAGGGCCATGAGTACTGATTT | *Oryza sativa L.* |
| AMPL2000485 | TTTGAGCTAACATGGCAGCTTCT | AAAACCACGACCTCCACCATTA | *Oryza sativa L.* |
| AMPL2000486 | GACAAGAGGGTCTACTACCTCAACCAT | CGCGACGGTGATGCGGAAT | *Oryza sativa L.* |
| AMPL2000487 | GAAAGGCTACTGTACCAAACTGTATGC | TGTGTGTAGGATTACCCATTTCATAGG | *Oryza sativa L.* |
| AMPL2000488 | GTGACTACAACTCAAATAGTCACACTATGGT | TGAGGAAACTGGAAAGGCGTTA | *Oryza sativa L.* |
| AMPL2000489 | CCCACACTCAGAGCATGGATTG | GGAAACCACTATTAGTGTTGGTTTTTGG | *Oryza sativa L.* |
| AMPL2000490 | TTGTTTTGCTCAAGCCAAGATCA | GCTGCTGCTAACTAAGCAAAGCCTT | *Oryza sativa L.* |
| AMPL2000491 | CCCCACGTTATACCCTACCATCCT | CTTCTCGTCCATACCTCACTAAAAAGC | *Oryza sativa L.* |
| AMPL2000492 | AATGGATTGTTCCTCGCTCGTAT | TTGAGGTTTGGTAGTTGTGGAATCTAA | *Oryza sativa L.* |
| AMPL2000493 | CCTGCAAAAGAGAGTCCATCTGGT | TCAAGTTGTATTGGCCAAGAGATACA | *Oryza sativa L.* |
| AMPL2000494 | TTTGCTCTCGTAACTGTCTCTGCTT | GGCTATTTGGCCTTGACAAGGTT | *Oryza sativa L.* |
| AMPL2000495 | TTTTCCTGAACCATTCATTATGCA | AAGCATTGGACCTGTTACAATTGAA | *Oryza sativa L.* |
| AMPL2000496 | CATTTGAGTGGTTAGAAGGCTTAGGA | TCACATATGCATTTATCTCTTTTCCCT | *Oryza sativa L.* |
| AMPL2000497 | AGTTGAAAGGAAAATCAGCTCTTACCT | TCAAAACATGCCTGCTCCTATGA | *Oryza sativa L.* |
| AMPL2000498 | CGACAGGTAACAGAGTCGCAAAAC | CATGTTACATGTGCGCTTTAGCCT | *Oryza sativa L.* |
| AMPL2000499 | CATAGCATGCTAGTGCTTCTTTGTCTC | CCTTGGAGTTCAGGTCGATGAAG | *Oryza sativa L.* |
| AMPL2000500 | GCCATAACCACGTTGGCAGACTA | ATGGGCTATGGAGAAGCTTTTGT | *Oryza sativa L.* |
| AMPL2000501 | CGCCTTCCTGGAGCAATATGAG | TGTGGCAAGTTCTTGATCTTTCC | *Oryza sativa L.* |
| AMPL2000502 | ACATGGAGAACGCATGTAGCAA | CGAAGCGTCGCTTCCTCTAAAT | *Oryza sativa L.* |
| AMPL2000503 | TGGAGAGATGGTTGTATATTATGCACC | CCTCTAGTGTTGACCAGGGAACAAC | *Oryza sativa L.* |
| AMPL2000504 | GACAGGGCTGGAGTGACTTCATA | TCAAGTACTACAAGGGCACTCACAGT | *Oryza sativa L.* |
| AMPL2000505 | GAACCGAACGAGCCCTTAACA | CAAAAGGCTAAAACCCTTTCTTCAA | *Oryza sativa L.* |
| AMPL2000506 | TCCACTAAAGAACCGTACCAAATTTTA | CAGATAAAACATGGAGGCTGCCTAT | *Oryza sativa L.* |
| AMPL2000508 | TTGTTATCTATATGGTTTGCATTGCCT | TTACTACTAGGTGTCTTGGGTCATGGT | *Oryza sativa L.* |
| AMPL2000509 | CCTGTGAGGTCGCAGAGGTACTCTA | CGGCTTCCATGTCGTAGATTTAA | *Oryza sativa L.* |
| AMPL2000510 | TCGTGCACTTCATGGCCAC | CTGTAGGTCTTGACCGCATATACG | *Oryza sativa L.* |
| AMPL2000511 | TGGAATATAGGTTTGTGTCCCTCTAGA | CCATAGTTAATGCTTGGATAAGCACAATT | *Oryza sativa L.* |
| AMPL2000513 | CTAGTGTTATCAAAGATATACTGCTGC | AGACTCGGCTACAGTAGGAATATTG | *Oryza sativa L.* |
| AMPL2000514 | GCATAAGGGAAGCCTTCATCTTTAC | AAATCAGCATAGAAAACTGGCCAAA | *Oryza sativa L.* |
| AMPL2000515 | CCGTTCCTAGGCAGATCTTTTATCT | TTCTGGTGTTTGGTGTGTGTTTTTA | *Oryza sativa L.* |
| AMPL2000516 | AGAAATTCAGGAAGATGGCATTTGG | CTACGGTAGTTGCAATGTCCTAAAA | *Oryza sativa L.* |
| AMPL2000517 | GTCTTATAACCTGAAACGGAGGGAA | GAATTATGACGACTCTCTCCAAGGA | *Oryza sativa L.* |
| AMPL2000518 | GAGGGAGAGGGAGAAGAGAAAAAG | ATAGCGAGATTGGAGAGGGAAC | *Oryza sativa L.* |
| AMPL2000519 | TCTACTCAGGACTTTTACCATTTGTT | ATACCATCGGACGTCCTATTTGTAG | *Oryza sativa L.* |
| AMPL2000520 | GCATCCATACATCAAGAAACCGAAT | AAAAAGACCAGTCAAATGATGCCAA | *Oryza sativa L.* |
| AMPL2000521 | CCATATGGCCATCTATAGTCACTTTG | TCCCATCAAGAAAGGGTACAATCAT | *Oryza sativa L.* |
| AMPL2000522 | TGTTTCCATTAATTAGTAGCCCTCCA | TCATTTAGATTTTCGTTAGCAGCCC | *Oryza sativa L.* |
| AMPL2000523 | ACTCCAGTAAAAGCATCAAAACCTG | TGACCATGACAGAGGTGAATAAGAG | *Oryza sativa L.* |
| AMPL2000524 | AAAAACTCCAATCTTGTCAAAGCGA | TGTTCTCCTGGTCTTTACTAACCTG | *Oryza sativa L.* |
| AMPL2000525 | TATATATTGGGCTTTGCTCGGTGAT | GGCCTATCTGTTGATCCTACGATAG | *Oryza sativa L.* |
| AMPL2000526 | TGTTAAAATGAAGGGACAATGGACG | TCCCAAAATACGTAAATAAGCGAAA | *Oryza sativa L.* |
| AMPL2000527 | GTCAGGTGCACCATCCTCAC | TGGAAGAACCTGCGAACGTC | *Oryza sativa L.* |
| AMPL2000529 | TCTCCACTGTAAGGTGTTCAATGTA | TGTTTTTCCTGTACGTTCTGACAAG | *Oryza sativa L.* |
| AMPL2000530 | GCAGATTTACAATACAGATGGGGTG | CCTCTAACTATTGAAAGTGGTCCCT | *Oryza sativa L.* |
| AMPL2000531 | TGCTTCTTCATTTGTTCAATCTTGT | CAAACATAAGACAGATGAGCGCTAG | *Oryza sativa L.* |
| AMPL2000532 | AACCCGTCAACATTTAACTTCATCC | ACTAAGTTCTAACTGGTAAGCACGA | *Oryza sativa L.* |
| AMPL2000533 | CGACGCTCGGGTTTAAAAATATGTA | TCCACATCCAGAAGTTAATCGTCTT | *Oryza sativa L.* |
| AMPL2000534 | TGTTGGCCTACATCATGATGCTATA | CATGTCGAACTTGAACCCCAATG | *Oryza sativa L.* |
| AMPL2000535 | TTCATTAACACGCGCATTACACTTA | TCGTTCGTAACTTAGCTAGGCAATA | *Oryza sativa L.* |
| AMPL2000539 | GAGTGGGTATGGATGTGCAAGG | CGAGGTTGAGGTTGAGGTTGAG | *Oryza sativa L.* |
| AMPL2000540 | GTATGGGGGTTGGTGGTGTATG | AGCTTCAATCAAGGTAACCAAACAC | *Oryza sativa L.* |
| AMPL2000541 | ATATACACACACACACATATGCCCT | CTACGTAACCCATTCTCAACTGGTA | *Oryza sativa L.* |
| AMPL2000542 | AATGGAAATCTGTTTGTGCGGAAT | TAGAAGTCAGTTGATATAGCCGCAA | *Oryza sativa L.* |
| AMPL2000543 | AACCCATATTGGCTATGTTTTGGTG | CTGGCGGGATCAGATATTGGATG | *Oryza sativa L.* |
| AMPL2000544 | AGAGATCTTGGAAATCTCGAAACCT | ACGCAAATAGACCCAGAATTTTGTT | *Oryza sativa L.* |
| AMPL2000545 | CTGGCTGCCAGCTTATTATATATGC | TCTCATAGTCACATGCATGTACTGT | *Oryza sativa L.* |
| AMPL2000546 | CAGGGGATTGATTTGCACATTTTTG | TTCTAGTCACTATCGCTATTGCCTC | *Oryza sativa L.* |
| AMPL2000547 | GGGTTAGTTACAGTTGGGCCTAAT | GCTCGATGTAAAACCAATGGTGAAT | *Oryza sativa L.* |
| AMPL2000550 | TAACTATAGCCAGACCTTGCCTTTT | TAGAGCTCACTCAAATCGATCTCTC | *Oryza sativa L.* |
| AMPL2000551 | GCTATGTGTTATCAACCGCTTGTTA | GCCTTTTCAATTGAATATGACGGGA | *Oryza sativa L.* |
| AMPL2000552 | AATACAGTGCGGGAAAAAGACAAG | ATCTAGCTGGAATAAATATCCCGGC | *Oryza sativa L.* |
| AMPL2000553 | TCCTCCAAAACCTTAAATTCACTGC | CAACATCTAGATTCGTGGGTCCTAT | *Oryza sativa L.* |
| AMPL2000554 | CTTGATGCAGAAACAAGCAAAGCT | ACTACCATGATTAGGCCTCTCTTTC | *Oryza sativa L.* |
| AMPL2000555 | CACAACAACAATCCCCAAAGTCATA | TTTGACTGTGCCAAGATCTCATCTA | *Oryza sativa L.* |
| AMPL2000556 | CCAATTTCAGTTTTTAGTTCTTACCGC | GTCGAAGTCCAGCACCATCG | *Oryza sativa L.* |
| AMPL2000557 | CCATATGTATGCAACTGGGGGTAAT | AAAGACACTTGTTGCAAGATAAGGG | *Oryza sativa L.* |
| AMPL2000558 | CTCTCCCTACGGTTGTTTCTACATT | CTGATATCCCCAATTGACACACATG | *Oryza sativa L.* |
| AMPL2000559 | CTGAGCTCGAGTATGTTCATGGTG | AAAACCACTCAATACTAAGTGCGTG | *Oryza sativa L.* |
| AMPL2000560 | AATACCAATAGTCGAGGACATCCAC | TACTCCCGAACAGGTTGATGAGAAG | *Oryza sativa L.* |
| AMPL2000561 | TCTCGGATTGGTTTGTTTTGAGTAC | GCAGCACAATCTTGAATAATCGGTA | *Oryza sativa L.* |
| AMPL2000562 | TGTTGTTTTACCCCACTTTTTAGGG | CTAGGGTTTTGGGTGTTCACATCTC | *Oryza sativa L.* |
| AMPL2000563 | AAAAGAAAAACCAACCCAATCACCT | CTCCACCATCATCGATCTTCAACA | *Oryza sativa L.* |
| AMPL2000564 | TTTAAACTCTCCATCTCCGTCCATC | ACAGGAGAAGAATTGCTAGGGTTAG | *Oryza sativa L.* |
| AMPL2000565 | GCAGTCAAAATCAGAAAAGAGAAGT | GTGGGTGCCTAGGTATGGATATATT | *Oryza sativa L.* |
| AMPL2000566 | CGATACTCTTTGTCATCCTCTCTCT | CTCCCCTCAACTTCATGCACTAG | *Oryza sativa L.* |
| AMPL2000567 | ACCCGTAGAAAAGATAAGAGAGACG | TTCGCCCATGTCAAGCTTATTTTT | *Oryza sativa L.* |
| AMPL2000568 | GACATAACGCGTGATGATTTACTGT | CTGCTCGATAACCTTCATCTGAAAT | *Oryza sativa L.* |
| AMPL2000570 | CAGTGCTATCCAAATTGACACAACT | CTCAATGCCCATTTACATTTGCTTG | *Oryza sativa L.* |
| AMPL2000571 | TTAGAAAGCATAACAATCCGTCACG | GCTTTATGCACGGAAAATTGACATA | *Oryza sativa L.* |
| AMPL2000572 | CTGAAATGACCTTATTGCATTTGGC | GACAAGAGAAACGTGGGCTTAAATG | *Oryza sativa L.* |
| AMPL2000573 | AGAGAGCTTACAAATTGATAATTTTCACA | CCACTCAAAAATGATGTGGAAGACA | *Oryza sativa L.* |

| | | | |
|---|---|---|---|
| AMPL2000574 | CCTTTTCAAGATAATGTAGGGGCTT | TTTCTACAGCATGTTCATCGACAAG | *Oryza sativa L.* |
| AMPL2000576 | TTGTAACAGAACGAAACATGACCTG | CAAAACTTCCTTGCAATTTCAGCAT | *Oryza sativa L.* |
| AMPL2000577 | ATTTGATACTGTGAGGTACCGGTAG | GAATGAAACTCAATGTGTTGTGCAG | *Oryza sativa L.* |
| AMPL2000578 | TCCCCAAAATATAGCAACTTCTGGA | TATCAAGGTGCACAAAGTTGCTAAG | *Oryza sativa L.* |
| AMPL2000579 | ACCACACACAATCTGATCTTTTGAG | ATAATTCACTGTGGTGTGTTTTCCC | *Oryza sativa L.* |
| AMPL2000580 | GAGATGTCTCATCATGGAACTCGAT | CATCAAGGTTTCCAAGACCCTTAAC | *Oryza sativa L.* |
| AMPL2000581 | TTCTATAGCTAAAAAGTTCTTAAAATCTGGA | ACCAGTTCATCCAGTTTATCCGTTA | *Oryza sativa L.* |
| AMPL2000582 | TCACCACTATCATTTAACCGGACTT | AAATCCCTAATTAAAACCGCGGTAC | *Oryza sativa L.* |
| AMPL2000583 | ACCAGATCGATGATCATGTTTAGGA | AGTGCGAGATTCATTTGTTTGTTTT | *Oryza sativa L.* |
| AMPL2000584 | GGGTCTCACACTTGGAAGAAAATTT | CTCACAGGACACGACATTCAGG | *Oryza sativa L.* |
| AMPL2000585 | GAAGAAGCGCGGCTCATTG | TCTTCTTCCTTATCTCATCTGCGAG | *Oryza sativa L.* |
| AMPL2000587 | CCACCCTTTTATCGTTGAAGTTTCA | TCCCTTATATTCACGGATAGAGGGA | *Oryza sativa L.* |
| AMPL2000588 | TGCCGAATTATGACCAAATTTCCAA | AACGGGATCCTAGGCAGTGTATC | *Oryza sativa L.* |
| AMPL2000589 | GCTCGATTGAAAAAGAAACTCTGGA | GATGCAAACCAGGACTATATAGGGT | *Oryza sativa L.* |
| AMPL2000590 | GATCGGTAGACGCTGGTGAG | ATATACCCTAATCGATGGCTCATGG | *Oryza sativa L.* |
| AMPL2000591 | CCTTGTAGATGTACTTTCCGAGGAA | CTTCGTCATCTTCATCCGCAC | *Oryza sativa L.* |
| AMPL2000593 | AGAAGCAATCACCCATAGCTACTTT | CCTTTAGGAACACATGAATTGACCC | *Oryza sativa L.* |
| AMPL2000594 | CTGCTCATAAACGAGTTCACGTCTT | CCCTAATCCTTGATTCCATCTTCCA | *Oryza sativa L.* |
| AMPL2000596 | AAGAGGGCATCATTCTTGTACTCTT | AATCTTTCTAATTTAATGCGGCCGT | *Oryza sativa L.* |
| AMPL2000597 | CCGGCTAATCATTCTGAAACTTGTT | AATTTGAGTCGTTTAAATGGGTCGG | *Oryza sativa L.* |
| AMPL2000598 | CTTCCTATATCCCTCCTCCTCTCC | ACCACAAGATAATAAATCGACCGAG | *Oryza sativa L.* |
| AMPL2000599 | CCCAAATTTTAGAACTACGGCTTCA | ATCCTGCACTCGTGTGTGG | *Oryza sativa L.* |
| AMPL2000600 | TTGCCATGCTAGATCTAGGAATTGA | CATTGCTACCGTTGTTCCAAATAGT | *Oryza sativa L.* |
| AMPL2000601 | GGATCTATGATTTGTTACTGCGCAA | AAATTCGATCCTATGCTAGCTCCAT | *Oryza sativa L.* |
| AMPL2000602 | TGATGTGTATCATACCACGGTCAG | GGGTTTATATGTCGTCAGGACCTAG | *Oryza sativa L.* |
| AMPL2000603 | AAATTTAACACCACCACCCACATAG | CACAACCACCCTTCCTTCTCC | *Oryza sativa L.* |
| AMPL2000604 | AGAAAAGACATATTGCAACATCGCA | TTTCTCTCTTATCTCCCCATGTGTG | *Oryza sativa L.* |
| AMPL2000606 | ACAAACTTATACATCCACATCTGCT | TATCGAGCTGGACATTTCCTTTTTG | *Oryza sativa L.* |
| AMPL2000607 | CGTAGGGCTGAAATAATGAGAGTCT | TGAACCGCTCCTTATTAAGAGTGAA | *Oryza sativa L.* |
| AMPL2000608 | TTTTGGACCAAATTACCCATGTGTT | GGCCTCGCTACCAATACCTG | *Oryza sativa L.* |
| AMPL2000609 | TTTTGTAGGTCGTCCGTTAAAGATC | GTTTGTTCCCTGGAGATAGGCATAG | *Oryza sativa L.* |
| AMPL2000611 | CTGCTTTTCCTTCAAATGTACCCTT | CATACATGCATATGGGTCCTAAGGT | *Oryza sativa L.* |
| AMPL2000613 | TTGTCGCATAAGTCAACAAAGGAAA | TAATTTTTGTTGGACATGATTCGCA | *Oryza sativa L.* |
| AMPL2000614 | AAAATTGCACACAAACTAAACACGG | GGAGGTATAGTGTTGCTCATGAGTT | *Oryza sativa L.* |
| AMPL2000615 | AGACATATTTTTCTACGTTATTGTTAGCT | GTCGGTTGCCCATATATTCTCAAAT | *Oryza sativa L.* |
| AMPL2000616 | AGAAATAGTTGCTATTGGGCATTGG | CGCTAGTTGTTTAGTCGTCAGAATC | *Oryza sativa L.* |
| AMPL2000617 | CGTGGCCAAATTTCCAATTGTTTAA | TACCCCATATCATTGAGGGCTAAAG | *Oryza sativa L.* |
| AMPL2000618 | TTATTTAACATTCGAGTTGGGCCAC | TAACACAATCTTTCTTTACGGCCAC | *Oryza sativa L.* |
| AMPL2000619 | TGAAATTCCAACCATATTCCAACCT | GCCAATATAAAGGTTCTGTTCAGGA | *Oryza sativa L.* |
| AMPL2000620 | TATTCTGAATGGCTGCAGACTATGG | TCTTTAGTCTCGATTTGTAACCCGG | *Oryza sativa L.* |
| AMPL2000621 | CTGCCTCCTTTCCCCTTTCC | CTACTCTTCCAAAATGCCGTTCTAC | *Oryza sativa L.* |
| AMPL2000622 | AAATTGAGGGGGATTTAGAGGTACC | GTAGCAGCAGGGAACATACAGAT | *Oryza sativa L.* |
| AMPL2000623 | GCCAGTGCCATCAAGTTTTTAATCA | CGATTCACCCAAAGAAGTACCAAAA | *Oryza sativa L.* |
| AMPL2000624 | CATGTAACGGTCCGATTTTCGTTC | CGGCAGTTGGATCGGGAAC | *Oryza sativa L.* |
| AMPL2000625 | CAAAGGAGTAGAGCTTCATGTTGG | TTACCCGTCAAGATGCCGAC | *Oryza sativa L.* |
| AMPL2000626 | GAGTATTGTCTTGTGTTCGCCAGTT | ACAAGCATGCAACTTCAGTAATAGC | *Oryza sativa L.* |
| AMPL2000627 | TATTCAAGAATGGCACCACTGAGAT | ATTTGTTATTTCTTGCCCTCTCTGC | *Oryza sativa L.* |
| AMPL2000629 | GTTGATGGATTATGTGACTCCGTTC | GCATGCTAATAATTCACCTCGTGAT | *Oryza sativa L.* |
| AMPL2000630 | ATGCTCTCATGGCATTTACAGTTTC | ATGAAACACTTTCTTGATGATGGCA | *Oryza sativa L.* |
| AMPL2000631 | GATCTTCTCGATCTCCAACTCGC | ACCGAAATAATATCGTATTGGAGGGT | *Oryza sativa L.* |
| AMPL2000632 | CGTACAGGTACCACCACCAC | ACGGTTTAGGTTCAATCCATATCCT | *Oryza sativa L.* |
| AMPL2000633 | GAAGCGGTACAGGTTGGTTTTC | AGAAAATTATGGCTACACTCCCTGT | *Oryza sativa L.* |
| AMPL2000634 | GAGAAGATCTACATGGACGCCC | TTCTATTTCCTTCGACTCTTCACCG | *Oryza sativa L.* |
| AMPL2000635 | ATGCGAGATCTTGTTATAAAGATTTATTCT | AAAAATCGGACTTCTGATCCAACAC | *Oryza sativa L.* |
| AMPL2000636 | TCCCATAACACATCAGTTAAGGAGG | TTAGGTGTAGGGTCTGGGATGATTA | *Oryza sativa L.* |
| AMPL2000637 | GATGGAGTCAGAGATATCGACGTAC | TTATCTCCCTGTCATGTCATGATCT | *Oryza sativa L.* |
| AMPL2000638 | CTTATGGGATCTATGTCTTGCAGCT | TAATAAACTTTGGCTGTCATTCGCA | *Oryza sativa L.* |
| AMPL2000639 | GAGAGACCTTCCTCCAAACATATCT | ACATCGCTAATGTTATAGAAGAACGA | *Oryza sativa L.* |
| AMPL2000640 | GTAGATGTTGGTGCTCATATAACGC | AATTGGAAGGGTTGGGGTAATACAA | *Oryza sativa L.* |
| AMPL2000641 | ACCCAACTCTTCAAGCAGAGAATAA | CATTTCAGGCTTCAGAGATGCTATG | *Oryza sativa L.* |
| AMPL2000642 | CGAAACCGATGCAGTCTCATCAA | CACCAAACCAATCTGTTTAGGTGTT | *Oryza sativa L.* |
| AMPL2000643 | AACGACCCACATATAAGGCCATAAT | ACATGGTCACTTTTAAAATTTGTAATTTACA | *Oryza sativa L.* |
| AMPL2000644 | GCTTTCTCCTGATTTATCTGCCAAA | TTAATTCCCTTTTCTCTCCGTTTGC | *Oryza sativa L.* |
| AMPL2000645 | ATCTGAAATATTTTGGACGATCGGC | TATACTCATACAGACGCACACTCAC | *Oryza sativa L.* |
| AMPL2000646 | GGGTTTGAATTGTTATATTGTGCGC | AATATATGGGCCTGACTCGTTACAA | *Oryza sativa L.* |
| AMPL2000648 | TCCACCCAAGAGATTAGAATCCATC | TCCAAAAATTATATACTCCCGCCCA | *Oryza sativa L.* |
| AMPL2000649 | CACCGCCCAGATCTTGTCTC | CTTTAGATCCAGACAAGGTGGGAC | *Oryza sativa L.* |
| AMPL2000650 | CAAACTCGAGGAATTCAAGACTTGG | CTAATTGTGAAAGAGCAGGACATGG | *Oryza sativa L.* |
| AMPL2000651 | CTAATAGACATCGAAAAACGTGCCA | GCCTTTGAGCTAAGGTTGAAGTAAA | *Oryza sativa L.* |
| AMPL2000652 | ACAAAAGCAGCAATGAAGACATGTA | CCACCAATTAGTACACCACAAAACT | *Oryza sativa L.* |
| AMPL2000653 | GAAATGGACGTACGGGAATATGC | TTGACGTTTTGTAGGCTAAAGGAAC | *Oryza sativa L.* |
| AMPL2000654 | AAAGTGTACTAGGGGTAAGGTGAAC | TCACTTGATATGCATGCATTTCACA | *Oryza sativa L.* |
| AMPL2000655 | GGTTGCCGATTATTTTTCCTAGTTT | AGGAAATATGTCGCTTTTCAAGTGA | *Oryza sativa L.* |
| AMPL2000657 | TAGAGAATGGCGTCAACTTTTGGAT | TGATTAACACAATGCAACCTCAACA | *Oryza sativa L.* |
| AMPL2000658 | TTCAAGAAGGTCATGGCATCATAGA | GAGTTTTGAGTTGGCAGAAGTTGTA | *Oryza sativa L.* |
| AMPL2000659 | ATATGGGTTCCTTGTAATAGTGGGG | TTGAACGCATGTTTCAATTTTCAGG | *Oryza sativa L.* |
| AMPL2000660 | ATTGAATTGCCCATACATGTGGATC | TTTTAGTGAGATTAGGGATGTGCCA | *Oryza sativa L.* |
| AMPL2000661 | GGATCAATGGCAATGGTAAATCGAT | TGCGCATTCACCACTATAAATTCAA | *Oryza sativa L.* |
| AMPL2000662 | TCATTAGTATCTGATAAGTGGGAAACA | ATGTCTGATGTGTAGGAATTCCTCA | *Oryza sativa L.* |
| AMPL2000663 | GTAGAAACTACAGTGCCTTTGGTTG | AGAGGAGAAACCCATCAAGTAGAAC | *Oryza sativa L.* |
| AMPL2000664 | AATCTTTCTTGTCACTATTTGCATAGG | CACAACCTTCGCTCATATTTCCTTT | *Oryza sativa L.* |

| | | | |
|---|---|---|---|
| AMPL2000665 | CTACGCATAGGTAATTGATCTGGGA | GGGCCACAAGTAGGAGTATATGATT | *Oryza sativa L.* |
| AMPL2000666 | TACCCCTCATTAATACTCCAAGCAG | AACAGAACACACCTACAGATTCAGA | *Oryza sativa L.* |
| AMPL2000667 | AGGGCAGAACATTTCCATTTCAAAT | TTCAATTCATCCTCCTTTTGAACCA | *Oryza sativa L.* |
| AMPL2000668 | AGAATTTCACAGTGCAAACCAAGAA | AGGATTCATGGCATAGAAACTTGTG | *Oryza sativa L.* |
| AMPL2000669 | AGCAGTGATTTGGTTTGTCTTACTG | GCAATAACCGTCTAATCGCACTAAA | *Oryza sativa L.* |
| AMPL2000670 | CATCCTACAAATCCTAGATCTGTCAC | GTTTATAATTCGCTGCAGATCCGG | *Oryza sativa L.* |
| AMPL2000671 | ACATGTCTTAAGCATATCGATTCCA | GCTTCAGTTAGACGAAACAATCCAA | *Oryza sativa L.* |
| AMPL2000672 | TGTCAAGTTACGTAACACATCAACTG | TAAGGGTGGATTCTTTATGGCATCT | *Oryza sativa L.* |
| AMPL2000673 | AGGACTAAAGATGTTTTTGGGACT | CGACATCTCGAACCTACACACG | *Oryza sativa L.* |
| AMPL2000675 | TGGTATCCACTTGATTCAAAGTTTTGA | AGCGAAGATTAATTGTTACCTTAGTGG | *Oryza sativa L.* |
| AMPL2000676 | TCTGCATTATTGGTTTTTGGACTTT | CCGCACAATAATCCAACCAGAATG | *Oryza sativa L.* |
| AMPL2000677 | ATAAGTCCCCAACGCTAATTACACT | AATAAACCCTTTTCACGATACCAGC | *Oryza sativa L.* |
| AMPL2000678 | CCGTACCTTTTCCTGGGGTTC | GGGAGGACGAAGGAGGAGAG | *Oryza sativa L.* |
| AMPL2000680 | TAGCTTCGACAGGGAACGCTTATC | CATCTGGCGAAGAATTTGGAGTAC | *Oryza sativa L.* |
| AMPL2000681 | TATTTGTGATTCACATGACCACGTG | GGAAAGGGAACTAAGTAAGAGCTGA | *Oryza sativa L.* |
| AMPL2000682 | CAAGGAGAAGGTGTTCGTCCAG | CCATATGACGACTGTACTTCTCAGA | *Oryza sativa L.* |
| AMPL2000683 | GGTGACAAACGTGATTTATTGCAAT | CATATCCGTATCCGTGTCATCAGTA | *Oryza sativa L.* |
| AMPL2000684 | CGACCTTTTATTCTTTGCTTATTGAAATGA | CTGATGATCGGTCACTGGAATTAGG | *Oryza sativa L.* |
| AMPL2000685 | ATGTGAAACTAAAAACAGCCTTTGT | CTATTAGCATGCGGCATACATGAAG | *Oryza sativa L.* |
| AMPL2000686 | TCTCCTTCCTCTCCCATCTCATC | AAAAAGAAAAACACACGAATCGCTC | *Oryza sativa L.* |
| AMPL2000687 | CGGCATAGGCGTTGGTGT | GATGATCTTTTGAGTTCGCCCTAAG | *Oryza sativa L.* |
| AMPL2000688 | AGAACATTTGCTAACCTCAACGTTT | TTTTAGGAGTTGGAGCTCTACCAAA | *Oryza sativa L.* |
| AMPL2000689 | GAGTAGATCTTGTAAGGGAGACGC | AAGTCTTCGTTTCAAGTTGGATACC | *Oryza sativa L.* |
| AMPL2000690 | CATGCCATGAATATCAACAAGACCA | AAGAGAAAATGACTGCTCCTTCTCT | *Oryza sativa L.* |
| AMPL2000691 | ACTGCATCTCTACAAGCTTATCAGT | AAAGCGTTTCTTCATCAAATTTGCC | *Oryza sativa L.* |
| AMPL2000692 | CTATTTCATCTGCGGCTTCCCT | TTTAAAAGACCATGAGACCAGCAAC | *Oryza sativa L.* |
| AMPL2000693 | TGCTCAAAGACAGGTCGAACATATA | GTGCTCTTTGTACAACCCATCTTAG | *Oryza sativa L.* |
| AMPL2000694 | GCGAAATTGAAATAGCGTGGGT | CGTAAGTACGTACAATACCTCTACCA | *Oryza sativa L.* |
| AMPL2000696 | TGAATCCCTTCATTTGTAGGAGTCA | TAGATCAAACACAAAGACAACAGGC | *Oryza sativa L.* |
| AMPL2000697 | GGAATCGAAGCTAACAATCTGCAAA | ACCTGATAAGAAGCATGGAGAAACA | *Oryza sativa L.* |
| AMPL2000698 | CGTAAGAAATACCAACAACCACCAA | GATCACTGCAACCACAACACTAAAT | *Oryza sativa L.* |
| AMPL2000699 | ATTGGGTTGAGGAGAGAAGGGAATA | CCGTTTCTCTCCAGTGTGTAGG | *Oryza sativa L.* |
| AMPL2000700 | TGGTTTTCGTAGTAGTGGTACTTGT | GGTACGTATCCAACATAGTAAGTACA | *Oryza sativa L.* |
| AMPL2000701 | TCAGGAACAGCTACACAAGAGTAAA | TATTCATGGATGAGCCTACATCTGG | *Oryza sativa L.* |
| AMPL2000702 | GTTGGATATAGAGGCCGGGTAAAAA | ATTGCCGAAATGCTGGTCCATC | *Oryza sativa L.* |
| AMPL2000703 | CACAATCACCTCTATTCACGGATTG | AGATGGAGCAATTACCTTATTGGGT | *Oryza sativa L.* |
| AMPL2000705 | TCCCTTGACTTCAAAATTGACGATG | TCTCCAGATTTTATGGATCCCAGAC | *Oryza sativa L.* |
| AMPL2000706 | AACTAAACAGGCCCAAAATAGAAGC | CGTCTCCACGTTTCTACTACCTTTA | *Oryza sativa L.* |
| AMPL2000707 | TTTTGGTAGATCATGCTCCTTTCAG | CCATTGCATTGACTTCATTTTACTGT | *Oryza sativa L.* |
| AMPL2000708 | TGAGGACCAAGTTCAAAAGAAATGA | CTCTGGTTTAGTGTATCGCAAATCA | *Oryza sativa L.* |
| AMPL2000709 | ACAATTAGACAGAGGGAGAGAGAGA | CCCATGATAAGCTCAATTGGTTACC | *Oryza sativa L.* |
| AMPL2000712 | GGCCCAACTTAAAGGTGATATAAGC | TCTTCTTCTTGGTTTCTACTGGTGG | *Oryza sativa L.* |
| AMPL2000713 | CCGGAGAAGACATCAAACACCTATA | GGTGCCTTTTACTCATTCAATAGGG | *Oryza sativa L.* |
| AMPL2000714 | GATATGGAGTTGGTGGGACGAC | GATGGCCACAGAATAGTCCATGA | *Oryza sativa L.* |
| AMPL2000715 | TGGTAAACACAAATTCAACACGGG | TTCTGGTTTTGAAAGGTTGCTTCTT | *Oryza sativa L.* |
| AMPL2000716 | TCAAGTTTGCTTTAAGGTACAGACA | GGACAACAGGGTACTCACATGAAAT | *Oryza sativa L.* |
| AMPL2000717 | AAAACTTTTGGCAAAACATGAACCG | GTTCCTGTATCAAGAATGAGACTTGT | *Oryza sativa L.* |
| AMPL2000718 | TGATCGATCACAGATTCACAGTTGA | TCGGACATGTACTCTTCTTGTTCAT | *Oryza sativa L.* |
| AMPL3258394 | GAATTAAAATAAGCTTCCCATCATGCCAAA | CTTTAACATATTCAAATCGACGTCCTCATA | *Gossypium spp* |
| AMPL3258835 | GGATAATGCACTTGAATACACTCGAACC | ACGACCCATATTTAAAGTAACTAACGAGACA | *Gossypium spp* |
| AMPL3258845 | AGAAACTGAAAACAGAGCCACCATTT | AAAATGGAGGTAAACCAACATGGAGTC | *Gossypium spp* |
| AMPL3258847 | CATCTTGTAGCTTTAGTTCAATCCATTGCA | GTCTTATCTCCCTGAAGTTGTAACAGAGC | *Gossypium spp* |
| AMPL3258849 | GTCATTTAAACAACCACATAGCCCTAGTT | AATAGGAGCTATTGGTGATGATGGTTGG | *Gossypium spp* |
| AMPL3258850 | AATAATAACTCGGTCATTCAGATCATGCAT | CCGTGTGCCTACTGTAGGAAATTATATTG | *Gossypium spp* |
| AMPL3258856 | CTTACCTCACACCGTACCCAAAG | TGTCATTTTATAATAGGAGGCAAGTTTGGTT | *Gossypium spp* |
| AMPL3258858 | ATGAGAAATTCGAGGGAAAAATTCCAAAGG | TTTAGAACAACATTGGAGCACCAAATGA | *Gossypium spp* |
| AMPL3258863 | TCAAATTCAGGCCTCAAACATCCAC | ACCACCTATACCAAGTATTTCTACTAGCTTG | *Gossypium spp* |
| AMPL3258864 | CAGACGTGTTTCTTGAAGAGTTGTCT | GTCTCATATTCGGCTACAATCATATCCTCC | *Gossypium spp* |
| AMPL3258871 | TGGCCAATTTTAGAGAAAGAGTGTGAA | CGAACATTATTGTATCCCTGGAAACTCAATT | *Gossypium spp* |
| AMPL3258872 | GTGGTACAAGTACCTATTTGAAGTGTATGAG | AGTGCTTCTAATGATAGCGTAATCGTAGC | *Gossypium spp* |
| AMPL3258875 | CAAGATGTAGTCGCAAATGAGTTTTGAT | ATCAATATTGATGTGGCGGAGATAAAAGTAA | *Gossypium spp* |
| AMPL3258877 | TAGTCGTGCTTCAGTACTATCATAAGGTG | ATGCGACAGTTTAGGTGCAAACA | *Gossypium spp* |
| AMPL3258888 | CAAGAGCTCCTCAAGCTCATCTC | CCGATTGATATTGAATATTTTGTGGCTCAAT | *Gossypium spp* |
| AMPL3258889 | GTCGAGCACCACCACTTTCATAC | CCACCTGCGATGTATGTTTTGTGC | *Gossypium spp* |
| AMPL3258943 | TATGGTGTCAATGAGGACACAATGATTAGG | GTGCTTAGACTATGTAACTCACTGTGAATAT | *Gossypium spp* |
| AMPL3258956 | CGTGTGTCCAAACACACAGTTGT | CCTGGCCTGGATATACTAGCGTG | *Gossypium spp* |
| AMPL3258966 | CCTACTTCCATTACTGAAGCACTTCAA | CCGAATAGTCATTGGTTTGACAACAGG | *Gossypium spp* |
| AMPL3258968 | GAGTATAACAATAGTCCGTCTCCTAGTCAA | GGCCTACCACTCATTAAATCACAAAGTAAAA | *Gossypium spp* |
| AMPL3258975 | AGTTATGATTCTGATTTGGGAAATCCTTGTA | AATGCATTGCATAAGGATTAAATTGTTGCT | *Gossypium spp* |
| AMPL3258981 | GAGGTTTAGGGACTAACTTGCAAATGAC | GGGATGAAAAGGATGATTAAGCTCCAAAG | *Gossypium spp* |
| AMPL3258985 | GCCACAGTTGTCTCACCAATGTC | GGCGACTCTATATTTCTGTTTTGCAAAAGT | *Gossypium spp* |
| AMPL3258989 | CCCATACTTGGCCCAAATGAAGG | ACATAAGAAGCATGTCTTGAGACTTGTTG | *Gossypium spp* |
| AMPL3258991 | TTTTACCCTTAAACTTTCGCATATTTACGCT | GGGTGTTAGATAAACGACTTTTGCTGAG | *Gossypium spp* |
| AMPL3258997 | TTTAGTAGGTGTTAAGGGAAACCCATTGA | CCTGATTTAAGGTTGTCCATTCTAAGAGAAG | *Gossypium spp* |
| AMPL3258998 | GAAAGGTGTTTAGGTCTTTTAAATGTACGCT | CAGTGTAGCTCACTGAGTTAGGTCA | *Gossypium spp* |
| AMPL3259003 | CCCTGAAGACCCTGAAATTGAAAGGA | GCAATTTTCAAAGAAATGGCCATCTCTATAG | *Gossypium spp* |
| AMPL3259004 | CTTGTTAGTGTATATGGCAGCAAAACCA | AGAAGCATCATAGATACATCTTGGACACTAT | *Gossypium spp* |
| AMPL3259014 | CACACACCAAGTATTTTAGTAAACGCTC | CGCTGCAGAACATGTTTTTAAATGAATAAAA | *Gossypium spp* |
| AMPL3259017 | AGAAACATGTTACAATTCTCAGACAGACAA | TTCCCTTTTCAAGTTCTGATTTCTGTTTTCT | *Gossypium spp* |
| AMPL3259025 | TTCAATATTGAAGGAGGAACATCACCTTTA | AGCTTAAGATTTTCATTGCGTTTGGTC | *Gossypium spp* |
| AMPL3259026 | CGATGTCAGACAATTTGGCCAAT | AGGGAACAGAATGATTGCTTCTTACTTA | *Gossypium spp* |

| | | | |
|---|---|---|---|
| AMPL3259029 | GGGATAGAGGATCCCTTTGACAGA | GTTGATGGCGCATGAGCTTTTAC | *Gossypium spp* |
| AMPL3259039 | CACTGCGGGTGAAGTATCCTACT | CTTTAACAAAGAGGCTAATGAGGGAGAAG | *Gossypium spp* |
| AMPL3259047 | CAACTCACCAACACAATGCATTATTTTTGT | TTTCGGTGTACTTGTTAGTTTACTTACCAAT | *Gossypium spp* |
| AMPL3259052 | CACTCCTCCAAATGGTAAAGAATCACAT | TTCAAAACGTTCTGGTTGAGTTTTTAAAGTA | *Gossypium spp* |
| AMPL3259060 | GGGTAAAATTGCTTTTTGAATACCTTTTGGA | CTCAATGCCGTTATGGATGATCATGG | *Gossypium spp* |
| AMPL3259064 | GATCCGATTCACTTAAATCAAGTCCTTCAA | CAAATTAAACCGTTCACCATTCCAACATT | *Gossypium spp* |
| AMPL3259066 | CCCTTGTGCCTTATCAATACAATTTTCCA | GTATTGATGGTTATCCGACATTGTTCTTTCA | *Gossypium spp* |
| AMPL3259070 | GCTGGATCATCTGTCTGTCAAAAGAAAA | GGAGCAATGGTCGTGCAGAATAG | *Gossypium spp* |
| AMPL3259072 | CTCAGATTTCCACACTCCCGTATG | GGTTAACGAGTTTATACGGAATTGATGGTAA | *Gossypium spp* |
| AMPL3259076 | TCGATCATTCGCTTGGTTACCAAC | CTTACCTCAAGGACAAAATAGCCCAAC | *Gossypium spp* |
| AMPL3259090 | CAAACACTCGAGGCAAATCTAAAATCAC | GCTCGATAATTGAAAAGAACCAATCCTACG | *Gossypium spp* |
| AMPL3259092 | CCTTTGAATCTCAAGCAATCATCTGAGT | GAGTTAGTGATTGACTATCATCTGGGAAAAG | *Gossypium spp* |
| AMPL3259098 | GTGGGTCCAATTAAGGGATATTTTGAATGA | GCATAGAGGTCATATGGCCTTTGTGG | *Gossypium spp* |
| AMPL3259099 | ATTCTTCTACTCTGCCACGCATTA | CTTTACCGTTGTGGGAGTTTTGACT | *Gossypium spp* |
| AMPL3259100 | CCGAAGCATAACTGCACACTCAC | AGGGCTCGACAAAACAGAAAGTAGA | *Gossypium spp* |
| AMPL3259110 | TCGATGTTGAAGTTAAACTCATTTGGGAA | GCCAATTAACTCGAGCTCTTTGCTC | *Gossypium spp* |
| AMPL3259111 | ATCTGAATTAACTGAAGTTGAAAGGGATGAT | GAAAATAAGATCGTATGACACCCTCCATTT | *Gossypium spp* |
| AMPL3259113 | TCTTTCTTATTTGGTGGGTGAACGATG | GGTATCAAACGTAATTGGTGATCTCAAAACC | *Gossypium spp* |
| AMPL3259123 | CCTGAATCACCCATTTGTCCTGTC | TGATAATAACCTAGCAGAGCTGATCGAC | *Gossypium spp* |
| AMPL3259129 | GGTTAAATCGGGCTTAAGGAAGCTC | GATTGGTCAAAATTGTTGAAACATCGTTGA | *Gossypium spp* |
| AMPL3259136 | GAATCAAATCTTACCGGTAAAAGTACCATGG | AAAAATGGGCAAATTAGTCTCTATACGGAT | *Gossypium spp* |
| AMPL3259139 | GGTTTAAAATACTAGTGCTTCTCCTCCTTA | GGGTGATAGAGACAATAAAGGGTTGAGA | *Gossypium spp* |
| AMPL3259149 | GGTCGGGATGATTTTGTTGTGGC | GCAAGATTGCCAGAGAATAACATTAGAGAAA | *Gossypium spp* |
| AMPL3259156 | GCTTGTTCCAAATATTCTTAGCCATAGGAC | GGTTCTTTCTAATTGAAAAACGCTTATGGTA | *Gossypium spp* |
| AMPL3259161 | CCCTTAAGGGTTGGTAGTAAATCTGGT | CAGCAAAAACATGACCTTAAAATTAGGACAA | *Gossypium spp* |
| AMPL3259163 | CGCTTCGATTAATCACATTCTGAATCGG | TGGTCAATCTCAAAAGCACACAAAATATGC | *Gossypium spp* |
| AMPL3259167 | CTTGAGTAACCACAACTTCCACAATCA | AGGCTAGTGTTGGAAAAGGACAGG | *Gossypium spp* |
| AMPL3259170 | CTAAGGCCGACTTTTGTCCCTAC | ATTCTACTCTAGGAGGAATGTTTTTATGCGG | *Gossypium spp* |
| AMPL3259176 | TCAAGTGGTGTTCCCTCTAAAGCA | GGTGGACTCAGATGAGCTATTCCC | *Gossypium spp* |
| AMPL3259179 | GCCTATCATTACCATTCAATTAAAGCCCAA | CCGATGAAGCAAGGGATTTCAAAATCA | *Gossypium spp* |
| AMPL3259180 | GTATGATGTGAGTGACTCAAAATATTGTGCT | GGAGAGGAAAAGAAGAAATTTGAGAGCAAA | *Gossypium spp* |
| AMPL3259184 | CTTAGGTAGCTTTAGGGTTTTTGTTTTGTTT | CCCGAATTAACAAAATCAGAATTGCCTTAAG | *Gossypium spp* |
| AMPL3259193 | TCCCAAATTGTTTCTAACTTGTGATTTGGG | AAAGAAGTTGTGGTTTCCACTATGGTT | *Gossypium spp* |
| AMPL3259195 | CAGAGTCTACTAAGATGATAGCCCAAGTG | CTTCAATTTGGCCCATTTTTGCTCG | *Gossypium spp* |
| AMPL3259197 | GAATAAGTGTTGAGTGCGGTACTCAT | TCTCTTGGAAGCCTCTCCACATG | *Gossypium spp* |
| AMPL3259203 | GTTAGCTAGAGTTTTGTCATCAGAGCAA | CATTGACCTAAGCCAATGAGTGACAA | *Gossypium spp* |
| AMPL3259206 | TGAATGCATTAGACAAGCATTAGGCAA | AGTGGACTCAGCCAATGGGATAG | *Gossypium spp* |
| AMPL3259207 | AAGCATTAGGAAAGAGGGATGAAAACAC | CGTTTCAAACAAAAATGGTGTCAACAATCA | *Gossypium spp* |
| AMPL3259210 | CATCCAATCTTTACTCGCTCCTATATTGAT | CATAGAGTCTGGAAAGAGGGTTGTTTTTT | *Gossypium spp* |
| AMPL3259211 | GTGAACAGCTTTGACGTGAATATAGGTAA | GGAGTTATTGTAGAAAGCACATTTCTCCTAA | *Gossypium spp* |
| AMPL3259213 | AAGATCAATGCTCAGAATGAATGTTATCCT | CTAAACCCAGCTGCAAAATGTTTCAA | *Gossypium spp* |
| AMPL3259224 | CATTTTTCAGGAACAAAGACCCGAAA | GTTGTGTCTCATGAATACGGTATTGCA | *Gossypium spp* |
| AMPL3259226 | CCCGTATGAGCTAACCAAATTTAGCTT | GGTTTCGACTTTTCCAATCTTTGAATTTTCA | *Gossypium spp* |
| AMPL3259227 | CACCTTTTCAAAAGCCTGCCATTC | CAAACTGATGCTCAATAAGCATCATCGC | *Gossypium spp* |
| AMPL3259234 | TAATAAAGGTAATTTTGTCCCGTGGTGATT | GAAAGTCAAATTTAGAAGTGTCCCTCCTATT | *Gossypium spp* |
| AMPL3259240 | TAGCATTATGCGCGTATATATTGTGATTCTC | CACCCTCAAGGTATTTGCATCTGATC | *Gossypium spp* |
| AMPL3259244 | TATGCCAACCCTATGGCTAATATTTGGT | GAGTACGGTAGGGTAACTAGCATATTTTTT | *Gossypium spp* |
| AMPL3259245 | AACGATAACTACACTGATCCCAAATGAGT | ATCGATTTTGTTTCTTCTTCCTACATCCTA | *Gossypium spp* |
| AMPL3259247 | CCTGTGTGACCCAGTTACATGGC | CATGTTCCTTATTCGAGCTTTCTTGACC | *Gossypium spp* |
| AMPL3259250 | TAGCTCAATTGTAGAGGATAAAGCTCCTT | GTTCGATCTTCGATGCTGAACTGT | *Gossypium spp* |
| AMPL3259253 | CTTGAGCGACGTCCTTCTCATTC | GTTTAGCGAAACGCCGTCAATTT | *Gossypium spp* |
| AMPL3259258 | CCACGTGTGAAGTCGAAAACATT | GCATTCATCGTCATTATGTTAGATAAGGCAT | *Gossypium spp* |
| AMPL3259264 | CTTGCACATCAGCAGAATGTTTGAA | CCCTTCAAATCGTTTACACATATCAGCC | *Gossypium spp* |
| AMPL3259279 | TAGTGAGAAAAATGGCAGACGTGTTT | GACGTCCTTTCAATGAAGGAACCTT | *Gossypium spp* |
| AMPL3259293 | CATTCGCCAATATCCACCTCACTT | AATCTTGCACATATAAAGCTCCCACTTATG | *Gossypium spp* |
| AMPL3259298 | TAACAAAACCTTCTTCTTAAGCCTCTTCATT | AAACTTTCAAGATGAAGCTCAAAGAGGATAT | *Gossypium spp* |
| AMPL3259302 | GTTTAATTTGTACCTTTTTCCTGACGCAAT | AACGAAGAAACCGTGACACTGTAG | *Gossypium spp* |
| AMPL3259306 | GAAGAGGGCTAATGCTAATTGGAGAT | GATACAAAAGGAGTTCCGACTACATCTATTT | *Gossypium spp* |
| AMPL3259309 | CTCACGCCCATTTTTATCTCGAATGA | CTCTTCGTACAGACTTGATGTGTATAACTTT | *Gossypium spp* |
| AMPL3259314 | ACTTCACTTGAACAAGGATAGTCCATGC | CTAATTCTCAGGAATTAGTTTGCCGTAACG | *Gossypium spp* |
| AMPL3259315 | CCAAAATCTGTATTCCAGAGAGGCTTG | TGTGATCATCAAACAATAATGCATAGGTCTT | *Gossypium spp* |
| AMPL3259318 | GACCTGAACTTGACAAGTGTTACCAC | GACAATTGTTTCCACATTGAAGTCTAAACTA | *Gossypium spp* |
| AMPL3259327 | GGGAAACAACCTATTTTACGAAATATGAGGT | CCCACTCTCATTTTCTTCCTTCTTTTCTTT | *Gossypium spp* |
| AMPL3259332 | CAACTAGACCCAATTTTGGCCTATTACATAA | GAGGTGTCCACATTTTGGTTACTTTGT | *Gossypium spp* |
| AMPL3259333 | AGATTCGTCCACTATAATGCAAGACTTTG | CTGTCAAGTGGCTTGTATTTCAAGGT | *Gossypium spp* |
| AMPL3259339 | CCCTAACGCCATTTAAGAAAGATTTAGTGT | CTCTGACTGGCCATCTATTTGAGGA | *Gossypium spp* |
| AMPL3259340 | AGAAGACAATGAATAGTGCAGCCTTTT | CGTGGGTCTACATACACGGGATT | *Gossypium spp* |
| AMPL3259341 | CTAGGGTTACTATTTTGGTTGAGATCCTACA | CCCGTTGAACCTATTAGAATATCGAGGAA | *Gossypium spp* |
| AMPL3259343 | TCAGAAACAAAAGGAATTCCTCAAGCTT | CCACTGACTCGGCTTTCTTCTTTTC | *Gossypium spp* |
| AMPL3259345 | CCATCACATCTTTAACGATTTCATTCAGCT | AACATGGTCATACATCGGATTTATCATCAT | *Gossypium spp* |
| AMPL3259346 | ACACAAAAAGAACTATCTGACGCACTAT | ATCGTACAAGAGCAAACAGAGATTTTTGG | *Gossypium spp* |
| AMPL3259350 | TCGTGTGAAAATATGACTAAAACATCCCTA | TTTGTGAAGTGTACCTCGTCTTTTGAA | *Gossypium spp* |
| AMPL3259353 | GGTCCTTTCTATTCCTGGGACGAATAT | CATTTGGTCTGACTAAGCATAAGGTGAAT | *Gossypium spp* |
| AMPL3259358 | CATCACTATTTTATCAGGGTGGCTCATC | AGTGGACAACGGTCACATGAAGG | *Gossypium spp* |
| AMPL3259371 | AAAATAATCTACACCAATCAGTGTACGACTA | TGGTAGGCTTTAACAAAGAAGGCATTG | *Gossypium spp* |
| AMPL3259377 | ATGGAAGAGTGGCCAAACTTGAG | CGTAGCCAACACCTCATTACACAAAG | *Gossypium spp* |
| AMPL3259380 | CCAGCAGACGTTCCTTTGTTTGT | CTTTGGTGTGTACGTGCTTGAAGA | *Gossypium spp* |
| AMPL3259384 | TGGAACGCTTGTGGTCTATTCGC | AGACATTCGTGTGGTTTACCCATG | *Gossypium spp* |
| AMPL3259398 | CAAATTAAAAATGCCACTTATCAACCTCCTT | CAATGTAGGTTCCTTGGCTTGATCATATT | *Gossypium spp* |
| AMPL3259406 | GGATTTTTGGGTGGGAGATGATGA | TTCACCTGTGATCAAAACTGTCAAAAAGA | *Gossypium spp* |
| AMPL3259429 | TGGTAACCGTGTAGAATTAGAGTCACAA | CCCTATTTAAGATTGGTTTTGGTGACTTGC | *Gossypium spp* |

| | | | |
|---|---|---|---|
| AMPL3259437 | AGGGATTGAATTTCCCGGTTCAAA | CAGGTTAGGATGTGTTCTTATGTAAGACGG | *Gossypium spp* |
| AMPL3259454 | CACGGGACTCGATCAAGCTTTTAT | CAAACACAATTCAAGTGTTGACAAACCT | *Gossypium spp* |
| AMPL3259458 | AAACAAGTTCGGATATTGACTTCTTATGGTT | CCACCCTAATTGGAAAAGATTCATGATGTAT | *Gossypium spp* |
| AMPL3259461 | TTGAGAGGGAGCTAAATTTGAAACTTTCTT | TCACCGTATTGGATCAAGATTGGACC | *Gossypium spp* |
| AMPL3259463 | CCCGCAAAATTCGGATGATAAGAA | TCGCACATGCAGCGATCTG | *Gossypium spp* |
| AMPL3259466 | TACGATAACTCGAGAGCTGAGTGAT | TACTTTTACCTCATGCGTACATACACAAAAG | *Gossypium spp* |
| AMPL3259471 | TATGTTTTGAAGTTGGCTTTCACATTTCAAA | GCCAGCCCATGTAACAATTGAAATATCT | *Gossypium spp* |
| AMPL3259472 | TGCACTTTAAGTTCGTATGACATTGATAGTT | CCTCCTTAGATGAATGAAAATTGTGCATTTA | *Gossypium spp* |
| AMPL3259475 | ATTTTTCACAATTTTCCATTTCAAGGGTTTTGT | CAAGTTCAACTTGACACTGACCAAGC | *Gossypium spp* |
| AMPL3259479 | GTGAAGTCAGTTCCTTAATAGTTGAAGGTA | CCCAACAATCAAAGATATGGTATCACTTTCC | *Gossypium spp* |
| AMPL3259482 | TTGCGTTCACAGAAGAGGCTAAG | GTTCCCATATGTCAAGCATAGTCCTACA | *Gossypium spp* |
| AMPL3259489 | GACCTGACCTCCTCTGTTTGATC | AGAGTGGTTTAAACTGAATACGGATGATGC | *Gossypium spp* |
| AMPL3259492 | ATAAGGTGGAGCGGACAGTAGAG | CATGTCCAATTATCACTCTACTTAAGCCTAT | *Gossypium spp* |
| AMPL3259501 | TCGAATGGCCAAATTTGAAAGAGAAAAGA | AAGCTAGGATCAGCGTGTTTTGTAG | *Gossypium spp* |
| AMPL3259504 | CTACCTATAGAAACTAGCTGGTCAACTTGG | TGAGTCCAGAGGTACTTACTTATTGTAGCC | *Gossypium spp* |
| AMPL3259506 | TGTTTTTGTTATACCTCCCACTTGGATTG | TCCTTTAAGACAATACTTGGTTATTGCACTC | *Gossypium spp* |
| AMPL3259516 | CAGAGAATGAGCTTCAGCTCTCATG | CGGATCATGGGAATAATTCGTGAATGTG | *Gossypium spp* |
| AMPL3259517 | AAACAAGAGAGGATTGACTGGCAAA | CCCTAGTTCCAAAATGGTGTTTGCA | *Gossypium spp* |
| AMPL3259518 | CAAAACATCATCAACTCACACACATTTCAT | AAATTGTTGTAGTTGTTTTGATGGTTGGATA | *Gossypium spp* |
| AMPL3259529 | AGCTCACCTAAAGTTGCAACAAAACC | TTTCGCTACCAAGCTTGCTTTGC | *Gossypium spp* |
| AMPL3259531 | CCAAAAACATGTTGGCGGATACTT | TAACAACCTGGAGCAAGAACACTATGT | *Gossypium spp* |
| AMPL3259548 | TAAAGTTGCAAAGTGAGTTCTCAGATATGTC | CTCTTAGCCAAGGTGCTTAGCTTAC | *Gossypium spp* |
| AMPL3259552 | GCATAACCTCTAGTTCATTCATTTGGTCA | GTTGGTCCCGAATGATTTCTAATGTGT | *Gossypium spp* |
| AMPL3259554 | TGTTGTACTGTGCAATGCGGTAG | GCACATATTTGTTCTGTTTGACTGTCTGA | *Gossypium spp* |
| AMPL3259560 | CCCATATTGGCCAATAGCACCAT | CGATACCCAAAAGTGGATATCACGATTCT | *Gossypium spp* |
| AMPL3259567 | TTGGTTCGATTAACGCCTAAGACTTT | CGTGTCCAACTAGATCCAACCTAAGG | *Gossypium spp* |
| AMPL3259570 | CTTCAACAAGCATTATTGGCACACTT | CTCTCATAGGTACTAAAGCCTCCAAGAG | *Gossypium spp* |
| AMPL3259573 | TCGTAACTTATCGGGCATTAGGAGAC | ACTAGCACTCTAGCGGACATACAG | *Gossypium spp* |
| AMPL3259576 | CAGAGAGGGTTTCGCAACACTAA | GCTGTGTTTTGATTGGGTAATGTGTTG | *Gossypium spp* |
| AMPL3259577 | GCTTTGGATGCAGGTTCATAGAGT | CCAATTACTACACCATCAAAGTTGAACTGAA | *Gossypium spp* |
| AMPL3259578 | TGGTTTTATGGAGAATAGGGTTGATTCGA | TAGTTGGTGTCGGAAATTTAATCATGTCAT | *Gossypium spp* |
| AMPL3259579 | ACGAATGGGAGTCATGTACATGCA | CCTACTCGAACCTCTAACAATCGAATGT | *Gossypium spp* |
| AMPL3259580 | GGAGGATAATGCACTTCAACATACTTGAATT | GTTGAATTAAGATTTACGTAGCCTGACCTAA | *Gossypium spp* |
| AMPL3259588 | CCCTCTGCTCTCCCAATTTCTCC | CGAATTCGTTTTTCGAAAACGGAGAC | *Gossypium spp* |
| AMPL3259591 | AGGCTCGTAATAGGGAACAATATGAATGA | CGATTTCTTATCAGTTGGAGCTACCTCTAG | *Gossypium spp* |
| AMPL3259592 | ATTCATAGAGGTAGGAGCCAGTCATT | ACGTCTCTTCTTCATTTACTTTTGTCATCTT | *Gossypium spp* |
| AMPL3259599 | CGAGGCCTTTGCAACATTTTAGAA | AGTTACATGGGCTGGCACATAGG | *Gossypium spp* |
| AMPL3259604 | ATAGGAAGCTCTAGGTATGAGGTTACATTTT | TCAAACTATCTCGGGTCACTTTTACTTTTTC | *Gossypium spp* |
| AMPL3259606 | CCCTGATCTAGATTTGAACTTTGTTTTTCTT | CCTAGTCTCGGGAACGGGTAAAG | *Gossypium spp* |
| AMPL3259608 | AGAAAAATGACACTTTGAGGTTGTGTACTA | CGATCCAAATATGGTTGAAACACTCTATTCA | *Gossypium spp* |
| AMPL3259610 | AGAAAAAGAAATTGAAATGACAGTGGCAAT | CCGGGTGTAGGGTGTTACAAACA | *Gossypium spp* |
| AMPL3259613 | GTAGATGGGTGGTGCTGCTTAAG | CCCAAAATCAGCAAACCACCATTTTC | *Gossypium spp* |
| AMPL3259617 | ACGAAAGTGGTTTTCGAGTTTTGAAAC | CATTTCCATATTGGCTATCCGACTCTT | *Gossypium spp* |
| AMPL3259629 | TTTTTGGCAACACTTCAATGTCAAATTCT | TAGGGTAACTTTGAAAACAACTGAAGGTAGA | *Gossypium spp* |
| AMPL3259633 | CCCGAACCACCTACATTGTATGTT | CTTGTAATCCCGTAAAAGAATCAAAGTGACC | *Gossypium spp* |
| AMPL3259643 | CACACAGACACATTTTCAGATCAGAAAC | GCTCTATTTTAGTACAATTGGGAAGTGAGTT | *Gossypium spp* |
| AMPL3259646 | GGTTTGGTTTTCGATTGTATCGCTA | CGGCCATTACACGAAAGTATCATGC | *Gossypium spp* |
| AMPL3259657 | TACTTAGGAGAGCATTTGGGAGAAGAA | TCAATCTTGGAGGAAATCTACTTCAGAGTAG | *Gossypium spp* |
| AMPL3259659 | CCGAATACCTCATACTCCTTTCCTTT | AAGTGAGCAACTTACTGAAATGTAGAACTA | *Gossypium spp* |
| AMPL3259666 | TTCAACACTACTCCTCCTTTAGTCAAATGT | GAGATAGTAGAATGGCGAAAGTATGGTTCC | *Gossypium spp* |
| AMPL3259677 | TGTGTTTAGGGCCTATTTAGGACCTT | TCGCCAAATAAGTGGTGTCTCTAAAATTT | *Gossypium spp* |
| AMPL3259681 | CAAAGTAATAGACCAAAGATTGGAAATGCTA | TCTAGTTGTGCTTGGGCACTTATCG | *Gossypium spp* |
| AMPL3259686 | ACACATATCAGATAATCTTTTGGTGGACAAG | CCCTCTGTTGCAGATCTTGCAATA | *Gossypium spp* |
| AMPL3259687 | AACAATTACGAAGCACTCAAAGAAACTCT | CTAACTCTATGGTAGGTGTATCTGCAGTAG | *Gossypium spp* |
| AMPL3259689 | CCACGCTCGTGACCAATCTAATA | CGTGGTGAAATTGCACCTTGATTTCC | *Gossypium spp* |
| AMPL3259691 | CCATACCCTTCTAGACTTTCACTTAACTACT | AGACCCTTAGTCTACCCATTCTTACATTTAT | *Gossypium spp* |
| AMPL3259699 | GCCTCCCACAGTTGCTACAAAGC | GGAAGAATTGCACTATAATCTGCAGGTTC | *Gossypium spp* |
| AMPL3259700 | CAGTACCACAAAACTTCAGTAGTTGCT | CCGTCCTAAGAAATGAGATAGATTGTATGGG | *Gossypium spp* |
| AMPL3259715 | AACAAAAGCCAAAATTCATCTTTAACCGAA | CATCAATGGCAACTCACAAATTCACAA | *Gossypium spp* |
| AMPL3259717 | ATGTGTTTTGAGTGTTTTGGTCATAAGATGG | GTGTGCTAGCCTGTGTAACCTACT | *Gossypium spp* |
| AMPL3259718 | CACCCGAAAAACCTCCTACCTAG | ACTCGAGCCCATTTTCCATTATAAATGTAAT | *Gossypium spp* |
| AMPL3259719 | AATCGAATTTCCATCCAGGGTCTTT | TCCATAGCCTCTAACCAGTACTAAACTAAGA | *Gossypium spp* |
| AMPL3259720 | GCCAAAAATAGAGAAAATGGGCCAAAATAAG | CCCTTTTTCCGTCTCCCTTTTCTTTC | *Gossypium spp* |
| AMPL3259722 | GGACAATGTAAACAACAATATCATATGGCAT | GTTTTGGGCTCGAGGGTTCATATAAA | *Gossypium spp* |
| AMPL3259726 | GCCTACATCGCGAAAATAGTTAGTTTTA | CAGTCCAGTTCTTGCCAAGGAAG | *Gossypium spp* |
| AMPL3259733 | CCAATTCCTTCAAATAGAAGAATATCGCCAT | TCAAATAAATGCCGCAAATTTTTGCGA | *Gossypium spp* |
| AMPL3259737 | CCTATAGAGACAAAATATACCAAACCGGTAG | ACGCGTTTAGGGTTACGTAGAATTTAGT | *Gossypium spp* |
| AMPL3259739 | CTGGCTTTTGAGGGAGTTAAGCT | ACCACCACAATATCATACACCATCTAGTAA | *Gossypium spp* |
| AMPL3259746 | CCATCTCGAATTCGCATACTTACCTT | GTTACTACGTTAACCGGGAATTCTTTCG | *Gossypium spp* |
| AMPL3259748 | AGAAATCGAGAAGTGTAAGTTGTCTAGATTT | TAGGGTTCAGCCAAAGTAGAGAGAAA | *Gossypium spp* |
| AMPL3259752 | CTTCTTCCTCCTCCTATTCTTCCTCAT | TCTGAGTTGTTCAGGGTCCATATATAACATA | *Gossypium spp* |
| AMPL3259760 | CTAGTGATTAATCAGTGGAACTTCGTGTT | CCTTTTTGCCTTTTGCTGAAACCTC | *Gossypium spp* |
| AMPL3259762 | AAAGAGAAAAAGTACCAGTAATCACATCCAG | GGCCAAAAGCAATAGTAAAGCCTG | *Gossypium spp* |
| AMPL3259768 | CCTTTTCCTTGTAGATCGTTACCTTGTAG | ACATGGAAATATAGCATATCATGCCGAAC | *Gossypium spp* |
| AMPL3259778 | AATCATAATCAAGCCATAACTTGACACAGAA | TGTTTTGGGCATACGACCATGTT | *Gossypium spp* |
| AMPL3259779 | GTTGGGTGAACTCAAGTTTATTGAATTGGT | GCCTCACACATGACCTATCACACG | *Gossypium spp* |
| AMPL3259780 | TCAATCATTAAGGTGAAAAAGAGACTGAGTG | TTGCAATTATAGTCGAATGCACATTCCT | *Gossypium spp* |
| AMPL3259781 | AAAATTGACAATACCAAAGTCCGAATTGAGC | AAATCCCGGTAGTGTTCATTCAGTAAGT | *Gossypium spp* |
| AMPL3259782 | TGTTTTAATGTCTCGTAGGACCTTACATGG | AAGTGGAGGAGTTTGAATGCAATTTTAAGG | *Gossypium spp* |
| AMPL3259787 | CCCAAGTTCTAGAAAACTGTACTAGCTA | GCCGCTTAGGCTCGTTTAAAAGG | *Gossypium spp* |
| AMPL3259802 | GGGTGCAGAGTTCAAACAATCCA | CCCTGCCACGAAAATTTAGCTATAATATTTT | *Gossypium spp* |

| | | | |
|---|---|---|---|
| AMPL3259807 | GGTTCGACCACAACAGGGATTTA | CAGATCTAGTAATCGTTCGAAGGGTAAGT | *Gossypium spp* |
| AMPL3259818 | GAAGAACCCTAAAATGCACAATAACACAC | TATGGCGTATGCATCAAGGGTTTTTATT | *Gossypium spp* |
| AMPL3259821 | GCTCAAGATGTTCAATGCAAATATCAAGAA | ACAGTTTGGTTAACAAATGATGGAAATGTAA | *Gossypium spp* |
| AMPL3259823 | GGAATCATGCATATAATTGGTATTTGGCAT | AAAGGAAACTATACGAACATTGCTAAGTCAA | *Gossypium spp* |
| AMPL3259827 | TGGAGAAAAGATTAATAGCTCTCATCCTTTA | ACATGTATGTCGTGGAGATTTCACCA | *Gossypium spp* |
| AMPL3259831 | CCGACAGTTTGCTTCAGTTGTGG | CTTTCAACAATCAAATGCACAAATCTGGA | *Gossypium spp* |
| AMPL3259839 | GTTCAGGACCAACGTTGTGATGT | GATTGTTTGGAGCAATTGAAATAGGTATCAA | *Gossypium spp* |
| AMPL3259840 | ATTTGTTTGGAGCTAACTATCCTCAACAA | CATAGTTGCTCACAAGAGCCTTGTAG | *Gossypium spp* |
| AMPL3259843 | TTTTTCTGTTTTTAAAGTCCGAATCCGATT | AAAATTATATGGTCGCCAAAATCCAAACATG | *Gossypium spp* |
| AMPL3259847 | CAAATTAGTTCGTGGACGTTAATTCATGTA | CCCATACCTAGCTGCAAAAAGTTGAATAA | *Gossypium spp* |
| AMPL3259849 | GAAACTTCAAGTTTATATTGGACATGGGTTG | CATTTAGTAGCCATGCAAATTCAGTATTGAT | *Gossypium spp* |
| AMPL3259851 | CGTTTTATGAAAACGGAGACTCAAAAATGC | AAAGAAGAATCCATTCTAGGACAAATGGAAG | *Gossypium spp* |
| AMPL3259888 | GGACTCGACAGCCATCGCTATATAT | GAAGGTGGATTAGGATTCAGAGACTTGG | *Gossypium spp* |
| AMPL3259892 | TCGAAATAGACCTAAGGCTAGCCATT | GATCCCTTCTGTGGACATATCTCGC | *Gossypium spp* |
| AMPL3259897 | TCTAGTGGACAATACAGTAGGTTACTCACT | AGCCTTGACTTTATCTAGGTCGATCTTAA | *Gossypium spp* |
| AMPL3259901 | CAGTACTACAAGCGAAATAGGTGTACTATA | AAGTGTAAAAACACCTACCAATAAGTCTGTA | *Gossypium spp* |
| AMPL3259905 | CAGTGGGAACTGGTGTTACTTTCTT | AAAGATGAAGCTCACACCTTCAACTG | *Gossypium spp* |
| AMPL3259907 | CAATTCATACTCAATCAATTGGAGGACTCTA | TTCAGCAGTGGTTAACGGATTCTAATC | *Gossypium spp* |
| AMPL3259911 | GATTTTTATTGTGACTGTGACACAAGTAGCT | GCTAACACTTTGCCTTCTTGCATAAGC | *Gossypium spp* |
| AMPL3259920 | CTTGAGCTCCAAGAGTAAGAAATCGAA | GGCAAGTATACTAACAATGTTGTTGTTTGAT | *Gossypium spp* |
| AMPL3259926 | CCCTTTTTCCCTATGGATTCAAATCACC | GGACACTCAAGAGGGCTTATAACACG | *Gossypium spp* |
| AMPL3259933 | CTTTTTGAAAGTTGAGTGGCTGAAATGAA | GCGTATTTCCAACAACAAGCAAATTTTTT | *Gossypium spp* |
| AMPL3259935 | GTGCTTGGGAGATTGATTTTTGATAATGTT | TCCTCTAGAAGGATGAAAAGTTTTTCCAATA | *Gossypium spp* |
| AMPL3259938 | GTTCGAGCTGGTTATCGATTGTTTGA | ACTAAACCATGAAACCCAACATTATGGTAA | *Gossypium spp* |
| AMPL3259942 | CCAACTTATGTAATGAGTAATCTGTCTGGAT | GCTTTACTAGTGGCCTGGATTGAAATAGG | *Gossypium spp* |
| AMPL3259944 | GCAGGCATCATCTTCAACTCTCTA | TACTCTTTCTTAGGGTACAGGAAACGTC | *Gossypium spp* |
| AMPL3259955 | GATTTTCTCCTCCGTCTACTTTGAATACAT | CATCCTTTAGGCTCTCAACTTGATCTTTC | *Gossypium spp* |
| AMPL3259957 | CCCAAGAGTGTTGAGAACGAGTG | GTGCTTCTTTGTCTTCTCTGATAGAGTAAGT | *Gossypium spp* |
| AMPL3259959 | CAATAGCCCAAAGACAAACAAATACAGATCC | TCAGGCCCTTTTTATAGCCAAAATAGAAAA | *Gossypium spp* |
| AMPL3259964 | AAAAACGCAACAGGGAATCTAATTTCTC | GTTACTTCAAACAAAACAGTGAACTCCAAAT | *Gossypium spp* |
| AMPL3259968 | CTACCATGAATATTGGTTGTTGCTGTTCG | TTAGTGAAGAACGAACAAAACAAAACAAGTA | *Gossypium spp* |
| AMPL3259969 | CAAACACGACCCTCTGAACCATAT | CACGAAACAACTATGGACTCCAAAATTTAAC | *Gossypium spp* |
| AMPL3259971 | ATGAGCAAAAATTAGAGCAAGTTTCAGGA | TCGACTTTAATGATATTTTTCGAGTCGTTTT | *Gossypium spp* |
| AMPL3259974 | GATTTCTAGGGTGTGAGCAGTTATTGTT | CCAAATTCTAAGTAGAGGTACCCAATCAAAA | *Gossypium spp* |
| AMPL3259977 | AGGCTCAACAAGACAGAGAGTTCG | TAGCTATTGAATTGCACCGGTAGTTTAAAA | *Gossypium spp* |
| AMPL3259985 | TCCACTCTTTTCCTTAAACTTGCTTTCAT | GATCCACCATACCAATTATCAATTTGTCGC | *Gossypium spp* |
| AMPL3259987 | GAACCAACTTCATCACAAACTTTTCTTTCTT | CAAGTGATATCAAAGCTAGACTTCAGACATA | *Gossypium spp* |
| AMPL3259995 | ACCCAAGTAGCTATTACTATAGATCAGCATT | CTTGTCAGGTGAAAAAGTGGTTTACATTTG | *Gossypium spp* |
| AMPL3259998 | TCATGGTCGTGTTTGCTCTGAAG | CAGACTCCTATCATCGAATAAGCTCAGT | *Gossypium spp* |
| AMPL3259999 | TGACTTCTCTGGTAACATTTCCATGTGA | TTCTCAGCCTTGTGATTACGCTAAAG | *Gossypium spp* |
| AMPL3260003 | TTCGGTCCAAACCGTACACTTTT | GATTGAAATAGGGTCGGAGGAGAAAATTG | *Gossypium spp* |
| AMPL3260004 | TAACCGTGGGATATATGTCGGAACTG | AAGTAAGGTTGTGTACTCCTCCATAGTA | *Gossypium spp* |
| AMPL3260008 | GTGATGTGCACCAATGCAGATGG | GCGGTAGGCAAGTATTATATTAGCATACTCG | *Gossypium spp* |
| AMPL3260014 | AATTGAATTTTACTCGACCCTTCATATCCTA | GCTCTTGTTCTATGAACCTTATATTCGACTT | *Gossypium spp* |
| AMPL3260029 | TACGAGAATATGATCTCAAAGGACCCTTC | TAATGTGACGTGCCTTCATAACTTATGTT | *Gossypium spp* |
| AMPL3260032 | CATGGCGTCGTATAGGGTGTTTT | GAGCTTGGGCCTACCCATTAAAG | *Gossypium spp* |
| AMPL3260034 | CTTATACATGAAAGGGTTGGTCTGAAGTG | ATTTGCAATATGCCAAACAAGTGTCATA | *Gossypium spp* |
| AMPL3260035 | GGCTTAAACCAGGCCTTGAAGAA | AAGTATCTTCCAACCCTTGTTGAGTAAGC | *Gossypium spp* |
| AMPL3260038 | TTCTCTCTTTTTCGGCTATAAAAGCCAA | TTTAACTCCGATTTCGACAAAGCAAAGG | *Gossypium spp* |
| AMPL3260041 | CCTTTTCCACCCTCTAGTCCATCA | ATTTATAGAGTGGAGTATGAATCCCTTCCAA | *Gossypium spp* |
| AMPL3260046 | CATGAGTTTTTAATGTCCCAATTTAGTGCTT | AACATTGAACAATTGATGACATAATGGCAAC | *Gossypium spp* |
| AMPL3260047 | GTTATAACACATGGTGAATTGACTTTGGTTC | AACAATACCCAAAGGGTGCAAGTATT | *Gossypium spp* |
| AMPL3260050 | AGCCATGGAGCCAAATGTGTAGG | GTAGTAAGGGAATAGTGGAAAAGGTTGTTC | *Gossypium spp* |
| AMPL3260056 | CGCCGAACTTGTAAGTACTTCTGC | CCAGATTAGGCATACTGCCCATAATGG | *Gossypium spp* |
| AMPL3260057 | GAGAAGCGAAGGTTCATCTCACAT | CAAAAGAGAGGACATGGTCATGTATAGGG | *Gossypium spp* |
| AMPL3260060 | GTCTTATCCAAAATTGCTGCAATAGCTT | CGAGAATGAAAACCACCACCTTTGG | *Gossypium spp* |
| AMPL3260063 | GGGTTTGACAAGTGTCTTATAGGGAT | GGGATGTTACAATGACATTATCATTGCCA | *Gossypium spp* |
| AMPL3260065 | CTTTGATAGGCTGGTGTAACCTATTGAG | AGGTTTTCATATTTGAAAGGTAGCCAGATTT | *Gossypium spp* |
| AMPL3260068 | AATGCTCAAAATTACGAATCTAGGTGAGAAT | TCTTCTTCCCATTTTGTGGTTTCTCATTG | *Gossypium spp* |
| AMPL3260069 | CATCGTGAAGTTGAGTTTGGGATTAAGC | TAAGCTCCTCAAAATTGGTCGAATAGATCA | *Gossypium spp* |
| AMPL3260070 | CAGGTTCAATATTGGGCATCAACTCA | CCACCGATATGTAGTGTCAATTAAACGAGA | *Gossypium spp* |
| AMPL3260072 | CCAAGTGCTGGCCAGCATAATAG | ATGATCCCTACAGGCATAGGAACATAA | *Gossypium spp* |
| AMPL3260074 | GAAGGGTTTACCTTGCGAGCTAG | AAACCATTGCATCTGGTTTAAACGGC | *Gossypium spp* |
| AMPL3260077 | GTCTGTTAAATAGTATCTGCCCAAGATCAA | AACTTACACCTTCACTACTCTGTTGACT | *Gossypium spp* |
| AMPL3260083 | ACATGCCGAAGTCTTCGAATTATCCG | AAAAGAGGGTCAGTTTCACACCTTAATG | *Gossypium spp* |
| AMPL3260085 | TTAGACTTGCGTTGCCACATCTC | GATGTTAAATAGCTAAGCTTAGTCCCAGATC | *Gossypium spp* |
| AMPL3260088 | CTTTATTTGATGTCTTAGCTATCCCTTCCAG | TTGTTAACCGGTTAACAACTACCCAAA | *Gossypium spp* |
| AMPL3260089 | GCCTCAACTGACAAAACCTTAGGTT | ACCCACTTTACATTCGTCTCCTACTC | *Gossypium spp* |
| AMPL3260094 | ATTTTTGTGTGTTTCAAATCTTGATCCGAG | CAAAAAGATTGGTGGTGACTACCTTGT | *Gossypium spp* |
| AMPL3260097 | TGTACGTTTTATGCCCTTTTGAGACTC | CACAGTAGTAGTGTAAGTGGGAGTTTGG | *Gossypium spp* |
| AMPL3260100 | GTGTAGTCACACTGTAAAGTTTGTTAATCGG | AAAATGAGCCCAGGCCCATAATA | *Gossypium spp* |
| AMPL3260101 | CAAAGAAAGAAAAGGAGCCTAACATCGA | GTGGAGATTGGGCTCACATCAAA | *Gossypium spp* |
| AMPL3260103 | AGGTGGGTAAGGAGAAGAAGGTG | TGTTTGTATCGTTGCCACCCATTTAT | *Gossypium spp* |
| AMPL3260104 | CGACGGTGAGTGTTTGGCTAATC | CCCGAAACGAGTGAAGGTAGAAGA | *Gossypium spp* |
| AMPL3260109 | CGATCAAATCTCTTTATAAAGATCCGGATGAC | CTCCTTCTCCTCAGGCCAACG | *Gossypium spp* |
| AMPL3260120 | CGTTGATCCAGACAAATAATATGGGTGT | CCTAATGCCAAAGCCTTAAATCAAAATAGAT | *Gossypium spp* |
| AMPL3260125 | TAATGTGAAATGAATGTAAGGCCAGTCATTG | CTATTCGTTGCCGGAATAGGGTAC | *Gossypium spp* |
| AMPL3260130 | TTTTTACCTTGAACCCTGAACCCTAA | AAACCATAAATCCTAAACCCTCGAACTCAAA | *Gossypium spp* |
| AMPL3260131 | TGGATCGAAGATCTATCTTTGAAAAGACGG | TGAATTAAGTTCGTTAACACCTTCTGCTC | *Gossypium spp* |
| AMPL3260133 | CAAACTCCCACAAGCTGTCAAAA | GAATTATGAAACGTGAGTTTGATTCGGATAA | *Gossypium spp* |
| AMPL3260143 | CTTTATTGCATTGGTGGAGGTTGATAAGA | GGTGAGTGAAGAAAATGACAATAATCGATTC | *Gossypium spp* |

| | | | |
|---|---|---|---|
| AMPL3260159 | CAAAGAGCCTATAGACTGTCGTCAC | GAGTACGGAGGAAATTACCTTAATGGGG | *Gossypium spp* |
| AMPL3260171 | TTTGAATTTGGCAATCGTTCATTTGCT | TCTCAGGCATGCTTACATCATCCC | *Gossypium spp* |
| AMPL3260197 | AACTAAGAAGAAAGGTGCAGTTCGAC | ATTTTCCTGAGGATCCCTTCTACCTT | *Gossypium spp* |
| AMPL3260199 | TGTGAGACGCAATTGTCAACCAA | ATGTTTCCTTGGAACTACAGAGTGTTTTAA | *Gossypium spp* |
| AMPL3260204 | CCATTTCCCGGTCTTAAAAATTTCAGTTT | ATTAATTACCGCATAACGACCATCAATTGA | *Gossypium spp* |
| AMPL3260211 | CCTAAGCTTCCGCCAATGGAATA | CAAGACAAAGAGGAAATTCAACCGATTATTG | *Gossypium spp* |
| AMPL3260226 | TCATCAAAGCTTGGAACGTGGAAG | TGGAGAATGGATACTGTGTGTGGATTAT | *Gossypium spp* |
| AMPL3260243 | CGTATGTCCCGTGCACCTTAATT | CCTAAAACATGCATTGGAAGTGATTCAGG | *Gossypium spp* |
| AMPL3260246 | TATCGTTGGTAAAAGAGCTTTGGCATAA | GAAATTTCATGCCAACCAAATTCATGCA | *Gossypium spp* |
| AMPL3260248 | CGGAAAAGTGTGTTGGTTGGATGG | AGGTGGTAAGCTACTTGAGCTCAG | *Gossypium spp* |
| AMPL3260254 | ATGCTTTAGAGTGTCTCACCTTTTTAATGA | TTCTCGGAGTGTTTATGCTCAGTTTTAG | *Gossypium spp* |
| AMPL3260256 | CTCAGATTCGAATGATCGAAGAATCGC | TTGGAACTTTTGTATCTCCATCCAACATATA | *Gossypium spp* |
| AMPL3260260 | TCTTGGAGGATTTGGAAACAAATATGTTGAT | ACAAACTCAATTACCAAAATCATCTCTCGTC | *Gossypium spp* |
| AMPL3260265 | AAGGTGAGTTTATGTACTCTACTTTTTGGGT | GCATTTCATGCATTTCACACACACAC | *Gossypium spp* |
| AMPL3260267 | GGGTGGTTCCGACAATAGTAGGG | TCCGCATCAACGTCTATTCCTTGT | *Gossypium spp* |
| AMPL3260272 | GTTGTTTCCAACCAACTAGGACAAGC | GGAGCATTCTGCATTTGTCATCGA | *Gossypium spp* |
| AMPL3260274 | GTACTCTAAATGGTATGATGCAAATGCTCA | CATCGGTTCTTTCTTCGGATACGTC | *Gossypium spp* |
| AMPL3260278 | GATTGGCTAAAAGTTCATTCCCATGTTA | AGCCAATCATTCTATCATACTGGTCTTAAAC | *Gossypium spp* |
| AMPL3260287 | GATATCTTTAAGAGAGGTGAGTGTATGCAGT | AGTGTCAAAGTTTGGTTTCATTGACAGC | *Gossypium spp* |
| AMPL3260290 | GAGCTAGTTAGACAAGTCTAAAAGAGATGGA | TATTGCTGAACAAGAAACAACAAGCACTT | *Gossypium spp* |
| AMPL3260295 | TCAAGGGACATGGATGTGTAATGGG | TCCAATCACATTACTATGTGTCCCTTTTTT | *Gossypium spp* |
| AMPL3260299 | CTTTGCTTCAAATCCGATTGTTTGCT | CTTTGGTGTTTCACAGGTCATAATCCT | *Gossypium spp* |
| AMPL3260301 | GGTTAATTGATAAATGTTGTGGCAAACAGAT | ATACGGTGCTATAGAGATAAGAGTAGTTCCC | *Gossypium spp* |
| AMPL3260305 | AAAAGGGAGCTCAAGAACTCTTACCA | GCTTCTACATCGCTTTGTTCCAAGC | *Gossypium spp* |
| AMPL3260308 | GTAACATCCGATATCCGAACTCGTTG | CCTCAATGTTGTCATTTTCGTAAAATGGGA | *Gossypium spp* |
| AMPL3260315 | AAACCTAAGGAGACAAGAAACTGTTTCAA | ACATTTAGTCCAATGGTTAAATCTGCTACTA | *Gossypium spp* |
| AMPL3260319 | CGTGCAAATTTGGATTGTGCTGTT | CAACCCAGGAAGCTCATTAGGAAAAA | *Gossypium spp* |
| AMPL3260324 | AGGTAACTTCTGGATTATTGCAGCTAGT | TCGTGATGTAAACTGTGGATCTTTATCAGA | *Gossypium spp* |
| AMPL3260343 | CCACCCTAAGGGTGAGACAAAAA | AGCCACATTGCAGAGTCTAATAAGGT | *Gossypium spp* |
| AMPL3260348 | TCAGTGCTCTCTGTTTAGCATATTGGT | GCCCTTTTTATGAAATTGAACACTCAAATGA | *Gossypium spp* |
| AMPL3260349 | CTAAATGGCTCTAGCTTTAAAAACCAAACTC | AAGGAAAATGAAGACTAGCTTAACGGAATC | *Gossypium spp* |
| AMPL3260355 | GCTGCATTAAAGTTGCAAACTCGAA | GATGCATCCCAATCACTATACAACTACTCC | *Gossypium spp* |
| AMPL3260364 | CCATCGTTCTCTGGGAAGATTATCGA | AAGCGAATGATCTCGTTAATCTCGTTTAA | *Gossypium spp* |
| AMPL3260371 | AGGTATATTATCGACCAATTTTCCCACAT | GGATCCGGCAATTCGGACAATAATT | *Gossypium spp* |
| AMPL3260374 | ATACATACAAGAGCGATTAGTGAGTTGTCG | CCTAAGTCTCGAACCTCATAATTAATGGTTA | *Gossypium spp* |
| AMPL3260377 | TTTAATGGAGATAAGCTTTGTGGTGGAAA | TCCAATAATTGTTGGGTTCAAATCTTCTCAT | *Gossypium spp* |
| AMPL3260380 | AGGTTGTTGGTATGTTTTAGGATAGCAAAA | GCATTTACAAAGAAATAGGTGGCCAAAATTA | *Gossypium spp* |
| AMPL3260383 | TGGCTGGAGCTAGCAACATTTTT | ATCCTCCTGCATGGCAACAATAC | *Gossypium spp* |
| AMPL3260385 | TGAAACTCATCTCTAGATGTTGCTTTTGTT | CCGTTTCTAAGCACTTAGTGCACTTG | *Gossypium spp* |
| AMPL3260387 | TCTATTGTTTAGCCAATTTCTAGTTGTGGTT | ATTCAAACAACCGCAACTATACTAAATTGGT | *Gossypium spp* |
| AMPL3260388 | CGTTTTGGGAAAGGACTGAACGAA | GGTTCCAAATTACTAGGTTTTAGTCCATCTC | *Gossypium spp* |
| AMPL3260390 | CCTGATGCTCCAAACCACCTATT | TCCCATTTCGTGGTTTCCTATTATCATCA | *Gossypium spp* |
| AMPL3260401 | ACATCAAATAAAATACCACACACCATGGAA | TAGCTTGAAAGTCCTCCCAATTCATTAGC | *Gossypium spp* |
| AMPL3260402 | CTTAATTAGTTGTTGCCCAAGGGTTAGG | ACTTTTGCTTTTTGAAATTCCAGTGCAC | *Gossypium spp* |
| AMPL3260441 | TAGCAAAATTTACCGAGAGCTTAAAGGAG | GTTGTAAGGTACGAGCGTGAAAAAA | *Gossypium spp* |
| AMPL3260442 | GGTATGAATTGGTTGGTGGAGCATC | CTCGACAACTTATCAGAAAAGACATCTGAAA | *Gossypium spp* |
| AMPL3260448 | AATTTACTCTCACAGCCTCTATTTCAAGTCT | GGGCGTGTGACCTAAGTAAGTGT | *Gossypium spp* |
| AMPL3260450 | GAGATGATAGTTTAAGGGAAGAGTAAAGCTT | TACAATGCTCGTAGCCTAAATTCCTCA | *Gossypium spp* |
| AMPL3260456 | CTAATGCAACGATTGCGTATAACAGATATGT | CGTGTCTTCTCTACACTTAATTTTAGCACTT | *Gossypium spp* |
| AMPL3260460 | CGTTCTATCTGTCCATTTGTCTATGGAT | GTGACACCAAGAAAGAAAGACTCGATC | *Gossypium spp* |
| AMPL3260463 | ACTTGTTTTGTGGTGTATTTGATGATGTTTA | CGATCGTGTCCCAGATTATGTCATAAACT | *Gossypium spp* |
| AMPL3260470 | GGGATCCTTTTATGAAGAGGTGCATG | TGCTCATCTTCAAAGTACCAAGGTTCT | *Gossypium spp* |
| AMPL3260471 | CATGCCTAGCCAAGATGAGGATG | TGCTCATTATTGCTCTATGATCGTTCATATA | *Gossypium spp* |
| AMPL3260475 | TGCATATACTTCTGATTGTGGTTGCAC | TGCCAGTATTTGTTCTTATACTCCTTGTGG | *Gossypium spp* |
| AMPL3260479 | TATCGTTTGAAGGACTCGTTACAAGTGC | CGTGGAAATTGCAAATTCAAGGTGTAGG | *Gossypium spp* |
| AMPL3260482 | GACACGGTAGCAGCTGAAGATTA | CACATGTAGTGGTGCTTGTGAAATAAACA | *Gossypium spp* |
| AMPL3260487 | GACCCGATTCTAAGAACTATTGGAGAAGC | TATGATGCCACCACAAAAGAGCAA | *Gossypium spp* |
| AMPL3260489 | GGTCATACGACCATGTGTCCTATA | ACAATCACAAACACCAAAGCAATAACAATAT | *Gossypium spp* |
| AMPL3260500 | CTTTTTGTTGATGAACACGGTACTCTA | AGGAGATCAACCAGTGTTCAATACTTTGT | *Gossypium spp* |
| AMPL3260507 | GGCACTAGAAGTTTCCAAACCCAT | GCTACACTGGACACCAACTTGAA | *Gossypium spp* |
| AMPL3260516 | AAGAAATTTAGTCCTCTCCCTCAGTATGAT | CGACTTCCTAAAAGACATTTCGACTGATTC | *Gossypium spp* |
| AMPL3260522 | GAAATTTTAGTCTTGATGCAACCGACAAT | CCCTGTCCTTTATACAAGTTGTGGATTTAGT | *Gossypium spp* |
| AMPL3260528 | GGACAGAAAACTCGGAATAGTCCAC | AGCTATACATTTCCATCTAATGAGCCTTTTT | *Gossypium spp* |
| AMPL3260532 | GAGCAAGCTACAATTCAACATCAACTG | GGTTCTCTTCTCAACTAGAACAAGTGTATA | *Gossypium spp* |
| AMPL3260538 | AATATCCCTAGTGCCAATAACCATTGTACC | GTAGGGTCATGTTGGGCTATTATCTTAGT | *Gossypium spp* |
| AMPL3260540 | AAAAATTTTAAATGCGTCCCTCTTTGTCAG | ATGGACCCACTTTGACTATTGGTCA | *Gossypium spp* |
| AMPL3260541 | CAAGTATGAAAGATAGTTGGTCAGCATCTG | TCAAAAAGCAAATTAAGAGAGCTTTCTGTTT | *Gossypium spp* |
| AMPL3260548 | GGTATTGTAGATCTCTATGACCATTGTCCAT | CCTCAGTAATAAAACCCTATGCTAGTTGCC | *Gossypium spp* |
| AMPL3260552 | AGTCCCTGTACGTTACTTAAAAGAACAAAT | TCTATGAGAGTTGATTGTTTTGTTGCCTT | *Gossypium spp* |
| AMPL3260558 | AATTTAGCTACACGACACTTCGAACTC | AGGACGGTCGATTGCATGAAAAT | *Gossypium spp* |
| AMPL3260567 | CATTTTCCCATAACTGCCATTTTTGTATCAT | GAGTGCTATAAATGCCACAAATTGGGG | *Gossypium spp* |
| AMPL3260571 | AAGGTTACACGGCCTAAAACACAA | CGGATTAAGGTTACGGAGCACTAC | *Gossypium spp* |
| AMPL3260580 | CGGGCTAGTAAGAGCATCCATCA | CCGATCAGACTATTTAGCTTAGATTCTGCC | *Gossypium spp* |
| AMPL3260589 | GCACTAGAGTAGCCGGAGATGAG | TAGACAGTAACGAAGACAAATGTGATGTTTG | *Gossypium spp* |
| AMPL3260598 | GATGGAAGAAAAAGATGCACTCCAAG | GAATGACCACCAGCCATTTGATAATTATTGA | *Gossypium spp* |
| AMPL3260601 | GTGGTCGTGGAGTGCATTTAAAA | CAGATCATTTTGTCAACTTAACCACCAACT | *Gossypium spp* |
| AMPL3260603 | AGACAGTGAGAGAAATCAACACACAATA | GCTGGCTATTTGTGTATCTTATACTTGGTAT | *Gossypium spp* |
| AMPL3260613 | GCTGTAAAAACCTTTTAAGGCATCGAA | CACGGTGAATATTGTGAATCATAAGGGATAA | *Gossypium spp* |
| AMPL3260614 | GTACTAACATTAGGCGCAGACATAACAT | TCAACAATGTACTCATTGAGTTGGGTTG | *Gossypium spp* |
| AMPL3260618 | GAGGGCATATAACCCTCTTGCTG | GGATCGGCAATTGGTGCAGTATG | *Gossypium spp* |
| AMPL3260621 | TTTCGTGTGACATATCAAACATAATGTTGGA | GCAATAAATCATGTAGATGCAAGTATTGGTA | *Gossypium spp* |

| | | | |
|---|---|---|---|
| AMPL3260625 | ATGCACCTTAAATAGCTTTCTAGGCAT | GTCAATAGAGTCACAAGTATGGGCATT | *Gossypium spp* |
| AMPL3260626 | GTAGGATTACCATACCATACCTATTGTGCA | CTATGCAAGTCAGAATGGATCGTGAG | *Gossypium spp* |
| AMPL3260630 | AATTTTGTGACATTGAGTGATGCGAAT | GGTAGGGTTGGCCGTTTGAATAGT | *Gossypium spp* |
| AMPL3260638 | AAAAATGATGCCAACAATCTATAATCATGAGG | AAAACACTATACCTACTTCCTTTTCTAGCAAAAG | *Gossypium spp* |
| AMPL3260640 | GAGCTACGGTTTAGCTTCCGTAG | CGAGGGCTCATATAAGGAACAATATGATTGA | *Gossypium spp* |
| AMPL3260642 | CTAAGGAAGCGCCTCAGTTTTGT | AACAATGGAAGGTATTGTCTCGCTTG | *Gossypium spp* |
| AMPL3260643 | TGTAATGAATGTCACGAGTATCATTTGATC | GAGATCCACTTGAAATGGTCTAAACTCTTAG | *Gossypium spp* |
| AMPL3260645 | GCAACAGTGATTCTAGTCACAGGAA | CGAGTGACAAAGTTTAGGGAACGATAGA | *Gossypium spp* |
| AMPL3260657 | CAATTGAATCCACTCATCTAAACCCATTGC | TACTTCATTCACCCTTCTCTTGATAGATGAC | *Gossypium spp* |
| AMPL3260665 | CAACAATAATTTCTGTACACGACATTCACAA | GTGCCGATGCAAGTCTCTAGTTT | *Gossypium spp* |
| AMPL3260670 | AAACCATCGTATCTTGGGTTCACTACC | CATTAATGGGTTGATGGATGCTTCTTGG | *Gossypium spp* |
| AMPL3260672 | AAGGTCTATCACTAATTTTCAAGCATCCAAT | CAAATGAGCTGAATTGCATATTTGAGGATC | *Gossypium spp* |
| AMPL3260675 | TCTTAATCATTCTGCTTCTTTAGATGCAGTA | CCCAATGTTACCATAATTCAAAATTTCCCTT | *Gossypium spp* |
| AMPL3260679 | GCCGTTTTCCTGTGGATCCAAAT | TCTGCTTTAAATTAGGGTTTGTTAGGGTTTT | *Gossypium spp* |
| AMPL3260680 | TCCAATTTGCCGGTTAAATAATAGGAAAACG | GTTGACCAAGTTTGCTCATTTCCTTCT | *Gossypium spp* |
| AMPL3260683 | TGATGGGTAATTGAATACGGGTTTTAGGA | GATTTGTCATTTTTCGAACGTTAAGTGACTA | *Gossypium spp* |
| AMPL3260688 | TACCAGTGAGATTGTTAAATGGGTTGACT | GGACTTTAAATGTTGTCGGATCTCCATAG | *Gossypium spp* |
| AMPL3260698 | GGTTTTTCAACCAAGGAAGCTTCTAGA | CGAGACATTACCACATGTCGAACATG | *Gossypium spp* |
| AMPL3260701 | TAAGTTGTGCAAAACAATTTAGCAACCTA | GCGTTTTGCTTAAATTGACTCACATCAATAT | *Gossypium spp* |
| AMPL3260707 | AATTGATGTCCTAAAAGATTGATGGAAGGAA | TAGCCTGCATCATTCAGAATCTGTTTA | *Gossypium spp* |
| AMPL3260712 | GAAAGCCTAATGAGGAAAAGGAAAAAGTGA | CTAAAAACCTGTTTTGCTATAATGTGGTGTG | *Gossypium spp* |
| AMPL3260716 | TCGTACCTTTTAGTCCAAAAAGAAATGGAC | GCTTGGCTTTTTCCCATTTAAGTTCTAGA | *Gossypium spp* |
| AMPL3260722 | TACAACTTTGTGCATGATTTGAATTGAATGAT | AAGCTTTTCATACAAGCTAACAAGTTGTCA | *Gossypium spp* |
| AMPL3260725 | CGCACTTGTCATGCATCATAAATAACCT | GACCGTTTGACTATGTATAGACACTTCTTA | *Gossypium spp* |
| AMPL3260727 | CCGCAATCATTTAGCGCGTATTT | GAAAGATCAATTAACGATAGAGGCCTCATG | *Gossypium spp* |
| AMPL3260729 | GGTACGATCATAGTCCTTAAGCAACTC | TGATGCATCTCATGCCTATCTTGGT | *Gossypium spp* |
| AMPL3260730 | AATTCTCATTTCAATGCCACCTAGACAA | CAAGTATGAACATTCACCCTTGAATCAATAA | *Gossypium spp* |
| AMPL3260789 | CCCTCTTTTCAATCTTAACCCTAACTACAA | GGATTCTCAAATTTTACAGGGTTCGAAACT | *Gossypium spp* |
| AMPL3260793 | CCTTATAGATCAAGATTCAATGTGCTGGAAT | TGGAGATTTTGAGGCTCCAAAGTTTTG | *Gossypium spp* |
| AMPL3260805 | CATCTCTAACGTCGACAGTTGAGATT | ACTTTAATGCATGGATTAAGAGTGGTCATC | *Gossypium spp* |
| AMPL3260816 | GGAAGACAATGTTCCCTTAGATTTATCCTG | CGATTCTTGTTTCATTCACCTGTTCTGG | *Gossypium spp* |
| AMPL3260822 | GTAAGGATGGCGTTTAATTCTTTTCCGA | CGGGCTAATCCCAGAAAACTTCTAACA | *Gossypium spp* |
| AMPL3260829 | TACGTGGGTTAACTAGTCAGTTAGTTAGTT | AAAAGGTCAAAGAGAGCCTCATTGAG | *Gossypium spp* |
| AMPL3260832 | TAGTCGATTTCAACTTTGGTAAGCCAA | CATTTGGAACATCCTCAAGCTCAATAAATA | *Gossypium spp* |
| AMPL3260833 | CTCCCGAGAGAGTTGTCAACATC | AGGGAATTTGTGTTGGTGTCTGTC | *Gossypium spp* |
| AMPL3260848 | GTTCGTTGGAAGCTAATGAATCCCT | GTCTATGCTCAATTTTATCTCCGAGCTTT | *Gossypium spp* |
| AMPL3260860 | CGGTGGCGACTCCAATTTCTATT | CATCCACTAACCATTTCCTATCCAACAAC | *Gossypium spp* |
| AMPL3260864 | GAGTTATGTTTCTATGATGGTTTGTTTGCAT | GGTTTGATTGGTTTTAGGCTGTTGTTG | *Gossypium spp* |
| AMPL3260867 | AAAGAGAGAGGAATCAACAATGGCAG | AGCCCTAACACATCACTAAAACATAGGT | *Gossypium spp* |
| AMPL3260871 | AACTGATCTCCCATTGAGCATTAAACAA | CAACCCTTTTTCCAAGTGAATCACAGC | *Gossypium spp* |
| AMPL3260873 | CGAAAAACGGTTTTAGAAATGATACGGTT | ACGTCTGAACATTTAATGAAACTTTATTAAGCAT | *Gossypium spp* |
| AMPL3260876 | GAAAAGGAACACCTAAATAATTGCCAAGATT | GGAACCAATTTGGTTATCTCGTACTGTC | *Gossypium spp* |
| AMPL3260889 | CAATAAGGGAATTGGCCTCCCTATG | ACGTTCTTTCTCATCAATAGTTAAGTTCGTT | *Gossypium spp* |
| AMPL3260892 | CATATTAATTGCTTGTGTGCGTTGAGTA | CAAAGTAGAAGGCCCTAGAGAAGATGC | *Gossypium spp* |
| AMPL3260900 | AAAAAGGCTTGAAATTGTGATCAAAGAAAAA | CAAAACTAATTGACTTTCAGTACGCATCAC | *Gossypium spp* |
| AMPL3260903 | GCGTCACCTCCTTTAGACCTACG | GAAAGCACTGTACAAGCTTGATCTGC | *Gossypium spp* |
| AMPL3260910 | TGAATGCATGGTGAGAAATTTTTCTTCCC | GTCATTACAACCCTTCCTGAAAAATTTGAGT | *Gossypium spp* |
| AMPL3260919 | AGATTGATGAACTATGTTTTGAGACCCTTC | GTTATGGGTTTCGACCATTCTTGTATGA | *Gossypium spp* |
| AMPL3260932 | TGTTGTACTCCCTATGACGTCCATC | CCAACTCATCCTTTACTGATCGGTC | *Gossypium spp* |
| AMPL3260939 | GAATAATGTTTGTGTAGGTAAGCAAATGCAT | GCTCGCATCCGTTTAACTAGTCATATTAATT | *Gossypium spp* |
| AMPL3260943 | AGGGTGGCAAATTATGTTACCACAG | TGTTCCAATTCGGATCCATAGAATGTACG | *Gossypium spp* |
| AMPL3260946 | ATTGTTCGTTCATGCGGTCATTCT | GAATCAGATTAGTCAGCTTGCCAAATTCG | *Gossypium spp* |
| AMPL3260948 | GCTTTTCTCTTTGGGTTTGGAAGACT | AAGGATAAGGCTTGGGATAAGTTGTATCG | *Gossypium spp* |
| AMPL3260950 | TGTGTACAAGATTGGTCAAGAAGATGGA | CTCAACACTGAGGCAGATCGTATCA | *Gossypium spp* |
| AMPL3260952 | TAAGGTGGCTAATTTCTCGTTGTTAGTGA | GCCCAAAATATGTAGGTTATCAAGGAAGATA | *Gossypium spp* |
| AMPL3260955 | GAGTTTCTTGTGGCTAAAATTGCTCAG | ACCAAATCTAACAAATACTACCTCACACAAC | *Gossypium spp* |
| AMPL3260961 | GAATTCTACGTGCTGGATACGACAA | TTCGGGAGTGTGTTGAATCGATTATTG | *Gossypium spp* |
| AMPL3260978 | TCCGCAAATCGATGGGAAAACTG | CCAAGTCGTTCTTTTCTTTCCAATCCT | *Gossypium spp* |
| AMPL3260988 | CTAAGTTACAGTACCCGTGCGAATT | ATCATTCATTCTATGCAAAGCCAGAAAATCA | *Gossypium spp* |
| AMPL3260999 | GCCCACAACATGGAGCTTATTCT | GATCGACTCGGGTCATAAGACGT | *Gossypium spp* |
| AMPL3261003 | CTTACTCCATATTTCCTTAACAGATGTGCAA | GAAGCGATGGAGGAGTACACTGT | *Gossypium spp* |
| AMPL3261028 | GGTTCATGTAAATGGAACACAACCATTG | CATTGAGTCGATCTACTAAAAGTAGCATGAC | *Gossypium spp* |
| AMPL3261056 | TATCCCAATTGTGATTTTCTACATGCACTG | CTTAACTTAGAGGTAAGCAGGTGTGTATCC | *Gossypium spp* |
| AMPL3261057 | TCACTTGACAGACTGTTCATGCAC | AGCTAACTCTTGCATGAATACTGTTCATA | *Gossypium spp* |
| AMPL3261063 | TTTTGACAAAACTCAGAGAATCGAGACTT | CATCATCTGTCTGTCACGCTTTTTGT | *Gossypium spp* |
| AMPL3261068 | CATCAGATGGAGCCAGCACTAAT | TGTAAGCTCGTATAAGACCTGGGAATG | *Gossypium spp* |
| AMPL3261070 | GCAATAGAAGCATGTTTACCAATGCTG | TGGAAGAGAGAAATTGGCAGCTATGC | *Gossypium spp* |
| AMPL3261080 | ATGGTTCAATGAGTTCAAACTTACACAAC | TTACCTAAACCCTCTCACATTTCGAACG | *Gossypium spp* |
| AMPL3261083 | AAAAATTTCAATGTGCAAGAGACAAACGC | TGCTTTGGATGTATTTTTGTATGCTTAGCT | *Gossypium spp* |
| AMPL3261091 | CCGGTTAACAGCAACAATGCTAG | TTTTTGGCACTAGCAGTTTAGTTGTTATTG | *Gossypium spp* |
| AMPL3261093 | GGATAAGTGACTTCCCAAATGAAACCTA | GCTTGAAAAGGTCACTCAAATCTTCTCG | *Gossypium spp* |
| AMPL3261095 | GGTCGAACCGGTTGAATTAAAAATCAA | GCCTAATGCATCACTTGAAAAGCTCG | *Gossypium spp* |
| AMPL3261097 | CCCTTCATGAGTATTGGGAGAGATTC | TTCAAGAAAGAGACATTTACCTTGTTCACTT | *Gossypium spp* |
| AMPL3261103 | CCAGGTGTGTAATGAAAATGTGAGAAAAC | CCCTAGGGAAAATTTTAGCATTTTGGACC | *Gossypium spp* |
| AMPL3261111 | CGAGAAAAGCTCCTTAAGTATACTGGGAA | GCCAAATGTCGAATCCTAAAAAGTTCTTTAC | *Gossypium spp* |
| AMPL3261114 | CGGAGTACTTACCTACTCTGTTGCAA | CCGTCCTCAGAGCTCAAATCATGT | *Gossypium spp* |
| AMPL3261115 | GGAGTATTAGCTCCAAAAGGTTTCACTG | CAAGACTTTGAAAACCAAGAGCTTGAAT | *Gossypium spp* |
| AMPL3261116 | CTTGTAGAGAATGGGATATTGTCCCAA | GTGGGTTGGGTATCCCTAAATCCT | *Gossypium spp* |
| AMPL3261138 | CTCGATTTGTGAAGGATTATTTGTGAAGGA | CCTAGGGTCCATATTTCATTCTGATCCC | *Gossypium spp* |
| AMPL3261142 | ATAGCAGACCATCCAATGGCTTACA | TGGCTTGCGAATTCTTTCCTTACTTT | *Gossypium spp* |
| AMPL3261145 | GGGTAGGTTTGGATGGGAGGTAC | TCTCCAACTTTTTGAGCCCAAAGAAC | *Gossypium spp* |

| | | | |
|---|---|---|---|
| AMPL3261147 | TAAAATGTACGAATCGATATCGCCACTTC | GGTACTTGATGCATAAATTGATAGGTTTGGT | *Gossypium spp* |
| AMPL3261151 | GCTCGTGAATTTTGGGTCATTACAGATT | TGTGTCTGGTTAATGTGGTTATAAACACATG | *Gossypium spp* |
| AMPL3261157 | CAAAAAGTCCAACTTACCTACCCATTTG | TTGCAGAATCCCACAAAGCACAAATA | *Gossypium spp* |
| AMPL3261164 | TTCTCTAAATGCCAAACCTACATGGTT | AGTTGAGTGGTAAGTAGTTTTGGAGTTTTT | *Gossypium spp* |
| AMPL3261168 | CTTAATATTGTTGATGCTTGGTTCGGAATT | CGTCGGAATTTACTACCAAACAAATTAACTC | *Gossypium spp* |
| AMPL3261169 | GCATCACTAGTAAAGACAATCTATTGACCAG | CCAATTGTATCAAGTATTACTGGAACCTCTT | *Gossypium spp* |
| AMPL3261172 | TGAGCTTTCTTTTGGCTAGAGTACAGA | AACCCAAATGATTCTATGGAGAATCCAAAG | *Gossypium spp* |
| AMPL3261174 | GAGTTTCATCGTCCTTGCTACTGAT | GTCATCTCGAGTCTATGTCATTTCAGTTTAA | *Gossypium spp* |
| AMPL3261178 | TGGACGCAGCCAACATAAGTTATT | TTGCAGGACGAACATAAAATACCAGATAAC | *Gossypium spp* |
| AMPL3261184 | CAAATTATTGATGTTCCCTGAGGAAATTCGC | TGTTCCGACAACAGTTTAGGGTTAGG | *Gossypium spp* |
| AMPL3261194 | AGTCATACTTAAACTTTGCCCTTTAAGGAA | GCTTGAATGGTGTAAGTCAATTTGGTACTAT | *Gossypium spp* |
| AMPL3261197 | AAAAATCCGGTTTGAAAACAGTTTTCCT | GACTTGGGTGTTTTTACTGTTGAATCCT | *Gossypium spp* |
| AMPL3261200 | TCTTGCACAGAGAAACAAACTCCAG | CCCTAATCGATCATTAACAATGAACAAGAAC | *Gossypium spp* |
| AMPL3261202 | AGAAATTACATAAGTGGTGTGTGAAGGTC | AGAAATTTCCCACAACCTCAGAATTAAAGAA | *Gossypium spp* |
| AMPL3261205 | CGTAGATTTTATGTGCTCGCATGTTTT | TAAGTTTCCAATGTGCCCTCTTAAAGTG | *Gossypium spp* |
| AMPL3261206 | CACATCGTTCCATCTGATTCATGCA | GGTTCGAGTTCATATTGATGTATGGTTACCT | *Gossypium spp* |
| AMPL3261210 | CTTTAGGTGTGATCGGACACGTAC | AACTCGTAATTTTGAAAATTAAGACAACGTG | *Gossypium spp* |
| AMPL3261216 | CCCAATCTCAGGTCTGTTAGAGGTA | TCTGATAGTACGGTAATGCTCCGTAATC | *Gossypium spp* |
| AMPL3261221 | TGCTTTTCACCTTAAAAATCCACTTACAACC | CAATTGTTCCTCTACTGCCAATTGCT | *Gossypium spp* |
| AMPL3261226 | GTTATTAGCATGTCATGTTGCATTGCATAG | GAGATAGCCAAATGTTTAGGAGTTCCTC | *Gossypium spp* |
| AMPL3261232 | CCTTTATTGCAGATCATTTAAGGCCCTAA | ATTGTGGAAGATATGGTCATATGAAGGAAGC | *Gossypium spp* |
| AMPL3261240 | CCCAAACCATCATTCATCGATTTTGAT | GCTAACACACCAATTACTTGTTCTAATGAAT | *Gossypium spp* |
| AMPL3261241 | CACCTGAAGATCAACTATCCCAATGAAA | CGCAGGAAGAACGTGTGAATTTTATATATTC | *Gossypium spp* |
| AMPL3261244 | GGTGGAAACCACAGCTTCTCTAC | CCTTTTGTTTGGCATCTCCATTTCATCT | *Gossypium spp* |
| AMPL3261246 | TTCGAATATTGCGTTATTTTGGAAAGCTT | ACCGATGGACAATTCACACATGAA | *Gossypium spp* |
| AMPL3261248 | CAAAGTTGAAGTTTTGTTTGGTTGAGTCC | TTCATGCATAGTTCTTGAAACCTTTTGATTATAA | *Gossypium spp* |
| AMPL3261250 | CTTTTGAAAATCACCTGGCAACCAAC | CCTCGCCGTGAATACTCTTAGAAGA | *Gossypium spp* |
| AMPL3261254 | TTTAAAGTGTTTGGTTTTGCCATGAAAAACT | GGGTCAGTTGAGGACATGAGAAGC | *Gossypium spp* |
| AMPL3261260 | CTCTTAGAGGTCCAGCTTTTACACCC | GAGGTCGTTTTCTGGATTGGTCTC | *Gossypium spp* |
| AMPL3261262 | CATTTTACTTCCAATCGCTTCATTCAAATGT | GATTTGTGTATAAGGCTTAGGAGCAGATG | *Gossypium spp* |
| AMPL3261263 | TGTTTGTACAGTTGGGCTTTACAGTG | CTCTAAAGAAGAGTCAGGCCATCTAAAATAC | *Gossypium spp* |
| AMPL3261269 | ACTAACGATTGGGTCACTTTGAGTACA | GAATTGGACCCTTTTGAAGACCTTTCT | *Gossypium spp* |
| AMPL3261271 | CCGTCGATGGATGGAACAGTTTT | CCAGCTGGTTCACCATCAGATTCT | *Gossypium spp* |
| AMPL3261280 | AATATGAGTGGAACTATGGAAGAGCAATCG | GACACGTTTTCCCTTTAAGTTGCATAT | *Gossypium spp* |
| AMPL3261283 | AGGCTTTTATAGCCGATTTCAATGACTC | ATTAAACCCAAACAGCCCAAAACTAACCCTG | *Gossypium spp* |
| AMPL3261288 | GATGGAGCTGCTCTTGGTACTTT | ATTTCAAGATGCCTGAGCTTGAGAG | *Gossypium spp* |
| AMPL3261290 | TTTGTACTATGCGCACGATTGTAGAT | GCCCATAGTTGTATTTGAGGAATTGTTGAA | *Gossypium spp* |
| AMPL3261291 | GATCCAAATTAAGTTTCTTGAAAGCCCAAT | GCAAGCAAAATTTATCTGTAAAAGGGATGTA | *Gossypium spp* |
| AMPL3261298 | CATAATAGCAGCCAGAGACCCAAA | CTGGCCTCGTTATACTTTCTTGAACC | *Gossypium spp* |
| AMPL3261303 | GATCGCAAAAACATATTCGTTTTTCAAAGTA | AGTTATAAAACTCACTCACTGTTTGTCTGTT | *Gossypium spp* |
| AMPL3261306 | TCAAATAATGGAACAAGGATGTCTATGGTTT | AATACGATTAAATTTCCTCCTCATAAGCGTA | *Gossypium spp* |
| AMPL3261307 | AAAATTATAGAATGTGACTCACCGGGAATC | GGGACTAACTTATTTATGGCTTTAACCTATC | *Gossypium spp* |
| AMPL3261313 | GCATACCAGGTTACTTAAGAAGTTGCTA | CGTGAAAAGTTGTCTAGAATCCTAACTCTAA | *Gossypium spp* |
| AMPL3261314 | AATACTTGCAGCACTGAGATAGCAAA | TCTTCTCTTCCCTCTCCTCTTGTGG | *Gossypium spp* |
| AMPL3261318 | TGAATTCAATATTTGGATCGGATTGAACGT | ACAAGAGCTAATCCATCCAAACACAC | *Gossypium spp* |
| AMPL3261325 | GTTATCCGGATTTAAACTGAGTGTCAAATT | CCCATCTGAAATGAATCTCCAAAGTGC | *Gossypium spp* |
| AMPL3261326 | CATCAGGCAATTTTTGGGTCCAAA | GAAAAGAATTGAACAAAACCTACACATTGCA | *Gossypium spp* |
| AMPL3261332 | TAAAGAAGGACTGGGTTGGAAAACAAA | GCCAAGCACAGGTATTAATTTGGGC | *Gossypium spp* |
| AMPL3261336 | CAAACAAGGCATGTTCATAGTATGAGTGC | GAGAGGCCATGGTGTCAAATGAC | *Gossypium spp* |
| AMPL3261342 | GCAGATTCTCGGGTTTTATATATCGAAGGT | TCACTAGATTTGGTGGCCTTGAACC | *Gossypium spp* |
| AMPL3261351 | GGCTGGTGAAACACAACATCAAA | AGAATTTGGTAAGCTGGTAAACCCTAAGC | *Gossypium spp* |
| AMPL3261362 | GATAACGAGACAATCTACGACCCAAC | GTTTGGTTTAAGGCTTTGTTCAATGTTCC | *Gossypium spp* |
| AMPL3261365 | GTTGTGTTAGGGAGATGGTCAATTTAGATAG | CTCTCTGTGGACCCGAATATTTCCA | *Gossypium spp* |
| AMPL3261369 | AGTTTCATTTGCATGACAACCCAAAAATTAT | CACGAACGAAATACGAATAACAACTTGAAC | *Gossypium spp* |
| AMPL3261380 | GATTATGAATGTCTCCTCGTTCGATGATC | ATTGATATGATTCGATTGGGTGTGTAGTTA | *Gossypium spp* |
| AMPL3261382 | GGACCATTTGTTTGGAAAATCAAAACCAA | GGCAGCTATCTGGTTTCTTGTGG | *Gossypium spp* |
| AMPL3261384 | GTCCTCCAAGGATTCTCCAAATACACA | CAACACATGGATTTATGACAAGATTCATCAT | *Gossypium spp* |
| AMPL3261393 | CCACCAGATCTAGAACCACTAAATGAACC | GTTGCGGTGGTCTACTCAGTTTC | *Gossypium spp* |
| AMPL3261394 | GTATTCGGAAGCTCGTAAGGGTTG | ACTTCCAACAATGAAAGTGTCGCTAC | *Gossypium spp* |
| AMPL3261396 | TGAGTATGATGACTTGCTTGCCAAC | TATCAATCTGTCGAAGGGACCAGTTAA | *Gossypium spp* |
| AMPL3261398 | AGTTGGAAGTGATGGTAAAAGAGGTGG | GAGTCTTTCTTTCTTGTTTCCAGCTTACC | *Gossypium spp* |
| AMPL3261400 | TTTTTATTTTTCGACTTTCTTCGATTGGAAA | ACCAGCGTGCACCCTTTTATC | *Gossypium spp* |
| AMPL3261404 | TGTAAGTAATATGAGGTCGAGTTAGAACCAC | GTTCTCTCTTAAAGATATGGGTGATCTTCAT | *Gossypium spp* |
| AMPL3261406 | TGGCATACTGTTGGGCATGTTTG | GGTGAGATGAAATGCATTTTTAACGCAC | *Gossypium spp* |
| AMPL3261409 | AAAAGCTAAATAGCAATGGTAGCATCAACT | TTCCTTTACTCCAATGACATGGAAATCTTC | *Gossypium spp* |
| AMPL3261412 | ACTTCTACGTATGGTTTATTTGTGCTTCTT | AGTTTGAAAGCTGGTTTTATGTTGTATCACT | *Gossypium spp* |
| AMPL3261413 | AGTGGGATGTGAGAGTATAGCAAGTT | CGGATGGATTACCTCCTAAGCTATCAAG | *Gossypium spp* |
| AMPL3261416 | CGATGGTAGAAAAGACATATCGTTTCGAA | AAAAACTTTCCAAAGCCATGATCGTAGC | *Gossypium spp* |
| AMPL3261418 | GCAACACGATTAGCGTAATTCAAATTTAGGC | CGTGTTGCGGACTGTTTGACTAA | *Gossypium spp* |
| AMPL3261419 | GACTTAAATGATGTGAGAAGTGGTCGTTAT | GTCCAACTGTATTGATAATTCCCATGAGTTA | *Gossypium spp* |
| AMPL3261429 | CCTAATTGAGTGTCTGACCTTCTTAGTAGG | TATGCATAGAGAACAAATGTTGTCGTTGAG | *Gossypium spp* |
| AMPL3261433 | ATCTCTCATTTATCTTTTGCTCTATCAACATGTC | CAACCAGAGGGTTTTACAATTTTAGAAAAAGAAG | *Gossypium spp* |
| AMPL3261437 | CTTAAAAACTTTATTGTCAGGGTTTTGGGTA | CGTTCGGACCACTCTGTTACTCT | *Gossypium spp* |
| AMPL3261441 | ACCCAAGTGGTTAAGCCTTTCATC | GTTTCCAAGAGATCAATTTCCAACAAAAGAA | *Gossypium spp* |
| AMPL3261445 | TTTCTCTTTTTGGTGTTTTTGCTACAAGTC | CCACATCCGAAAAGGTAACCCTTTAATTAAT | *Gossypium spp* |
| AMPL3261446 | CGGCAAAGTATCCTAGAAAGATGCAT | GAATGAGCAGAACCTTTCTCGAAAAGG | *Gossypium spp* |
| AMPL3261453 | GCTCATCATATATCAAAACATGTCAAAGCAT | GCACACCGCCATATATTGGTGTG | *Gossypium spp* |
| AMPL3261454 | CGAAAATCCCATTTCCATAATCCTAACTTCT | GCGTTTAAGTTACTTTTTGTCTCATGCTACA | *Gossypium spp* |
| AMPL3261456 | GGGCCTGATTTTCCTTCATTCTACCT | GGCTACTAGTAGGGCTCAATAAATATGGAG | *Gossypium spp* |
| AMPL3261458 | CCTTATACATTTCTTATGGCATCATTGGGTC | TCCTGCTTCTCAATACAACCACTATTAATGC | *Gossypium spp* |
| AMPL3261479 | AAGAGAGTAAAGAAATGATTCCTCCCAAAGC | AAAAACAAAGAGAAAGGGTAGATGGAAAGAT | *Gossypium spp* |

| | | | |
|---|---|---|---|
| AMPL3261482 | CTCAACTGCATACACACAAAGTATGTCG | GTAAACTACACTAGCAGTCACCCAAC | *Gossypium spp* |
| AMPL4000000 | CCTCCCTTACTCTGCCACTTTTT | TCCCATGTTTCTCTTGTGGGTTACG | *Gossypium spp* |
| AMPL4000001 | CTGGTCGGATCCCACTTATGAGT | TGTTCGTATGCTAAAGAGGTGTGATTAAATT | *Gossypium spp* |
| AMPL4000002 | TTTTTGGCATTGGTAAAACAACATTTAGGG | CAATGAAGCATGACAGTCGCAAC | *Gossypium spp* |
| AMPL4000003 | CCAGCTAACCAAATAGGATCTGGTT | AGGCCACCTCTGATTTGTTCTTTAAGA | *Gossypium spp* |
| AMPL4000004 | GCTTGGGAGATCCTACATGTACATACA | AAAATACCGGGTTCTATCTTAGGCTTGG | *Gossypium spp* |
| AMPL4000005 | GGCATGGAAAAGAAGGCATGAGAA | CTGGAGAAGAAGTCCTACTTGAAGTCTAA | *Gossypium spp* |
| AMPL4000006 | GGCAACTACAACGGTGATGAGAG | CTTACCTTGTTGCGTGATACATTAAACCT | *Gossypium spp* |
| AMPL4000007 | TTAATACTAGTACTTTTCGTCATTGTAGATACGA | AACCAAAATTGCATATAACACTATATTGAACCGA | *Gossypium spp* |
| AMPL4000008 | AAAAATTCAGATGTTTGGCCCACATG | GAGGGTCCATTTAAGTGACGTTTCG | *Gossypium spp* |
| AMPL4000009 | ATTGATTTGTGGATTTGGTTGTGAAATGAT | CCCTAATTCAACAACCAGTGAGTTACAC | *Gossypium spp* |
| AMPL4000010 | GTTATCATGCACACCCGATGAGTG | ATGACGCAGATTGATAGCTTGACTAGA | *Gossypium spp* |
| AMPL4000011 | GAAATAAAAATGACGACCCTTGAGCAAA | GCGAAGTAGACTACTTAGTCAAAGAGCT | *Gossypium spp* |
| AMPL4000013 | ACACTCTGTACGATTTGACTACAAATCCG | AAATTTCAGTGGCTTTTCCCAAATTTTCG | *Gossypium spp* |
| AMPL4000014 | ACTTTATTCCACTCCATTCAACCAAACAA | ACAAGAATTGATTTGGGTCAAGGTTCTAG | *Gossypium spp* |
| AMPL4000015 | TTAGGTAAAATTTGGGTCCACACGAAC | AAAGTCATGAAAATTTGGAAACAAAGGACAC | *Gossypium spp* |
| AMPL4000016 | TTCGAGAGTCCAATATAGGGACTTATTTACTA | GACAGAGAGTAAAAATGGAAAATACTTTTGGAAA | *Gossypium spp* |
| AMPL4000018 | GATGCAGCACAAAATTAAGATTTTGGAAGT | GTAGTAAGGCGTCAATGAATACATGCC | *Gossypium spp* |
| AMPL4000019 | TTTGGAGATGTTTGTGCTTGGTTTCA | CACTGTTTAGGGATTAGTTGTTTTTCGAAAG | *Gossypium spp* |
| AMPL4000020 | TTACACAAGGTTTGATTATAGACGAGCAAG | CCATTTTTACTACTAAAAACAGACGTGGTTG | *Gossypium spp* |
| AMPL4000021 | CCACCAAGCATTTCTGTAGTTTCCA | GGTGAAGGTAGTAAGGTTTAAAGATAGGAGG | *Gossypium spp* |
| AMPL4000023 | CTCTCTGTGAACCCTAGTATTTCCAAACT | GACTTGATTTGGAAAGTCTTTTGAGAGATGG | *Gossypium spp* |
| AMPL4000024 | CGACTACCATTGGTGAACTTGCAG | GAAGTGGGAAAGTCCTATGAGACTATCTT | *Gossypium spp* |
| AMPL4000025 | TTCAGAAAAGAGATTTTCAGAGAGGGATC | TCATTTGCACCTCTATCTAGCTCCTAAAA | *Gossypium spp* |
| AMPL4000026 | GATAGATGGTTGACTGTGATTATTTGCTGA | CACTACCCTAAAGCTTATAATCCACTTACCT | *Gossypium spp* |
| AMPL4000027 | GCCTAATTTCACTGAATCAATTGCTAAGGT | CAAAATATAATCAACCCAACCCAATGCAATG | *Gossypium spp* |
| AMPL4000028 | ATTGAGCAATTAGAAAGACAATCACGATTGT | CTCTCATAGGCACATTCTTCAGAGAAGA | *Gossypium spp* |
| AMPL4000029 | TGTACGACATTCCTATCATAATCCAAACCA | GTTTTACGATGCAACACTAACTACTTGAATA | *Gossypium spp* |
| AMPL4000030 | CTTCCGATCATTTTTGAGAATTCTTTGTTCAT | TATATATCCAATCGGTCTCTCCGCTAGC | *Gossypium spp* |
| AMPL4000033 | AACCAGATACTGGGAAGCACTACTAA | TTGAGCCTCTCTGTATTTGACCTTGG | *Gossypium spp* |
| AMPL4000034 | TCTTGCCAACTGTTGAGAGGTTATCA | CTCGAGAATTGGAGGGACGATAGG | *Gossypium spp* |
| AMPL4000035 | CGTAACATTAAGACCTGTGAAACACAGTT | TGTAAAGTCCAAACCCAAAGGAGTT | *Gossypium spp* |
| AMPL4000036 | GTCCATTTACAGACCTTGACCCAAT | CTGCTTTGTTTGTCTTGTTTGTTTGCA | *Gossypium spp* |
| AMPL4000038 | AATAATTGACTTTCATGTCCTCAAGTCCAA | GTCTGATGGAATGCTGAATTTACTTCCT | *Gossypium spp* |
| AMPL4000039 | GAATCTGAACATACATCCGTGTCTGAA | CTTAATTTATGAACCGAGAGTCAATCCGAC | *Gossypium spp* |
| AMPL4000040 | TGAATCTAAGAGGTGCAATGTTTATGGAGA | GACTTGAAAATACACTTTTCCGATACCAAAA | *Gossypium spp* |
| AMPL4000041 | CATTATCCACCACCTCTATAAAACTACCCT | CATTGTGTCTGAATTACGTTAACAAAGCTTG | *Gossypium spp* |
| AMPL4000042 | CGCTCATTACACAACTACAACGAACA | CAAAGTTGTGGTGGAATTAAAATTCTGAGAA | *Gossypium spp* |
| AMPL4000043 | GATTTCCTTTTGGGAAATTCTATCACACTT | TGTCAGCAAATCCTTCGAGACTATGG | *Gossypium spp* |
| AMPL4000044 | TCCGTGATAATTAAATGGCTTTGGTCTC | TTTTGGTTGGTTTTTAGTGACCCTTTTATGA | *Gossypium spp* |
| AMPL4000045 | ATTCCTCCTCTTTTCATGTCACAGTTTG | GGTACACACCTAGTTCGTTTTTGCTG | *Gossypium spp* |
| AMPL4000046 | TCAGAAATATGGTCATGTGGCCAAAT | GTCGTGTGTCTTCTTGATTTTAAGTACAGG | *Gossypium spp* |
| AMPL4000047 | TGTGACCCTGTCCTCTTCTACAC | GAAAGAGCTGTACAATGTGGACATGC | *Gossypium spp* |
| AMPL4000048 | CTATGATATGGCACTCTGGTCCAAA | GATTTAAGAGCTTTCGTAAACTCGTTTGAAA | *Gossypium spp* |
| AMPL4000049 | ACTTCTAACGTGGAGCAAAATATTTGTTGC | ATTGTACAGATTGGTAATGTCCTGTAATCCT | *Gossypium spp* |
| AMPL4000051 | AGAACATGGCTTCGCAATCTCAA | TGCTAAGGGAACAATTTCAGTTGGTG | *Gossypium spp* |
| AMPL4000052 | GGTTGAAATAATGATGGTCTGGTTAAGCC | GAACACACTTTGTAGACTTATTTGGACAAAG | *Gossypium spp* |
| AMPL4000053 | ACATGTATCTGTGGCTATAACAATGCAT | TTAACATGGCTGACGAAATAACCAGATAAT | *Gossypium spp* |
| AMPL4000054 | TTTAGGTGAAAGTCGTGAGACGTTG | TTTATATAGTCTGACACACGGTCATATTGCG | *Gossypium spp* |
| AMPL4000055 | CCTTTTCCCTTAGCCCACTTCTT | AGATGGTGCTGGATAGATGAAAAGGG | *Gossypium spp* |
| AMPL4000056 | CCCATATTGGTGATGGGATGATCAG | GATCCGCCTAAGTTTGTGGAGTAC | *Gossypium spp* |
| AMPL4000057 | TGGAATTTGTAGAAGCTATGGAAAATGACTA | ATTCTTTCTAGAACAACTCATTGACGGAAC | *Gossypium spp* |
| AMPL4000058 | ATTTGAACCAGCTTCAGCAATGTGA | TGACCCATGTGCAATATCTTGTACGA | *Gossypium spp* |
| AMPL4000059 | TCAAATTTGTTCTCAAACTTCAAAACATTCCA | TTAAAGCTTTTGTAAAAGTTTCGAAAAACCT | *Gossypium spp* |
| AMPL4000060 | GTGAACTGTCGCATGAGTAGTCAC | ATCTCCACAATACTTCAACTAGGAAAGCT | *Gossypium spp* |
| AMPL4000061 | GGAGCGATTTGCAATTGGGTTAA | CCTATTGAGCTAAAATGATGTTCTCGTTGAT | *Gossypium spp* |
| AMPL4000062 | ATTGAGTTCCACGATCATTTTGAGATGTAA | AACAAGATTGATGTGGTAACTTGTAACATCC | *Gossypium spp* |
| AMPL4000065 | CCAAGTCCATTCTATTAACCCAGTCTA | CGTTTTATTTTCTTATGGATCGCTAAACACTTTT | *Gossypium spp* |
| AMPL4000066 | AATCTGAATCCTACTTCTAAACCTTCATGGT | TCCATTAGTTTTAATGGCAAGAGTTGGAAT | *Gossypium spp* |
| AMPL4000068 | ACACATTTCATATGTCCTTATTAAGTAGTTTCAA | GGCATAGTTTTGGGATGAAAATTGGTTTATTG | *Gossypium spp* |
| AMPL4000069 | GATTAATCAGTCGATGGTCGAATTTAGTTGA | GAATTGACAGAAGAAGCAAACTGTTAGAGA | *Gossypium spp* |
| AMPL4000070 | CGTCGGATTTGAAGTTGAAAGAGGG | GGTAGCAAACTGGATCAGGTCAG | *Gossypium spp* |
| AMPL4000071 | CTTAATCCTTCTTAACCTCGCAATCAACT | CCCTACTAGAACAAAATCAAACCATAGGTTG | *Gossypium spp* |
| AMPL4000072 | AGTGCCACACTCACCAAGAAAAT | TGACAGCTTCGTCTGTTCTTAGTTAATAACT | *Gossypium spp* |
| AMPL4000073 | AGATAAACTAGTTGTAGTTGCTCACCTTG | AAACTGTATGGAACATAAGTGGATGAACATT | *Gossypium spp* |
| AMPL4000074 | GTCCATCTGCAAACATGTGCAAA | ATGACCAGATGGCTTTTAGGGTTTATTT | *Gossypium spp* |
| AMPL4000075 | GTATTTGATCGAGACCTGAGGTTCAC | TTGACCATGACTCGTATTTTAAACTCAGTG | *Gossypium spp* |
| AMPL4000076 | GTTGCAAATTGAGCTTCTTGAGCTG | CCTTTCCACAAATGAGATGCATTCCC | *Gossypium spp* |
| AMPL4000078 | TGTTTGCACAATAATCTCGTCTCTTGAG | TACTGCCACATCACATTTTTGAATAACAAGC | *Gossypium spp* |
| AMPL4000079 | ATGGAAGAGAGGTGCAAGAGGAGAA | ACACGATCGTATGGCCTAACACG | *Gossypium spp* |
| AMPL4000080 | GCCTTGGCTGTGTGGATTATACCG | GGGCAGTTTTCTCAGTAAATAGAAACTAAGT | *Gossypium spp* |
| AMPL4000081 | CCAGGGTGGATTACCGAGGTAAT | TATGTGACAATTTCCTAGATGATTCGTCACA | *Gossypium spp* |
| AMPL4000082 | GTGAGGCTTGGTTTTTATGGTTAGGG | CCCAACATGGGTTTCACAAGTCC | *Gossypium spp* |
| AMPL4000083 | CAGCTTGTCTGGAAAGTGTGATATAAACT | GAACATGCAAAACCATTTCATGCTTTGT | *Gossypium spp* |
| AMPL4000084 | GTGCCACACAAGCTGGACACAAG | CGGGATTAAATGTTACGGAGTGGTAAAC | *Gossypium spp* |
| AMPL4000085 | CCGGTTATCTACCTAATTTTGAACTGAACTT | GTAATCAATCTTTGAAAGATCCGAACACTTC | *Gossypium spp* |
| AMPL4000086 | GAGTTTGTGGTTATGAGTTTGGTGTTTA | CAAGCCCATAAGCTTAAACATTAAACTCAAA | *Gossypium spp* |
| AMPL4000087 | TACAAATATGAGCGAGCTTGAACAAAACT | GCAATGACTAATTCCCTCGTAGTGAG | *Gossypium spp* |
| AMPL4000088 | GGGCATTGGAAAGAGGAGTTGAG | AAAAAGGAGTGAATTGAAACTGGTTGAAAAA | *Gossypium spp* |
| AMPL4000089 | ATTGGGTTGTAGTTATTTGGATGTTTGTGG | GAACGTCGTCGTGACATCATCTG | *Gossypium spp* |
| AMPL4000090 | ACTAGCTGGTAACTAATACCTGCACTA | ACAATGTGTTAGTTGCGGTTAGTGTAAA | *Gossypium spp* |

| | | | |
|---|---|---|---|
| AMPL4000091 | GATGTGGCTGATGCAAAATATTTGTCTA | GACTATGGACGAGTTCAAGCAGTG | *Gossypium spp* |
| AMPL4000092 | TTTAAAACAAAAGGGTGGAGGAGACATT | TTTCAGTCCAATTGATCCAATCGACTAATC | *Gossypium spp* |
| AMPL4000095 | GCGTCACATCATCGAGACGATAG | AGTGCACAAGTCAAAAGCTAGAAGTATTC | *Gossypium spp* |
| AMPL4000096 | TGATTAGCAAGACACATCTTGAGACAATC | CTGACCCTCGCTGGTACTAATCC | *Gossypium spp* |
| AMPL4000097 | AAGCTAACTCGGAAGGATTCACTGC | CATTGAGACTTGAGTTTCTCGAGATCAAA | *Gossypium spp* |
| AMPL4000098 | GGGTGTTACATACTCTCAATCTAGACTCT | GGTGGCTCACAATACAGCCAAAG | *Gossypium spp* |
| AMPL4000099 | CGTGTCCTACTGCTAGCAGTTATAATTAA | ATATTTGTTCACTGTATGTTTGACTGCCTAT | *Gossypium spp* |
| AMPL4000100 | TGATTCATTTTCCCAAGTTCTCTAACAGTC | GGGAGGTGAGATTTCCAGTTAGAATTAAGC | *Gossypium spp* |
| AMPL4000101 | CTTTATCCTCCAGTTCCTATATCAGATCGAT | TACAACATATCACTCGCAAACAGATGAG | *Gossypium spp* |
| AMPL4000102 | ACTCGAGAAATGAATTTTTAAGCAATCGTGA | GTCCAACTCACTCGAATCGACATTC | *Gossypium spp* |
| AMPL4000105 | GATTGAATATGAGATTAGTCCGGTAATGCTT | CCTTTGAGTCCCTAAAAGGACATCAAAAAT | *Gossypium spp* |
| AMPL4000106 | CTACTTTGGATTCATGATTTCAAAACCACTA | TACTACTTGAATGCAGGATAGGTTAAGACTC | *Gossypium spp* |
| AMPL4000107 | TAACCAAGCAGAACACACACAAATTCTAA | TAGCCATATCACATGGCATATGTAACACC | *Gossypium spp* |
| AMPL4000108 | AAAAATTGAACCGATGCCTGCAAC | CGTCGGTACCACCCTTTATCATCC | *Gossypium spp* |
| AMPL4000109 | TTGGCGCAACATTGTTATTACCACT | GCCGCTTGATAATGAGGCATAAAAAA | *Gossypium spp* |
| AMPL4000110 | GAGGGACTAGATAGAAATTTTCCCATTTCCC | AATTTGGTTTTGACTTTGAGACTTCTCAACT | *Gossypium spp* |
| AMPL4000111 | GCCCTCTTGGCGTATTATCGTAT | TCTAGGCTATGTAAAAAGGTGAGAAAGAGAT | *Gossypium spp* |
| AMPL4000112 | CTGAAGTTGTACTTTGACAACAACAACAA | ATTCCGTTCTCAATGGTTTTTGAATGGG | *Gossypium spp* |
| AMPL4000113 | TTAGTAGAGAATTGTACGATGATCGGAGATA | AGTGGATCCCACTCACTTATCGTC | *Gossypium spp* |
| AMPL4000114 | TCAATTTTGTAAGCTCCCAACAAATCTTGA | GGCTAGAGCAAATTGGTTAAAGAATGATGA | *Gossypium spp* |
| AMPL4000115 | AATTACATTCTTTGGCTTTGGCTTTTGTAG | ATCTCTTTAACAGCTTCAGAAACCCTAGG | *Gossypium spp* |
| AMPL4000116 | GTCACACCTGTTACGGCAATCTT | AACTCTCCACTCTACATCTTCCTCTTAGA | *Gossypium spp* |
| AMPL4000117 | CTTTGCAAATAACGAGATCTCCATTTAGCT | CTCTAGATGATAGGGTTGCCAAAATTGAG | *Gossypium spp* |
| AMPL4000118 | ACAGTCAAACAGTATGGATTGTTTTGCT | CCATTCGAGAAACCTTCCTCATTGTTT | *Gossypium spp* |
| AMPL4000120 | TTTTTCTTATGTCACATGTTGCCCATT | CCCTAGTGATGCTTCACAATCTGGT | *Gossypium spp* |
| AMPL4000121 | GCTAGCAATCTCCATTTCAGATGTACTC | ACCTTCATGTATTTCTTTATTTCGACCTGTC | *Gossypium spp* |
| AMPL4000122 | AACATCATTTAGACTCCCATTGAAAGGTG | GGCTATGTGTGATGCCATAAGCC | *Gossypium spp* |
| AMPL4000123 | CGTGAACTTTGCTTTGGCTGATT | CGTTCCTCATGGTAGATTTGAAACTTACAC | *Gossypium spp* |
| AMPL4000125 | GTAATGCTTCGGTTTCCAATAAGGAATCG | CGTATTCTCAAGCCAGTATTTAGCTCTGG | *Gossypium spp* |
| AMPL4000126 | GCCATGCATAAGACACGACATGT | TATTTGACCATCCCTTTACCAAACACAT | *Gossypium spp* |
| AMPL4000127 | AACAACAATTCATTCAAGGCAAATAACAGAA | AGGGATAAGCGTAGTCGTTGAAAACT | *Gossypium spp* |
| AMPL4000128 | CGAACTCATAAATTTTTGCATTGGAAACAAT | CCCATTTTGATCGTGCAAAAGACCA | *Gossypium spp* |
| AMPL4000129 | TGTCACAACCTTTTAATGACAGTATCATGAA | GCAATTTAGGGATCTCAGGGTCATAATACC | *Gossypium spp* |
| AMPL4000130 | TCTTGCTCCTTATAACTCAAGCATGAAATC | GGAGTGGTAGCTCATAGTTGCTAGG | *Gossypium spp* |
| AMPL4000133 | GTTGTTGGCGTGCCAATGAATTATA | AAAAGTCTCAAAATGTTGAACAGAGAACTCT | *Gossypium spp* |
| AMPL4000135 | CCACTGTAAGATATGGAAGCACTGTA | TTGGGTTGGAACTAGTGATGAACCC | *Gossypium spp* |
| AMPL4000136 | CATCCGTCGTGCCATTTCTTTCT | CAGCATGGTAAGGATGATCCTATAAGCA | *Gossypium spp* |
| AMPL4000138 | GGCCTGGTCTACAAAAATGGAGTT | CTCATTGTCCAGAATTTATGTCAGAGGTG | *Gossypium spp* |
| AMPL4000139 | AGCTCATGAAGACCCTATCATACTGTT | AGCAAATTGCCTATGTCCTCGATC | *Gossypium spp* |
| AMPL4000140 | TGTCAAGGGCATAGGAACTCAATATTGG | CGTGTTTAACTTTTGATTGCAAAAACCAA | *Gossypium spp* |
| AMPL4000141 | CGATTACGAGAATTGGCAAATGGATGT | CATACAACCACTAAGCCAGACACTC | *Gossypium spp* |
| AMPL4000142 | GAGATTCATGCACGACTAGGTACATCG | TAGTGGTGTTTGGCTGAAAAGACAC | *Gossypium spp* |
| AMPL4000143 | AGAGGATAGGCTATCATTTTCAGGAACTT | GCATTAAACTTTCCAACTCTACACACAGC | *Gossypium spp* |
| AMPL4000144 | CTTGGTACCTAGAGCCTCATTTTTCTT | CCCAATGGAAATGGGAGCCTATTACT | *Gossypium spp* |
| AMPL4000146 | CATGATTGTGTCTTTGCCCATGTG | AGCGATTAGGTTCTTTTATCAAATTGAGCA | *Gossypium spp* |
| AMPL4000147 | AGAAATATTTTGATCACCAGCAGTGACAA | GCCTCTGCTCCTCCTCTTTTCTTAT | *Gossypium spp* |
| AMPL4000149 | TAGTTTTACTCCTCATACTTGGAGGTCTTT | AAGGAGACTCCAGTTTAAATTTCGTGTT | *Gossypium spp* |
| AMPL4000150 | CACCACAAGCATTACGAGCATCA | CAATTCTCTTTGCGCCTTGCTTG | *Gossypium spp* |
| AMPL4000151 | AACCGCTCCCAATTTATGTTTTGGG | CCATCTTCAGCAGTAGCCCTAAATTCT | *Gossypium spp* |
| AMPL4000152 | TTTGTTCGAACGAACTCCGTAAACTAA | ATGTTACACGTAGCGTGTGTGAG | *Gossypium spp* |
| AMPL4000154 | ACCAATACCAATGGACATCCAAGAGC | GAGAGAAGGAAATCTGAAGGCGAAG | *Gossypium spp* |
| AMPL4000155 | GGTTAGAAGAAAAATGGAAATATGGGCAT | TCCCTTTAAATCAGATTTGTGGTTACGAAAC | *Gossypium spp* |
| AMPL4000156 | CAGGGTAAGAAACTCACTTATGATGTAGAAC | CACCTCCATCTCAATAGAGCCTTTA | *Gossypium spp* |
| AMPL4000157 | AAAAATCTGCGAAGTCAACAGAATTACAT | GCAAGTTTTGTGACCTCACCAAAGA | *Gossypium spp* |
| AMPL4000158 | CATCCAGTCCATGTACCTTAACAACTC | GTCTAATAGGAAATGAGCAAACCGTTTGA | *Gossypium spp* |
| AMPL4000160 | AGAAAGGGTTTTTCAACGATATCACATTTT | GGAATTTGGAGATGGTTTTATGCCACTATG | *Gossypium spp* |
| AMPL4000161 | AGTACAAGTGGTTTGTCCCTAAAGTTACG | TCATCAACCGTCCAAATTAATTTCCAAGT | *Gossypium spp* |
| AMPL4000163 | GGCCAAAGAGTAAAACATTGTTTCGAG | CTTAGTGGTGTCACGACGTCAAG | *Gossypium spp* |
| AMPL4000164 | CTTTATATGGTCGGAAATGCAGAAGTCT | CGATCCAATAAATTTTGGATTGAGCTTTCAT | *Gossypium spp* |
| AMPL4000165 | GACATTTTGATCTACTCCGGTGATGAATC | GTCTTACGCTTTTTATATGCGAATAAGGTCT | *Gossypium spp* |
| AMPL4000166 | TCAGGGTCTCGAGGGTTCTTTTTAG | TACAAGGGTTCTACCTTCTCTTCCTAAAAA | *Gossypium spp* |
| AMPL4000167 | GCCATTACGACACGTATCAGATATGGT | GGGTCAGTAAGGGTTAAATAAACCCTATTCT | *Gossypium spp* |
| AMPL4000168 | GGATTATATCATGCCATTGTCTACTTAGCAT | TTTTATCCAAATATTCAAGTTTCGATTTAGTGGT | *Gossypium spp* |
| AMPL4000169 | CAACAGGAAGAAGTCATGGGAGATT | ACTTCACTCATGGTTTTCCCTTAAAAGAA | *Gossypium spp* |
| AMPL4000170 | AAGCCCAAACCTATAGGGACCTAAA | CCTGTTACAATATGTTCTTGTGGCTTTA | *Gossypium spp* |
| AMPL4000171 | CCTAACACAGGCGTGAGAGAAAC | GAAGAAGATGATGAATAGTGAAGGGTCAAG | *Gossypium spp* |
| AMPL4000172 | AGCTTTACCATATGGCATTAGAGCAAT | GAAGGAGAGAAATATCCACGAACTAAGTA | *Gossypium spp* |
| AMPL4000173 | GTCGAATAGAGAGATAACTTGTCTTGGAAAT | CATTTGCCTAATGTACTAGCTTTCAATCCA | *Gossypium spp* |
| AMPL4000174 | ACTTCTTGAAGAGGTAAATGACTTACGATGA | GGGCTAAAAATGCCTTACCTGAGAC | *Gossypium spp* |
| AMPL4000175 | TCTTTAGGTATACCATTTACAACCCTCGATA | GGGATAGTCTAGTTGACAAGATCCATTGAT | *Gossypium spp* |
| AMPL4000176 | ATGATTGAACAAAATTTTGTAACCACGAGAC | AACCAATGACTTTTCATACACAAGTTGTCT | *Gossypium spp* |
| AMPL4000177 | GTTGGCCGAGGAATCCATCTTAT | TCAAAAACTCCGATAACTTATTGGGCTG | *Gossypium spp* |
| AMPL4000178 | AAAGAAGCTTTGTATCAAGACATGGAGAC | GGATATATTTGGTGCTTTCAAGACACACC | *Gossypium spp* |
| AMPL4000179 | TGACTAATGTCCAAATCATTTCTAGCGTA | TAACTAGCTTATGTGGCCAGTTAACTTGC | *Gossypium spp* |
| AMPL4000180 | TCTATCTCGGATTCTGTGGTCATGAA | CTTCAATGCCCATTATGTTGCTGTG | *Gossypium spp* |
| AMPL4000181 | GAGGGATAAAATTGACGTCGAATAGCT | CAAAAAGATAAACATCACCACATTTGAGATCAG | *Gossypium spp* |
| AMPL4000182 | CAAACAGTAAACTTCGAGCTATTTGTTGAT | TGTATAAATGGGAAATAGGAGGTCCCTAATG | *Gossypium spp* |
| AMPL4000183 | CTTTGGATCATTTGGGACACACAAT | GCAAACATTGTAACAAATGCCTTAAGTAGGA | *Gossypium spp* |
| AMPL4000184 | GGTTGTGTGCGATGACATGACTG | CCGATTAGGACGACCCATGTAACC | *Gossypium spp* |
| AMPL4000185 | AAGGGAAGTTTCTATCCCAAATGATTGAAA | ATGAGGCCAATTGTTAAAACAATGTTGATA | *Gossypium spp* |
| AMPL4000186 | TTTTTGCAAAAACTCTTTCTCAAGTTTTGAC | CCAAAGCAATTAGATTGTTGCAGGAAC | *Gossypium spp* |

| | | | |
|---|---|---|---|
| AMPL4000187 | CAAAAGGCCTAATTTGTGCCAAAAATA | AAGTTCATTATACAAACTTTCTCGGGTCTTA | *Gossypium spp* |
| AMPL4000188 | CCAAAATTTTCCACACGTCTTCTACTTTT | TGGGTTCAATATAACGGTTGCAAATATTTGT | *Gossypium spp* |
| AMPL4000189 | GCGACTTTTCGAGATGCGATGTAA | GTGCATTCTATCGTATAAACCTAACGAAAGG | *Gossypium spp* |
| AMPL4000190 | AAAAGATGATGCTTTTGAGACCTCTGAC | CTGAATAACACTTGTGACCGAATGTTTG | *Gossypium spp* |
| AMPL4000191 | CAATGGTAATCAGCATTCTCCATTGTTTT | CCGATAAATGTCTAGGATCAACTATATGCAT | *Gossypium spp* |
| AMPL4000192 | GAACGCTTGGTTTATTAGATTCTTATCACCC | GGAGCGAAAAAGAGAGTAAGTTTTACATTTT | *Gossypium spp* |
| AMPL4000193 | CCAGCAACTTCCTCGATGAGAAA | GGGAGATTAATAAAGGGAAGCAAGTTCATTT | *Gossypium spp* |
| AMPL4000194 | CTGATTTTCTCTCGTTTTTCGCTCTT | CATATGCAAAAACTCATGACTTGGCTTG | *Gossypium spp* |
| AMPL4000195 | CATTCAACAGTTCATTATGGCCAAATGA | GTGTCAAAGTTAGAAATGAGGTAGAATGGAT | *Gossypium spp* |
| AMPL4000197 | CCCACATGTCATATTATTATGCTGCCAAA | CAGATGGGAACTAGGTTGGTGTTG | *Gossypium spp* |
| AMPL4000198 | ATCAACAACAAATTAATTGCTCTACACCTC | GAGGAGAGATGACTACTTCAAGTAATGCA | *Gossypium spp* |
| AMPL4000199 | CAATAAATCATTGTCACATCAGCAACCAA | TGAACTACCCACATGACATATGATACTCAC | *Gossypium spp* |
| AMPL4000200 | GGTGCACTGAGTCAACCCATAGA | TACGTAAACGCTGCCCTTACAAAA | *Gossypium spp* |
| AMPL4000201 | AATTTAGGAATCACTCGAGGACAAAACAG | CCACACGAGTGTGTGTCCTTACC | *Gossypium spp* |
| AMPL4000203 | CAAGCTCTTCAAATCTTAATTCATCACCTTA | CCATGATGTTTCAACTATTCTTGGAAGACC | *Gossypium spp* |
| AMPL4000204 | TAATGAACGAATGTAGGTTTCATGAGTTCCA | TATGGCCCAAAAAGCTAGTTTGTCTG | *Gossypium spp* |
| AMPL4000205 | TTGGACACATTGCAGACACTTTATAAGAG | CAAGGCCCTTTAAATAGGCTAAGATTAAAGT | *Gossypium spp* |
| AMPL4000206 | GGCTTGTATTCGTTTCATGCGAT | GGTCATGTTAATGTAAGGTTCACACGAG | *Gossypium spp* |
| AMPL4000208 | GTTATGCGGGTGGTGTAGGAAAC | TGCATCATTGCGATGGAGGAAAA | *Gossypium spp* |
| AMPL4000209 | GTCCACAATTATTTGTTGGTGTGATTTAGTT | GGGTGAGCTATAAAGCTCAGTGTATACTAAA | *Gossypium spp* |
| AMPL4000210 | GTGTTGTGTGGAGGTTTTGTGTTTG | AAGAAGATCGAAATCAAAGGCAAAACTTAGA | *Gossypium spp* |
| AMPL4000211 | ATCAGTCAGGGCACTTAAGTATGATTGC | ACTCAGTCATAGGATCCAGATTGGTG | *Gossypium spp* |
| AMPL4000212 | ACGAGTGAGTTAGAAGCCCAGTAT | AATTGGATGGATGTACTCATGTAGTCCTAG | *Gossypium spp* |
| AMPL4000214 | ATTGTAAAGTTCAACCTCCTATGTGGAAAC | CAAGAGTTTTCATATTGGGACTTGAACTTTG | *Gossypium spp* |
| AMPL4000215 | AGGCCTACTATCTATGTTTGACCCAT | TCAGTTAAGAGCGTCAATTCTATGAACAATA | *Gossypium spp* |
| AMPL4000216 | GGTGAGCATGCATTCTTGTGATATTGG | AGAAAGGGATTCTTTTTCTTGTCTTGTACAT | *Gossypium spp* |
| AMPL4000218 | TTTTTGGAATGTGAAGCATGTCACTT | GGACAAACACTAAAGTGAGTTCTTATCTCAG | *Gossypium spp* |
| AMPL4000219 | TTTTTATTTTGGCTATTGGGCTAATTGGG | TATCCAATCGAATAGCATCACATTTGCTC | *Gossypium spp* |
| AMPL4000220 | TGATTTAACATGTCATATCAGCTTACCGTTA | TGAATCCAAATTTTGGTTAGGTTCATTTCGT | *Gossypium spp* |
| AMPL4000221 | TTCCTAATTCCTATCATTGCTTGGAAATCTT | GACAATATTCTCAAAGATGGAAAGACAACGG | *Gossypium spp* |
| AMPL4000222 | AAGGGTGGACAAAAAGCTTGGTAA | CTAAGCCGTGTGGGTGATTTAATTTTTC | *Gossypium spp* |
| AMPL4000223 | ATACCACTTTTCGAGTCTTACTTAAGCAGG | TGTTTCTAACTCGAAACCACAATGAAACA | *Gossypium spp* |
| AMPL4000224 | GGTGAAGAGTAGGCTCAGAGATGG | ATGACCACCTCTCATAGACCACAC | *Gossypium spp* |
| AMPL4000225 | AAGCGAAGATCATCGAAAATGTTAAGCG | CGGACAGTCTCTAATATGGTACTCTAAGAA | *Gossypium spp* |
| AMPL4000226 | GGCGAGCTAGGAGACATACAAAG | CCAAACCATGAACATTCCAACTGAGG | *Gossypium spp* |
| AMPL4000227 | CAGTCCAAATGCCGGTTAGAATCTG | CCACTTATTATCCCATCGGTACATGAGAT | *Gossypium spp* |
| AMPL4000228 | GGTATGTATCATACCCATTCACGAATCAAG | CTCCCAATACAATGATGGCTAATGGAAG | *Gossypium spp* |
| AMPL4000229 | GACTTCTAGTATAATGGCCTTCATGGTACT | GAGCTCATCATGTATGGAAGTTTTAGTCTAG | *Gossypium spp* |
| AMPL4000230 | GCTTTAACTTTACCTCCATAATACGCGAG | TAACGCAATGGGTAAAGAGCCTTG | *Gossypium spp* |
| AMPL4000231 | AGGGAAATGAGTTATGAGTTAAGCCTTTTA | CTTTATATCGGTGAACCATGCCATTAGTATA | *Gossypium spp* |
| AMPL4000232 | TTGGAAGGAATGGTCCATGCAAT | CCACACACATGCTCCTGTTTCTT | *Gossypium spp* |
| AMPL4000233 | GTGGCCTCAAAACTACCATTTTCGA | GCCGTAAGCTCATGTATCATATAACATTTCA | *Gossypium spp* |
| AMPL4000234 | GACATTATCTAAGGTGCAATAGCATTCCC | AATCAGGTCTACGTTGAATCGTCTTCT | *Gossypium spp* |
| AMPL4000235 | CCAAACCCATTCCAGAAGTATCATATTCAA | GTCATCCATGGTGGTGGAGTAAAGG | *Gossypium spp* |
| AMPL4000236 | TTCAGACCCATGCCAAACAAAGAT | AAAAACATGAGAGATAGAGAGAGCCTTTTTT | *Gossypium spp* |
| AMPL4000238 | TAGGCAGGGTTCCGTCTAAGATG | ATCTTAACCACTCGGCAAATGGAG | *Gossypium spp* |
| AMPL4000240 | ATGTCAATGAACTAAAACACTCCTTCCAC | TAAGGAAGCTGTTGCACGGAATAAG | *Gossypium spp* |
| AMPL4000241 | GTAATCTTGATATAAGCCTTCTCGAGCATA | TCAAATCATGTAACAGGATGCTCACAAATAG | *Gossypium spp* |
| AMPL4000242 | CAGGCTCAAAACGAAGTCGTTT | GATTAAAATAAAATGCGCAAAAACATACGGGG | *Gossypium spp* |
| AMPL4000243 | CAAACACCTACCGCGTTGTTTTT | GGCCGAATATGTGCCATAAGATGG | *Gossypium spp* |
| AMPL4000244 | TGTTTCGCCTAACCTGGACAATC | GTATTTGGGCAATTGTGAAGTTTTTGGT | *Gossypium spp* |
| AMPL4000245 | ATAATATTTGCAAACCTACTCATCGGATTTAGTT | CCATACTCAGATCCAATTATTGGATTGTGTAAAG | *Gossypium spp* |
| AMPL4000246 | CCAAGTGTGTTGCCTTATAATGATTTGGT | TGCATGAACTAAGTGTTGTAGTTTACTTCTA | *Gossypium spp* |
| AMPL4000247 | GCTGGAATAAAAGGGATTGATCTATCAGCG | GTCGCATATATTCCCATTCATAATCTACCAT | *Gossypium spp* |
| AMPL4000248 | TGGAGCAAGTTTTAGGCTTTAACAAAAAG | TCCAATCACTTTCCAACAATCACCTAAAT | *Gossypium spp* |
| AMPL4000249 | ATCGACACGGGCTTGAGAGAGTT | GTCGTGGTATTTATTTCACTCAAGATTGGG | *Gossypium spp* |
| AMPL4000250 | TAGACTCGGTGAATTCCGATTTTTGAAT | GTCCCAGAAAATACAAACTCTCGTATCTTAA | *Gossypium spp* |
| AMPL4000251 | CTCTCGTTGACGAGCTGCATAAA | GCCATCCGAATGAATGTTGAAGGAC | *Gossypium spp* |
| AMPL4000252 | CCGGTACACACATGATATGTTTTGATGC | GTGAGTTTTCATTGGTTTTCTTCATATCGTT | *Gossypium spp* |
| AMPL4000253 | AGAAGTTCTCTAAAGTTGGAATGATCTGTGT | GCTTGAGTAACCAAATCTTCATGTAATAGGG | *Gossypium spp* |
| AMPL4000255 | TACAATTAGGACACCAAATCCAACATCAA | TTTCAACAGTTGTACGTTAAGATGTTGCC | *Gossypium spp* |
| AMPL4000257 | AGTAGTGTGGATCTACGTTTTTGTTGAG | TGCTTACACACGGAACTTATGTAGGT | *Gossypium spp* |
| AMPL4000258 | CTCGTTACCTCCACAACCAACAC | CCAAGGCATGATTTAACCTAAACATCACA | *Gossypium spp* |
| AMPL4000259 | CTTTTAGCTTGTTTCCTCCAAAACTGTT | TATAACGACGGTGTGATTGAGTGTGT | *Gossypium spp* |
| AMPL4000260 | CGACTTTATCTTTTGATGCAGTGATGAC | GCTCTGCATTGAACGTTTTACCTCTAT | *Gossypium spp* |
| AMPL4000261 | CATATTGATATTGCTGGACATAGTGACGAC | CGTTGCCCAACATCTCTATGAGGA | *Gossypium spp* |
| AMPL4000262 | GCCCTGGTCAAGAAGACCAAAAT | TCCATTGTCTCAATATGAACAAAAATCACCC | *Gossypium spp* |
| AMPL4000263 | GGGCCAAATCTAAAATAAAAATAAGCCACTG | TCCAGGATCCGCGTGTTTTTAATT | *Gossypium spp* |
| AMPL4000264 | GGGTGTGTCCCACAAACAGTAAC | GGGAGGGTGACTATAATGGGAGTTC | *Gossypium spp* |
| AMPL4000265 | CTTAAAAACCAAGTCTGAAGTGCTTCATG | CACTCTAGTAGACCAGAGGCATAGAGG | *Gossypium spp* |
| AMPL4000266 | GAAATGAGAGATCCAGTTATCTTCAACGAA | CTCTCTAACGTTCTTCCTCTTCCAAACC | *Gossypium spp* |
| AMPL4000267 | ATGGGTATCATAAAATTCGACAACCCATC | GGCATTAATGTTCAGATTCCCTAACATTCTT | *Gossypium spp* |
| AMPL4000268 | TCTTTTCCAATCTTCAATACTTACCGTTGAT | GTGAAATTTTGTTAGGACAAACTACACCAAG | *Gossypium spp* |
| AMPL4000269 | CACATGAACGATCAAAATTTATCAAGCCAA | ACAAAAGTGACTACTTTTGAGTTTACCCTTT | *Gossypium spp* |
| AMPL4000270 | CTTCTCCAAGGTTTGTTGATGTCAAAGT | TGGAACAATTCCATTATAGGTTTTGGTCATA | *Gossypium spp* |
| AMPL4000271 | GGAGCAGACCAGTTTTGTGCTTAA | GCTTCAACGCATTATCTTCTTATATAAGGGT | *Gossypium spp* |
| AMPL4000272 | TGTTTATTGCATGTTATGCTCATTTTGAGAC | CGGTATCGTCGGCAATTTCAAGATG | *Gossypium spp* |
| AMPL4000273 | ATGTTCTTCACGTTCCCGATGTG | TAGTCGACAATGCCATAGTGAAGATGA | *Gossypium spp* |
| AMPL4000274 | CAAGTGAAGCTTGGCGAGAAAAAC | ATTAAAGATTTGCTCCATCGTACGAATATGC | *Gossypium spp* |
| AMPL4000275 | ATAGAAGGTTTTTACCTGAACACTGCTTC | CGAAAAAGCCTATTTTAGCCGTTTTATCTCC | *Gossypium spp* |
| AMPL4000276 | TCCGTGACGTCTTTGTCTTCACG | TAGGTATTGTGGTAGACGTCATCCAA | *Gossypium spp* |

| | | | |
|---|---|---|---|
| AMPL4000277 | GGAAGCTTATTTACTGTAGTTGTGTCAGTA | AATAAGAGACAGAGGTAGCAGAACAAACA | *Gossypium spp* |
| AMPL4000279 | TGTCATATTATAGTCCAAAAGTGTGGCATA | TAAGTATAAGACACCTAAGGTACAAGGCATA | *Gossypium spp* |
| AMPL4000280 | TCATTTATGACGATACACGAAGCATAACAG | GAGTGAACCCTTGAACAGAAGCTTAA | *Gossypium spp* |
| AMPL4000281 | AGTCTATACCAAATGGTTTGCACTTAAGGT | TTCCACGCGACATCAATGAGTTAC | *Gossypium spp* |
| AMPL4000282 | GCCTCAATTGGTAATGCCCACTTC | ACGTTTCTAACTGAAACAACTTCAAACAACT | *Gossypium spp* |
| AMPL4000283 | TCACCAGCAGGTCTAAGAAAGAAAGG | ACGAGCGATGATATGACTCTATATACGGA | *Gossypium spp* |
| AMPL4000284 | GGTAACGTGTTGTTTGTCGCAG | GAGCTCATTGGCGAAACTTTGAAAA | *Gossypium spp* |
| AMPL4000285 | TGCGGTTTGGTTTCTGTGTGTAG | AACATATGAAAGTTCTAAGTTGCATACCCAT | *Gossypium spp* |
| AMPL4000286 | GGGAGGACTTCAGTCATTTAAAGGTT | AACAGGTTCAGTGTCCAAGGAAGC | *Gossypium spp* |
| AMPL4000287 | CGGTTTAGACATCACAACAAGTTATACCTC | CCTTAAACACAAATAAATCTTCCAGTCCAAA | *Gossypium spp* |
| AMPL4000288 | ATTGAAATTGGGTAACTTCTTCGCATTCA | AATCTCTCAACGCAGGGAATGAGG | *Gossypium spp* |
| AMPL4000290 | AAGTGCAAAAGATACAGGGAGTGAAAG | CAACGCCAGGATTGTTTAATGCTTTC | *Gossypium spp* |
| AMPL4000292 | TACTGCAGAAGAGACCGGTATAGTATTTA | CTCTCAATTAACCAATCTTGGTATGGCTTAA | *Gossypium spp* |
| AMPL4000293 | TACAAATCGACGAATGGAAATGCTAAGAA | TTTCGTAGCACTTTGTTCTATCGAGTC | *Gossypium spp* |
| AMPL4000295 | GAGAAAGTACTGATAATCTCATATGGTGCAT | GCTTGTTTCAGATGTGGATCTAAAGATCATA | *Gossypium spp* |
| AMPL4000296 | GTAAAGGTTCTTTGGCAACGATATGGG | ACCTAATCAAGCAAATAAGGTGGTTGAA | *Gossypium spp* |
| AMPL4000297 | CTCCTAGAACACATAACCACTAAAGTAGCA | TCCAAGCTTACGTCCGAGTAGTT | *Gossypium spp* |
| AMPL4000298 | CGTTAGTTCTTTCTTTTTCCACTATTACGCC | ACGCTCCTCTCCTAACAGAACAA | *Gossypium spp* |
| AMPL4000301 | GATAAAATGGAGTGTGAAACTCAACATGTT | AGATTCCTTCGTTAAAACCATCTTTGCTAG | *Gossypium spp* |
| AMPL4000303 | CATCAAATTCAACATGATACAACCACCTA | ACGTTACCTGGAATACGGATGAACTTATAA | *Gossypium spp* |
| AMPL4000304 | GAGATCATTCGGTTTTGGGTCTTAAATTTA | AAGAGAGTTACATTGTTCATGCACATAGTC | *Gossypium spp* |
| AMPL4000305 | TATCGGTACCCAACAGCCAAGTA | CACCCTTAACTCAAGTTTACGACAGTC | *Gossypium spp* |
| AMPL4000309 | TTTTCTTCCGTTCTTCATCTTCTTCAGTA | TGGCTGCTAAGCATTACAACTAAATATCTA | *Gossypium spp* |
| AMPL4000310 | ATACCACTATTTGTTGTTCACATCTAAGCAC | TGTAATGGTAATTAATCGGCCTTTTAGTGAA | *Gossypium spp* |
| AMPL4000311 | TTTTAGTACCCTAGCACCCATATCTGAT | GTGCTTCCACAATGCCCTATACAG | *Gossypium spp* |
| AMPL4000312 | GGTATGTTTTGGTGCCTTGGTCATATG | GAGGAAATAGGGAAACATAGTCGTGTG | *Gossypium spp* |
| AMPL4000313 | TTATTTTCTCTTCTCCTCCACTGTAGCTAC | CGATGGTGGTGGTCCTATGGTAG | *Gossypium spp* |
| AMPL4000314 | CTGAAATTATGTTGTGTACTCACAACCATCC | ACTCATCTATCCTACACACCGAGTTG | *Gossypium spp* |
| AMPL4000315 | ATTTTCAAAATCTGCTATTGTGGTTGCTG | CGGAGTACTCTCAATACTTCGTTGTACTAAT | *Gossypium spp* |
| AMPL4000316 | CCTCAAGCCACCAACCTTTCATT | GAAGAACCGATACGATTGGTAATTGTCATAA | *Gossypium spp* |
| AMPL4000317 | GCAAAAGAAATCAACAAGAGGTGTATCAAA | GCCTACCATCCTTATTGTTGGTGAC | *Gossypium spp* |
| AMPL4000318 | TCAACCGAAATGACGTGTCCAAC | GGTTGATTTTGCATCAGAATAGAACCAGG | *Gossypium spp* |
| AMPL4000319 | GTAGGCCCAAACCCAGCAGTATG | CGGCTGCTGGAATATCATTTTCGAC | *Gossypium spp* |
| AMPL4000320 | CGAGCTTAACTCGAGATATAATCGAGCT | CTCACAAGCTACTCTAGTTCGACTAAAAAA | *Gossypium spp* |
| AMPL4000321 | ACATCGCTGACATTCTTGTCTATTGGC | AAGAGTTTGTTGTCTTGGTCGATAGAGC | *Gossypium spp* |
| AMPL4000322 | ACAAGTTAAGTGCTTTCTCCCAAATACAT | GAACATTTGGCAGGCTATAGTAACTTGT | *Gossypium spp* |
| AMPL4000323 | GTAAGATGATAAGAAAATGTGCCGTAGAGTA | CAAAGCTTCAAGAGAGATTAAGGTTATCCAG | *Gossypium spp* |
| AMPL4000324 | TGAAAGGGCAATGAAGATTTTGAGGTAA | AATTTCTAAAAGGGCTTCCTTAGCAGTC | *Gossypium spp* |
| AMPL4000325 | ACAGACTATAAGAAGTTACGTCTCTCGATA | CTTAAACTCGCTTTCAGCTCTCTTACG | *Gossypium spp* |
| AMPL4000326 | CATCCAAACTAAGCCTAAGTACCCAA | CCCTAGGAGATAAGCATTTAAGGTGAAAAAT | *Gossypium spp* |
| AMPL4000327 | ACTCTATGTGTCCCTACTATGGTATGGG | TTGTGTTAGTTTAAGAGCTTTCATAAGCTCA | *Gossypium spp* |
| AMPL4000328 | CTGTCAGGTGATCTGGCATAACAC | ACTTCCAATGGAGATGAAGATGATACATGC | *Gossypium spp* |
| AMPL4000329 | TACGTTGCTCTCAAGTAAACCAAGGT | GGAGGATGATATAAAGTGGAGACTGATAGAG | *Gossypium spp* |
| AMPL4000330 | CCTCCTAAATAGTTTCTGCCAAAAATGACT | AACCTTTACCTATTCCTAGTAAAGTCTGGTA | *Gossypium spp* |
| AMPL4000331 | AAACTCACCGGTGGAGAAGATGA | TGACGTCTGAACTCAAATTGATATTACTCAT | *Gossypium spp* |
| AMPL4000332 | GGTAAGGGCATAAGATCTCAAGTACGA | TCTTCTACTGACTTCCCAGTAGCAG | *Gossypium spp* |
| AMPL4000333 | ATTGAATTGGTTGAAAAGTAAACTCGCATT | GCTTTTATTGCTTGTAAACCTCTCTTTCATA | *Gossypium spp* |
| AMPL4000334 | GAAGGCGTGACCAATGTTACCTAT | CCCAACACTAATGTAAGGAGAGAGGATTATC | *Gossypium spp* |
| AMPL4000335 | TGACTCTAGAATTCCTTCCACAATCCTA | TCGCTCCTTGTTAGGGTAGTCAA | *Gossypium spp* |
| AMPL4000336 | AATTGCAATGGAATAGCTTTTGGATAGCA | AATGATTTTCTAAAGTGAGAACAAGGGACTT | *Gossypium spp* |
| AMPL4000337 | GGAGAAACATGTAAACTTGTAAATGCGTT | TCTAGGAATGTGACGAATTCTTGTCTTTCA | *Gossypium spp* |
| AMPL4000338 | TCGTCAAACTAGCTGTAACATCCAAGG | CCTTAAGGAAGGGTGAATGTAACACCC | *Gossypium spp* |
| AMPL4000339 | CACGTCTTCCACGTACCTACAGC | CCCAAAGAGAAAATCACCTCTCCAATTTAC | *Gossypium spp* |
| AMPL4000340 | AAGTGGTGGGTTCCAAGATTCAAG | TCTCATCCTCACTAAGCTTTAGCTTTTGC | *Gossypium spp* |
| AMPL4000341 | ACAATCTGGAAATATTTAAGATGGAGGAGGATAA | TAGACACTACTCAAAGCTACTAACAATCCTT | *Gossypium spp* |
| AMPL4000342 | AGAAAACCTTCCTACAACTACTTGAAACTTT | GGGTCAAGCAAGACCTTGTTAAACC | *Gossypium spp* |
| AMPL4000344 | ATTAAGGTTTGCTTGTAAGTTTGGGATTTCG | AGTAACGTGGGCAAAAATTTTGGGG | *Gossypium spp* |
| AMPL4000345 | TGCATGATCTAGTTCTCTAGATGCATAAAA | GCACACGGTCGTGTATCTCTTGC | *Gossypium spp* |
| AMPL4000346 | ACAAACACGAACTAACTCAACATTTCACT | TTTTGACGGGTGAGTTTTGCAATTT | *Gossypium spp* |
| AMPL4000348 | GCATGTACCTGCAACGAACAACA | AAAAAGCAAGCAAATTAACACAAATAAAATGTAA | *Gossypium spp* |
| AMPL4000349 | TGGAAAATCATTAATCAATCAGCTGATCGAT | GAAATGACCTACAAGATGACTTGACAAAAAA | *Gossypium spp* |
| AMPL4000350 | CGCTAATGATGGGCAGGGTAAAT | TCTATCAAGGTGGTTCATCGTCCTAAC | *Gossypium spp* |
| AMPL4000351 | CTAATTAAATTCCTACGCAAGGCAAGTACA | AGCTTCTTCTACTCTCTCTCATTCTTTCTA | *Gossypium spp* |
| AMPL4000352 | GTAAAAGGGAGTGGTCGAAAGTAAGAAT | TCATACTTTGCATCGAACTTGCATTGG | *Gossypium spp* |
| AMPL4000353 | TACAGGAGAACCAATGATTACTAATGCTTGA | GGCCATTTACCCACTAGCTTTATATATTCTC | *Gossypium spp* |
| AMPL4000354 | TCCTTTTCTGTTTCTTCCTATTTTTCCCTG | AAGAATTTTCCGTTGGCAACGTTTAAA | *Gossypium spp* |
| AMPL4000355 | ATCCACTGCAATAAGAAGGCTTCTT | GATCCTGACTTCATATTTTCTCAGGGTATTT | *Gossypium spp* |
| AMPL4000356 | AAGAGCTTTTGTTTTAACATCTGCCATTTG | GCAAAGTACAAAGCTTTAGCTCAAAGTATAT | *Gossypium spp* |
| AMPL4000357 | GAATGCATGTTATGGTAAGGAATCTGGTAA | ACCGTTCTTTAGTTCTTTTCAGAAGGTG | *Gossypium spp* |
| AMPL4000358 | AACTGGGAGTGTTATCGTCGTTCG | ACCGCTACAGACATGGAAATCAACC | *Gossypium spp* |
| AMPL4000359 | CTTGGTAGTGGCAAGGAAAGGTC | AAGTTGATTTAGGAAAGGCCTTAGGAAATT | *Gossypium spp* |
| AMPL4000361 | CGACCCTAAAACGCGATAGAAATTTG | CCAAAAATTGGCCAAAATGGCTTACA | *Gossypium spp* |
| AMPL4000362 | CTGGACAAAGAAACTTGGACTGTTGG | TGCTGAGAGAGAATAATTGGTGTAAGTTTTT | *Gossypium spp* |
| AMPL4000363 | TGGTTTCCTCCCTAAATTCATTTATGCAT | TTTTAGGGTTTTCTCCTCTCAATTTCTCATC | *Gossypium spp* |
| AMPL4000364 | GTTTCTTGCTATTTTCTCATGGATGACAT | AGATGAGCACATGCGCTTCTATCA | *Gossypium spp* |
| AMPL4000365 | TCTAACCGTTACAGTATCCAATCTCATCAC | TGCGAAGGAATGATCTTTGTCCAAG | *Gossypium spp* |
| AMPL4000366 | CTGGGAAATTTAGTTCCTCTAGTCTTTTGAT | TTTTGTTTTTGCCTCATGTGCACTG | *Gossypium spp* |
| AMPL4000367 | CCCTTAACTTAGGCATCCCTCTAATAGC | GTGACCAGGGCATGCTAGAAAAA | *Gossypium spp* |
| AMPL4000368 | AACTGTCAGCTTATATGTTAAATGGTCCAT | TAGAGGGTCTCAGGTCTCTACCTG | *Gossypium spp* |
| AMPL4000372 | TTTAGAGTGGCTTTAATGGGCGATG | CTCACTAGGCTATTCACATTCAATCACATT | *Gossypium spp* |
| AMPL4000373 | TTGTTATTCGAAAAGGATGTAGTTAGGGTTTT | GTGTTGGGTCCTTGAGTACTATCATGC | *Gossypium spp* |

| | | | |
|---|---|---|---|
| AMPL4000374 | TTCTTTTCTTAGTCATCGGCCTCAAAT | GAATTTTTGGAAGATCTAGTTCTTTTGCGAG | *Gossypium spp* |
| AMPL4000376 | GTGGACATAGCCACTAGAGATTGCA | CAGCCTGTCCTACATGTCTGTCA | *Gossypium spp* |
| AMPL4000377 | TTGTGCCCTAACTTTTGCTTAAATTGTTC | AAAGTTGGGAACGCCTAGTATCCG | *Gossypium spp* |
| AMPL4000378 | GTCTTTGCAACCAAAACAAAAAGGCC | AACATGTTTCGCATGATTCTACAACAATACA | *Gossypium spp* |
| AMPL4000379 | GTGTCTGACTCCGTGAGATCATCA | CATGCCTTAATTCCCTATTTGTCTTAAGGA | *Gossypium spp* |
| AMPL4000380 | GGACTCACTTGTCGAAATTGTGGT | CATAAACTGATGAAACCAACCATTCATCATT | *Gossypium spp* |
| AMPL4000381 | GTCTTAACTAAACTCTAACACCACTTACCAC | GGAGGATAATGTATCAAGTTATTGTTTGGCA | *Gossypium spp* |
| AMPL4000383 | TCAGTCTAGACGGGCTTTTTGACA | TGCCTACCAAGACTATTCCTCTTTGAG | *Gossypium spp* |
| AMPL4000384 | TGATTGTGTATCATGTTATGTACGCGTAAT | AGAAATTTAGTCCACAGACTCATTCATGTA | *Gossypium spp* |
| AMPL4000385 | GAGCAACATTTGTTTCTATTTTTCTTTGCGT | GGAATAATTCAATCCACCCTCCACAAG | *Gossypium spp* |
| AMPL4000386 | TGTCTGGTTCAGACTTTTATGTTCCCC | GAAACAAATTCCTTGTATTGCGAGATGTC | *Gossypium spp* |
| AMPL4000387 | GATCCACAGAAAACCCAATCTTTCTTATTCT | TCTTACGCAGGTGTATAACTCCTTTGC | *Gossypium spp* |
| AMPL4000388 | CTACTAGAGCCTTTGCGGGTTTC | GGAGTGTAAAACAACCACATGCAGTAC | *Gossypium spp* |
| AMPL4000389 | TAGATGTGAAGGTGGGATTAGGGTC | CTTTCCAAGCAGTGAGTCAGTTCTG | *Gossypium spp* |
| AMPL4000390 | TCTTAAGGCCTCTTGGTATCACCTT | AAAATAATGCGAAACCGAACATCGATACA | *Gossypium spp* |
| AMPL4000391 | ACTGAAGAATGGAACCGAAAGCAG | AAGTACAGGGACTACATTACAAATTTTCGTA | *Gossypium spp* |
| AMPL4000392 | ACTTCAAGTTGAAAAGGGTTCTGGTTC | GTGTGATTTACCTAAATACTGAGCTAGTCAT | *Gossypium spp* |
| AMPL4000393 | CGATTATTCTGTAAGTCGCACCTGTTT | TGTTATTTTTCATTTGCAGCTCAACTTCAAA | *Gossypium spp* |
| AMPL4000394 | GGGCTGCATTTCAAGGTCTAAAAGA | TGACTTAGGCGAGTTGGAGACAA | *Gossypium spp* |
| AMPL4000395 | CTAAGGGCCAATTCGGGTAGTTC | ATTAGAAATAGAGGGACTTGTAATTCGGTTA | *Gossypium spp* |
| AMPL4000396 | TATCTCATGCCAACAACTTGTACCAAG | CATCACAGTCCAAGGTTCCAGTTAC | *Gossypium spp* |
| AMPL4000397 | TGCTAAAGTAGCACGCCAATTTTGA | GTTAACACCTCTTCCATGATGAGCTT | *Gossypium spp* |
| AMPL4000398 | CAATTTTCCCAAGGCAAGGCTTG | GAGGGCAATACACTTATGATAAGGTCATTG | *Gossypium spp* |
| AMPL4000399 | AGAGTGAGAATAGGATCGAGCTCAAG | CTCTAACCAGTTTTCAGTTAAAGAAAGGCC | *Gossypium spp* |
| AMPL4000400 | CGCCATAGCATTTTTAGTTGTTTGCT | CAAGTAATAAAGTGATGAGAGAGATTGCCTC | *Gossypium spp* |
| AMPL4000401 | CTCACTCCAAATTCCTTTTTAACTCATACGT | GTGTTACAAAATTCCATAACCATACACGTG | *Gossypium spp* |
| AMPL4000402 | AGGAAGTATGACTGGTAAGATTATGAGTGAA | ACATATTTGCCACACTGGTATTTTCTCATT | *Gossypium spp* |
| AMPL4000403 | AAGGATGAATCATGCTTTCTTCTTGAAAAA | GAGGTAAATGAAGTGGGTTTAGAGGTAAGA | *Gossypium spp* |
| AMPL4000405 | CCTTAGTCTTAGGTGAAGTGCAATGTT | CAAAACTAAAGAGCATGCTTCAGATACGG | *Gossypium spp* |
| AMPL4000406 | AACATATGATGGACTTGCACTTGGTTAA | AAATGGTGGTGAAAATAAGGAGGCTAG | *Gossypium spp* |
| AMPL4000407 | GCTTGATCAAAGCAACGATTTTCTAGCG | TGATCTATGAACCTTAGCCCAGTGAAT | *Gossypium spp* |
| AMPL4000408 | GTTTTGGGTCCCTCTAGGTGAAGT | AGTTTCAAGTTAGAATCACACAACTGGTG | *Gossypium spp* |
| AMPL4000409 | CGAATGTGGTTACGTTCGGATTTT | CCCTTAACAGGCCAAAATTTTCTAAAGAAAC | *Gossypium spp* |
| AMPL4000410 | TCGTTTCCTCTCTCCGACGAATAT | TGGGAGATTGAGTTTCAGAGTTCACG | *Gossypium spp* |
| AMPL4000412 | CTCGATTCTTGTTTCTTGTGCCAAAAC | AAACCAACATTTCCTTAAACAAACCCAAA | *Gossypium spp* |
| AMPL4000413 | GATGTTTTCCTTGCTGTATTGAAGTGGT | CGCCTTCACCATTGATAAAATAATGTCAAAA | *Gossypium spp* |
| AMPL4000414 | AAGGCAGCAAGGTATAATGAGTAAGACC | GCATTAGTCCCATAGTGGGCTCTATG | *Gossypium spp* |
| AMPL4000415 | CCAACGGCATCACACGAAATTCA | TAAGAAAAAGGGTGGATCGATGAGATTGT | *Gossypium spp* |
| AMPL4000416 | CGTTTGAACCTCAGAATGAGAAGGG | GTAGGTTCTGCCCAATCTTCCAC | *Gossypium spp* |
| AMPL4000417 | CGATGCCGATATCATATCTTGCATCC | TGGGTAAGGATTCCATCCTATGATCCC | *Gossypium spp* |
| AMPL4000418 | CCCACTGGAATGGGTTTTGAAACA | CTGGACACATATCTTACTCAAAAAGAGGTTC | *Gossypium spp* |
| AMPL4000419 | CGGACGTCTGATTTCTCTATTAATCCCT | ACCTTAAACACGGTTATAAGAGGTTAAAGGG | *Gossypium spp* |
| AMPL4000420 | CCTGTTGTCGAGGAGAATGTGTG | GACCTTGGAGGAAGATGATAGTAGTGC | *Gossypium spp* |
| AMPL4000421 | GTGTATTTATGGTAAGAAGGCACCAAGTA | CTCAAAGAGAATAACGACCATGGGTAAAA | *Gossypium spp* |
| AMPL4000422 | CTCTTTGCTTCTCCAATTGTCATGGT | GCCAAAAGGCATAACCAACAACTCA | *Gossypium spp* |
| AMPL4000423 | TCCATGGATGCTCGTTGGAAAAC | AAGTTGTCTGATAGGTCCAGTAATGCC | *Gossypium spp* |
| AMPL4000424 | AGCAAAACTACCAAATTGCACTAACATATCT | CATTAGAGGGATAATTTTGTTACTAGACCGATAT | *Gossypium spp* |
| AMPL4000425 | CGATTGGAAAAGTTTGCGAGATTTGAT | CCGTTAAAATAAGCTAGCAAGAGTTGTTCA | *Gossypium spp* |
| AMPL4000426 | ACTTCCTCTATGATATCTAATGCTGGACAT | GGGCTTGTGTTGCCTTCCTTTAG | *Gossypium spp* |
| AMPL4000427 | CATCCAAACTCACCCTTAGAATGCT | TATTGGAATGATGGAGTGGGTAAAACATGA | *Gossypium spp* |
| AMPL4000428 | GGGAAGGAGACAAAGATATAGTTGAATAGTT | GGCGGAAGCCTTAGAACCTATGA | *Gossypium spp* |
| AMPL4000429 | ACATGCCACACAAAAGATCAACATCA | TACGCGGTTTAGTCACGACTTGT | *Gossypium spp* |
| AMPL4000430 | TTGCCCAGGCGAACTAAACAATT | ATGTCTTCCTTGTAAGAAGTTGGATGTACA | *Gossypium spp* |
| AMPL4000431 | CACTAGACGATTTCTTCTGGTAACCAAA | AAGGTTTGATGGAATGTATTGTGAGAAGTAT | *Gossypium spp* |
| AMPL4000432 | GGTACAATTTTCATGGTGCGTGTG | AATAAAAATACAGTTTCACAATTTGAATAGCCTA | *Gossypium spp* |
| AMPL4000434 | ATGGCTTAGACCTACAATTGTATGAATCCT | GGCTATAGGATGCTACAACAATTAACGGA | *Gossypium spp* |
| AMPL4000435 | GAACAAGTCAAGCTCAACATGTTGAA | GCTCCACCACCGGATACTATGTG | *Gossypium spp* |
| AMPL4000437 | ACACAAAACACCCTTACAACATGCC | ATGAGGTTCTTATTGGAATTGTCAAGGTATG | *Gossypium spp* |
| AMPL4000438 | TGGTAGGTCAAATCACAAAACAAATTGTGT | TACCTGTCTAGGATGATTTACAACTAGGAGT | *Gossypium spp* |
| AMPL4000439 | GTCGCCACATGACACCTTAAAAGC | ATTAAGGTATTCACCGTCTCAGATGTGG | *Gossypium spp* |
| AMPL4000440 | TTAGATGTGGTCAAGCCATATTCAAAATCTT | TTAGGGTCACTATAGGAGAGAATGAAGTGA | *Gossypium spp* |
| AMPL4000441 | CTTATGTGAAAAGTGGTGGACGTCTAA | CGAGGGTCAATAGAGGGATGTTATGATTATT | *Gossypium spp* |
| AMPL4000442 | CCAAGGCCTCTACTTTGGGAATC | TACGAGTTGGAATGAAGTCTTGATCGA | *Gossypium spp* |
| AMPL4000443 | TCAAACGTCAAAATTTTCGAACGAAATTTGC | GGTGACCTGTCGGAGGAAATTTATATGC | *Gossypium spp* |
| AMPL4000445 | CTTCCACTTCCACATAGCCGTTG | CAACGGACGAAATGATTTTGGAGTTG | *Gossypium spp* |
| AMPL4000447 | TGAGATTGCCCTCAGTTGCAAAG | ATCTAGGGTCAATCAATGCATACACAATAAC | *Gossypium spp* |
| AMPL4000448 | TTCTCGTTAAATTCTCATTATCGTTGGACTT | AAAAATTTAGTCAAATTCTGGCTTGGCTAG | *Gossypium spp* |
| AMPL4000449 | GGTATTACTTGAATCATAGGTTTGCTTGGTA | AAAACAAGTTCGAAACTGAATGAAGACCA | *Gossypium spp* |
| AMPL4000450 | ACAAGCAATTTCTCTTGACTATTTTCGTATA | GTGAGCTGCTTTCTTTCAAGTGGC | *Gossypium spp* |
| AMPL4000452 | ATGCGTCCACACCCAAAACAAAAGA | TGACTGCTCCCATTGAGCAATCC | *Gossypium spp* |
| AMPL4000453 | TTAAACACCAACAATTTCTGTCATTTGCTT | CAAAAATTGAAGAATCGGGTGTTTCTGTC | *Gossypium spp* |
| AMPL4000454 | GGGTCATCGATGACGCTATTAATCTT | CCACTCAAGTAATCCGTAATACGGTTATTTT | *Gossypium spp* |
| AMPL4000455 | GTCCGTGTTTTTGTGAGATAATAGTTGTT | TTACTGATATGACATGCTAGGAGGTTCTA | *Gossypium spp* |
| AMPL4000456 | CAATGCAAAAGTATGACACACCGAAA | TCTGAGGGAATCTGATGAGTGGTTAG | *Gossypium spp* |
| AMPL4000457 | CGATTGAAGTAGAAGAAGAACCCTTAACCA | AACACCATTGTTATACCCTCGAAGTTTTAA | *Gossypium spp* |
| AMPL4000458 | GCATCGACATAAAATAGTGTTGATTCCAAA | TCAACCAAATTCATCCAATTGGCAAGT | *Gossypium spp* |
| AMPL4000459 | TTACAACCTGGCTCCTCTATGTTAAATGT | GCTCCTGCAATGCTGGAGTTAGC | *Gossypium spp* |
| AMPL4000460 | AAATGGGCTCGGTTTGGACTTAT | GAAATTAACCCACTTTGATGAGTAGTGGAC | *Gossypium spp* |
| AMPL4000461 | CTCTTTATAGAGTCCAAACTTAAGCTCGTT | GGTCAAATGAGGCCCATTTAATTAATTGACC | *Gossypium spp* |
| AMPL4000462 | TTTTTAGGTTTAAATTTGCCTGAGTCATCAT | TGCATGGTGAATGAAGATCTGTGTTC | *Gossypium spp* |
| AMPL4000463 | CGTGTCTGCCCTGCACTTTATTT | AGGTCGAAACACACAGGAATACGA | *Gossypium spp* |

| | | | |
|---|---|---|---|
| AMPL4000464 | AAGAGGCAGACTGTTCATTATTCATATTTGA | AACCAAGCATAGACACCTTAAGCATAATA | *Gossypium spp* |
| AMPL4000465 | GATTTTCATCTAACAACCTCAGAGGTTCAG | GCTAAGCGTTTTGGATGTGTGTTTGA | *Gossypium spp* |
| AMPL4000466 | CTCGGCGTTGTAGTGGTTGTTATAT | TGAGAGTGTGTTGACACTTTGAACAG | *Gossypium spp* |
| AMPL4000467 | CCAAATCTCTAATTGATACTCCTCCAATGTG | GCATAGCCGTCAACCTCTTATAAGTATTTTT | *Gossypium spp* |
| AMPL4000468 | AGGAAAGCACTCAACTGTAACTTACAAATA | CAATTTGGTCCTTTCCATCTAATTAGACACG | *Gossypium spp* |
| AMPL4000469 | CGAGTGACTAGAAAACGATAGGATTGATGA | TGGTGATGAAAAGCGTAAATGGTATAGGT | *Gossypium spp* |
| AMPL4000470 | TAAAACAAAACAAAGGCTAATTGGTCGAAAG | AAAATGCTAATGGCAGCCTACTCTATAGA | *Gossypium spp* |
| AMPL4000471 | ATTTCAAATAGGACACGGTCCTTAAAAGAT | ATCGAGATTGGAGATGAGCTAGGTAAT | *Gossypium spp* |
| AMPL4000472 | GGAAATACATTTTCAAGCAACATCATTGGA | AATTTAACGTACAGGGACTAATTTGTCCAT | *Gossypium spp* |
| AMPL4000473 | AAAACCCAGTTTGAAGAAACAAATAAACCTT | CGTATGGGATGCGAAGGGTAAAGG | *Gossypium spp* |
| AMPL4000474 | AAATACAATGCCTCACGATATGCTTTTTAA | AAACTCACTGAAATTATGACATGTCTACGTG | *Gossypium spp* |
| AMPL4000476 | TGTATGATAGACACGAAAACTCTAAACCACC | GTACTTGAGGAACAAGGAATATCAGATGTA | *Gossypium spp* |
| AMPL4000477 | ATCTAAAGCTTCGAGCTCAACCTAAGT | GTGGAGTGGTTACAAGTAATCAACTAATCTT | *Gossypium spp* |
| AMPL4000478 | CCTGGTCGCAAGTCTAGAATTTTGA | GACTCGCATGGGAGTTAGAGATCG | *Gossypium spp* |
| AMPL4000479 | AGCGCCAGATGCAATTATTTGTCG | ATATGTGCACCATTAGTCCGCATG | *Gossypium spp* |
| AMPL4000480 | TGTGTTCGAAAACCATATCGGATTGTG | GTAAAACTCGTTTGCCGATGCATG | *Gossypium spp* |
| AMPL4000481 | CAAAATGACTGAGACACGAGGTAAGTT | CCTTAAGGCAATCCGATTGAATTAACATACT | *Gossypium spp* |
| AMPL4000482 | GGAAAACTTCAGATCTGGCCCTTTT | CACCTGTGTTCGGCTTCTAACAC | *Gossypium spp* |
| AMPL4000483 | ACACTAATTCGTTGATTTCATTGCTATGCT | TTTTGAAACACGATAATTTTTGAACACGAG | *Gossypium spp* |
