## Supplementary table 2 for "Multiple nucleotide polymorphism DNA markers for the accurate evaluation of genetic variations"

**Supplementary Table 2. Statistics of sequencing data for rice and contton varieties**

| Sample code | Species | Number of paired reads | MNP loci detected | | | Replicate * |
|---|---|---|---|---|---|---|
| | | | Average number | Average coverage folds | Ratio of the detected MNP loci | |
| OSD205093 | cotton | 232,437 | 935 | 248.60 | 99.15% | replicate1 |
| OSD205094 | cotton | 390,325 | 941 | 414.80 | 99.79% | replicate1 |
| OSD205095 | cotton | 333,586 | 939 | 355.26 | 99.58% | replicate1 |
| OSD205096 | cotton | 361,051 | 874 | 413.10 | 92.68% | replicate1 |
| OSD205097 | cotton | 274,898 | 914 | 300.76 | 96.92% | replicate1 |
| OSD205098 | cotton | 585,269 | 941 | 621.96 | 99.79% | replicate1 |
| OSD205099 | cotton | 593,207 | 941 | 630.40 | 99.79% | replicate1 |
| OSD205100 | cotton | 227,901 | 936 | 243.48 | 99.26% | replicate1 |
| OSD205101 | cotton | 294,534 | 941 | 313.00 | 99.79% | replicate1 |
| OSD205102 | cotton | 590,566 | 941 | 627.59 | 99.79% | replicate1 |
| OSD205103 | cotton | 503,385 | 941 | 534.95 | 99.79% | replicate1 |
| OSD205104 | cotton | 452,966 | 934 | 484.97 | 99.05% | replicate1 |
| OSD205105 | cotton | 341,247 | 941 | 362.64 | 99.79% | replicate1 |
| OSD205106 | cotton | 402,534 | 939 | 428.68 | 99.58% | replicate1 |
| OSD205107 | cotton | 292,027 | 938 | 311.33 | 99.47% | replicate1 |
| OSD205108 | cotton | 280,032 | 936 | 299.18 | 99.26% | replicate1 |
| OSD205109 | cotton | 371,614 | 942 | 394.49 | 99.89% | replicate1 |
| OSD205110 | cotton | 231,321 | 937 | 246.87 | 99.36% | replicate1 |
| OSD205111 | cotton | 499,939 | 941 | 531.28 | 99.79% | replicate1 |
| OSD205112 | cotton | 386,923 | 939 | 412.06 | 99.58% | replicate1 |
| OSD101198 | cotton | 420,930 | 942 | 446.85 | 99.89% | replicate1 |
| OSD101201 | cotton | 553,338 | 942 | 587.41 | 99.89% | replicate1 |
| OSD101202 | cotton | 310,546 | 942 | 329.67 | 99.89% | replicate1 |
| OSD101203 | cotton | 345,218 | 940 | 367.25 | 99.68% | replicate1 |
| OSD101204 | cotton | 390,010 | 938 | 415.79 | 99.47% | replicate1 |
| OSD101205 | cotton | 513,701 | 940 | 546.49 | 99.68% | replicate1 |
| OSD101207 | cotton | 598,186 | 941 | 635.69 | 99.79% | replicate1 |
| OSD101208 | cotton | 349,084 | 940 | 371.37 | 99.68% | replicate1 |
| OSD101209 | cotton | 301,508 | 942 | 320.07 | 99.89% | replicate1 |
| OSD101210 | cotton | 336,919 | 942 | 357.66 | 99.89% | replicate1 |
| OSD101211 | cotton | 384,104 | 942 | 407.75 | 99.89% | replicate1 |
| OSD101212 | cotton | 438,595 | 942 | 465.60 | 99.89% | replicate1 |
| OSD101213 | cotton | 675,672 | 942 | 717.27 | 99.89% | replicate1 |
| OSD101214 | cotton | 432,203 | 940 | 459.79 | 99.68% | replicate1 |
| OSD101216 | cotton | 251,641 | 938 | 268.27 | 99.47% | replicate1 |
| OSD101217 | cotton | 191,873 | 937 | 204.77 | 99.36% | replicate1 |
| OSD101223 | cotton | 118,067 | 856 | 137.93 | 90.77% | replicate1 |
| OSD101224 | cotton | 455,564 | 942 | 483.61 | 99.89% | replicate1 |
| OSD101225 | cotton | 340,994 | 874 | 390.15 | 92.68% | replicate1 |
| OSD101231 | cotton | 313,284 | 940 | 333.28 | 99.68% | replicate1 |
| OSD101232 | cotton | 287,830 | 938 | 306.86 | 99.47% | replicate1 |
| OSD101233 | cotton | 485,152 | 939 | 516.67 | 99.58% | replicate1 |
| OSD101235 | cotton | 254,459 | 940 | 270.70 | 99.68% | replicate1 |
| OSD205093 | cotton | 721,415 | 940 | 767.46 | 99.68% | replicate2 |
| OSD205094 | cotton | 946,404 | 941 | 1005.74 | 99.79% | replicate2 |
| OSD205095 | cotton | 557,804 | 941 | 592.78 | 99.79% | replicate2 |
| OSD205096 | cotton | 888,419 | 880 | 1009.57 | 93.32% | replicate2 |
| OSD205097 | cotton | 748,746 | 942 | 794.85 | 99.89% | replicate2 |
| OSD205098 | cotton | 780,898 | 941 | 829.86 | 99.79% | replicate2 |
| OSD205099 | cotton | 719,015 | 941 | 764.10 | 99.79% | replicate2 |
| OSD205100 | cotton | 841,327 | 937 | 897.89 | 99.36% | replicate2 |
| OSD205101 | cotton | 844,991 | 942 | 897.02 | 99.89% | replicate2 |
| OSD205102 | cotton | 977,127 | 941 | 1038.39 | 99.79% | replicate2 |
| OSD205103 | cotton | 856,472 | 941 | 910.17 | 99.79% | replicate2 |
| OSD205104 | cotton | 772,623 | 934 | 827.22 | 99.05% | replicate2 |
| OSD205105 | cotton | 776,420 | 941 | 825.10 | 99.79% | replicate2 |
| OSD205106 | cotton | 422,377 | 939 | 449.82 | 99.58% | replicate2 |
| OSD205107 | cotton | 623,458 | 939 | 663.96 | 99.58% | replicate2 |
| OSD205108 | cotton | Fail | / | / | / | replicate2 |
| OSD205109 | cotton | 622,739 | 942 | 661.08 | 99.89% | replicate2 |
| OSD205110 | cotton | 771,477 | 940 | 820.72 | 99.68% | replicate2 |
| OSD205111 | cotton | 869,691 | 941 | 924.22 | 99.79% | replicate2 |
| OSD205112 | cotton | 918,233 | 940 | 976.84 | 99.68% | replicate2 |
| OSD101198 | cotton | 954,461 | 943 | 1012.15 | 100.00% | replicate2 |
| OSD101201 | cotton | 992,194 | 943 | 1052.17 | 100.00% | replicate2 |
| OSD101202 | cotton | 922,140 | 942 | 978.92 | 99.89% | replicate2 |
| OSD101203 | cotton | 711,172 | 940 | 756.57 | 99.68% | replicate2 |
| OSD101204 | cotton | 839,683 | 938 | 895.18 | 99.47% | replicate2 |

| | | | | | | |
|---|---|---|---|---|---|---|
| OSD101205 | cotton | Fail | / | / | / | replicate2 |
| OSD101207 | cotton | 823,381 | 941 | 875.01 | 99.79% | replicate2 |
| OSD101208 | cotton | 659,679 | 940 | 701.79 | 99.68% | replicate2 |
| OSD101209 | cotton | 732,089 | 942 | 777.16 | 99.89% | replicate2 |
| OSD101210 | cotton | 647,186 | 943 | 686.31 | 100.00% | replicate2 |
| OSD101211 | cotton | 721,154 | 942 | 765.56 | 99.89% | replicate2 |
| OSD101212 | cotton | 886,401 | 942 | 940.98 | 99.89% | replicate2 |
| OSD101213 | cotton | 1,044,369 | 942 | 1108.67 | 99.89% | replicate2 |
| OSD101214 | cotton | 639,930 | 942 | 679.33 | 99.89% | replicate2 |
| OSD101216 | cotton | 695,234 | 940 | 739.61 | 99.68% | replicate2 |
| OSD101217 | cotton | 804,302 | 940 | 855.64 | 99.68% | replicate2 |
| OSD101223 | cotton | 886,321 | 881 | 1006.04 | 93.43% | replicate2 |
| OSD101224 | cotton | 652,458 | 942 | 692.63 | 99.89% | replicate2 |
| OSD101225 | cotton | 698,792 | 879 | 794.99 | 93.21% | replicate2 |
| OSD101231 | cotton | 587,161 | 941 | 623.98 | 99.79% | replicate2 |
| OSD101232 | cotton | 642,958 | 939 | 684.73 | 99.58% | replicate2 |
| OSD101233 | cotton | 310,432 | 939 | 330.60 | 99.58% | replicate2 |
| OSD101235 | cotton | 516,787 | 940 | 549.77 | 99.68% | replicate2 |
| OSD205093 | cotton | 709,550 | 940 | 754.84 | 99.68% | replicate3 |
| OSD205094 | cotton | 711,161 | 941 | 755.75 | 99.79% | replicate3 |
| OSD205095 | cotton | 500,448 | 940 | 532.39 | 99.68% | replicate3 |
| OSD205096 | cotton | 546,792 | 876 | 624.19 | 92.90% | replicate3 |
| OSD205097 | cotton | 631,654 | 942 | 670.55 | 99.89% | replicate3 |
| OSD205098 | cotton | 830,338 | 941 | 882.40 | 99.79% | replicate3 |
| OSD205099 | cotton | 416,194 | 941 | 442.29 | 99.79% | replicate3 |
| OSD205100 | cotton | 330,387 | 937 | 352.60 | 99.36% | replicate3 |
| OSD205101 | cotton | 994,570 | 942 | 1055.81 | 99.89% | replicate3 |
| OSD205102 | cotton | 631,976 | 941 | 671.60 | 99.79% | replicate3 |
| OSD205103 | cotton | 664,790 | 941 | 706.47 | 99.79% | replicate3 |
| OSD205104 | cotton | 826,992 | 934 | 885.43 | 99.05% | replicate3 |
| OSD205105 | cotton | 789,244 | 941 | 838.73 | 99.79% | replicate3 |
| OSD205106 | cotton | 640,905 | 940 | 681.81 | 99.68% | replicate3 |
| OSD205107 | cotton | 428,083 | 939 | 455.89 | 99.58% | replicate3 |
| OSD205108 | cotton | 594,698 | 942 | 631.31 | 99.89% | replicate3 |
| OSD205109 | cotton | 895,196 | 942 | 950.31 | 99.89% | replicate3 |
| OSD205110 | cotton | 624,647 | 940 | 664.52 | 99.68% | replicate3 |
| OSD205111 | cotton | 671,289 | 941 | 713.38 | 99.79% | replicate3 |
| OSD205112 | cotton | 880,223 | 940 | 936.41 | 99.68% | replicate3 |
| OSD101198 | cotton | 994,882 | 943 | 1055.02 | 100.00% | replicate3 |
| OSD101201 | cotton | 437,867 | 942 | 464.83 | 99.89% | replicate3 |
| OSD101202 | cotton | 695,137 | 942 | 737.94 | 99.89% | replicate3 |
| OSD101203 | cotton | 890,815 | 940 | 947.68 | 99.68% | replicate3 |
| OSD101204 | cotton | 789,499 | 938 | 841.68 | 99.47% | replicate3 |
| OSD101205 | cotton | 679,402 | 940 | 722.77 | 99.68% | replicate3 |
| OSD101207 | cotton | 762,593 | 941 | 810.41 | 99.79% | replicate3 |
| OSD101208 | cotton | 820,285 | 940 | 872.64 | 99.68% | replicate3 |
| OSD101209 | cotton | 522,423 | 942 | 554.59 | 99.89% | replicate3 |
| OSD101210 | cotton | 848,812 | 943 | 900.12 | 100.00% | replicate3 |
| OSD101211 | cotton | 823,489 | 942 | 874.19 | 99.89% | replicate3 |
| OSD101212 | cotton | 961,440 | 942 | 1020.64 | 99.89% | replicate3 |
| OSD101213 | cotton | 655,075 | 942 | 695.41 | 99.89% | replicate3 |
| OSD101214 | cotton | 750,771 | 942 | 797.00 | 99.89% | replicate3 |
| OSD101216 | cotton | 909,205 | 941 | 966.21 | 99.79% | replicate3 |
| OSD101217 | cotton | 218,033 | 937 | 232.69 | 99.36% | replicate3 |
| OSD101223 | cotton | 576,855 | 877 | 657.76 | 93.00% | replicate3 |
| OSD101224 | cotton | 709,132 | 942 | 752.79 | 99.89% | replicate3 |
| OSD101225 | cotton | 665,494 | 877 | 758.83 | 93.00% | replicate3 |
| OSD101231 | cotton | 858,028 | 941 | 911.83 | 99.79% | replicate3 |
| OSD101232 | cotton | 691,280 | 939 | 736.19 | 99.58% | replicate3 |
| OSD101233 | cotton | 624,266 | 939 | 664.82 | 99.58% | replicate3 |
| OSD101235 | cotton | 555,727 | 941 | 590.57 | 99.79% | replicate3 |
| OSD202062 | rice | 303,049 | 911 | 332.66 | 97.96% | replicate1 |
| OSD202175 | rice | 402,689 | 924 | 435.81 | 99.35% | replicate1 |
| OSD202180 | rice | 396,367 | 911 | 435.09 | 97.96% | replicate1 |
| OSD202183 | rice | 538,057 | 926 | 581.06 | 99.57% | replicate1 |
| OSD202200 | rice | 618,220 | 926 | 667.62 | 99.57% | replicate1 |
| OSD202201 | rice | 496,946 | 928 | 535.50 | 99.78% | replicate1 |
| OSD202227 | rice | 670,784 | 930 | 721.27 | 100.00% | replicate1 |
| OSD202237 | rice | 670,762 | 929 | 722.03 | 99.89% | replicate1 |
| OSD202243 | rice | 556,322 | 930 | 598.20 | 100.00% | replicate1 |
| OSD202246 | rice | 670,344 | 927 | 723.13 | 99.68% | replicate1 |
| OSD202358 | rice | 544,589 | 926 | 588.11 | 99.57% | replicate1 |

| | | | | | | |
|---|---|---|---|---|---|---|
| OSD202412 | rice | 352,660 | 923 | 382.08 | 99.25% | replicate1 |
| OSD202440 | rice | 755,686 | 928 | 814.32 | 99.78% | replicate1 |
| OSD202557 | rice | 748,547 | 929 | 805.76 | 99.89% | replicate1 |
| OSD202663 | rice | 655,677 | 921 | 711.92 | 99.03% | replicate1 |
| OSD202674 | rice | 507,996 | 930 | 546.23 | 100.00% | replicate1 |
| OSD202680 | rice | 988,794 | 926 | 1067.81 | 99.57% | replicate1 |
| OSD202712 | rice | 820,428 | 929 | 883.13 | 99.89% | replicate1 |
| OSD202723 | rice | 663,429 | 929 | 714.13 | 99.89% | replicate1 |
| OSD202740 | rice | 542,908 | 927 | 585.66 | 99.68% | replicate1 |
| OSD202745 | rice | 362,139 | 887 | 408.27 | 95.38% | replicate1 |
| OSD202748 | rice | 439,447 | 917 | 479.22 | 98.60% | replicate1 |
| OSD202751 | rice | 308,123 | 871 | 353.76 | 93.66% | replicate1 |
| OSD202763 | rice | 477,353 | 924 | 516.62 | 99.35% | replicate1 |
| OSD202766 | rice | 379,146 | 924 | 410.33 | 99.35% | replicate1 |
| OSD202767 | rice | 454,011 | 927 | 489.76 | 99.68% | replicate1 |
| OSD202768 | rice | 578,260 | 929 | 622.45 | 99.89% | replicate1 |
| OSD202781 | rice | 593,573 | 924 | 642.40 | 99.35% | replicate1 |
| OSD202792 | rice | 819,131 | 928 | 882.68 | 99.78% | replicate1 |
| OSD202799 | rice | 641,780 | 929 | 690.83 | 99.89% | replicate1 |
| OSD202801 | rice | 810,646 | 930 | 871.66 | 100.00% | replicate1 |
| OSD202802 | rice | 468,903 | 929 | 504.74 | 99.89% | replicate1 |
| OSD202809 | rice | 979,949 | 930 | 1053.71 | 100.00% | replicate1 |
| OSD202811 | rice | 680,703 | 929 | 732.73 | 99.89% | replicate1 |
| OSD202815 | rice | 526,156 | 927 | 567.59 | 99.68% | replicate1 |
| OSD202904 | rice | 472,398 | 927 | 509.60 | 99.68% | replicate1 |
| OSD202906 | rice | 420,322 | 920 | 456.87 | 98.92% | replicate1 |
| OSD202912 | rice | 841,007 | 929 | 905.28 | 99.89% | replicate1 |
| OSD202914 | rice | 331,105 | 917 | 361.07 | 98.60% | replicate1 |
| OSD202936 | rice | 513,294 | 921 | 557.32 | 99.03% | replicate1 |
| OSD202938 | rice | 602,626 | 927 | 650.08 | 99.68% | replicate1 |
| OSD202958 | rice | 457,730 | 923 | 495.92 | 99.25% | replicate1 |
| OSD203003 | rice | Fail | / | / | / | replicate1 |
| OSD203044 | rice | 809,476 | 930 | 870.40 | 100.00% | replicate1 |
| OSD203080 | rice | 1,162,907 | 929 | 1251.78 | 99.89% | replicate1 |
| OSD203081 | rice | 588,327 | 926 | 635.34 | 99.57% | replicate1 |
| OSD203095 | rice | 716,153 | 929 | 770.89 | 99.89% | replicate1 |
| OSD202062 | rice | 1,115,040 | 929 | 1200.26 | 99.89% | replicate2 |
| OSD202175 | rice | 755,735 | 927 | 815.25 | 99.68% | replicate2 |
| OSD202180 | rice | 516,945 | 928 | 557.05 | 99.78% | replicate2 |
| OSD202183 | rice | 108,890 | 811 | 134.27 | 87.20% | replicate2 |
| OSD202200 | rice | 1,043,521 | 929 | 1123.27 | 99.89% | replicate2 |
| OSD202201 | rice | 953,852 | 929 | 1026.75 | 99.89% | replicate2 |
| OSD202227 | rice | 1,255,184 | 929 | 1351.11 | 99.89% | replicate2 |
| OSD202237 | rice | 1,181,223 | 930 | 1270.13 | 100.00% | replicate2 |
| OSD202243 | rice | 1,253,160 | 930 | 1347.48 | 100.00% | replicate2 |
| OSD202246 | rice | 1,487,740 | 927 | 1604.90 | 99.68% | replicate2 |
| OSD202358 | rice | 1,276,554 | 929 | 1374.12 | 99.89% | replicate2 |
| OSD202412 | rice | 804,160 | 929 | 865.62 | 99.89% | replicate2 |
| OSD202440 | rice | 815,322 | 928 | 878.58 | 99.78% | replicate2 |
| OSD202557 | rice | 895,099 | 929 | 963.51 | 99.89% | replicate2 |
| OSD202663 | rice | 857,358 | 928 | 923.88 | 99.78% | replicate2 |
| OSD202674 | rice | 100,329 | 799 | 125.57 | 85.91% | replicate2 |
| OSD202680 | rice | 1,472,493 | 926 | 1590.17 | 99.57% | replicate2 |
| OSD202712 | rice | 1,222,849 | 929 | 1316.31 | 99.89% | replicate2 |
| OSD202723 | rice | 348,033 | 927 | 375.44 | 99.68% | replicate2 |
| OSD202740 | rice | 917,164 | 928 | 988.32 | 99.78% | replicate2 |
| OSD202745 | rice | 1,162,807 | 929 | 1251.68 | 99.89% | replicate2 |
| OSD202748 | rice | 1,063,203 | 928 | 1145.69 | 99.78% | replicate2 |
| OSD202751 | rice | 877,206 | 928 | 945.27 | 99.78% | replicate2 |
| OSD202763 | rice | 626,831 | 927 | 676.19 | 99.68% | replicate2 |
| OSD202766 | rice | 619,970 | 929 | 667.35 | 99.89% | replicate2 |
| OSD202767 | rice | 1,161,285 | 929 | 1250.04 | 99.89% | replicate2 |
| OSD202768 | rice | 875,463 | 930 | 941.36 | 100.00% | replicate2 |
| OSD202781 | rice | 1,109,736 | 927 | 1197.13 | 99.68% | replicate2 |
| OSD202792 | rice | 1,139,736 | 929 | 1226.84 | 99.89% | replicate2 |
| OSD202799 | rice | 1,170,908 | 929 | 1260.40 | 99.89% | replicate2 |
| OSD202801 | rice | 1,219,861 | 930 | 1311.68 | 100.00% | replicate2 |
| OSD202802 | rice | 1,217,929 | 930 | 1309.60 | 100.00% | replicate2 |
| OSD202809 | rice | 965,136 | 929 | 1038.90 | 99.89% | replicate2 |
| OSD202811 | rice | 976,844 | 929 | 1051.50 | 99.89% | replicate2 |
| OSD202815 | rice | 1,021,648 | 930 | 1098.55 | 100.00% | replicate2 |
| OSD202904 | rice | 618,565 | 930 | 665.12 | 100.00% | replicate2 |

| | | | | | | |
|---|---|---|---|---|---|---|
| OSD202906 | rice | 681,119 | 928 | 733.96 | 99.78% | replicate2 |
| OSD202912 | rice | 871,150 | 929 | 937.73 | 99.89% | replicate2 |
| OSD202914 | rice | 1,102,886 | 930 | 1185.90 | 100.00% | replicate2 |
| OSD202936 | rice | 1,036,704 | 929 | 1115.94 | 99.89% | replicate2 |
| OSD202938 | rice | 1,182,972 | 930 | 1272.01 | 100.00% | replicate2 |
| OSD202958 | rice | 701,520 | 929 | 755.13 | 99.89% | replicate2 |
| OSD203003 | rice | 812,656 | 929 | 874.76 | 99.89% | replicate2 |
| OSD203044 | rice | 1,236,385 | 930 | 1329.45 | 100.00% | replicate2 |
| OSD203080 | rice | 918,517 | 928 | 989.78 | 99.78% | replicate2 |
| OSD203081 | rice | 798,776 | 928 | 860.75 | 99.78% | replicate2 |
| OSD203095 | rice | 874,857 | 929 | 941.72 | 99.89% | replicate2 |
| OSD202062 | rice | 770,403 | 928 | 830.18 | 99.78% | replicate3 |
| OSD202175 | rice | 656,593 | 927 | 708.30 | 99.68% | replicate3 |
| OSD202180 | rice | 309,659 | 914 | 338.80 | 98.28% | replicate3 |
| OSD202183 | rice | 861,079 | 929 | 926.89 | 99.89% | replicate3 |
| OSD202200 | rice | 683,009 | 928 | 736.00 | 99.78% | replicate3 |
| OSD202201 | rice | 825,293 | 929 | 888.37 | 99.89% | replicate3 |
| OSD202227 | rice | 853,829 | 929 | 919.08 | 99.89% | replicate3 |
| OSD202237 | rice | 1,059,006 | 930 | 1138.72 | 100.00% | replicate3 |
| OSD202243 | rice | 1,160,878 | 930 | 1248.26 | 100.00% | replicate3 |
| OSD202246 | rice | 917,180 | 927 | 989.41 | 99.68% | replicate3 |
| OSD202358 | rice | 1,579,234 | 929 | 1699.93 | 99.89% | replicate3 |
| OSD202412 | rice | 1,159,900 | 929 | 1248.55 | 99.89% | replicate3 |
| OSD202440 | rice | 1,116,630 | 929 | 1201.97 | 99.89% | replicate3 |
| OSD202557 | rice | 1,014,763 | 929 | 1092.32 | 99.89% | replicate3 |
| OSD202663 | rice | 880,536 | 927 | 949.88 | 99.68% | replicate3 |
| OSD202674 | rice | 1,107,602 | 930 | 1190.97 | 100.00% | replicate3 |
| OSD202680 | rice | 1,250,425 | 926 | 1350.35 | 99.57% | replicate3 |
| OSD202712 | rice | 1,155,524 | 929 | 1243.84 | 99.89% | replicate3 |
| OSD202723 | rice | 945,385 | 929 | 1017.64 | 99.89% | replicate3 |
| OSD202740 | rice | 1,276,198 | 928 | 1375.21 | 99.78% | replicate3 |
| OSD202745 | rice | 775,920 | 928 | 836.12 | 99.78% | replicate3 |
| OSD202748 | rice | 1,144,210 | 928 | 1232.98 | 99.78% | replicate3 |
| OSD202751 | rice | 1,127,893 | 929 | 1214.09 | 99.89% | replicate3 |
| OSD202763 | rice | 865,300 | 927 | 933.44 | 99.68% | replicate3 |
| OSD202766 | rice | 679,298 | 929 | 731.21 | 99.89% | replicate3 |
| OSD202767 | rice | 1,160,658 | 929 | 1249.36 | 99.89% | replicate3 |
| OSD202768 | rice | 937,814 | 929 | 1009.49 | 99.89% | replicate3 |
| OSD202781 | rice | 1,371,857 | 927 | 1479.89 | 99.68% | replicate3 |
| OSD202792 | rice | 1,515,379 | 928 | 1632.95 | 99.78% | replicate3 |
| OSD202799 | rice | 1,484,599 | 929 | 1598.06 | 99.89% | replicate3 |
| OSD202801 | rice | 1,305,290 | 930 | 1403.54 | 100.00% | replicate3 |
| OSD202802 | rice | 1,371,409 | 929 | 1476.22 | 99.89% | replicate3 |
| OSD202809 | rice | 916,260 | 930 | 985.23 | 100.00% | replicate3 |
| OSD202811 | rice | 726,056 | 929 | 781.55 | 99.89% | replicate3 |
| OSD202815 | rice | 1,174,481 | 930 | 1262.88 | 100.00% | replicate3 |
| OSD202904 | rice | 1,051,356 | 930 | 1130.49 | 100.00% | replicate3 |
| OSD202906 | rice | 636,951 | 926 | 687.85 | 99.57% | replicate3 |
| OSD202912 | rice | 972,959 | 929 | 1047.32 | 99.89% | replicate3 |
| OSD202914 | rice | 640,558 | 929 | 689.51 | 99.89% | replicate3 |
| OSD202936 | rice | 1,136,067 | 929 | 1222.89 | 99.89% | replicate3 |
| OSD202938 | rice | 1,199,552 | 930 | 1289.84 | 100.00% | replicate3 |
| OSD202958 | rice | 1,148,646 | 929 | 1236.43 | 99.89% | replicate3 |
| OSD203003 | rice | 1,310,683 | 929 | 1410.85 | 99.89% | replicate3 |
| OSD203044 | rice | 832,068 | 927 | 897.59 | 99.68% | replicate3 |
| OSD203080 | rice | 1,257,173 | 929 | 1353.25 | 99.89% | replicate3 |
| OSD203081 | rice | 918,203 | 928 | 989.44 | 99.78% | replicate3 |
| OSD203095 | rice | 120,199 | 821 | 146.41 | 88.28% | replicate3 |
| 200383 | rice | 1,046,809 | 926 | 1130.46 | 99.57% | |
| 200384 | rice | 667,157 | 923 | 722.81 | 99.25% | |
| 200385 | rice | 932,718 | 926 | 1007.25 | 99.57% | |
| 200386 | rice | 987,321 | 927 | 1065.07 | 99.68% | |
| 200387 | rice | 883,882 | 926 | 954.52 | 99.57% | |
| 200388 | rice | 980,980 | 923 | 1062.82 | 99.25% | |
| 200389 | rice | 1,025,523 | 927 | 1106.28 | 99.68% | |
| 200390 | rice | 1,083,022 | 926 | 1169.57 | 99.57% | |
| 200391 | rice | 811,493 | 927 | 875.40 | 99.68% | |
| 200392 | rice | 1,009,613 | 926 | 1090.29 | 99.57% | |
| 200393 | rice | 916,326 | 924 | 991.69 | 99.35% | |
| 200394 | rice | 838,230 | 925 | 906.19 | 99.46% | |
| 200395 | rice | 716,715 | 927 | 773.16 | 99.68% | |
| 200396 | rice | 786,381 | 927 | 848.31 | 99.68% | |

| | | | | | |
|---|---|---|---|---|---|
| 200398 | rice | 1,043,660 | 926 | 1127.06 | 99.57% |
| 200399 | rice | 1,010,062 | 925 | 1091.96 | 99.46% |
| 200400 | rice | 875,685 | 923 | 948.74 | 99.25% |
| 200401 | rice | 947,389 | 926 | 1023.10 | 99.57% |
| 200402 | rice | 1,093,804 | 926 | 1181.21 | 99.57% |
| 200403 | rice | 926,613 | 924 | 1002.83 | 99.35% |
| 200404 | rice | 868,145 | 925 | 938.54 | 99.46% |
| 200405 | rice | 725,201 | 925 | 784.00 | 99.46% |
| 200406 | rice | 922,145 | 925 | 996.91 | 99.46% |
| 200407 | rice | 1,081,597 | 925 | 1169.29 | 99.46% |
| 200408 | rice | 907,492 | 927 | 978.96 | 99.68% |
| 201110 | rice | 1,112,284 | 926 | 1201.17 | 99.57% |
| 201111 | rice | 1,177,312 | 926 | 1271.40 | 99.57% |
| 201112 | rice | 1,254,762 | 926 | 1355.03 | 99.57% |
| 201113 | rice | 1,075,240 | 927 | 1159.91 | 99.68% |
| 201114 | rice | 1,132,768 | 927 | 1221.97 | 99.68% |
| market1 | rice | 640,621 | 891 | 718.99 | 95.70% |
| market12 | rice | 506,385 | 887 | 570.90 | 95.27% |
| market13 | rice | 140,216 | 658 | 213.09 | 70.68% |
| market14 | rice | 657,392 | 922 | 713.01 | 99.03% |
| market15 | rice | 532,212 | 919 | 579.12 | 98.71% |
| market16 | rice | 255,754 | 891 | 287.04 | 95.70% |
| market17 | rice | 364,856 | 906 | 402.71 | 97.31% |
| market18 | rice | 598,688 | 914 | 655.02 | 98.17% |
| market19 | rice | 216,322 | 816 | 265.10 | 87.65% |
| market2 | rice | 362,341 | 886 | 408.96 | 95.17% |
| market20 | rice | 228,740 | 810 | 282.40 | 87.00% |
| market21 | rice | 356,100 | 866 | 411.20 | 93.02% |
| market22 | rice | 321,762 | 831 | 387.20 | 89.26% |
| market23 | rice | 970,794 | 909 | 1067.98 | 97.64% |
| market24 | rice | 143,029 | 772 | 185.27 | 82.92% |
| market25 | rice | 769,990 | 907 | 848.94 | 97.42% |
| market26 | rice | 358,013 | 904 | 396.03 | 97.10% |
| market27 | rice | 404,414 | 902 | 448.35 | 96.89% |
| market28 | rice | 307,522 | 832 | 369.62 | 89.37% |
| market29 | rice | 608,615 | 909 | 669.54 | 97.64% |
| market3 | rice | 562,330 | 917 | 613.23 | 98.50% |
| market4 | rice | 180,592 | 845 | 213.72 | 90.76% |
| market5 | rice | 186,970 | 868 | 215.40 | 93.23% |
| market6 | rice | 1,231,005 | 923 | 1333.70 | 99.14% |
| market7 | rice | 306,904 | 905 | 339.12 | 97.21% |
| market8 | rice | 207,412 | 879 | 235.96 | 94.41% |
| market9 | rice | 222,159 | 842 | 263.85 | 90.44% |
| OSD10524 | rice | 289,685 | 865 | 334.90 | 92.91% |
| OSD1089 | rice | 124,097 | 781 | 158.90 | 83.89% |
| OSD1116 | rice | 1,096,002 | 918 | 1193.90 | 98.60% |
| OSD112 | rice | 123,817 | 765 | 161.85 | 82.17% |
| OSD1140 | rice | 651,509 | 919 | 708.93 | 98.71% |
| OSD1230 | rice | 209,425 | 807 | 259.51 | 86.68% |
| OSD1343 | rice | 227,564 | 842 | 270.27 | 90.44% |
| OSD1371 | rice | 303,610 | 906 | 335.11 | 97.31% |
| OSD1378 | rice | 378,180 | 869 | 435.19 | 93.34% |
| OSD1408 | rice | 518,105 | 906 | 571.86 | 97.31% |
| OSD1412 | rice | 296,454 | 874 | 339.19 | 93.88% |
| OSD1545 | rice | 272,780 | 880 | 309.98 | 94.52% |
| OSD1571 | rice | 137,314 | 797 | 172.29 | 85.61% |
| OSD1574 | rice | 280,283 | 859 | 326.29 | 92.27% |
| OSD1578 | rice | 279,516 | 878 | 318.36 | 94.31% |
| OSD1592 | rice | 341,014 | 877 | 388.84 | 94.20% |
| OSD1615 | rice | 435,236 | 912 | 477.23 | 97.96% |
| OSD1738 | rice | 379,974 | 886 | 428.86 | 95.17% |
| OSD1756 | rice | 552,189 | 927 | 595.67 | 99.57% |
| OSD1823 | rice | 317,149 | 905 | 350.44 | 97.21% |
| OSD1841 | rice | 145,106 | 803 | 180.70 | 86.25% |
| OSD1875 | rice | 332,381 | 893 | 372.21 | 95.92% |
| OSD1876 | rice | 198,024 | 882 | 224.52 | 94.74% |
| OSD1893 | rice | 188,824 | 844 | 223.73 | 90.66% |
| OSD1901 | rice | 317,283 | 890 | 356.50 | 95.60% |
| OSD1959 | rice | 130,316 | 744 | 175.16 | 79.91% |
| OSD1995 | rice | 462,991 | 912 | 507.67 | 97.96% |
| OSD200322 | rice | 666,195 | 913 | 729.68 | 98.07% |
| OSD2030 | rice | 283,451 | 882 | 321.37 | 94.74% |

| | | | | | |
|---|---|---|---|---|---|
| OSD2036 | rice | 313,870 | 877 | 357.89 | 94.20% |
| OSD2107 | rice | 177,845 | 831 | 214.01 | 89.26% |
| OSD2237 | rice | 227,552 | 835 | 272.52 | 89.69% |
| OSD2253 | rice | 596,101 | 915 | 651.48 | 98.28% |
| OSD2260 | rice | 238,160 | 901 | 264.33 | 96.78% |
| OSD2293 | rice | 606,506 | 915 | 662.85 | 98.28% |
| OSD2307 | rice | 356,311 | 891 | 399.90 | 95.70% |
| OSD2406 | rice | 433,548 | 901 | 481.19 | 96.78% |
| OSD2407 | rice | 277,962 | 868 | 320.23 | 93.23% |
| OSD2477 | rice | 817,033 | 917 | 890.98 | 98.50% |
| OSD2484 | rice | 202,681 | 844 | 240.14 | 90.66% |
| OSD2488 | rice | 298,424 | 880 | 339.12 | 94.52% |
| OSD2812 | rice | 157,058 | 775 | 202.66 | 83.24% |
| OSD2841 | rice | 115,483 | 753 | 153.36 | 80.88% |
| OSD2877 | rice | 221,529 | 842 | 263.10 | 90.44% |
| OSD30084 | rice | 338,229 | 880 | 384.35 | 94.52% |
| OSD30336 | rice | 314,521 | 897 | 350.64 | 96.35% |
| OSD3232 | rice | 236,252 | 837 | 282.26 | 89.90% |
| OSD3301 | rice | 376,147 | 894 | 420.75 | 96.03% |
| OSD3416 | rice | 599,370 | 886 | 676.49 | 95.17% |
| OSD3435 | rice | 259,627 | 832 | 312.05 | 89.37% |
| OSD3534 | rice | 515,536 | 916 | 562.81 | 98.39% |
| OSD3628 | rice | 251,477 | 873 | 288.06 | 93.77% |
| OSD3676 | rice | 459,964 | 908 | 506.57 | 97.53% |
| OSD3808 | rice | 266,092 | 806 | 330.14 | 86.57% |
| OSD3810 | rice | 155,128 | 771 | 201.20 | 82.81% |
| OSD3819 | rice | 620,808 | 905 | 685.98 | 97.21% |
| OSD3847 | rice | 340,958 | 895 | 380.96 | 96.13% |
| OSD3865 | rice | 378,875 | 899 | 421.44 | 96.56% |
| OSD4085 | rice | 136,745 | 797 | 171.57 | 85.61% |
| OSD4248 | rice | 268,813 | 864 | 311.13 | 92.80% |
| OSD4249 | rice | 287,519 | 887 | 324.15 | 95.27% |
| OSD4253 | rice | 617,297 | 904 | 682.85 | 97.10% |
| OSD4281 | rice | 241,316 | 862 | 279.95 | 92.59% |
| OSD4339 | rice | 178,493 | 780 | 228.84 | 83.78% |
| OSD4341 | rice | 122,315 | 746 | 163.96 | 80.13% |
| OSD4366 | rice | 478,927 | 915 | 523.42 | 98.28% |
| OSD4369 | rice | 800,962 | 921 | 869.67 | 98.93% |
| OSD4371 | rice | 291,781 | 872 | 334.61 | 93.66% |
| OSD4384 | rice | 214,240 | 816 | 262.55 | 87.65% |
| OSD4386 | rice | 112,674 | 764 | 147.48 | 82.06% |
| OSD4416 | rice | 296,391 | 862 | 343.84 | 92.59% |
| OSD4435 | rice | 244,486 | 854 | 286.28 | 91.73% |
| OSD4501 | rice | 386,021 | 902 | 427.96 | 96.89% |
| OSD4507 | rice | 306,658 | 863 | 355.34 | 92.70% |
| OSD4519 | rice | 535,997 | 906 | 591.61 | 97.31% |
| OSD4559 | rice | 363,789 | 902 | 403.31 | 96.89% |
| OSD4560 | rice | 243,384 | 857 | 284.00 | 92.05% |
| OSD4564 | rice | 311,203 | 901 | 345.40 | 96.78% |
| OSD4628 | rice | 213,285 | 854 | 249.75 | 91.73% |
| OSD4728 | rice | 205,016 | 836 | 245.23 | 89.80% |
| OSD4738 | rice | 231,750 | 843 | 274.91 | 90.55% |
| OSD4740 | rice | 106,770 | 761 | 140.30 | 81.74% |
| OSD4756 | rice | 186,862 | 824 | 226.77 | 88.51% |
| OSD4760 | rice | 302,945 | 887 | 341.54 | 95.27% |
| OSD4843 | rice | 210,764 | 845 | 249.42 | 90.76% |
| OSD5028 | rice | 758,990 | 917 | 827.69 | 98.50% |
| OSD5059 | rice | 178,989 | 844 | 212.07 | 90.66% |
| OSD5070 | rice | 236,549 | 848 | 278.95 | 91.08% |
| OSD5117 | rice | 541,416 | 919 | 589.14 | 98.71% |
| OSD5242 | rice | 254,101 | 866 | 293.42 | 93.02% |
| OSD5267 | rice | 414,211 | 910 | 455.18 | 97.74% |
| OSD5446 | rice | 832,466 | 921 | 903.87 | 98.93% |
| OSD5460 | rice | 338,215 | 903 | 374.55 | 96.99% |
| OSD5749 | rice | 314,557 | 888 | 354.23 | 95.38% |
| OSD5791 | rice | 352,138 | 885 | 397.90 | 95.06% |
| OSD5804 | rice | 169,495 | 778 | 217.86 | 83.57% |
| OSD5846 | rice | 623,460 | 899 | 693.50 | 96.56% |
| OSD5919 | rice | 556,347 | 915 | 608.03 | 98.28% |
| OSD6046 | rice | 644,993 | 902 | 715.07 | 96.89% |
| OSD6048 | rice | 315,070 | 900 | 350.08 | 96.67% |
| OSD6229 | rice | 152,743 | 773 | 197.60 | 83.03% |

| | | | | | |
|---|---|---|---|---|---|
| OSD6239 | rice | 301,154 | 866 | 347.75 | 93.02% |
| OSD6314 | rice | 289,447 | 881 | 328.54 | 94.63% |
| OSD6383 | rice | 381,795 | 915 | 417.26 | 98.28% |
| OSD6385 | rice | 214,714 | 834 | 257.45 | 89.58% |
| OSD6387 | rice | 229,957 | 844 | 272.46 | 90.66% |
| OSD6392 | rice | 243,885 | 889 | 274.34 | 95.49% |
| OSD6394 | rice | 152,385 | 780 | 195.37 | 83.78% |
| OSD6402 | rice | 1,160,753 | 914 | 1269.97 | 98.17% |
| OSD6471 | rice | 367,777 | 899 | 409.10 | 96.56% |
| OSD6473 | rice | 411,579 | 881 | 467.17 | 94.63% |
| OSD6482 | rice | 267,130 | 873 | 305.99 | 93.77% |
| OSD6489 | rice | 265,499 | 879 | 302.05 | 94.41% |
| OSD6512 | rice | 297,237 | 865 | 343.63 | 92.91% |
| OSD6662 | rice | 154,708 | 769 | 201.18 | 82.60% |
| OSD6826 | rice | 331,648 | 907 | 365.65 | 97.42% |
| OSD685 | rice | 322,658 | 879 | 367.07 | 94.41% |
| OSD6869 | rice | 330,359 | 881 | 374.98 | 94.63% |
| OSD7294 | rice | 135,689 | 749 | 181.16 | 80.45% |
| OSD7363 | rice | 319,162 | 875 | 364.76 | 93.98% |
| OSD7365 | rice | 404,859 | 898 | 450.85 | 96.46% |
| OSD7372 | rice | 638,531 | 894 | 714.24 | 96.03% |
| OSD7377 | rice | 197,693 | 841 | 235.07 | 90.33% |
| OSD7410 | rice | 143,052 | 755 | 189.47 | 81.10% |
| OSD7532 | rice | 356,586 | 894 | 398.87 | 96.03% |
| OSD7612 | rice | 179,101 | 761 | 235.35 | 81.74% |
| OSD7619 | rice | 204,648 | 827 | 247.46 | 88.83% |
| OSD7650 | rice | 390,332 | 893 | 437.10 | 95.92% |
| OSD7658 | rice | 164,516 | 809 | 203.36 | 86.90% |
| OSD7662 | rice | 317,048 | 858 | 369.52 | 92.16% |
| OSD7674 | rice | 243,571 | 849 | 286.89 | 91.19% |
| OSD7782 | rice | 105,779 | 745 | 141.99 | 80.02% |
| OSD7806 | rice | 218,792 | 831 | 263.29 | 89.26% |
| OSD7843 | rice | 196,999 | 859 | 229.34 | 92.27% |
| OSD7865 | rice | 180,679 | 768 | 235.26 | 82.49% |
| OSD8059 | rice | 304,876 | 902 | 338.00 | 96.89% |
| OSD8148 | rice | 119,813 | 769 | 155.80 | 82.60% |
| OSD8202 | rice | 361,223 | 889 | 406.33 | 95.49% |
| OSD8205 | rice | 508,554 | 924 | 550.38 | 99.25% |
| OSD8207 | rice | 472,192 | 916 | 515.49 | 98.39% |
| OSD8218 | rice | 416,103 | 918 | 453.27 | 98.60% |
| OSD8241 | rice | 775,052 | 917 | 845.20 | 98.50% |
| OSD8250 | rice | 602,245 | 913 | 659.63 | 98.07% |
| OSD8316 | rice | 487,132 | 901 | 540.66 | 96.78% |
| OSD8371 | rice | 135,533 | 747 | 181.44 | 80.24% |
| OSD8600 | rice | 398,282 | 886 | 449.53 | 95.17% |
| OSD889 | rice | 275,877 | 893 | 308.93 | 95.92% |
| OSD890 | rice | 371,615 | 887 | 418.96 | 95.27% |
| OSD9010 | rice | 264,260 | 847 | 312.00 | 90.98% |
| OSD9013 | rice | 187,496 | 836 | 224.28 | 89.80% |

*Only the samples used for estimation of MNP-seq reproducibility have replicates.
