## Supplementary table 3 for "Multiple nucleotide polymorphism DNA markers for the accurate evaluation of genetic variations"

**Supplementary Table 3. The allele number for each of SSR and MNP loci**

| Locus code | Alleles number | Marker type |
|---|---|---|
| AMPL1563062 | 4 | MNP |
| AMPL1563095 | 3 | MNP |
| AMPL1563096 | 6 | MNP |
| AMPL1563097 | 3 | MNP |
| AMPL1563100 | 4 | MNP |
| AMPL1563102 | 3 | MNP |
| AMPL1563103 | 4 | MNP |
| AMPL1563116 | 5 | MNP |
| AMPL1563121 | 5 | MNP |
| AMPL1563123 | 2 | MNP |
| AMPL1563126 | 4 | MNP |
| AMPL1563133 | 2 | MNP |
| AMPL1563134 | 3 | MNP |
| AMPL1563137 | 3 | MNP |
| AMPL1563138 | 5 | MNP |
| AMPL1563143 | 5 | MNP |
| AMPL1563147 | 5 | MNP |
| AMPL1563153 | 2 | MNP |
| AMPL1563154 | 4 | MNP |
| AMPL1563157 | 2 | MNP |
| AMPL1563158 | 5 | MNP |
| AMPL1563159 | 3 | MNP |
| AMPL1563161 | 4 | MNP |
| AMPL1563166 | 4 | MNP |
| AMPL1563168 | 2 | MNP |
| AMPL1563169 | 2 | MNP |
| AMPL1563172 | 4 | MNP |
| AMPL1563173 | 3 | MNP |
| AMPL1563175 | 3 | MNP |
| AMPL1563176 | 4 | MNP |
| AMPL1563178 | 4 | MNP |
| AMPL1563182 | 5 | MNP |
| AMPL1563184 | 3 | MNP |
| AMPL1563186 | 2 | MNP |
| AMPL1563187 | 3 | MNP |
| AMPL1563189 | 3 | MNP |
| AMPL1563193 | 2 | MNP |
| AMPL1563194 | 2 | MNP |
| AMPL1563196 | 2 | MNP |
| AMPL1563200 | 3 | MNP |
| AMPL1563203 | 5 | MNP |
| AMPL1563205 | 3 | MNP |
| AMPL1563212 | 6 | MNP |
| AMPL1563213 | 7 | MNP |
| AMPL1563214 | 4 | MNP |

| | | |
|---|---|---|
| AMPL1563215 | 6 | MNP |
| AMPL1563218 | 3 | MNP |
| AMPL1563222 | 3 | MNP |
| AMPL1563223 | 5 | MNP |
| AMPL1563234 | 2 | MNP |
| AMPL1563238 | 3 | MNP |
| AMPL1563244 | 4 | MNP |
| AMPL1563256 | 3 | MNP |
| AMPL1563261 | 5 | MNP |
| AMPL1563263 | 3 | MNP |
| AMPL1563267 | 2 | MNP |
| AMPL1563268 | 3 | MNP |
| AMPL1563272 | 3 | MNP |
| AMPL1563275 | 6 | MNP |
| AMPL1563276 | 5 | MNP |
| AMPL1563279 | 5 | MNP |
| AMPL1563282 | 5 | MNP |
| AMPL1563285 | 6 | MNP |
| AMPL1563286 | 4 | MNP |
| AMPL1563289 | 5 | MNP |
| AMPL1563296 | 3 | MNP |
| AMPL1563298 | 2 | MNP |
| AMPL1563299 | 3 | MNP |
| AMPL1563304 | 2 | MNP |
| AMPL1563308 | 3 | MNP |
| AMPL1563310 | 5 | MNP |
| AMPL1563311 | 5 | MNP |
| AMPL1563315 | 2 | MNP |
| AMPL1563317 | 3 | MNP |
| AMPL1563318 | 5 | MNP |
| AMPL1563326 | 3 | MNP |
| AMPL1563332 | 3 | MNP |
| AMPL1563333 | 5 | MNP |
| AMPL1563334 | 3 | MNP |
| AMPL1563337 | 6 | MNP |
| AMPL1563345 | 4 | MNP |
| AMPL1563353 | 4 | MNP |
| AMPL1563357 | 3 | MNP |
| AMPL1563358 | 4 | MNP |
| AMPL1563362 | 2 | MNP |
| AMPL1563365 | 3 | MNP |
| AMPL1563366 | 2 | MNP |
| AMPL1563369 | 6 | MNP |
| AMPL1563373 | 2 | MNP |
| AMPL1563375 | 4 | MNP |
| AMPL1563376 | 2 | MNP |
| AMPL1563377 | 4 | MNP |
| AMPL1563381 | 4 | MNP |

| | | |
|---|---|---|
| AMPL1563385 | 5 | MNP |
| AMPL1563387 | 4 | MNP |
| AMPL1563389 | 6 | MNP |
| AMPL1563390 | 4 | MNP |
| AMPL1563392 | 5 | MNP |
| AMPL1563396 | 3 | MNP |
| AMPL1563397 | 3 | MNP |
| AMPL1563398 | 4 | MNP |
| AMPL1563399 | 5 | MNP |
| AMPL1563409 | 3 | MNP |
| AMPL1563410 | 4 | MNP |
| AMPL1563412 | 6 | MNP |
| AMPL1563418 | 6 | MNP |
| AMPL1563419 | 3 | MNP |
| AMPL1563422 | 4 | MNP |
| AMPL1563424 | 3 | MNP |
| AMPL1563425 | 4 | MNP |
| AMPL1563427 | 3 | MNP |
| AMPL1563429 | 4 | MNP |
| AMPL1563435 | 2 | MNP |
| AMPL1563436 | 4 | MNP |
| AMPL1563437 | 6 | MNP |
| AMPL1563438 | 3 | MNP |
| AMPL1563443 | 3 | MNP |
| AMPL1563445 | 2 | MNP |
| AMPL1563447 | 5 | MNP |
| AMPL1563449 | 4 | MNP |
| AMPL1563451 | 4 | MNP |
| AMPL1563454 | 4 | MNP |
| AMPL1563456 | 3 | MNP |
| AMPL1563457 | 4 | MNP |
| AMPL1563460 | 4 | MNP |
| AMPL1563462 | 4 | MNP |
| AMPL1563464 | 3 | MNP |
| AMPL1563469 | 3 | MNP |
| AMPL1563472 | 5 | MNP |
| AMPL1563474 | 3 | MNP |
| AMPL1563482 | 4 | MNP |
| AMPL1563488 | 4 | MNP |
| AMPL1563489 | 5 | MNP |
| AMPL1563491 | 5 | MNP |
| AMPL1563493 | 2 | MNP |
| AMPL1563495 | 2 | MNP |
| AMPL1563497 | 4 | MNP |
| AMPL1563501 | 2 | MNP |
| AMPL1563509 | 5 | MNP |
| AMPL1563511 | 5 | MNP |
| AMPL1563512 | 5 | MNP |

| | | |
|---|---|---|
| AMPL1563513 | 7 | MNP |
| AMPL1563520 | 4 | MNP |
| AMPL1563524 | 5 | MNP |
| AMPL1563525 | 3 | MNP |
| AMPL1563529 | 5 | MNP |
| AMPL1563533 | 3 | MNP |
| AMPL1563534 | 3 | MNP |
| AMPL1563535 | 3 | MNP |
| AMPL1563536 | 2 | MNP |
| AMPL1563539 | 2 | MNP |
| AMPL1563543 | 2 | MNP |
| AMPL1563545 | 3 | MNP |
| AMPL1563548 | 3 | MNP |
| AMPL1563549 | 2 | MNP |
| AMPL1563550 | 3 | MNP |
| AMPL1563552 | 5 | MNP |
| AMPL1563555 | 3 | MNP |
| AMPL1563557 | 3 | MNP |
| AMPL1563559 | 2 | MNP |
| AMPL1563561 | 2 | MNP |
| AMPL1563563 | 3 | MNP |
| AMPL1563567 | 2 | MNP |
| AMPL1563568 | 4 | MNP |
| AMPL1563569 | 3 | MNP |
| AMPL1563576 | 3 | MNP |
| AMPL1563577 | 5 | MNP |
| AMPL1563587 | 3 | MNP |
| AMPL1563589 | 2 | MNP |
| AMPL1563591 | 5 | MNP |
| AMPL1563595 | 3 | MNP |
| AMPL1563598 | 4 | MNP |
| AMPL1563603 | 4 | MNP |
| AMPL1563605 | 3 | MNP |
| AMPL1563608 | 4 | MNP |
| AMPL1563609 | 4 | MNP |
| AMPL1563611 | 3 | MNP |
| AMPL1563614 | 3 | MNP |
| AMPL1563616 | 5 | MNP |
| AMPL1563620 | 4 | MNP |
| AMPL1563623 | 3 | MNP |
| AMPL1563626 | 2 | MNP |
| AMPL1563633 | 2 | MNP |
| AMPL1563635 | 3 | MNP |
| AMPL1563636 | 3 | MNP |
| AMPL1563637 | 4 | MNP |
| AMPL1563639 | 3 | MNP |
| AMPL1563641 | 2 | MNP |
| AMPL1563643 | 4 | MNP |

| | | |
|---|---|---|
| AMPL1563648 | 3 | MNP |
| AMPL1563651 | 3 | MNP |
| AMPL1563653 | 2 | MNP |
| AMPL1563656 | 2 | MNP |
| AMPL1563657 | 3 | MNP |
| AMPL1563658 | 3 | MNP |
| AMPL1563663 | 3 | MNP |
| AMPL1563664 | 3 | MNP |
| AMPL1563669 | 4 | MNP |
| AMPL1563670 | 5 | MNP |
| AMPL1563672 | 3 | MNP |
| AMPL1563673 | 5 | MNP |
| AMPL1563675 | 3 | MNP |
| AMPL1563684 | 2 | MNP |
| AMPL1563685 | 3 | MNP |
| AMPL1563686 | 5 | MNP |
| AMPL1563687 | 3 | MNP |
| AMPL1563689 | 3 | MNP |
| AMPL1563690 | 2 | MNP |
| AMPL1563695 | 2 | MNP |
| AMPL1563698 | 3 | MNP |
| AMPL1563701 | 3 | MNP |
| AMPL1563704 | 4 | MNP |
| AMPL1563705 | 3 | MNP |
| AMPL1563712 | 2 | MNP |
| AMPL1563713 | 4 | MNP |
| AMPL1563716 | 3 | MNP |
| AMPL1563718 | 4 | MNP |
| AMPL1563721 | 3 | MNP |
| AMPL1563722 | 3 | MNP |
| AMPL1563723 | 4 | MNP |
| AMPL1563725 | 4 | MNP |
| AMPL1563728 | 3 | MNP |
| AMPL1563730 | 3 | MNP |
| AMPL1563733 | 3 | MNP |
| AMPL1563737 | 2 | MNP |
| AMPL1563738 | 5 | MNP |
| AMPL1563741 | 2 | MNP |
| AMPL1563742 | 2 | MNP |
| AMPL1563745 | 2 | MNP |
| AMPL1563748 | 5 | MNP |
| AMPL1563751 | 3 | MNP |
| AMPL1563752 | 2 | MNP |
| AMPL1563755 | 2 | MNP |
| AMPL1563756 | 3 | MNP |
| AMPL1563757 | 5 | MNP |
| AMPL1563759 | 3 | MNP |
| AMPL1563764 | 2 | MNP |

| | | |
|---|---|---|
| AMPL1563767 | 4 | MNP |
| AMPL1563773 | 4 | MNP |
| AMPL1563776 | 3 | MNP |
| AMPL1563779 | 5 | MNP |
| AMPL1563785 | 3 | MNP |
| AMPL1563788 | 5 | MNP |
| AMPL1563790 | 4 | MNP |
| AMPL1563792 | 3 | MNP |
| AMPL1563797 | 2 | MNP |
| AMPL1563798 | 3 | MNP |
| AMPL1563800 | 3 | MNP |
| AMPL1563805 | 4 | MNP |
| AMPL1563808 | 3 | MNP |
| AMPL1563811 | 7 | MNP |
| AMPL1563814 | 5 | MNP |
| AMPL1563817 | 4 | MNP |
| AMPL1563818 | 4 | MNP |
| AMPL1563819 | 3 | MNP |
| AMPL1563831 | 3 | MNP |
| AMPL1563832 | 2 | MNP |
| AMPL1563833 | 3 | MNP |
| AMPL1563835 | 3 | MNP |
| AMPL1563843 | 3 | MNP |
| AMPL1563845 | 6 | MNP |
| AMPL1563851 | 3 | MNP |
| AMPL1563855 | 5 | MNP |
| AMPL1563865 | 3 | MNP |
| AMPL1563870 | 3 | MNP |
| AMPL1563871 | 2 | MNP |
| AMPL1563874 | 4 | MNP |
| AMPL1563878 | 3 | MNP |
| AMPL1563882 | 4 | MNP |
| AMPL1563886 | 2 | MNP |
| AMPL1563888 | 4 | MNP |
| AMPL1563889 | 5 | MNP |
| AMPL1563894 | 3 | MNP |
| AMPL1563895 | 8 | MNP |
| AMPL1563898 | 6 | MNP |
| AMPL1563899 | 4 | MNP |
| AMPL1563900 | 4 | MNP |
| AMPL1563902 | 4 | MNP |
| AMPL1563907 | 3 | MNP |
| AMPL1563909 | 3 | MNP |
| AMPL1563910 | 5 | MNP |
| AMPL2000000 | 3 | MNP |
| AMPL2000001 | 3 | MNP |
| AMPL2000002 | 3 | MNP |
| AMPL2000003 | 5 | MNP |

| | | |
|---|---|---|
| AMPL2000004 | 6 | MNP |
| AMPL2000005 | 4 | MNP |
| AMPL2000006 | 5 | MNP |
| AMPL2000007 | 4 | MNP |
| AMPL2000008 | 3 | MNP |
| AMPL2000009 | 4 | MNP |
| AMPL2000010 | 3 | MNP |
| AMPL2000011 | 5 | MNP |
| AMPL2000012 | 2 | MNP |
| AMPL2000013 | 3 | MNP |
| AMPL2000014 | 3 | MNP |
| AMPL2000015 | 3 | MNP |
| AMPL2000016 | 2 | MNP |
| AMPL2000017 | 3 | MNP |
| AMPL2000018 | 4 | MNP |
| AMPL2000019 | 2 | MNP |
| AMPL2000020 | 3 | MNP |
| AMPL2000021 | 4 | MNP |
| AMPL2000022 | 2 | MNP |
| AMPL2000023 | 3 | MNP |
| AMPL2000024 | 3 | MNP |
| AMPL2000026 | 4 | MNP |
| AMPL2000027 | 3 | MNP |
| AMPL2000028 | 3 | MNP |
| AMPL2000029 | 3 | MNP |
| AMPL2000030 | 3 | MNP |
| AMPL2000031 | 4 | MNP |
| AMPL2000032 | 3 | MNP |
| AMPL2000033 | 3 | MNP |
| AMPL2000035 | 3 | MNP |
| AMPL2000036 | 4 | MNP |
| AMPL2000037 | 6 | MNP |
| AMPL2000038 | 3 | MNP |
| AMPL2000039 | 3 | MNP |
| AMPL2000040 | 3 | MNP |
| AMPL2000041 | 3 | MNP |
| AMPL2000042 | 3 | MNP |
| AMPL2000043 | 3 | MNP |
| AMPL2000044 | 4 | MNP |
| AMPL2000045 | 3 | MNP |
| AMPL2000046 | 5 | MNP |
| AMPL2000047 | 4 | MNP |
| AMPL2000048 | 4 | MNP |
| AMPL2000049 | 3 | MNP |
| AMPL2000050 | 1 | MNP |
| AMPL2000051 | 3 | MNP |
| AMPL2000052 | 4 | MNP |
| AMPL2000053 | 3 | MNP |

| | | |
|---|---|---|
| AMPL2000054 | 2 | MNP |
| AMPL2000055 | 4 | MNP |
| AMPL2000056 | 3 | MNP |
| AMPL2000057 | 4 | MNP |
| AMPL2000058 | 3 | MNP |
| AMPL2000059 | 4 | MNP |
| AMPL2000061 | 3 | MNP |
| AMPL2000062 | 2 | MNP |
| AMPL2000063 | 2 | MNP |
| AMPL2000064 | 3 | MNP |
| AMPL2000065 | 2 | MNP |
| AMPL2000066 | 4 | MNP |
| AMPL2000067 | 3 | MNP |
| AMPL2000068 | 4 | MNP |
| AMPL2000069 | 2 | MNP |
| AMPL2000070 | 3 | MNP |
| AMPL2000071 | 2 | MNP |
| AMPL2000072 | 3 | MNP |
| AMPL2000073 | 4 | MNP |
| AMPL2000074 | 4 | MNP |
| AMPL2000075 | 6 | MNP |
| AMPL2000076 | 6 | MNP |
| AMPL2000077 | 2 | MNP |
| AMPL2000078 | 4 | MNP |
| AMPL2000079 | 4 | MNP |
| AMPL2000080 | 3 | MNP |
| AMPL2000081 | 6 | MNP |
| AMPL2000083 | 5 | MNP |
| AMPL2000085 | 2 | MNP |
| AMPL2000086 | 2 | MNP |
| AMPL2000087 | 3 | MNP |
| AMPL2000088 | 3 | MNP |
| AMPL2000089 | 5 | MNP |
| AMPL2000090 | 3 | MNP |
| AMPL2000091 | 3 | MNP |
| AMPL2000093 | 3 | MNP |
| AMPL2000094 | 3 | MNP |
| AMPL2000095 | 4 | MNP |
| AMPL2000096 | 3 | MNP |
| AMPL2000097 | 2 | MNP |
| AMPL2000098 | 6 | MNP |
| AMPL2000099 | 6 | MNP |
| AMPL2000100 | 4 | MNP |
| AMPL2000101 | 4 | MNP |
| AMPL2000103 | 3 | MNP |
| AMPL2000104 | 2 | MNP |
| AMPL2000105 | 3 | MNP |
| AMPL2000106 | 3 | MNP |

| | | |
|---|---|---|
| AMPL2000107 | 2 | MNP |
| AMPL2000108 | 4 | MNP |
| AMPL2000109 | 3 | MNP |
| AMPL2000110 | 3 | MNP |
| AMPL2000112 | 5 | MNP |
| AMPL2000113 | 5 | MNP |
| AMPL2000114 | 6 | MNP |
| AMPL2000115 | 3 | MNP |
| AMPL2000117 | 5 | MNP |
| AMPL2000118 | 2 | MNP |
| AMPL2000119 | 2 | MNP |
| AMPL2000120 | 3 | MNP |
| AMPL2000121 | 4 | MNP |
| AMPL2000122 | 3 | MNP |
| AMPL2000123 | 3 | MNP |
| AMPL2000124 | 3 | MNP |
| AMPL2000125 | 4 | MNP |
| AMPL2000126 | 3 | MNP |
| AMPL2000127 | 3 | MNP |
| AMPL2000128 | 3 | MNP |
| AMPL2000129 | 3 | MNP |
| AMPL2000130 | 3 | MNP |
| AMPL2000131 | 4 | MNP |
| AMPL2000132 | 3 | MNP |
| AMPL2000133 | 6 | MNP |
| AMPL2000134 | 3 | MNP |
| AMPL2000135 | 3 | MNP |
| AMPL2000136 | 7 | MNP |
| AMPL2000137 | 6 | MNP |
| AMPL2000138 | 5 | MNP |
| AMPL2000139 | 5 | MNP |
| AMPL2000141 | 3 | MNP |
| AMPL2000142 | 3 | MNP |
| AMPL2000143 | 6 | MNP |
| AMPL2000144 | 3 | MNP |
| AMPL2000145 | 3 | MNP |
| AMPL2000146 | 2 | MNP |
| AMPL2000147 | 6 | MNP |
| AMPL2000148 | 4 | MNP |
| AMPL2000149 | 3 | MNP |
| AMPL2000150 | 4 | MNP |
| AMPL2000151 | 7 | MNP |
| AMPL2000152 | 2 | MNP |
| AMPL2000153 | 3 | MNP |
| AMPL2000154 | 4 | MNP |
| AMPL2000155 | 3 | MNP |
| AMPL2000156 | 5 | MNP |
| AMPL2000157 | 3 | MNP |

| | | |
|---|---|---|
| AMPL2000158 | 4 | MNP |
| AMPL2000159 | 2 | MNP |
| AMPL2000160 | 4 | MNP |
| AMPL2000161 | 4 | MNP |
| AMPL2000162 | 3 | MNP |
| AMPL2000163 | 5 | MNP |
| AMPL2000165 | 4 | MNP |
| AMPL2000166 | 4 | MNP |
| AMPL2000168 | 3 | MNP |
| AMPL2000169 | 3 | MNP |
| AMPL2000170 | 2 | MNP |
| AMPL2000171 | 5 | MNP |
| AMPL2000172 | 2 | MNP |
| AMPL2000173 | 2 | MNP |
| AMPL2000174 | 2 | MNP |
| AMPL2000175 | 4 | MNP |
| AMPL2000176 | 3 | MNP |
| AMPL2000177 | 3 | MNP |
| AMPL2000178 | 3 | MNP |
| AMPL2000179 | 3 | MNP |
| AMPL2000180 | 4 | MNP |
| AMPL2000181 | 2 | MNP |
| AMPL2000182 | 3 | MNP |
| AMPL2000183 | 5 | MNP |
| AMPL2000184 | 9 | MNP |
| AMPL2000185 | 3 | MNP |
| AMPL2000186 | 3 | MNP |
| AMPL2000187 | 3 | MNP |
| AMPL2000188 | 2 | MNP |
| AMPL2000189 | 3 | MNP |
| AMPL2000190 | 2 | MNP |
| AMPL2000191 | 2 | MNP |
| AMPL2000192 | 3 | MNP |
| AMPL2000193 | 2 | MNP |
| AMPL2000194 | 3 | MNP |
| AMPL2000195 | 3 | MNP |
| AMPL2000196 | 4 | MNP |
| AMPL2000198 | 4 | MNP |
| AMPL2000199 | 3 | MNP |
| AMPL2000200 | 3 | MNP |
| AMPL2000201 | 2 | MNP |
| AMPL2000202 | 3 | MNP |
| AMPL2000203 | 3 | MNP |
| AMPL2000204 | 2 | MNP |
| AMPL2000205 | 3 | MNP |
| AMPL2000206 | 3 | MNP |
| AMPL2000207 | 3 | MNP |
| AMPL2000208 | 4 | MNP |

| | | |
|---|---|---|
| AMPL2000209 | 4 | MNP |
| AMPL2000210 | 4 | MNP |
| AMPL2000211 | 3 | MNP |
| AMPL2000212 | 4 | MNP |
| AMPL2000213 | 4 | MNP |
| AMPL2000214 | 4 | MNP |
| AMPL2000215 | 5 | MNP |
| AMPL2000216 | 2 | MNP |
| AMPL2000217 | 2 | MNP |
| AMPL2000218 | 3 | MNP |
| AMPL2000219 | 2 | MNP |
| AMPL2000220 | 3 | MNP |
| AMPL2000222 | 3 | MNP |
| AMPL2000223 | 3 | MNP |
| AMPL2000224 | 3 | MNP |
| AMPL2000225 | 3 | MNP |
| AMPL2000226 | 4 | MNP |
| AMPL2000228 | 4 | MNP |
| AMPL2000229 | 3 | MNP |
| AMPL2000230 | 4 | MNP |
| AMPL2000231 | 3 | MNP |
| AMPL2000232 | 3 | MNP |
| AMPL2000233 | 3 | MNP |
| AMPL2000234 | 4 | MNP |
| AMPL2000235 | 3 | MNP |
| AMPL2000236 | 3 | MNP |
| AMPL2000237 | 8 | MNP |
| AMPL2000238 | 3 | MNP |
| AMPL2000239 | 4 | MNP |
| AMPL2000240 | 3 | MNP |
| AMPL2000241 | 2 | MNP |
| AMPL2000242 | 5 | MNP |
| AMPL2000243 | 4 | MNP |
| AMPL2000244 | 2 | MNP |
| AMPL2000245 | 2 | MNP |
| AMPL2000246 | 3 | MNP |
| AMPL2000247 | 3 | MNP |
| AMPL2000248 | 3 | MNP |
| AMPL2000249 | 5 | MNP |
| AMPL2000250 | 2 | MNP |
| AMPL2000251 | 2 | MNP |
| AMPL2000252 | 3 | MNP |
| AMPL2000253 | 5 | MNP |
| AMPL2000254 | 4 | MNP |
| AMPL2000255 | 3 | MNP |
| AMPL2000256 | 3 | MNP |
| AMPL2000257 | 5 | MNP |
| AMPL2000258 | 2 | MNP |

| | | |
|---|---|---|
| AMPL2000259 | 4 | MNP |
| AMPL2000260 | 2 | MNP |
| AMPL2000261 | 3 | MNP |
| AMPL2000262 | 2 | MNP |
| AMPL2000263 | 3 | MNP |
| AMPL2000264 | 2 | MNP |
| AMPL2000265 | 2 | MNP |
| AMPL2000266 | 3 | MNP |
| AMPL2000267 | 2 | MNP |
| AMPL2000268 | 3 | MNP |
| AMPL2000269 | 3 | MNP |
| AMPL2000272 | 6 | MNP |
| AMPL2000273 | 3 | MNP |
| AMPL2000274 | 2 | MNP |
| AMPL2000275 | 3 | MNP |
| AMPL2000277 | 3 | MNP |
| AMPL2000278 | 3 | MNP |
| AMPL2000279 | 2 | MNP |
| AMPL2000280 | 2 | MNP |
| AMPL2000281 | 2 | MNP |
| AMPL2000282 | 2 | MNP |
| AMPL2000283 | 2 | MNP |
| AMPL2000284 | 3 | MNP |
| AMPL2000285 | 2 | MNP |
| AMPL2000286 | 3 | MNP |
| AMPL2000287 | 1 | MNP |
| AMPL2000288 | 4 | MNP |
| AMPL2000289 | 2 | MNP |
| AMPL2000290 | 3 | MNP |
| AMPL2000291 | 2 | MNP |
| AMPL2000292 | 5 | MNP |
| AMPL2000293 | 6 | MNP |
| AMPL2000294 | 4 | MNP |
| AMPL2000295 | 7 | MNP |
| AMPL2000296 | 5 | MNP |
| AMPL2000297 | 4 | MNP |
| AMPL2000298 | 3 | MNP |
| AMPL2000300 | 3 | MNP |
| AMPL2000302 | 2 | MNP |
| AMPL2000303 | 4 | MNP |
| AMPL2000304 | 3 | MNP |
| AMPL2000305 | 4 | MNP |
| AMPL2000306 | 8 | MNP |
| AMPL2000307 | 4 | MNP |
| AMPL2000308 | 3 | MNP |
| AMPL2000309 | 3 | MNP |
| AMPL2000311 | 3 | MNP |
| AMPL2000312 | 2 | MNP |

| | | |
|---|---|---|
| AMPL2000313 | 4 | MNP |
| AMPL2000314 | 2 | MNP |
| AMPL2000315 | 5 | MNP |
| AMPL2000316 | 3 | MNP |
| AMPL2000319 | 3 | MNP |
| AMPL2000320 | 3 | MNP |
| AMPL2000321 | 2 | MNP |
| AMPL2000323 | 3 | MNP |
| AMPL2000324 | 2 | MNP |
| AMPL2000325 | 4 | MNP |
| AMPL2000326 | 2 | MNP |
| AMPL2000327 | 4 | MNP |
| AMPL2000328 | 3 | MNP |
| AMPL2000329 | 2 | MNP |
| AMPL2000330 | 4 | MNP |
| AMPL2000331 | 3 | MNP |
| AMPL2000332 | 3 | MNP |
| AMPL2000333 | 5 | MNP |
| AMPL2000335 | 3 | MNP |
| AMPL2000336 | 2 | MNP |
| AMPL2000337 | 4 | MNP |
| AMPL2000338 | 4 | MNP |
| AMPL2000339 | 5 | MNP |
| AMPL2000340 | 3 | MNP |
| AMPL2000341 | 3 | MNP |
| AMPL2000342 | 2 | MNP |
| AMPL2000343 | 5 | MNP |
| AMPL2000344 | 3 | MNP |
| AMPL2000345 | 4 | MNP |
| AMPL2000346 | 3 | MNP |
| AMPL2000347 | 3 | MNP |
| AMPL2000349 | 3 | MNP |
| AMPL2000350 | 3 | MNP |
| AMPL2000351 | 3 | MNP |
| AMPL2000352 | 3 | MNP |
| AMPL2000353 | 2 | MNP |
| AMPL2000354 | 3 | MNP |
| AMPL2000355 | 2 | MNP |
| AMPL2000356 | 3 | MNP |
| AMPL2000357 | 3 | MNP |
| AMPL2000358 | 3 | MNP |
| AMPL2000359 | 4 | MNP |
| AMPL2000360 | 4 | MNP |
| AMPL2000362 | 4 | MNP |
| AMPL2000363 | 5 | MNP |
| AMPL2000365 | 2 | MNP |
| AMPL2000366 | 3 | MNP |
| AMPL2000367 | 6 | MNP |

| | | |
|---|---|---|
| AMPL2000368 | 4 | MNP |
| AMPL2000369 | 5 | MNP |
| AMPL2000370 | 2 | MNP |
| AMPL2000371 | 5 | MNP |
| AMPL2000372 | 4 | MNP |
| AMPL2000373 | 2 | MNP |
| AMPL2000374 | 4 | MNP |
| AMPL2000375 | 3 | MNP |
| AMPL2000376 | 2 | MNP |
| AMPL2000378 | 3 | MNP |
| AMPL2000379 | 4 | MNP |
| AMPL2000380 | 3 | MNP |
| AMPL2000381 | 3 | MNP |
| AMPL2000382 | 4 | MNP |
| AMPL2000383 | 2 | MNP |
| AMPL2000384 | 4 | MNP |
| AMPL2000385 | 2 | MNP |
| AMPL2000386 | 2 | MNP |
| AMPL2000388 | 3 | MNP |
| AMPL2000389 | 4 | MNP |
| AMPL2000390 | 4 | MNP |
| AMPL2000391 | 5 | MNP |
| AMPL2000392 | 3 | MNP |
| AMPL2000393 | 4 | MNP |
| AMPL2000394 | 5 | MNP |
| AMPL2000396 | 4 | MNP |
| AMPL2000399 | 3 | MNP |
| AMPL2000400 | 4 | MNP |
| AMPL2000401 | 4 | MNP |
| AMPL2000403 | 4 | MNP |
| AMPL2000404 | 3 | MNP |
| AMPL2000406 | 3 | MNP |
| AMPL2000407 | 5 | MNP |
| AMPL2000408 | 3 | MNP |
| AMPL2000409 | 4 | MNP |
| AMPL2000410 | 3 | MNP |
| AMPL2000411 | 4 | MNP |
| AMPL2000412 | 4 | MNP |
| AMPL2000414 | 3 | MNP |
| AMPL2000415 | 3 | MNP |
| AMPL2000416 | 3 | MNP |
| AMPL2000417 | 2 | MNP |
| AMPL2000420 | 2 | MNP |
| AMPL2000421 | 2 | MNP |
| AMPL2000422 | 3 | MNP |
| AMPL2000423 | 3 | MNP |
| AMPL2000425 | 2 | MNP |
| AMPL2000426 | 3 | MNP |

| | | |
|---|---|---|
| AMPL2000427 | 3 | MNP |
| AMPL2000428 | 3 | MNP |
| AMPL2000429 | 4 | MNP |
| AMPL2000430 | 3 | MNP |
| AMPL2000431 | 4 | MNP |
| AMPL2000433 | 2 | MNP |
| AMPL2000434 | 4 | MNP |
| AMPL2000435 | 4 | MNP |
| AMPL2000436 | 2 | MNP |
| AMPL2000437 | 4 | MNP |
| AMPL2000438 | 4 | MNP |
| AMPL2000439 | 3 | MNP |
| AMPL2000441 | 4 | MNP |
| AMPL2000442 | 6 | MNP |
| AMPL2000443 | 3 | MNP |
| AMPL2000444 | 3 | MNP |
| AMPL2000445 | 3 | MNP |
| AMPL2000447 | 3 | MNP |
| AMPL2000448 | 3 | MNP |
| AMPL2000450 | 4 | MNP |
| AMPL2000451 | 3 | MNP |
| AMPL2000452 | 4 | MNP |
| AMPL2000453 | 3 | MNP |
| AMPL2000454 | 3 | MNP |
| AMPL2000455 | 2 | MNP |
| AMPL2000456 | 4 | MNP |
| AMPL2000457 | 3 | MNP |
| AMPL2000458 | 3 | MNP |
| AMPL2000459 | 3 | MNP |
| AMPL2000460 | 4 | MNP |
| AMPL2000461 | 4 | MNP |
| AMPL2000462 | 3 | MNP |
| AMPL2000463 | 4 | MNP |
| AMPL2000464 | 2 | MNP |
| AMPL2000465 | 3 | MNP |
| AMPL2000466 | 4 | MNP |
| AMPL2000467 | 3 | MNP |
| AMPL2000468 | 2 | MNP |
| AMPL2000469 | 3 | MNP |
| AMPL2000470 | 3 | MNP |
| AMPL2000471 | 3 | MNP |
| AMPL2000472 | 3 | MNP |
| AMPL2000473 | 3 | MNP |
| AMPL2000475 | 6 | MNP |
| AMPL2000476 | 6 | MNP |
| AMPL2000477 | 5 | MNP |
| AMPL2000478 | 3 | MNP |
| AMPL2000479 | 3 | MNP |

| | | |
|---|---|---|
| AMPL2000480 | 3 | MNP |
| AMPL2000481 | 1 | MNP |
| AMPL2000483 | 4 | MNP |
| AMPL2000484 | 4 | MNP |
| AMPL2000485 | 3 | MNP |
| AMPL2000486 | 3 | MNP |
| AMPL2000487 | 3 | MNP |
| AMPL2000488 | 3 | MNP |
| AMPL2000489 | 5 | MNP |
| AMPL2000490 | 2 | MNP |
| AMPL2000491 | 4 | MNP |
| AMPL2000492 | 3 | MNP |
| AMPL2000493 | 3 | MNP |
| AMPL2000494 | 4 | MNP |
| AMPL2000495 | 3 | MNP |
| AMPL2000496 | 3 | MNP |
| AMPL2000497 | 3 | MNP |
| AMPL2000498 | 3 | MNP |
| AMPL2000499 | 3 | MNP |
| AMPL2000500 | 2 | MNP |
| AMPL2000501 | 5 | MNP |
| AMPL2000502 | 3 | MNP |
| AMPL2000503 | 2 | MNP |
| AMPL2000504 | 4 | MNP |
| AMPL2000505 | 4 | MNP |
| AMPL2000506 | 5 | MNP |
| AMPL2000508 | 4 | MNP |
| AMPL2000509 | 4 | MNP |
| AMPL2000510 | 1 | MNP |
| AMPL2000511 | 3 | MNP |
| AMPL2000513 | 3 | MNP |
| AMPL2000514 | 3 | MNP |
| AMPL2000515 | 3 | MNP |
| AMPL2000516 | 3 | MNP |
| AMPL2000517 | 1 | MNP |
| AMPL2000518 | 3 | MNP |
| AMPL2000519 | 3 | MNP |
| AMPL2000520 | 3 | MNP |
| AMPL2000521 | 4 | MNP |
| AMPL2000522 | 3 | MNP |
| AMPL2000523 | 3 | MNP |
| AMPL2000524 | 3 | MNP |
| AMPL2000525 | 2 | MNP |
| AMPL2000526 | 2 | MNP |
| AMPL2000527 | 3 | MNP |
| AMPL2000529 | 2 | MNP |
| AMPL2000530 | 2 | MNP |
| AMPL2000531 | 2 | MNP |

| | | |
|---|---|---|
| AMPL2000532 | 3 | MNP |
| AMPL2000533 | 3 | MNP |
| AMPL2000534 | 3 | MNP |
| AMPL2000535 | 3 | MNP |
| AMPL2000539 | 4 | MNP |
| AMPL2000540 | 1 | MNP |
| AMPL2000541 | 3 | MNP |
| AMPL2000542 | 4 | MNP |
| AMPL2000543 | 4 | MNP |
| AMPL2000544 | 3 | MNP |
| AMPL2000545 | 3 | MNP |
| AMPL2000546 | 3 | MNP |
| AMPL2000547 | 4 | MNP |
| AMPL2000550 | 1 | MNP |
| AMPL2000551 | 3 | MNP |
| AMPL2000552 | 4 | MNP |
| AMPL2000553 | 4 | MNP |
| AMPL2000554 | 5 | MNP |
| AMPL2000555 | 4 | MNP |
| AMPL2000556 | 4 | MNP |
| AMPL2000557 | 2 | MNP |
| AMPL2000558 | 2 | MNP |
| AMPL2000559 | 3 | MNP |
| AMPL2000560 | 3 | MNP |
| AMPL2000561 | 3 | MNP |
| AMPL2000562 | 3 | MNP |
| AMPL2000563 | 4 | MNP |
| AMPL2000564 | 4 | MNP |
| AMPL2000565 | 3 | MNP |
| AMPL2000566 | 6 | MNP |
| AMPL2000567 | 5 | MNP |
| AMPL2000568 | 4 | MNP |
| AMPL2000570 | 3 | MNP |
| AMPL2000571 | 2 | MNP |
| AMPL2000572 | 3 | MNP |
| AMPL2000573 | 3 | MNP |
| AMPL2000574 | 4 | MNP |
| AMPL2000576 | 3 | MNP |
| AMPL2000577 | 2 | MNP |
| AMPL2000578 | 2 | MNP |
| AMPL2000579 | 3 | MNP |
| AMPL2000580 | 4 | MNP |
| AMPL2000581 | 5 | MNP |
| AMPL2000582 | 2 | MNP |
| AMPL2000583 | 3 | MNP |
| AMPL2000584 | 2 | MNP |
| AMPL2000585 | 2 | MNP |
| AMPL2000587 | 4 | MNP |

| | | |
|---|---|---|
| AMPL2000588 | 4 | MNP |
| AMPL2000589 | 4 | MNP |
| AMPL2000590 | 4 | MNP |
| AMPL2000591 | 6 | MNP |
| AMPL2000593 | 3 | MNP |
| AMPL2000594 | 3 | MNP |
| AMPL2000596 | 3 | MNP |
| AMPL2000597 | 3 | MNP |
| AMPL2000598 | 2 | MNP |
| AMPL2000599 | 2 | MNP |
| AMPL2000600 | 3 | MNP |
| AMPL2000601 | 2 | MNP |
| AMPL2000602 | 2 | MNP |
| AMPL2000603 | 2 | MNP |
| AMPL2000604 | 3 | MNP |
| AMPL2000606 | 2 | MNP |
| AMPL2000607 | 2 | MNP |
| AMPL2000608 | 4 | MNP |
| AMPL2000609 | 3 | MNP |
| AMPL2000611 | 2 | MNP |
| AMPL2000613 | 4 | MNP |
| AMPL2000614 | 3 | MNP |
| AMPL2000615 | 2 | MNP |
| AMPL2000616 | 3 | MNP |
| AMPL2000617 | 3 | MNP |
| AMPL2000618 | 7 | MNP |
| AMPL2000619 | 4 | MNP |
| AMPL2000620 | 3 | MNP |
| AMPL2000621 | 4 | MNP |
| AMPL2000622 | 3 | MNP |
| AMPL2000623 | 2 | MNP |
| AMPL2000624 | 3 | MNP |
| AMPL2000625 | 3 | MNP |
| AMPL2000626 | 3 | MNP |
| AMPL2000627 | 3 | MNP |
| AMPL2000629 | 1 | MNP |
| AMPL2000630 | 2 | MNP |
| AMPL2000631 | 2 | MNP |
| AMPL2000632 | 3 | MNP |
| AMPL2000633 | 5 | MNP |
| AMPL2000634 | 3 | MNP |
| AMPL2000635 | 2 | MNP |
| AMPL2000636 | 4 | MNP |
| AMPL2000637 | 3 | MNP |
| AMPL2000638 | 5 | MNP |
| AMPL2000639 | 3 | MNP |
| AMPL2000640 | 4 | MNP |
| AMPL2000641 | 3 | MNP |

| | | |
|---|---|---|
| AMPL2000642 | 3 | MNP |
| AMPL2000643 | 4 | MNP |
| AMPL2000644 | 4 | MNP |
| AMPL2000645 | 4 | MNP |
| AMPL2000646 | 3 | MNP |
| AMPL2000648 | 2 | MNP |
| AMPL2000649 | 4 | MNP |
| AMPL2000650 | 2 | MNP |
| AMPL2000651 | 3 | MNP |
| AMPL2000652 | 2 | MNP |
| AMPL2000653 | 3 | MNP |
| AMPL2000654 | 3 | MNP |
| AMPL2000655 | 5 | MNP |
| AMPL2000657 | 2 | MNP |
| AMPL2000658 | 3 | MNP |
| AMPL2000659 | 3 | MNP |
| AMPL2000660 | 3 | MNP |
| AMPL2000661 | 3 | MNP |
| AMPL2000662 | 2 | MNP |
| AMPL2000663 | 3 | MNP |
| AMPL2000664 | 3 | MNP |
| AMPL2000665 | 3 | MNP |
| AMPL2000666 | 2 | MNP |
| AMPL2000667 | 2 | MNP |
| AMPL2000668 | 3 | MNP |
| AMPL2000669 | 3 | MNP |
| AMPL2000670 | 3 | MNP |
| AMPL2000671 | 2 | MNP |
| AMPL2000672 | 2 | MNP |
| AMPL2000673 | 2 | MNP |
| AMPL2000675 | 4 | MNP |
| AMPL2000676 | 3 | MNP |
| AMPL2000677 | 4 | MNP |
| AMPL2000678 | 2 | MNP |
| AMPL2000680 | 3 | MNP |
| AMPL2000681 | 2 | MNP |
| AMPL2000682 | 2 | MNP |
| AMPL2000683 | 4 | MNP |
| AMPL2000684 | 3 | MNP |
| AMPL2000685 | 3 | MNP |
| AMPL2000686 | 3 | MNP |
| AMPL2000687 | 3 | MNP |
| AMPL2000688 | 2 | MNP |
| AMPL2000689 | 6 | MNP |
| AMPL2000690 | 3 | MNP |
| AMPL2000691 | 3 | MNP |
| AMPL2000692 | 4 | MNP |
| AMPL2000693 | 4 | MNP |

| | | |
|---|---|---|
| AMPL2000694 | 3 | MNP |
| AMPL2000696 | 4 | MNP |
| AMPL2000697 | 3 | MNP |
| AMPL2000698 | 3 | MNP |
| AMPL2000699 | 4 | MNP |
| AMPL2000700 | 1 | MNP |
| AMPL2000701 | 3 | MNP |
| AMPL2000702 | 2 | MNP |
| AMPL2000703 | 4 | MNP |
| AMPL2000705 | 4 | MNP |
| AMPL2000706 | 4 | MNP |
| AMPL2000707 | 2 | MNP |
| AMPL2000708 | 3 | MNP |
| AMPL2000709 | 3 | MNP |
| AMPL2000712 | 4 | MNP |
| AMPL2000713 | 2 | MNP |
| AMPL2000714 | 2 | MNP |
| AMPL2000715 | 2 | MNP |
| AMPL2000716 | 2 | MNP |
| AMPL2000717 | 2 | MNP |
| AMPL2000718 | 3 | MNP |
| AMPL1563850_1_1 | 2 | SSR |
| AMPL1563551_1_1 | 4 | SSR |
| AMPL1563795_1_2 | 1 | SSR |
| AMPL1563350_1_1 | 3 | SSR |
| AMPL1563506_1_2 | 2 | SSR |
| AMPL1563769_1_1 | 2 | SSR |
| AMPL1563321_1_2 | 3 | SSR |
| AMPL1563384_2_1 | 1 | SSR |
| AMPL1563719_2_1 | 1 | SSR |
| AMPL1563715_1_2 | 1 | SSR |
| AMPL1563642_2_1 | 1 | SSR |
| AMPL1563243_1_1 | 3 | SSR |
| AMPL1563699_2_1 | 1 | SSR |
| AMPL1563475_1_1 | 1 | SSR |
| AMPL1563821_1_2 | 2 | SSR |
| AMPL1563171_2_2 | 2 | SSR |
| AMPL1563305_1_1 | 5 | SSR |
| AMPL1563781_1_1 | 2 | SSR |
| AMPL1563247_1_1 | 5 | SSR |
| AMPL1563269_1_1 | 4 | SSR |
| AMPL1563523_1_1 | 4 | SSR |
| AMPL1563354_2_1 | 2 | SSR |
| AMPL1563249_1_1 | 6 | SSR |
| AMPL1563271_1_1 | 3 | SSR |
| AMPL1563801_1_1 | 4 | SSR |
| AMPL1563164_1_1 | 4 | SSR |
| AMPL1563476_1_1 | 2 | SSR |

| | | |
|---|---|---|
| AMPL1563273_1_1 | 5 | SSR |
| AMPL1563135_1_1 | 6 | SSR |
| AMPL1563483_1_1 | 1 | SSR |
| AMPL1563531_1_3 | 2 | SSR |
| AMPL1563331_2_1 | 1 | SSR |
| AMPL1563556_1_1 | 3 | SSR |
| AMPL1563787_1_1 | 5 | SSR |
| AMPL1563596_2_1 | 4 | SSR |
| AMPL1563413_1_1 | 5 | SSR |
| AMPL1563346_1_1 | 1 | SSR |
| AMPL1563901_1_1 | 3 | SSR |
| AMPL1563131_1_1 | 6 | SSR |
| AMPL1563475_1_2 | 1 | SSR |
| AMPL1563278_1_1 | 2 | SSR |
| AMPL1563125_1_1 | 4 | SSR |
| AMPL1563293_1_2 | 1 | SSR |
| AMPL1563145_1_1 | 7 | SSR |
| AMPL1563204_1_1 | 2 | SSR |
| AMPL1563331_1_1 | 1 | SSR |
| AMPL1563505_1_1 | 6 | SSR |
| AMPL1563809_1_2 | 2 | SSR |
| AMPL1563554_1_1 | 3 | SSR |
| AMPL1563812_1_1 | 3 | SSR |
| AMPL1563224_2_1 | 1 | SSR |
| AMPL1563896_1_2 | 2 | SSR |
| AMPL1563241_1_2 | 5 | SSR |
| AMPL1563335_1_2 | 2 | SSR |
| AMPL1563354_3_1 | 1 | SSR |
| AMPL1563715_1_3 | 4 | SSR |
| AMPL1563813_1_1 | 3 | SSR |
| AMPL1563796_1_1 | 4 | SSR |
| AMPL1563590_1_2 | 3 | SSR |
| AMPL1563668_1_1 | 6 | SSR |
| AMPL1563141_1_1 | 6 | SSR |
| AMPL1563775_1_1 | 6 | SSR |
| AMPL1563224_1_1 | 3 | SSR |
| AMPL1563124_1_1 | 3 | SSR |
| AMPL1563321_1_1 | 1 | SSR |
| AMPL1563204_1_3 | 2 | SSR |
| AMPL1563473_1_2 | 3 | SSR |
| AMPL1563717_1_1 | 3 | SSR |
| AMPL1563606_1_1 | 1 | SSR |
| AMPL1563816_1_1 | 2 | SSR |
| AMPL1563903_1_1 | 4 | SSR |
| AMPL1563146_1_1 | 1 | SSR |
| AMPL1563228_1_2 | 1 | SSR |
| AMPL1563599_1_1 | 5 | SSR |
| AMPL1563323_2_1 | 2 | SSR |

| | | |
|---|---|---|
| AMPL1563408_1_1 | 2 | SSR |
| AMPL1563341_1_1 | 4 | SSR |
| AMPL1563825_1_1 | 4 | SSR |
| AMPL1563407_1_1 | 6 | SSR |
| AMPL1563406_1_1 | 4 | SSR |
| AMPL1563132_1_1 | 1 | SSR |
| AMPL1563372_1_2 | 2 | SSR |
| AMPL1563508_2_1 | 1 | SSR |
| AMPL1563160_1_1 | 7 | SSR |
| AMPL1563699_1_2 | 2 | SSR |
| AMPL1563807_1_1 | 1 | SSR |
| AMPL1563594_1_1 | 3 | SSR |
| AMPL1563502_1_1 | 6 | SSR |
| AMPL1563496_1_1 | 4 | SSR |
| AMPL1563530_1_1 | 2 | SSR |
| AMPL1563144_1_1 | 5 | SSR |
| AMPL1563514_1_1 | 4 | SSR |
| AMPL1563262_1_2 | 1 | SSR |
| AMPL1563265_1_1 | 2 | SSR |
| AMPL1563293_2_1 | 1 | SSR |
| AMPL1563379_1_1 | 3 | SSR |
| AMPL1563483_2_1 | 1 | SSR |
| AMPL1563224_2_2 | 1 | SSR |
| AMPL1563816_1_2 | 3 | SSR |
| AMPL1563211_1_1 | 1 | SSR |
| AMPL1563597_3_1 | 1 | SSR |
| AMPL1563139_1_1 | 4 | SSR |
| AMPL1563248_1_2 | 5 | SSR |
| AMPL1563546_2_1 | 3 | SSR |
| AMPL1563891_1_1 | 3 | SSR |
| AMPL1563226_1_1 | 3 | SSR |
| AMPL1563532_1_2 | 3 | SSR |
| AMPL1563905_1_1 | 3 | SSR |
| AMPL1563688_1_1 | 2 | SSR |
| AMPL1563155_2_1 | 3 | SSR |
| AMPL1563402_1_1 | 4 | SSR |
| AMPL1563271_2_1 | 1 | SSR |
| AMPL1563228_1_1 | 3 | SSR |
| AMPL1563876_1_1 | 6 | SSR |
| AMPL1563171_2_1 | 3 | SSR |
| AMPL1563879_1_1 | 2 | SSR |
| AMPL1563699_2_2 | 2 | SSR |
| AMPL1563719_1_1 | 5 | SSR |
| AMPL1563829_2_1 | 2 | SSR |
| AMPL1563416_1_1 | 3 | SSR |
| AMPL1563597_1_1 | 3 | SSR |
| AMPL1563235_1_1 | 2 | SSR |
| AMPL1563822_1_1 | 4 | SSR |

| | | |
|---|---|---|
| AMPL1563877_1_1 | 4 | SSR |
| AMPL1563791_2_2 | 1 | SSR |
| AMPL1563803_1_2 | 1 | SSR |
| AMPL1563867_2_1 | 1 | SSR |
| AMPL1563726_1_1 | 4 | SSR |
| AMPL1563665_3_1 | 1 | SSR |
| AMPL1563475_2_1 | 7 | SSR |
| AMPL1563618_1_1 | 5 | SSR |
| AMPL1563171_1_1 | 1 | SSR |
| AMPL1563151_1_1 | 2 | SSR |
| AMPL1563774_1_1 | 5 | SSR |
| AMPL1563129_1_1 | 3 | SSR |
| AMPL1563530_2_1 | 1 | SSR |
| AMPL1563642_1_1 | 1 | SSR |
| AMPL1563338_1_1 | 5 | SSR |
| AMPL1563830_1_1 | 6 | SSR |
| AMPL1563125_1_2 | 4 | SSR |
| AMPL1563778_1_1 | 3 | SSR |
| AMPL1563578_2_1 | 3 | SSR |
| AMPL1563354_1_1 | 2 | SSR |
| AMPL1563355_1_1 | 4 | SSR |
| AMPL1563823_1_2 | 4 | SSR |
| AMPL1563596_1_1 | 3 | SSR |
| AMPL1563372_1_1 | 3 | SSR |
| AMPL1563371_1_1 | 7 | SSR |
| AMPL1563330_1_1 | 1 | SSR |
| AMPL1563732_1_2 | 2 | SSR |
| AMPL1563262_1_1 | 2 | SSR |
| AMPL1563823_1_1 | 1 | SSR |
| AMPL1563791_2_1 | 1 | SSR |
| AMPL1563190_1_2 | 1 | SSR |
| AMPL1563151_2_1 | 3 | SSR |
| AMPL1563230_2_1 | 3 | SSR |
| AMPL1563531_1_2 | 3 | SSR |
| AMPL1563531_1_1 | 2 | SSR |
| AMPL1563789_1_2 | 3 | SSR |
| AMPL1563503_1_1 | 5 | SSR |
| AMPL1563780_2_2 | 1 | SSR |
| AMPL1563277_1_1 | 3 | SSR |
| AMPL1563868_2_1 | 2 | SSR |
| AMPL1563606_2_1 | 2 | SSR |
| AMPL1563149_1_1 | 8 | SSR |
| AMPL1563883_1_1 | 3 | SSR |
| AMPL1563229_1_1 | 6 | SSR |
| AMPL1563820_1_1 | 3 | SSR |
| AMPL1563791_1_3 | 2 | SSR |
| AMPL1563804_2_1 | 4 | SSR |
| AMPL1562999_1_1 | 5 | SSR |

| | | |
|---|---|---|
| AMPL1563170_1_1 | 2 | SSR |
| AMPL1563854_2_1 | 4 | SSR |
| AMPL1563384_1_1 | 2 | SSR |
| AMPL1563854_2_2 | 3 | SSR |
| AMPL1563510_1_1 | 2 | SSR |
| AMPL1563715_1_1 | 2 | SSR |
| AMPL1563155_1_1 | 1 | SSR |
| AMPL1563450_2_1 | 1 | SSR |
| AMPL1563167_1_1 | 3 | SSR |
| AMPL1563906_1_1 | 5 | SSR |
| AMPL1563590_1_1 | 2 | SSR |
| AMPL1563638_2_1 | 2 | SSR |
| AMPL1563241_1_1 | 1 | SSR |
| AMPL1563201_1_1 | 3 | SSR |
| AMPL1563374_1_1 | 1 | SSR |
| AMPL1563884_1_1 | 1 | SSR |
| AMPL1563148_1_1 | 5 | SSR |
| AMPL1563128_1_1 | 7 | SSR |
| AMPL1563546_1_1 | 1 | SSR |
| AMPL1563566_1_1 | 3 | SSR |
| AMPL1563140_1_1 | 4 | SSR |
| AMPL1563572_1_1 | 5 | SSR |
| AMPL1563104_1_1 | 4 | SSR |
| AMPL1563391_1_1 | 7 | SSR |
| AMPL1563542_1_1 | 4 | SSR |
| AMPL1562757_1_1 | 3 | SSR |
| AMPL1563338_2_1 | 2 | SSR |
| AMPL1563789_1_1 | 2 | SSR |
| AMPL1563856_1_1 | 2 | SSR |
| AMPL1563393_1_1 | 3 | SSR |
| AMPL1563335_1_1 | 2 | SSR |
| AMPL1563300_1_1 | 6 | SSR |
| AMPL1563280_2_1 | 5 | SSR |
| AMPL1563644_1_1 | 4 | SSR |
| AMPL1563323_1_1 | 2 | SSR |
| AMPL1563526_1_1 | 2 | SSR |
| AMPL1563829_1_2 | 2 | SSR |
| AMPL1563309_2_1 | 2 | SSR |
| AMPL1563816_1_3 | 2 | SSR |
| AMPL1563487_1_1 | 5 | SSR |
| AMPL1563383_2_1 | 3 | SSR |
| AMPL1563452_1_2 | 2 | SSR |
| AMPL1563854_1_1 | 4 | SSR |
| AMPL1563650_1_1 | 3 | SSR |
| AMPL1563264_1_1 | 3 | SSR |
| AMPL1563368_1_1 | 5 | SSR |
| AMPL1563349_2_1 | 2 | SSR |
| AMPL1563216_1_1 | 3 | SSR |

| | | |
|---|---|---|
| AMPL1563825_1_2 | 1 | SSR |
| AMPL1563731_1_1 | 3 | SSR |
| AMPL1563540_1_1 | 3 | SSR |
| AMPL1563204_1_2 | 2 | SSR |
| AMPL1563810_1_1 | 7 | SSR |
| AMPL1563132_2_1 | 5 | SSR |
| AMPL1563804_1_1 | 1 | SSR |
| AMPL1563446_1_2 | 3 | SSR |
| AMPL1563575_1_1 | 1 | SSR |
| AMPL1563237_1_1 | 5 | SSR |
| AMPL1563791_1_1 | 3 | SSR |
| AMPL1563124_2_1 | 1 | SSR |
| AMPL1563671_1_1 | 4 | SSR |
| AMPL1563309_1_1 | 1 | SSR |
| AMPL1563638_1_1 | 1 | SSR |
| AMPL1563393_2_1 | 2 | SSR |
| AMPL1563471_1_1 | 3 | SSR |
| AMPL1563270_1_1 | 2 | SSR |
| AMPL1563750_1_1 | 4 | SSR |
| AMPL1563217_1_1 | 2 | SSR |
| AMPL1563642_2_2 | 3 | SSR |
| AMPL1563141_2_1 | 3 | SSR |
| AMPL1563370_1_1 | 4 | SSR |
| AMPL1563897_1_1 | 4 | SSR |
| AMPL1563807_1_2 | 4 | SSR |
| AMPL1563911_1_1 | 3 | SSR |
| AMPL1563339_2_1 | 1 | SSR |
| AMPL1563183_1_1 | 2 | SSR |
| AMPL1563780_1_1 | 3 | SSR |
| AMPL1563405_1_1 | 5 | SSR |
| AMPL1563287_1_1 | 6 | SSR |
| AMPL1563640_2_1 | 2 | SSR |
| AMPL1563867_1_1 | 2 | SSR |
| AMPL1563177_1_1 | 2 | SSR |
| AMPL1563446_1_1 | 3 | SSR |
| AMPL1563473_1_1 | 2 | SSR |
| AMPL1563699_1_1 | 2 | SSR |
| AMPL1563421_1_1 | 6 | SSR |
| AMPL1563221_1_1 | 3 | SSR |
| AMPL1563640_3_1 | 1 | SSR |
| AMPL1563283_1_1 | 2 | SSR |
| AMPL1563775_2_1 | 2 | SSR |
| AMPL1563597_2_1 | 2 | SSR |
| AMPL1563239_1_1 | 1 | SSR |
| AMPL1563414_1_1 | 1 | SSR |
| AMPL1563866_1_1 | 2 | SSR |
| AMPL1563846_1_1 | 4 | SSR |
| AMPL1563248_1_1 | 4 | SSR |

| | | |
|---|---|---|
| AMPL1563880_1_1 | 4 | SSR |
| AMPL1563893_1_1 | 3 | SSR |
| AMPL1563732_1_1 | 2 | SSR |
| AMPL1563821_1_1 | 3 | SSR |
| AMPL1563270_2_1 | 4 | SSR |
| AMPL1563760_1_1 | 3 | SSR |
| AMPL1563640_1_1 | 2 | SSR |
| AMPL1563665_2_2 | 2 | SSR |
| AMPL1563803_1_1 | 1 | SSR |
| AMPL1563809_1_1 | 1 | SSR |
| AMPL1563795_1_1 | 4 | SSR |
| AMPL1563246_1_1 | 3 | SSR |
| AMPL1563806_1_1 | 1 | SSR |
| AMPL1563867_4_1 | 1 | SSR |
| AMPL1563302_1_1 | 3 | SSR |
| AMPL1563367_1_1 | 3 | SSR |
| AMPL1563850_2_1 | 4 | SSR |
| AMPL1563531_1_4 | 4 | SSR |
| AMPL1563834_1_1 | 1 | SSR |
| AMPL1563826_1_1 | 6 | SSR |
| AMPL1563130_1_1 | 2 | SSR |
| AMPL1563868_1_1 | 2 | SSR |
| AMPL1563450_1_1 | 1 | SSR |
| AMPL1563578_1_1 | 1 | SSR |
| AMPL1563150_1_1 | 2 | SSR |
| AMPL1563601_1_1 | 4 | SSR |
| AMPL1563481_1_1 | 3 | SSR |
| AMPL1563794_1_1 | 2 | SSR |
| AMPL1563506_1_1 | 2 | SSR |
| AMPL1563452_1_1 | 3 | SSR |
| AMPL1563386_1_1 | 2 | SSR |
| AMPL1563891_2_1 | 4 | SSR |
| AMPL1563866_2_1 | 2 | SSR |
| AMPL1563411_1_1 | 1 | SSR |
| AMPL1563211_1_2 | 1 | SSR |
| AMPL1563459_1_1 | 4 | SSR |
| AMPL1563852_1_1 | 4 | SSR |
| AMPL1563872_1_2 | 2 | SSR |
| AMPL1563405_1_2 | 6 | SSR |
| AMPL1563868_1_2 | 2 | SSR |
| AMPL1563206_1_1 | 6 | SSR |
| AMPL1563868_1_3 | 2 | SSR |
| AMPL1563145_1_2 | 3 | SSR |
| AMPL1563241_1_3 | 2 | SSR |
| AMPL1563593_1_1 | 4 | SSR |
| AMPL1563246_1_2 | 1 | SSR |
| AMPL1563780_2_1 | 1 | SSR |
| AMPL1563483_3_1 | 4 | SSR |

| | | |
|---|---|---|
| AMPL1563734_1_1 | 3 | SSR |
| AMPL1563339_1_1 | 4 | SSR |
| AMPL1563896_1_1 | 2 | SSR |
| AMPL1563165_1_1 | 3 | SSR |
| AMPL1563642_3_1 | 1 | SSR |
| AMPL1563521_2_1 | 3 | SSR |
| AMPL1563401_1_1 | 3 | SSR |
| AMPL1563349_1_1 | 3 | SSR |
| AMPL1563872_1_1 | 1 | SSR |
| AMPL1563127_1_1 | 2 | SSR |
| AMPL1563183_1_2 | 2 | SSR |
| AMPL1563237_2_1 | 1 | SSR |
| AMPL1563735_1_1 | 3 | SSR |
| AMPL1563426_1_1 | 3 | SSR |
| AMPL1563775_1_2 | 1 | SSR |
| AMPL1563521_1_1 | 1 | SSR |
| AMPL1563273_1_2 | 3 | SSR |
| AMPL1563844_1_1 | 4 | SSR |
| AMPL1563142_1_1 | 3 | SSR |
| AMPL1563328_1_1 | 5 | SSR |
| AMPL1563508_2_2 | 2 | SSR |
| AMPL1563660_1_1 | 1 | SSR |
| AMPL1563532_1_1 | 5 | SSR |
| AMPL1563665_2_1 | 2 | SSR |
| AMPL1563830_1_2 | 1 | SSR |
| AMPL1563270_1_2 | 3 | SSR |
| AMPL1563508_1_1 | 1 | SSR |
| AMPL1563486_1_1 | 5 | SSR |
| AMPL1562814_1_1 | 2 | SSR |
| AMPL1563697_1_1 | 6 | SSR |
| AMPL1563590_2_1 | 3 | SSR |
| AMPL1563665_1_1 | 1 | SSR |
| AMPL1563136_1_1 | 3 | SSR |
| AMPL1563098_1_1 | 3 | SSR |
| AMPL1563104_2_1 | 2 | SSR |
| AMPL1563739_1_1 | 3 | SSR |
| AMPL1563293_1_1 | 1 | SSR |
| AMPL1563740_1_1 | 4 | SSR |
| AMPL1563829_1_1 | 3 | SSR |
| AMPL1563494_1_1 | 2 | SSR |
| AMPL1563280_1_1 | 4 | SSR |
| AMPL1563867_3_1 | 2 | SSR |
| AMPL1563791_1_2 | 1 | SSR |
| AMPL1563181_1_1 | 2 | SSR |
| AMPL1563812_1_2 | 1 | SSR |
| AMPL1563383_1_1 | 4 | SSR |
| AMPL1563210_1_1 | 5 | SSR |
| AMPL1563578_2_2 | 3 | SSR |

| | | |
|---|---|---|
| AMPL1563230_1_1 | 2 | SSR |
| AMPL1563494_1_2 | 2 | SSR |
| AMPL1563617_1_1 | 2 | SSR |
| AMPL1563209_1_1 | 4 | SSR |
| AMPL1563592_1_1 | 3 | SSR |
| AMPL1563890_1_1 | 2 | SSR |
| AMPL1563853_1_1 | 4 | SSR |
| AMPL1563146_1_2 | 6 | SSR |
| AMPL1563571_1_1 | 2 | SSR |
| AMPL1563794_2_1 | 1 | SSR |
| AMPL1563202_1_1 | 2 | SSR |
| AMPL1563190_1_1 | 4 | SSR |
| AMPL1563606_3_1 | 3 | SSR |
| AMPL1563266_1_1 | 4 | SSR |
