## Supplementary table 4 for "Multiple nucleotide polymorphism DNA markers for the accurate evaluation of genetic variations"

**Supplementary Table 4. Nipponbare rice lines used in this study**

| Sample code | Species | Line name | Ploidy | Hybrid/Inbred | Source |
|---|---|---|---|---|---|
| 200383 | *Oryza sativa L.* | Nipponbare | Diploid | Inbred | Wuhan University |
| 200384 | *Oryza sativa L.* | Nipponbare | Diploid | Inbred | China National Rice Research Institute |
| 200385 | *Oryza sativa L.* | Nipponbare | Diploid | Inbred | China National Rice Research Institute |
| 200386 | *Oryza sativa L.* | Nipponbare | Diploid | Inbred | China National Rice Research Institute |
| 200387 | *Oryza sativa L.* | Nipponbare | Diploid | Inbred | China National Rice Research Institute |
| 200388 | *Oryza sativa L.* | Nipponbare | Diploid | Inbred | China National Rice Research Institute |
| 200389 | *Oryza sativa L.* | Nipponbare | Diploid | Inbred | China National Rice Research Institute |
| 200390 | *Oryza sativa L.* | Nipponbare | Diploid | Inbred | China National Rice Research Institute |
| 200391 | *Oryza sativa L.* | Nipponbare | Diploid | Inbred | China National Rice Research Institute |
| 200392 | *Oryza sativa L.* | Nipponbare | Diploid | Inbred | Huazhong Agricultural University |
| 200393 | *Oryza sativa L.* | Nipponbare | Diploid | Inbred | HunanAgricultural University |
| 200394 | *Oryza sativa L.* | Nipponbare | Diploid | Inbred | HunanAgricultural University |
| 200395 | *Oryza sativa L.* | Nipponbare | Diploid | Inbred | HunanAgricultural University |
| 200396 | *Oryza sativa L.* | Nipponbare | Diploid | Inbred | HunanAgricultural University |
| 200398 | *Oryza sativa L.* | Nipponbare | Diploid | Inbred | Northeast Agricultural University |
| 200399 | *Oryza sativa L.* | Nipponbare | Diploid | Inbred | China National Rice Research Institute |
| 200400 | *Oryza sativa L.* | Nipponbare | Diploid | Inbred | China National Rice Research Institute |
| 200401 | *Oryza sativa L.* | Nipponbare | Diploid | Inbred | Hunan Hybrid Rice Research Center |
| 200402 | *Oryza sativa L.* | Nipponbare | Diploid | Inbred | Hunan Hybrid Rice Research Center |
| 200403 | *Oryza sativa L.* | Nipponbare | Diploid | Inbred | Hunan Hybrid Rice Research Center |
| 200404 | *Oryza sativa L.* | Nipponbare | Diploid | Inbred | Hunan Hybrid Rice Research Center |
| 200405 | *Oryza sativa L.* | Nipponbare | Diploid | Inbred | Anhui Academy of Agricultural Science |
| 200406 | *Oryza sativa L.* | Nipponbare | Diploid | Inbred | The Northeast Institute of Geography and Agroecology |
| 200407 | *Oryza sativa L.* | Nipponbare | Diploid | Inbred | Yangzhou University |
| 200408 | *Oryza sativa L.* | Nipponbare | Diploid | Inbred | Fujian Agriculture and Forestry University |
| 201110 | *Oryza sativa L.* | Nipponbare | Diploid | Inbred | Institute of Genetics and Developmental Biology |
| 201111 | *Oryza sativa L.* | Nipponbare | Diploid | Inbred | Institute of Genetics and Developmental Biology |
| 201112 | *Oryza sativa L.* | Nipponbare | Diploid | Inbred | Institute of Genetics and Developmental Biology |
| 201113 | *Oryza sativa L.* | Nipponbare | Diploid | Inbred | Institute of Genetics and Developmental Biology |
| 201114 | *Oryza sativa L.* | Nipponbare | Diploid | Inbred | Institute of Genetics and Developmental Biology |
