## Supplementary table 5 for "Multiple nucleotide polymorphism DNA markers for the accurate evaluation of genetic variations"

**Supplementary Table 5. Distinct MNP genotypes for each pair of Nipponbare lines**

| Line pair | Number of MNPs compared | Number of distinct MNPs genotypes | Ratio of distinct MNPs genotypes |
|---|---|---|---|
| 201113 vs 201114 | 927 | 2 | 0.22% |
| 201112 vs 201113 | 926 | 2 | 0.22% |
| 201112 vs 201114 | 926 | 2 | 0.22% |
| 201111 vs 201112 | 925 | 1 | 0.11% |
| 201111 vs 201113 | 926 | 2 | 0.22% |
| 201111 vs 201114 | 926 | 1 | 0.11% |
| 201110 vs 201111 | 926 | 0 | 0 |
| 201110 vs 201112 | 925 | 2 | 0.22% |
| 201110 vs 201113 | 926 | 2 | 0.22% |
| 201110 vs 201114 | 926 | 1 | 0.11% |
| 200408 vs 201110 | 926 | 3 | 0.32% |
| 200408 vs 201111 | 926 | 3 | 0.32% |
| 200408 vs 201112 | 926 | 4 | 0.43% |
| 200408 vs 201113 | 927 | 5 | 0.54% |
| 200408 vs 201114 | 927 | 4 | 0.43% |
| 200407 vs 200408 | 925 | 4 | 0.43% |
| 200407 vs 201110 | 924 | 3 | 0.32% |
| 200407 vs 201111 | 924 | 3 | 0.32% |
| 200407 vs 201112 | 924 | 3 | 0.32% |
| 200407 vs 201113 | 925 | 4 | 0.43% |
| 200407 vs 201114 | 925 | 4 | 0.43% |
| 200406 vs 200407 | 924 | 3 | 0.32% |
| 200406 vs 200408 | 925 | 4 | 0.43% |
| 200406 vs 201110 | 924 | 1 | 0.11% |
| 200406 vs 201111 | 924 | 1 | 0.11% |
| 200406 vs 201112 | 924 | 0 | 0 |
| 200406 vs 201113 | 925 | 3 | 0.32% |
| 200406 vs 201114 | 925 | 2 | 0.22% |
| 200405 vs 200406 | 925 | 3 | 0.32% |
| 200405 vs 200407 | 924 | 5 | 0.54% |
| 200405 vs 200408 | 925 | 5 | 0.54% |
| 200405 vs 201110 | 924 | 2 | 0.22% |
| 200405 vs 201111 | 924 | 2 | 0.22% |
| 200405 vs 201112 | 924 | 3 | 0.32% |
| 200405 vs 201113 | 925 | 3 | 0.32% |
| 200405 vs 201114 | 925 | 2 | 0.22% |
| 200404 vs 200405 | 924 | 4 | 0.43% |
| 200404 vs 200406 | 924 | 3 | 0.32% |
| 200404 vs 200407 | 924 | 1 | 0.11% |
| 200404 vs 200408 | 925 | 3 | 0.32% |
| 200404 vs 201110 | 924 | 2 | 0.22% |
| 200404 vs 201111 | 924 | 2 | 0.22% |
| 200404 vs 201112 | 924 | 4 | 0.43% |
| 200404 vs 201113 | 925 | 4 | 0.43% |
| 200404 vs 201114 | 925 | 3 | 0.32% |
| 200403 vs 200404 | 924 | 2 | 0.22% |
| 200403 vs 200405 | 923 | 4 | 0.43% |
| 200403 vs 200406 | 923 | 4 | 0.43% |
| 200403 vs 200407 | 923 | 2 | 0.22% |
| 200403 vs 200408 | 924 | 4 | 0.43% |
| 200403 vs 201110 | 923 | 4 | 0.43% |
| 200403 vs 201111 | 923 | 2 | 0.22% |
| 200403 vs 201112 | 923 | 2 | 0.22% |
| 200403 vs 201113 | 924 | 4 | 0.43% |
| 200403 vs 201114 | 924 | 2 | 0.22% |
| 200402 vs 200403 | 924 | 6 | 0.65% |
| 200402 vs 200404 | 925 | 4 | 0.43% |
| 200402 vs 200405 | 925 | 4 | 0.43% |
| 200402 vs 200406 | 925 | 2 | 0.22% |
| 200402 vs 200407 | 925 | 5 | 0.54% |
| 200402 vs 200408 | 926 | 5 | 0.54% |
| 200402 vs 201110 | 925 | 2 | 0.22% |
| 200402 vs 201111 | 925 | 2 | 0.22% |
| 200402 vs 201112 | 925 | 3 | 0.32% |
| 200402 vs 201113 | 926 | 5 | 0.54% |
| 200402 vs 201114 | 926 | 3 | 0.32% |
| 200401 vs 200402 | 926 | 4 | 0.43% |
| 200401 vs 200403 | 924 | 2 | 0.22% |
| 200401 vs 200404 | 925 | 4 | 0.43% |

| | | | |
|---|---|---|---|
| 200401 vs 200405 | 925 | 3 | 0.32% |
| 200401 vs 200406 | 925 | 2 | 0.22% |
| 200401 vs 200407 | 925 | 2 | 0.22% |
| 200401 vs 200408 | 926 | 4 | 0.43% |
| 200401 vs 201110 | 925 | 2 | 0.22% |
| 200401 vs 201111 | 925 | 1 | 0.11% |
| 200401 vs 201112 | 925 | 2 | 0.22% |
| 200401 vs 201113 | 926 | 2 | 0.22% |
| 200401 vs 201114 | 926 | 2 | 0.22% |
| 200400 vs 200401 | 923 | 2 | 0.22% |
| 200400 vs 200402 | 923 | 5 | 0.54% |
| 200400 vs 200403 | 922 | 2 | 0.22% |
| 200400 vs 200404 | 922 | 5 | 0.54% |
| 200400 vs 200405 | 922 | 5 | 0.54% |
| 200400 vs 200406 | 922 | 4 | 0.43% |
| 200400 vs 200407 | 923 | 2 | 0.22% |
| 200400 vs 200408 | 923 | 5 | 0.54% |
| 200400 vs 201110 | 922 | 4 | 0.43% |
| 200400 vs 201111 | 922 | 2 | 0.22% |
| 200400 vs 201112 | 923 | 4 | 0.43% |
| 200400 vs 201113 | 923 | 4 | 0.43% |
| 200400 vs 201114 | 923 | 4 | 0.43% |
| 200399 vs 200400 | 923 | 4 | 0.43% |
| 200399 vs 200401 | 925 | 3 | 0.32% |
| 200399 vs 200402 | 925 | 4 | 0.43% |
| 200399 vs 200403 | 924 | 2 | 0.22% |
| 200399 vs 200404 | 924 | 3 | 0.32% |
| 200399 vs 200405 | 924 | 4 | 0.43% |
| 200399 vs 200406 | 924 | 2 | 0.22% |
| 200399 vs 200407 | 924 | 5 | 0.54% |
| 200399 vs 200408 | 925 | 5 | 0.54% |
| 200399 vs 201110 | 924 | 2 | 0.22% |
| 200399 vs 201111 | 924 | 2 | 0.22% |
| 200399 vs 201112 | 924 | 1 | 0.11% |
| 200399 vs 201113 | 925 | 3 | 0.32% |
| 200399 vs 201114 | 925 | 2 | 0.22% |
| 200398 vs 200399 | 925 | 5 | 0.54% |
| 200398 vs 200400 | 923 | 6 | 0.65% |
| 200398 vs 200401 | 926 | 4 | 0.43% |
| 200398 vs 200402 | 926 | 5 | 0.54% |
| 200398 vs 200403 | 924 | 7 | 0.76% |
| 200398 vs 200404 | 925 | 6 | 0.65% |
| 200398 vs 200405 | 925 | 6 | 0.65% |
| 200398 vs 200406 | 925 | 3 | 0.32% |
| 200398 vs 200407 | 925 | 4 | 0.43% |
| 200398 vs 200408 | 926 | 7 | 0.76% |
| 200398 vs 201110 | 925 | 4 | 0.43% |
| 200398 vs 201111 | 925 | 4 | 0.43% |
| 200398 vs 201112 | 925 | 4 | 0.43% |
| 200398 vs 201113 | 926 | 8 | 0.86% |
| 200398 vs 201114 | 926 | 5 | 0.54% |
| 200396 vs 200398 | 926 | 3 | 0.32% |
| 200396 vs 200399 | 925 | 2 | 0.22% |
| 200396 vs 200400 | 923 | 3 | 0.33% |
| 200396 vs 200401 | 926 | 2 | 0.22% |
| 200396 vs 200402 | 926 | 2 | 0.22% |
| 200396 vs 200403 | 924 | 3 | 0.32% |
| 200396 vs 200404 | 925 | 2 | 0.22% |
| 200396 vs 200405 | 925 | 2 | 0.22% |
| 200396 vs 200406 | 925 | 0 | 0 |
| 200396 vs 200407 | 925 | 3 | 0.32% |
| 200396 vs 200408 | 927 | 3 | 0.32% |
| 200396 vs 201110 | 926 | 0 | 0 |
| 200396 vs 201111 | 926 | 0 | 0 |
| 200396 vs 201112 | 926 | 1 | 0.11% |
| 200396 vs 201113 | 927 | 2 | 0.22% |
| 200396 vs 201114 | 927 | 1 | 0.11% |
| 200395 vs 200396 | 927 | 0 | 0 |
| 200395 vs 200398 | 926 | 3 | 0.32% |
| 200395 vs 200399 | 925 | 2 | 0.22% |
| 200395 vs 200400 | 923 | 4 | 0.43% |
| 200395 vs 200401 | 926 | 3 | 0.32% |

| | | | |
|---|---|---|---|
| 200395 vs 200402 | 926 | 2 | 0.22% |
| 200395 vs 200403 | 924 | 4 | 0.43% |
| 200395 vs 200404 | 925 | 3 | 0.32% |
| 200395 vs 200405 | 925 | 3 | 0.32% |
| 200395 vs 200406 | 925 | 0 | 0 |
| 200395 vs 200407 | 925 | 3 | 0.32% |
| 200395 vs 200408 | 927 | 4 | 0.43% |
| 200395 vs 201110 | 926 | 1 | 0.11% |
| 200395 vs 201111 | 926 | 1 | 0.11% |
| 200395 vs 201112 | 926 | 1 | 0.11% |
| 200395 vs 201113 | 927 | 3 | 0.32% |
| 200395 vs 201114 | 927 | 2 | 0.22% |
| 200394 vs 200395 | 925 | 1 | 0.11% |
| 200394 vs 200396 | 925 | 0 | 0 |
| 200394 vs 200398 | 924 | 4 | 0.43% |
| 200394 vs 200399 | 923 | 1 | 0.11% |
| 200394 vs 200400 | 923 | 3 | 0.33% |
| 200394 vs 200401 | 924 | 1 | 0.11% |
| 200394 vs 200402 | 924 | 2 | 0.22% |
| 200394 vs 200403 | 922 | 1 | 0.11% |
| 200394 vs 200404 | 923 | 2 | 0.22% |
| 200394 vs 200405 | 923 | 2 | 0.22% |
| 200394 vs 200406 | 923 | 1 | 0.11% |
| 200394 vs 200407 | 924 | 3 | 0.32% |
| 200394 vs 200408 | 925 | 3 | 0.32% |
| 200394 vs 201110 | 924 | 0 | 0 |
| 200394 vs 201111 | 924 | 0 | 0 |
| 200394 vs 201112 | 925 | 1 | 0.11% |
| 200394 vs 201113 | 925 | 1 | 0.11% |
| 200394 vs 201114 | 925 | 1 | 0.11% |
| 200393 vs 200394 | 924 | 1 | 0.11% |
| 200393 vs 200395 | 924 | 4 | 0.43% |
| 200393 vs 200396 | 924 | 3 | 0.32% |
| 200393 vs 200398 | 924 | 7 | 0.76% |
| 200393 vs 200399 | 923 | 2 | 0.22% |
| 200393 vs 200400 | 923 | 3 | 0.33% |
| 200393 vs 200401 | 924 | 3 | 0.32% |
| 200393 vs 200402 | 924 | 5 | 0.54% |
| 200393 vs 200403 | 922 | 0 | 0 |
| 200393 vs 200404 | 923 | 2 | 0.22% |
| 200393 vs 200405 | 923 | 4 | 0.43% |
| 200393 vs 200406 | 923 | 3 | 0.33% |
| 200393 vs 200407 | 924 | 3 | 0.32% |
| 200393 vs 200408 | 924 | 3 | 0.32% |
| 200393 vs 201110 | 923 | 4 | 0.43% |
| 200393 vs 201111 | 923 | 2 | 0.22% |
| 200393 vs 201112 | 924 | 2 | 0.22% |
| 200393 vs 201113 | 924 | 3 | 0.32% |
| 200393 vs 201114 | 924 | 2 | 0.22% |
| 200392 vs 200393 | 924 | 5 | 0.54% |
| 200392 vs 200394 | 924 | 5 | 0.54% |
| 200392 vs 200395 | 926 | 4 | 0.43% |
| 200392 vs 200396 | 926 | 4 | 0.43% |
| 200392 vs 200398 | 926 | 7 | 0.76% |
| 200392 vs 200399 | 925 | 6 | 0.65% |
| 200392 vs 200400 | 923 | 7 | 0.76% |
| 200392 vs 200401 | 926 | 6 | 0.65% |
| 200392 vs 200402 | 926 | 6 | 0.65% |
| 200392 vs 200403 | 924 | 5 | 0.54% |
| 200392 vs 200404 | 925 | 3 | 0.32% |
| 200392 vs 200405 | 925 | 7 | 0.76% |
| 200392 vs 200406 | 925 | 4 | 0.43% |
| 200392 vs 200407 | 925 | 3 | 0.32% |
| 200392 vs 200408 | 926 | 6 | 0.65% |
| 200392 vs 201110 | 925 | 5 | 0.54% |
| 200392 vs 201111 | 925 | 5 | 0.54% |
| 200392 vs 201112 | 925 | 4 | 0.43% |
| 200392 vs 201113 | 926 | 6 | 0.65% |
| 200392 vs 201114 | 926 | 6 | 0.65% |
| 200391 vs 200392 | 926 | 5 | 0.54% |
| 200391 vs 200393 | 924 | 2 | 0.22% |
| 200391 vs 200394 | 925 | 0 | 0 |

| | | | |
|---|---|---|---|
| 200391 vs 200395 | 927 | 1 | 0.11% |
| 200391 vs 200396 | 927 | 0 | 0 |
| 200391 vs 200398 | 926 | 4 | 0.43% |
| 200391 vs 200399 | 925 | 2 | 0.22% |
| 200391 vs 200400 | 923 | 2 | 0.22% |
| 200391 vs 200401 | 926 | 1 | 0.11% |
| 200391 vs 200402 | 926 | 2 | 0.22% |
| 200391 vs 200403 | 924 | 2 | 0.22% |
| 200391 vs 200404 | 925 | 2 | 0.22% |
| 200391 vs 200405 | 925 | 2 | 0.22% |
| 200391 vs 200406 | 925 | 1 | 0.11% |
| 200391 vs 200407 | 925 | 3 | 0.32% |
| 200391 vs 200408 | 927 | 3 | 0.32% |
| 200391 vs 201110 | 926 | 0 | 0 |
| 200391 vs 201111 | 926 | 0 | 0 |
| 200391 vs 201112 | 926 | 1 | 0.11% |
| 200391 vs 201113 | 927 | 1 | 0.11% |
| 200391 vs 201114 | 927 | 1 | 0.11% |
| 200390 vs 200391 | 926 | 1 | 0.11% |
| 200390 vs 200392 | 926 | 6 | 0.65% |
| 200390 vs 200393 | 924 | 3 | 0.32% |
| 200390 vs 200394 | 924 | 1 | 0.11% |
| 200390 vs 200395 | 926 | 3 | 0.32% |
| 200390 vs 200396 | 926 | 2 | 0.22% |
| 200390 vs 200398 | 926 | 6 | 0.65% |
| 200390 vs 200399 | 925 | 3 | 0.32% |
| 200390 vs 200400 | 923 | 3 | 0.33% |
| 200390 vs 200401 | 926 | 2 | 0.22% |
| 200390 vs 200402 | 926 | 4 | 0.43% |
| 200390 vs 200403 | 924 | 3 | 0.32% |
| 200390 vs 200404 | 925 | 4 | 0.43% |
| 200390 vs 200405 | 925 | 3 | 0.32% |
| 200390 vs 200406 | 925 | 2 | 0.22% |
| 200390 vs 200407 | 925 | 4 | 0.43% |
| 200390 vs 200408 | 926 | 3 | 0.32% |
| 200390 vs 201110 | 925 | 2 | 0.22% |
| 200390 vs 201111 | 925 | 1 | 0.11% |
| 200390 vs 201112 | 925 | 2 | 0.22% |
| 200390 vs 201113 | 926 | 2 | 0.22% |
| 200390 vs 201114 | 926 | 2 | 0.22% |
| 200389 vs 200390 | 926 | 2 | 0.22% |
| 200389 vs 200391 | 927 | 1 | 0.11% |
| 200389 vs 200392 | 926 | 4 | 0.43% |
| 200389 vs 200393 | 924 | 3 | 0.32% |
| 200389 vs 200394 | 925 | 1 | 0.11% |
| 200389 vs 200395 | 927 | 0 | 0 |
| 200389 vs 200396 | 927 | 0 | 0 |
| 200389 vs 200398 | 926 | 3 | 0.32% |
| 200389 vs 200399 | 925 | 2 | 0.22% |
| 200389 vs 200400 | 923 | 4 | 0.43% |
| 200389 vs 200401 | 926 | 2 | 0.22% |
| 200389 vs 200402 | 926 | 2 | 0.22% |
| 200389 vs 200403 | 924 | 3 | 0.32% |
| 200389 vs 200404 | 925 | 3 | 0.32% |
| 200389 vs 200405 | 925 | 3 | 0.32% |
| 200389 vs 200406 | 925 | 0 | 0 |
| 200389 vs 200407 | 925 | 3 | 0.32% |
| 200389 vs 200408 | 927 | 4 | 0.43% |
| 200389 vs 201110 | 926 | 1 | 0.11% |
| 200389 vs 201111 | 926 | 1 | 0.11% |
| 200389 vs 201112 | 926 | 0 | 0 |
| 200389 vs 201113 | 927 | 2 | 0.22% |
| 200389 vs 201114 | 927 | 2 | 0.22% |
| 200388 vs 200389 | 923 | 2 | 0.22% |
| 200388 vs 200390 | 923 | 2 | 0.22% |
| 200388 vs 200391 | 923 | 1 | 0.11% |
| 200388 vs 200392 | 923 | 6 | 0.65% |
| 200388 vs 200393 | 921 | 3 | 0.33% |
| 200388 vs 200394 | 921 | 1 | 0.11% |
| 200388 vs 200395 | 923 | 3 | 0.33% |
| 200388 vs 200396 | 923 | 2 | 0.22% |
| 200388 vs 200398 | 923 | 6 | 0.65% |

| | | | |
|---|---|---|---|
| 200388 vs 200399 | 923 | 3 | 0.33% |
| 200388 vs 200400 | 921 | 4 | 0.43% |
| 200388 vs 200401 | 923 | 2 | 0.22% |
| 200388 vs 200402 | 923 | 4 | 0.43% |
| 200388 vs 200403 | 923 | 3 | 0.33% |
| 200388 vs 200404 | 923 | 4 | 0.43% |
| 200388 vs 200405 | 923 | 1 | 0.11% |
| 200388 vs 200406 | 923 | 2 | 0.22% |
| 200388 vs 200407 | 922 | 4 | 0.43% |
| 200388 vs 200408 | 923 | 4 | 0.43% |
| 200388 vs 201110 | 922 | 3 | 0.33% |
| 200388 vs 201111 | 922 | 1 | 0.11% |
| 200388 vs 201112 | 922 | 2 | 0.22% |
| 200388 vs 201113 | 923 | 2 | 0.22% |
| 200388 vs 201114 | 923 | 1 | 0.11% |
| 200387 vs 200388 | 923 | 3 | 0.33% |
| 200387 vs 200389 | 926 | 3 | 0.32% |
| 200387 vs 200390 | 925 | 2 | 0.22% |
| 200387 vs 200391 | 926 | 1 | 0.11% |
| 200387 vs 200392 | 925 | 7 | 0.76% |
| 200387 vs 200393 | 923 | 4 | 0.43% |
| 200387 vs 200394 | 924 | 2 | 0.22% |
| 200387 vs 200395 | 926 | 3 | 0.32% |
| 200387 vs 200396 | 926 | 2 | 0.22% |
| 200387 vs 200398 | 925 | 5 | 0.54% |
| 200387 vs 200399 | 925 | 4 | 0.43% |
| 200387 vs 200400 | 923 | 1 | 0.11% |
| 200387 vs 200401 | 925 | 1 | 0.11% |
| 200387 vs 200402 | 925 | 5 | 0.54% |
| 200387 vs 200403 | 924 | 3 | 0.32% |
| 200387 vs 200404 | 924 | 5 | 0.54% |
| 200387 vs 200405 | 924 | 4 | 0.43% |
| 200387 vs 200406 | 924 | 3 | 0.32% |
| 200387 vs 200407 | 924 | 2 | 0.22% |
| 200387 vs 200408 | 926 | 5 | 0.54% |
| 200387 vs 201110 | 925 | 3 | 0.32% |
| 200387 vs 201111 | 925 | 1 | 0.11% |
| 200387 vs 201112 | 925 | 3 | 0.32% |
| 200387 vs 201113 | 926 | 3 | 0.32% |
| 200387 vs 201114 | 926 | 3 | 0.32% |
| 200386 vs 200387 | 925 | 12 | 1.30% |
| 200386 vs 200388 | 923 | 14 | 1.52% |
| 200386 vs 200389 | 926 | 15 | 1.62% |
| 200386 vs 200390 | 926 | 14 | 1.51% |
| 200386 vs 200391 | 926 | 13 | 1.40% |
| 200386 vs 200392 | 926 | 19 | 2.05% |
| 200386 vs 200393 | 924 | 16 | 1.73% |
| 200386 vs 200394 | 924 | 14 | 1.52% |
| 200386 vs 200395 | 926 | 14 | 1.51% |
| 200386 vs 200396 | 926 | 13 | 1.40% |
| 200386 vs 200398 | 926 | 17 | 1.84% |
| 200386 vs 200399 | 925 | 15 | 1.62% |
| 200386 vs 200400 | 923 | 12 | 1.30% |
| 200386 vs 200401 | 926 | 14 | 1.51% |
| 200386 vs 200402 | 926 | 16 | 1.73% |
| 200386 vs 200403 | 924 | 15 | 1.62% |
| 200386 vs 200404 | 925 | 16 | 1.73% |
| 200386 vs 200405 | 925 | 16 | 1.73% |
| 200386 vs 200406 | 925 | 15 | 1.62% |
| 200386 vs 200407 | 925 | 15 | 1.62% |
| 200386 vs 200408 | 926 | 17 | 1.84% |
| 200386 vs 201110 | 925 | 13 | 1.41% |
| 200386 vs 201111 | 925 | 12 | 1.30% |
| 200386 vs 201112 | 925 | 15 | 1.62% |
| 200386 vs 201113 | 926 | 15 | 1.62% |
| 200386 vs 201114 | 926 | 15 | 1.62% |
| 200385 vs 200386 | 926 | 16 | 1.73% |
| 200385 vs 200387 | 925 | 2 | 0.22% |
| 200385 vs 200388 | 923 | 4 | 0.43% |
| 200385 vs 200389 | 926 | 4 | 0.43% |
| 200385 vs 200390 | 926 | 3 | 0.32% |
| 200385 vs 200391 | 926 | 3 | 0.32% |

| | | | |
|---|---|---|---|
| 200385 vs 200392 | 926 | 8 | 0.86% |
| 200385 vs 200393 | 924 | 5 | 0.54% |
| 200385 vs 200394 | 924 | 3 | 0.32% |
| 200385 vs 200395 | 926 | 4 | 0.43% |
| 200385 vs 200396 | 926 | 3 | 0.32% |
| 200385 vs 200398 | 926 | 5 | 0.54% |
| 200385 vs 200399 | 925 | 4 | 0.43% |
| 200385 vs 200400 | 923 | 3 | 0.33% |
| 200385 vs 200401 | 926 | 2 | 0.22% |
| 200385 vs 200402 | 926 | 5 | 0.54% |
| 200385 vs 200403 | 924 | 4 | 0.43% |
| 200385 vs 200404 | 925 | 5 | 0.54% |
| 200385 vs 200405 | 925 | 6 | 0.65% |
| 200385 vs 200406 | 925 | 4 | 0.43% |
| 200385 vs 200407 | 925 | 4 | 0.43% |
| 200385 vs 200408 | 926 | 6 | 0.65% |
| 200385 vs 201110 | 925 | 3 | 0.32% |
| 200385 vs 201111 | 925 | 3 | 0.32% |
| 200385 vs 201112 | 925 | 4 | 0.43% |
| 200385 vs 201113 | 926 | 2 | 0.22% |
| 200385 vs 201114 | 926 | 4 | 0.43% |
| 200384 vs 200385 | 923 | 8 | 0.87% |
| 200384 vs 200386 | 923 | 4 | 0.43% |
| 200384 vs 200387 | 922 | 5 | 0.54% |
| 200384 vs 200388 | 920 | 4 | 0.43% |
| 200384 vs 200389 | 923 | 6 | 0.65% |
| 200384 vs 200390 | 923 | 6 | 0.65% |
| 200384 vs 200391 | 923 | 5 | 0.54% |
| 200384 vs 200392 | 923 | 10 | 1.08% |
| 200384 vs 200393 | 923 | 6 | 0.65% |
| 200384 vs 200394 | 923 | 5 | 0.54% |
| 200384 vs 200395 | 923 | 6 | 0.65% |
| 200384 vs 200396 | 923 | 5 | 0.54% |
| 200384 vs 200398 | 923 | 9 | 0.98% |
| 200384 vs 200399 | 922 | 5 | 0.54% |
| 200384 vs 200400 | 922 | 6 | 0.65% |
| 200384 vs 200401 | 923 | 6 | 0.65% |
| 200384 vs 200402 | 923 | 7 | 0.76% |
| 200384 vs 200403 | 921 | 4 | 0.43% |
| 200384 vs 200404 | 922 | 6 | 0.65% |
| 200384 vs 200405 | 922 | 7 | 0.76% |
| 200384 vs 200406 | 922 | 6 | 0.65% |
| 200384 vs 200407 | 923 | 8 | 0.87% |
| 200384 vs 200408 | 923 | 8 | 0.87% |
| 200384 vs 201110 | 922 | 5 | 0.54% |
| 200384 vs 201111 | 922 | 4 | 0.43% |
| 200384 vs 201112 | 923 | 6 | 0.65% |
| 200384 vs 201113 | 923 | 6 | 0.65% |
| 200384 vs 201114 | 923 | 6 | 0.65% |
| 200383 vs 200384 | 923 | 47 | 5.09% |
| 200383 vs 200385 | 925 | 117 | 12.65% |
| 200383 vs 200386 | 926 | 42 | 4.54% |
| 200383 vs 200387 | 924 | 116 | 12.55% |
| 200383 vs 200388 | 922 | 114 | 12.36% |
| 200383 vs 200389 | 925 | 116 | 12.54% |
| 200383 vs 200390 | 925 | 117 | 12.65% |
| 200383 vs 200391 | 925 | 115 | 12.43% |
| 200383 vs 200392 | 925 | 118 | 12.76% |
| 200383 vs 200393 | 924 | 116 | 12.55% |
| 200383 vs 200394 | 924 | 114 | 12.34% |
| 200383 vs 200395 | 925 | 116 | 12.54% |
| 200383 vs 200396 | 925 | 115 | 12.43% |
| 200383 vs 200398 | 925 | 118 | 12.76% |
| 200383 vs 200399 | 924 | 116 | 12.55% |
| 200383 vs 200400 | 923 | 114 | 12.35% |
| 200383 vs 200401 | 925 | 116 | 12.54% |
| 200383 vs 200402 | 925 | 115 | 12.43% |
| 200383 vs 200403 | 923 | 115 | 12.46% |
| 200383 vs 200404 | 924 | 114 | 12.34% |
| 200383 vs 200405 | 924 | 116 | 12.55% |
| 200383 vs 200406 | 924 | 115 | 12.45% |
| 200383 vs 200407 | 924 | 114 | 12.34% |

| | | | |
|---|---|---|---|
| 200383 vs 200408 | 925 | 117 | 12.65% |
| 200383 vs 201110 | 924 | 114 | 12.34% |
| 200383 vs 201111 | 924 | 114 | 12.34% |
| 200383 vs 201112 | 925 | 117 | 12.65% |
| 200383 vs 201113 | 925 | 117 | 12.65% |
| 200383 vs 201114 | 925 | 116 | 12.54% |
