## Supplementary table 6 for "Multiple nucleotide polymorphism DNA markers for the accurate evaluation of genetic variations"

**Supplementary Table 6. Rice and cotton varieties used in this study**

| Variety code | Species | Variety name | Ploidy | Hybrid/Inbred | Used in | | | |
|---|---|---|---|---|---|---|---|---|
| | | | | | Reproducibility experiments | SSR genotyping | MNP fingerprint database | Infringement investigation |
| OSD202062 | *Oryza sativa L. Indica* | HLiangyou518 | Diploid | Hybrid | ✔ | ✔ | ✔ | |
| OSD202180 | *Oryza sativa L. Indica* | YLiangyou115 | Diploid | Hybrid | ✔ | ✔ | ✔ | |
| OSD202183 | *Oryza sativa L. Indica* | Chuanyou8377 | Diploid | Hybrid | ✔ | ✔ | ✔ | |
| OSD202200 | *Oryza sativa L. Indica* | Xieliangyou273 | Diploid | Hybrid | ✔ | ✔ | ✔ | |
| OSD202201 | *Oryza sativa L. Indica* | Anyou402 | Diploid | Hybrid | ✔ | ✔ | ✔ | |
| OSD202227 | *Oryza sativa L. Indica* | Chuanyou727 | Diploid | Hybrid | ✔ | ✔ | ✔ | |
| OSD202246 | *Oryza sativa L. Indica* | Fuhui7206 | Diploid | Inbred | ✔ | ✔ | ✔ | |
| OSD202358 | *Oryza sativa L. Indica* | YLiangyou916 | Diploid | Hybrid | ✔ | ✔ | ✔ | |
| OSD202557 | *Oryza sativa L. Japonica* | Yongyou13 | Diploid | Hybrid | ✔ | ✔ | ✔ | |
| OSD202663 | *Oryza sativa L. Japonica* | Liaoyou9906 | Diploid | Hybrid | ✔ | ✔ | ✔ | |
| OSD202674 | *Oryza sativa L. Indica* | Liangyou289 | Diploid | Hybrid | ✔ | ✔ | ✔ | |
| OSD202680 | *Oryza sativa L. Indica* | Wufengyou66 | Diploid | Hybrid | ✔ | ✔ | ✔ | |
| OSD202723 | *Oryza sativa L. Indica* | Yangxianyou68 | Diploid | Hybrid | ✔ | ✔ | ✔ | |
| OSD202745 | *Oryza sativa L. Indica* | Guyoumingzhan | Diploid | Hybrid | ✔ | ✔ | ✔ | |
| OSD202748 | *Oryza sativa L. Indica* | Xiannong7 | Diploid | Hybrid | ✔ | ✔ | ✔ | |
| OSD202751 | *Oryza sativa L. Indica* | IIYou3027 | Diploid | Hybrid | ✔ | ✔ | ✔ | |
| OSD202763 | *Oryza sativa L. Indica* | IIYou3216 | Diploid | Hybrid | ✔ | ✔ | ✔ | |
| OSD202766 | *Oryza sativa L. Indica* | Qianyou2 | Diploid | Hybrid | ✔ | ✔ | ✔ | |
| OSD202767 | *Oryza sativa L. Indica* | Gangyou1577 | Diploid | Hybrid | ✔ | ✔ | ✔ | |
| OSD202768 | *Oryza sativa L. Indica* | Guangyouming118 | Diploid | Hybrid | ✔ | ✔ | ✔ | |
| OSD202781 | *Oryza sativa L. Indica* | Tianyou81 | Diploid | Hybrid | ✔ | ✔ | ✔ | |
| OSD202792 | *Oryza sativa L. Indica* | TYou111 | Diploid | Hybrid | ✔ | ✔ | ✔ | |
| OSD202799 | *Oryza sativa L. Indica* | Fengyou416 | Diploid | Hybrid | ✔ | ✔ | ✔ | |
| OSD202801 | *Oryza sativa L. Indica* | Qyou8 | Diploid | Hybrid | ✔ | ✔ | ✔ | |
| OSD202802 | *Oryza sativa L. Indica* | Chuanfeng6 | Diploid | Hybrid | ✔ | ✔ | ✔ | |
| OSD202809 | *Oryza sativa L. Indica* | Boyoushuangqing | Diploid | Hybrid | ✔ | ✔ | ✔ | |
| OSD202811 | *Oryza sativa L. Indica* | Xiangfengyou402 | Diploid | Hybrid | ✔ | ✔ | ✔ | |
| OSD202815 | *Oryza sativa L. Indica* | IIYou416 | Diploid | Hybrid | ✔ | ✔ | ✔ | |
| OSD202904 | *Oryza sativa L. Japonica* | Longyou1715 | Diploid | Hybrid | ✔ | ✔ | ✔ | |
| OSD202906 | *Oryza sativa L. Indica* | Jinyou928 | Diploid | Hybrid | ✔ | ✔ | ✔ | |
| OSD202912 | *Oryza sativa L. Indica* | Yixiangyou7808 | Diploid | Hybrid | ✔ | ✔ | ✔ | |
| OSD202914 | *Oryza sativa L. Indica* | Fengyouxiangzhan | Diploid | Hybrid | ✔ | ✔ | ✔ | |
| OSD202936 | *Oryza sativa L. Indica* | Fyou498 | Diploid | Hybrid | ✔ | ✔ | ✔ | |
| OSD202938 | *Oryza sativa L. Indica* | YLiangyouNo2 | Diploid | Hybrid | ✔ | ✔ | ✔ | |
| OSD202958 | *Oryza sativa L. Indica* | Chuanyou673 | Diploid | Hybrid | ✔ | ✔ | ✔ | |
| OSD203003 | *Oryza sativa L. Indica* | Shenyou9586 | Diploid | Hybrid | ✔ | ✔ | ✔ | |
| OSD203044 | *Oryza sativa L. Indica* | Liangyou6507 | Diploid | Hybrid | ✔ | ✔ | ✔ | |
| OSD203080 | *Oryza sativa L. Indica* | Hnayou73 | Diploid | Hybrid | ✔ | ✔ | ✔ | |
| OSD203081 | *Oryza sativa L. Indica* | Hnayou113 | Diploid | Hybrid | ✔ | | ✔ | |
| OSD202243 | *Oryza sativa L. Indica* | Liangyou8901 | Diploid | Hybrid | ✔ | | ✔ | |
| OSD202412 | *Oryza sativa L. Indica* | Liangyou6816 | Diploid | Hybrid | ✔ | | ✔ | |
| OSD202440 | *Oryza sativa L. Indica* | DLiangyou71 | Diploid | Hybrid | ✔ | | ✔ | |
| OSD202712 | *Oryza sativa L. Indica* | Huiliangyou1898 | Diploid | Hybrid | ✔ | | ✔ | |
| OSD202237 | *Oryza sativa L. Indica* | Tianlong8You177 | Diploid | Hybrid | ✔ | | ✔ | |
| OSD202740 | *Oryza sativa L. Indica* | Zhenxianyou184 | Diploid | Hybrid | ✔ | | ✔ | |
| OSD203095 | *Oryza sativa L. Indica* | Teyou7166 | Diploid | Hybrid | ✔ | | ✔ | |
| OSD202175 | *Oryza sativa L. Indica* | Yuanliangyou473 | Diploid | Hybrid | ✔ | | ✔ | |
| OSD205106 | *Gossypium spp* | K638 | Tetraploid | Hybrid | ✔ | | | |
| OSD101231 | *Gossypium spp* | sGK958 | Tetraploid | Hybrid | ✔ | | | |
| OSD101235 | *Gossypium spp* | Bianmian6 | Tetraploid | Hybrid | ✔ | | | |
| OSD205097 | *Gossypium spp* | Dangza02-3 | Tetraploid | Hybrid | ✔ | | | |
| OSD101223 | *Gossypium spp* | Dejiachang2 | Tetraploid | Hybrid | ✔ | | | |
| OSD101209 | *Gossypium spp* | Ezamian23F1 | Tetraploid | Hybrid | ✔ | | | |
| OSD101207 | *Gossypium spp* | Fengzaimian2 | Tetraploid | Hybrid | ✔ | | | |
| OSD101210 | *Gossypium spp* | Fumian2 | Tetraploid | Hybrid | ✔ | | | |
| OSD101208 | *Gossypium spp* | Fumian289 | Tetraploid | Hybrid | ✔ | | | |
| OSD101216 | *Gossypium spp* | Fuquan10 | Tetraploid | Hybrid | ✔ | | | |
| OSD205101 | *Gossypium spp* | Jiza708 | Tetraploid | Hybrid | ✔ | | | |
| OSD205098 | *Gossypium spp* | Lu05H9 | Tetraploid | Hybrid | ✔ | | | |
| OSD205110 | *Gossypium spp* | Lumianyan21 | Tetraploid | Hybrid | ✔ | | | |
| OSD205094 | *Gossypium spp* | Lumianyan23 | Tetraploid | Hybrid | ✔ | | | |
| OSD205102 | *Gossypium spp* | Lumianyan24 | Tetraploid | Hybrid | ✔ | | | |
| OSD205103 | *Gossypium spp* | Lumianyan27 | Tetraploid | Hybrid | ✔ | | | |
| OSD205111 | *Gossypium spp* | Lumianyan30 | Tetraploid | Hybrid | ✔ | | | |

| | | | | | | | |
|---|---|---|---|---|---|---|---|
| OSD205107 | *Gossypium spp* | Lumianyan36 | Tetraploid | Hybrid | ✔ | | |
| OSD101213 | *Gossypium spp* | Lvyimian8 | Tetraploid | Hybrid | ✔ | | |
| OSD101201 | *Gossypium spp* | Ningzamian3 | Tetraploid | Hybrid | ✔ | | |
| OSD101233 | *Gossypium spp* | Shannongshengza3 | Tetraploid | Hybrid | ✔ | | |
| OSD205108 | *Gossypium spp* | Tianyunzayihao | Tetraploid | Hybrid | ✔ | | |
| OSD101205 | *Gossypium spp* | Xiangzamian12 | Tetraploid | Hybrid | ✔ | | |
| OSD101202 | *Gossypium spp* | Xiangzamian9 | Tetraploid | Hybrid | ✔ | | |
| OSD205096 | *Gossypium spp* | Xinhai24 | Tetraploid | Hybrid | ✔ | | |
| OSD101225 | *Gossypium spp* | Xinhai25 | Tetraploid | Hybrid | ✔ | | |
| OSD205093 | *Gossypium spp* | Xinluzao24 | Tetraploid | Hybrid | ✔ | | |
| OSD205104 | *Gossypium spp* | Xinluzao25 | Tetraploid | Hybrid | ✔ | | |
| OSD101212 | *Gossypium spp* | Xinluzao26 | Tetraploid | Hybrid | ✔ | | |
| OSD101203 | *Gossypium spp* | Xinluzao33 | Tetraploid | Hybrid | ✔ | | |
| OSD205112 | *Gossypium spp* | Xinluzao35 | Tetraploid | Hybrid | ✔ | | |
| OSD101204 | *Gossypium spp* | Xinluzao36 | Tetraploid | Hybrid | ✔ | | |
| OSD101217 | *Gossypium spp* | Xinluzhong26 | Tetraploid | Hybrid | ✔ | | |
| OSD101232 | *Gossypium spp* | Xinluzhong28 | Tetraploid | Hybrid | ✔ | | |
| OSD101224 | *Gossypium spp* | Xinluzhong32 | Tetraploid | Hybrid | ✔ | | |
| OSD205100 | *Gossypium spp* | Xinluzhong47 | Tetraploid | Hybrid | ✔ | | |
| OSD101211 | *Gossypium spp* | Xinzhi5 | Tetraploid | Hybrid | ✔ | | |
| OSD101198 | *Gossypium spp* | Xumian18 | Tetraploid | Hybrid | ✔ | | |
| OSD205109 | *Gossypium spp* | Yinmian2 | Tetraploid | Hybrid | ✔ | | |
| OSD205095 | *Gossypium spp* | Yinrui361 | Tetraploid | Hybrid | ✔ | | |
| OSD205105 | *Gossypium spp* | Zhongmiansuo64 | Tetraploid | Hybrid | ✔ | | |
| OSD205099 | *Gossypium spp* | Zhongmiansuo66 | Tetraploid | Hybrid | | | |
| OSD101214 | *Gossypium spp* | Zhongzhimian2 | Tetraploid | Hybrid | | | |
| OSD30084 | *Oryza sativa L. Japonica* | Wuyoudao4 | Diploid | Inbred | | | ✔ |
| OSD200322 | *Oryza sativa L. Indica* | Ezhong5 | Diploid | Inbred | | | ✔ |
| OSD30336 | *Oryza sativa L. Indica* | Huanghuazhan | Diploid | Inbred | | | ✔ |
| OSD9013 | *Oryza sativa L. Indica* | Tianliangyou16 | Diploid | hybrid | | | ✔ |
| OSD6048 | *Oryza sativa L. Japonica* | Ejingza3 | Diploid | hybrid | | | ✔ |
| OSD8218 | *Oryza sativa L. Japonica* | DianZa35 | Diploid | hybrid | | | ✔ |
| OSD8207 | *Oryza sativa L. Japonica* | Chunyou172 | Diploid | hybrid | | | ✔ |
| OSD8205 | *Oryza sativa L. Japonica* | Chunyou59 | Diploid | hybrid | | | ✔ |
| OSD7294 | *Oryza sativa L. Japonica* | Chunyou58 | Diploid | hybrid | | | ✔ |
| OSD6402 | *Oryza sativa L. Japonica* | Shenyou8 | Diploid | hybrid | | | ✔ |
| OSD4760 | *Oryza sativa L. Japonica* | Jinyou2003 | Diploid | hybrid | | | ✔ |
| OSD4369 | *Oryza sativa L. Japonica* | DianZa32 | Diploid | hybrid | | | ✔ |
| OSD1756 | *Oryza sativa L. Japonica* | Shenyou4 | Diploid | hybrid | | | ✔ |
| OSD7782 | *Oryza sativa L. Japonica* | QiuYouJinFeng | Diploid | hybrid | | | ✔ |
| OSD6314 | *Oryza sativa L. Japonica* | Yongyou6 | Diploid | hybrid | | | ✔ |
| OSD5446 | *Oryza sativa L. Japonica* | Sujingyou3 | Diploid | hybrid | | | ✔ |
| OSD8202 | *Oryza sativa L. Indica* | Guangliangyou100 | Diploid | hybrid | | | ✔ |
| OSD7843 | *Oryza sativa L. Indica* | Yliangyou599 | Diploid | hybrid | | | ✔ |
| OSD6392 | *Oryza sativa L. Indica* | Liangyouhang2 | Diploid | hybrid | | | ✔ |
| OSD6239 | *Oryza sativa L. Indica* | Cliangyou396 | Diploid | hybrid | | | ✔ |
| OSD6046 | *Oryza sativa L. Indica* | Zhutwoyou30 | Diploid | hybrid | | | ✔ |
| OSD5919 | *Oryza sativa L. Indica* | Zhutwoyou706 | Diploid | hybrid | | | ✔ |
| OSD5117 | *Oryza sativa L. Indica* | Liangyou108 | Diploid | hybrid | | | ✔ |
| OSD5059 | *Oryza sativa L. Indica* | Zhuliangyou124 | Diploid | hybrid | | | ✔ |
| OSD4341 | *Oryza sativa L. Indica* | Zhuliangyou99 | Diploid | hybrid | | | ✔ |
| OSD3847 | *Oryza sativa L. Indica* | liangyou276 | Diploid | hybrid | | | ✔ |
| OSD3301 | *Oryza sativa L. Indica* | LuLiangyou996 | Diploid | hybrid | | | ✔ |
| OSD2030 | *Oryza sativa L. Indica* | Peiliangyou500 | Diploid | hybrid | | | ✔ |
| OSD1893 | *Oryza sativa L. Indica* | liangyou932 | Diploid | hybrid | | | ✔ |
| OSD1140 | *Oryza sativa L. Indica* | YangLiangyou6 | Diploid | hybrid | | | ✔ |
| OSD8148 | *Oryza sativa L. Indica* | YiSwan2 | Diploid | hybrid | | | ✔ |
| OSD8059 | *Oryza sativa L. Indica* | HuiliangyouNo.6 | Diploid | hybrid | | | ✔ |
| OSD7674 | *Oryza sativa L. Indica* | ZhunLiangyou2 | Diploid | hybrid | | | ✔ |
| OSD6383 | *Oryza sativa L. Indica* | Twoyou0293 | Diploid | hybrid | | | ✔ |
| OSD5846 | *Oryza sativa L. Indica* | Newtwoexcellent6380 | Diploid | hybrid | | | ✔ |
| OSD2307 | *Oryza sativa L. Indica* | Anliangyouqingzhan | Diploid | hybrid | | | ✔ |
| OSD1230 | *Oryza sativa L. Indica* | TwoyouE32 | Diploid | hybrid | | | ✔ |
| OSD8600 | *Oryza sativa L. Indica* | Hanyou3hao | Diploid | hybrid | | | ✔ |
| OSD890 | *Oryza sativa L. Indica* | Gangyou3551 | Diploid | hybrid | | | ✔ |
| OSD8371 | *Oryza sativa L. Indica* | BoIIyou859 | Diploid | hybrid | | | ✔ |
| OSD8316 | *Oryza sativa L. Indica* | WenFu7 | Diploid | hybrid | | | ✔ |
| OSD8250 | *Oryza sativa L. Indica* | XianNong2058 | Diploid | hybrid | | | ✔ |
| OSD8241 | *Oryza sativa L. Indica* | Nonghuayou5365 | Diploid | hybrid | | | ✔ |

| | | | | | | | |
|---|---|---|---|---|---|---|---|
| OSD7865 | *Oryza sativa L. Indica* | Yuyou600 | Diploid | hybrid | | | ✔ |
| OSD7806 | *Oryza sativa L. Indica* | Shenyou9734 | Diploid | hybrid | | | ✔ |
| OSD7662 | *Oryza sativa L. Indica* | Yixiang2239 | Diploid | hybrid | | | ✔ |
| OSD7658 | *Oryza sativa L. Indica* | Yixiang2084 | Diploid | hybrid | | | ✔ |
| OSD7650 | *Oryza sativa L. Indica* | Boyou768 | Diploid | hybrid | | | ✔ |
| OSD7619 | *Oryza sativa L. Indica* | Xinongyou2 | Diploid | hybrid | | | ✔ |
| OSD7612 | *Oryza sativa L. Indica* | XiNongyou30 | Diploid | hybrid | | | ✔ |
| OSD7532 | *Oryza sativa L. Indica* | Dyou158 | Diploid | hybrid | | | ✔ |
| OSD7410 | *Oryza sativa L. Indica* | Jinyou167 | Diploid | hybrid | | | ✔ |
| OSD7377 | *Oryza sativa L. Indica* | Nyou69 | Diploid | hybrid | | | ✔ |
| OSD7372 | *Oryza sativa L. Indica* | NeiXiang8156 | Diploid | hybrid | | | ✔ |
| OSD7365 | *Oryza sativa L. Indica* | NeiXiang10 | Diploid | hybrid | | | ✔ |
| OSD7363 | *Oryza sativa L. Indica* | NeiXiang2128 | Diploid | hybrid | | | ✔ |
| OSD6869 | *Oryza sativa L. Indica* | Liuyou105 | Diploid | hybrid | | | ✔ |
| OSD685 | *Oryza sativa L. Indica* | Dyou13 | Diploid | hybrid | | | ✔ |
| OSD6826 | *Oryza sativa L. Indica* | IIyou1069 | Diploid | hybrid | | | ✔ |
| OSD6512 | *Oryza sativa L. Indica* | Fenghuayou1 | Diploid | hybrid | | | ✔ |
| OSD6482 | *Oryza sativa L. Indica* | Zhong9you3190 | Diploid | hybrid | | | ✔ |
| OSD6473 | *Oryza sativa L. Indica* | Fengyoudazhan | Diploid | hybrid | | | ✔ |
| OSD6471 | *Oryza sativa L. Indica* | Fenghuayou2 | Diploid | hybrid | | | ✔ |
| OSD6394 | *Oryza sativa L. Indica* | IIyouhang2 | Diploid | hybrid | | | ✔ |
| OSD6387 | *Oryza sativa L. Indica* | Tyou618 | Diploid | hybrid | | | ✔ |
| OSD6385 | *Oryza sativa L. Indica* | Tyou608 | Diploid | hybrid | | | ✔ |
| OSD6229 | *Oryza sativa L. Indica* | Zijindao6 | Diploid | hybrid | | | ✔ |
| OSD5791 | *Oryza sativa L. Indica* | Jinyou898 | Diploid | hybrid | | | ✔ |
| OSD5749 | *Oryza sativa L. Indica* | IIyou640 | Diploid | hybrid | | | ✔ |
| OSD5460 | *Oryza sativa L. Indica* | Feiyou600 | Diploid | hybrid | | | ✔ |
| OSD5242 | *Oryza sativa L. Indica* | Jinyou188 | Diploid | hybrid | | | ✔ |
| OSD5028 | *Oryza sativa L. Indica* | Huayou2 | Diploid | hybrid | | | ✔ |
| OSD4843 | *Oryza sativa L. Indica* | YangxinyouNo.1 | Diploid | hybrid | | | ✔ |
| OSD4738 | *Oryza sativa L. Indica* | Yixiang3728 | Diploid | hybrid | | | ✔ |
| OSD4728 | *Oryza sativa L. Indica* | Shanyou161 | Diploid | hybrid | | | ✔ |
| OSD4628 | *Oryza sativa L. Indica* | IIyou107 | Diploid | hybrid | | | ✔ |
| OSD4564 | *Oryza sativa L. Indica* | Guyou3119 | Diploid | hybrid | | | ✔ |
| OSD4560 | *Oryza sativa L. Indica* | IIyou3169 | Diploid | hybrid | | | ✔ |
| OSD4559 | *Oryza sativa L. Indica* | IIyou3139 | Diploid | hybrid | | | ✔ |
| OSD4507 | *Oryza sativa L. Indica* | TeYou627 | Diploid | hybrid | | | ✔ |
| OSD4435 | *Oryza sativa L. Indica* | ThreeXiangyou714 | Diploid | hybrid | | | ✔ |
| OSD4386 | *Oryza sativa L. Indica* | IIyou550 | Diploid | hybrid | | | ✔ |
| OSD4384 | *Oryza sativa L. Indica* | Boyou781 | Diploid | hybrid | | | ✔ |
| OSD4371 | *Oryza sativa L. Indica* | Jinyou540 | Diploid | hybrid | | | ✔ |
| OSD4366 | *Oryza sativa L. Indica* | Zhongyou177 | Diploid | hybrid | | | ✔ |
| OSD4281 | *Oryza sativa L. Indica* | GanXin6 | Diploid | hybrid | | | ✔ |
| OSD4253 | *Oryza sativa L. Indica* | Xieyou332 | Diploid | hybrid | | | ✔ |
| OSD4248 | *Oryza sativa L. Indica* | Xieyou336 | Diploid | hybrid | | | ✔ |
| OSD4085 | *Oryza sativa L. Indica* | Fengyuanyou6135 | Diploid | hybrid | | | ✔ |
| OSD3865 | *Oryza sativa L. Indica* | XianNong4 | Diploid | hybrid | | | ✔ |
| OSD3810 | *Oryza sativa L. Indica* | Fuyou4 | Diploid | hybrid | | | ✔ |
| OSD3676 | *Oryza sativa L. Indica* | IIyou131 | Diploid | hybrid | | | ✔ |
| OSD3628 | *Oryza sativa L. Indica* | Jinyou217 | Diploid | hybrid | | | ✔ |
| OSD3435 | *Oryza sativa L. Indica* | Yixiang19 | Diploid | hybrid | | | ✔ |
| OSD3416 | *Oryza sativa L. Indica* | Gangyou7954 | Diploid | hybrid | | | ✔ |
| OSD2877 | *Oryza sativa L. Indica* | Xieyou218 | Diploid | hybrid | | | ✔ |
| OSD2841 | *Oryza sativa L. Indica* | Jinyou527 | Diploid | hybrid | | | ✔ |
| OSD2812 | *Oryza sativa L. Indica* | Chuan7you89 | Diploid | hybrid | | | ✔ |
| OSD2477 | *Oryza sativa L. Indica* | Neixiangyou9 | Diploid | hybrid | | | ✔ |
| OSD2260 | *Oryza sativa L. Indica* | TeYou898 | Diploid | hybrid | | | ✔ |
| OSD2253 | *Oryza sativa L. Indica* | JinYouming100 | Diploid | hybrid | | | ✔ |
| OSD2036 | *Oryza sativa L. Indica* | Zhongyou63 | Diploid | hybrid | | | ✔ |
| OSD1995 | *Oryza sativa L. Indica* | Xieyou527 | Diploid | hybrid | | | ✔ |
| OSD1901 | *Oryza sativa L. Indica* | Kyou66 | Diploid | hybrid | | | ✔ |
| OSD1876 | *Oryza sativa L. Indica* | Jinyou706 | Diploid | hybrid | | | ✔ |
| OSD1841 | *Oryza sativa L. Indica* | Gangyou19 | Diploid | hybrid | | | ✔ |
| OSD1738 | *Oryza sativa L. Indica* | XianNong5 | Diploid | hybrid | | | ✔ |
| OSD1615 | *Oryza sativa L. Indica* | Jinyou299 | Diploid | hybrid | | | ✔ |
| OSD1578 | *Oryza sativa L. Indica* | Yixiang2308 | Diploid | hybrid | | | ✔ |
| OSD1574 | *Oryza sativa L. Indica* | Yixiang11 | Diploid | hybrid | | | ✔ |
| OSD1412 | *Oryza sativa L. Indica* | Jinyou718 | Diploid | hybrid | | | ✔ |
| OSD1408 | *Oryza sativa L. Indica* | Qyou1 | Diploid | hybrid | | | ✔ |

| | | | | | | | | |
|---|---|---|---|---|---|---|---|---|
| OSD1343 | *Oryza sativa L. Indica* | Chuanxiangyou2 | Diploid | hybrid | | | ✔ | |
| OSD1116 | *Oryza sativa L. Indica* | Yueyou938 | Diploid | hybrid | | | ✔ | |
| OSD10524 | *Oryza sativa L. Indica* | Fuyou151 | Diploid | hybrid | | | ✔ | |
| OSD1959 | *Oryza sativa L. Indica* | Jinyou182 | Diploid | hybrid | | | ✔ | |
| OSD9010 | *Oryza sativa L. Indica* | Yixiang99E-4 | Diploid | hybrid | | | ✔ | |
| OSD889 | *Oryza sativa L. Indica* | Yixiangyou1577 | Diploid | hybrid | | | ✔ | |
| OSD6662 | *Oryza sativa L. Indica* | Yixiang481 | Diploid | hybrid | | | ✔ | |
| OSD6489 | *Oryza sativa L. Indica* | GuohaoGuoxiang12 | Diploid | hybrid | | | ✔ | |
| OSD5804 | *Oryza sativa L. Indica* | TeYou399 | Diploid | hybrid | | | ✔ | |
| OSD5070 | *Oryza sativa L. Indica* | IIyouhang148 | Diploid | hybrid | | | ✔ | |
| OSD4756 | *Oryza sativa L. Indica* | GuoDao1 | Diploid | hybrid | | | ✔ | |
| OSD4740 | *Oryza sativa L. Indica* | Mian5you838 | Diploid | hybrid | | | ✔ | |
| OSD4519 | *Oryza sativa L. Indica* | Xieyou085 | Diploid | hybrid | | | ✔ | |
| OSD4501 | *Oryza sativa L. Indica* | Shennongdafengrice10 | Diploid | hybrid | | | ✔ | |
| OSD4416 | *Oryza sativa L. Indica* | Jinyou2155 | Diploid | hybrid | | | ✔ | |
| OSD4339 | *Oryza sativa L. Indica* | Yuxiangyou164 | Diploid | hybrid | | | ✔ | |
| OSD3808 | *Oryza sativa L. Indica* | Gangyou825 | Diploid | hybrid | | | ✔ | |
| OSD3232 | *Oryza sativa L. Indica* | TianYou218 | Diploid | hybrid | | | ✔ | |
| OSD2484 | *Oryza sativa L. Indica* | Neixiangyou1 | Diploid | hybrid | | | ✔ | |
| OSD2407 | *Oryza sativa L. Indica* | Mian2you151 | Diploid | hybrid | | | ✔ | |
| OSD2293 | *Oryza sativa L. Indica* | Jingchuyou148 | Diploid | hybrid | | | ✔ | |
| OSD2107 | *Oryza sativa L. Indica* | IIyou87 | Diploid | hybrid | | | ✔ | |
| OSD1875 | *Oryza sativa L. Indica* | Tyou706 | Diploid | hybrid | | | ✔ | |
| OSD1571 | *Oryza sativa L. Indica* | Yixiang2292 | Diploid | hybrid | | | ✔ | |
| OSD1371 | *Oryza sativa L. Indica* | Wandao81 | Diploid | hybrid | | | ✔ | |
| OSD112 | *Oryza sativa L. Indica* | IIyou98 | Diploid | hybrid | | | ✔ | |
| OSD5267 | *Oryza sativa L. Indica* | NongHuaYou808 | Diploid | hybrid | | | ✔ | |
| OSD4249 | *Oryza sativa L. Indica* | YangXianYou22 | Diploid | hybrid | | | ✔ | |
| OSD3819 | *Oryza sativa L. Indica* | F3027 | Diploid | hybrid | | | ✔ | |
| OSD3534 | *Oryza sativa L. Indica* | Hongmiao1 | Diploid | hybrid | | | ✔ | |
| OSD2488 | *Oryza sativa L. Indica* | Neixiangyou13 | Diploid | hybrid | | | ✔ | |
| OSD2406 | *Oryza sativa L. Indica* | IIyou085 | Diploid | hybrid | | | ✔ | |
| OSD2237 | *Oryza sativa L. Indica* | Jinyou9059 | Diploid | hybrid | | | ✔ | |
| OSD1823 | *Oryza sativa L. Indica* | Fengyuan299 | Diploid | hybrid | | | ✔ | |
| OSD1592 | *Oryza sativa L. Indica* | Ningteyou420 | Diploid | hybrid | | | ✔ | |
| OSD1545 | *Oryza sativa L. Indica* | Yixiangyou725 | Diploid | hybrid | | | ✔ | |
| OSD1378 | *Oryza sativa L. Indica* | Zhongyou752 | Diploid | hybrid | | | ✔ | |
| OSD1089 | *Oryza sativa L. Indica* | GuofengNo.1 | Diploid | hybrid | | | ✔ | |
| market1 | *Oryza sativa L.* | Tianliangyou16 | Diploid | / | | | | ✔ |
| market2 | *Oryza sativa L.* | Taifengyou3301 | Diploid | / | | | | ✔ |
| market3 | *Oryza sativa L.* | NeiXiang2128 | Diploid | / | | | | ✔ |
| market4 | *Oryza sativa L.* | Taiyou390 | Diploid | / | | | | ✔ |
| market5 | *Oryza sativa L.* | Meixiangzhan2 | Diploid | / | | | | ✔ |
| market6 | *Oryza sativa L.* | Yliangyou800 | Diploid | / | | | | ✔ |
| market7 | *Oryza sativa L.* | Anliangyou166 | Diploid | / | | | | ✔ |
| market8 | *Oryza sativa L.* | HuangHuaZhan | Diploid | / | | | | ✔ |
| market9 | *Oryza sativa L.* | Fenghuazhan | Diploid | / | | | | ✔ |
| market10 | *Oryza sativa L.* | GuyouMingzhan | Diploid | / | | | | ✔ |
| market11 | *Oryza sativa L.* | HuiLiangYousiMiao | Diploid | / | | | | ✔ |
| market12 | *Oryza sativa L.* | liangyou602 | Diploid | / | | | | ✔ |
| market13 | *Oryza sativa L.* | Quanliangyou1 | Diploid | / | | | | ✔ |
| market14 | *Oryza sativa L.* | TianlongLiangyou140 | Diploid | / | | | | ✔ |
| market15 | *Oryza sativa L.* | Wuyudao4 | Diploid | / | | | | ✔ |
| market16 | *Oryza sativa L.* | Ezhong5 | Diploid | / | | | | ✔ |
| market17 | *Oryza sativa L.* | Cliangyou396 | Diploid | / | | | | ✔ |
| market18 | *Oryza sativa L.* | LongJing21 | Diploid | / | | | | ✔ |
| market19 | *Oryza sativa L.* | Yanquanyou1393 | Diploid | / | | | | ✔ |
| market20 | *Oryza sativa L.* | Jingliangyou2847 | Diploid | / | | | | ✔ |
| market21 | *Oryza sativa L.* | Cliangyouhuazhan | Diploid | / | | | | ✔ |
| market22 | *Oryza sativa L.* | Fengliangyouwansan | Diploid | / | | | | ✔ |
| market23 | *Oryza sativa L.* | Lingliangyou211 | Diploid | / | | | | ✔ |
| market24 | *Oryza sativa L.* | liangyou234 | Diploid | / | | | | ✔ |
| market25 | *Oryza sativa L.* | Yliangyou102 | Diploid | / | | | | ✔ |
| market26 | *Oryza sativa L.* | Fengyuanyouhuazhan | Diploid | / | | | | ✔ |
| market27 | *Oryza sativa L.* | ZaofengYouhuazhuan | Diploid | / | | | | ✔ |
| market28 | *Oryza sativa L.* | Yixiangyou2115 | Diploid | / | | | | ✔ |
| market29 | *Oryza sativa L.* | Gangyou3165 | Diploid | / | | | | ✔ |
