## Supplementary table 7 for "Multiple nucleotide polymorphism DNA markers for the accurate evaluation of genetic variations"

### Supplementary Table 7. Identity determination of blind samples

| Blind sample code | Compared to the correct identity of the blind variety | | | | Other varieties with <1% distinct MNP genotypes compared to the blind variety | Species |
|---|---|---|---|---|---|---|
| | Variey code | Number of MNP genotypes compared | Number of distinct MNP genotypes | Ratio of distinct MNP genotypes | | |
| OSD202062-2-2 | OSD202062-3-1 | 928 | 0 | 0 | None | Rice |
| OSD202175-2-2 | OSD202175-3-1 | 926 | 0 | 0 | None | Rice |
| OSD202180-2-2 | OSD202180-3-1 | 914 | 0 | 0 | None | Rice |
| OSD202183-2-2 | OSD202183-3-1 | 810 | 0 | 0 | None | Rice |
| OSD202200-2-2 | OSD202200-3-1 | 928 | 0 | 0 | None | Rice |
| OSD202201-2-2 | OSD202201-3-1 | 929 | 0 | 0 | None | Rice |
| OSD202227-2-2 | OSD202227-3-1 | 929 | 0 | 0 | None | Rice |
| OSD202237-2-2 | OSD202237-3-1 | 930 | 0 | 0 | None | Rice |
| OSD202243-2-2 | OSD202243-3-1 | 930 | 0 | 0 | None | Rice |
| OSD202246-2-2 | OSD202246-3-1 | 927 | 0 | 0 | None | Rice |
| OSD202358-2-2 | OSD202358-3-1 | 929 | 0 | 0 | None | Rice |
| OSD202412-2-2 | OSD202412-3-1 | 929 | 0 | 0 | None | Rice |
| OSD202440-2-2 | OSD202440-3-1 | 928 | 0 | 0 | None | Rice |
| OSD202557-2-2 | OSD202557-3-1 | 929 | 0 | 0 | None | Rice |
| OSD202663-2-2 | OSD202663-3-1 | 927 | 0 | 0 | None | Rice |
| OSD202674-2-2 | OSD202674-3-1 | 799 | 0 | 0 | None | Rice |
| OSD202680-2-2 | OSD202680-3-1 | 926 | 0 | 0 | None | Rice |
| OSD202712-2-2 | OSD202712-3-1 | 929 | 0 | 0 | None | Rice |
| OSD202723-2-2 | OSD202723-3-1 | 927 | 0 | 0 | None | Rice |
| OSD202740-2-2 | OSD202740-3-1 | 928 | 0 | 0 | None | Rice |
| OSD202745-2-2 | OSD202745-3-1 | 928 | 0 | 0 | None | Rice |
| OSD202748-2-2 | OSD202748-3-1 | 928 | 0 | 0 | None | Rice |
| OSD202751-2-2 | OSD202751-3-1 | 928 | 0 | 0 | None | Rice |
| OSD202763-2-2 | OSD202763-3-1 | 927 | 0 | 0 | None | Rice |
| OSD202766-2-2 | OSD202766-3-1 | 929 | 0 | 0 | None | Rice |
| OSD202767-2-2 | OSD202767-3-1 | 929 | 0 | 0 | None | Rice |
| OSD202768-2-2 | OSD202768-3-1 | 929 | 0 | 0 | None | Rice |
| OSD202781-2-2 | OSD202781-3-1 | 927 | 0 | 0 | None | Rice |
| OSD202792-2-2 | OSD202792-3-1 | 928 | 0 | 0 | None | Rice |
| OSD202799-2-2 | OSD202799-3-1 | 929 | 0 | 0 | None | Rice |
| OSD202801-2-2 | OSD202801-3-1 | 930 | 0 | 0 | None | Rice |
| OSD202802-2-2 | OSD202802-3-1 | 929 | 0 | 0 | None | Rice |
| OSD202809-2-2 | OSD202809-3-1 | 929 | 0 | 0 | None | Rice |
| OSD202811-2-2 | OSD202811-3-1 | 929 | 0 | 0 | None | Rice |
| OSD202815-2-2 | OSD202815-3-1 | 930 | 0 | 0 | None | Rice |
| OSD202904-2-2 | OSD202904-3-1 | 930 | 0 | 0 | None | Rice |
| OSD202906-2-2 | OSD202906-3-1 | 926 | 0 | 0 | None | Rice |
| OSD202912-2-2 | OSD202912-3-1 | 929 | 0 | 0 | None | Rice |
| OSD202914-2-2 | OSD202914-3-1 | 929 | 0 | 0 | None | Rice |
| OSD202936-2-2 | OSD202936-3-1 | 929 | 0 | 0 | None | Rice |
| OSD202938-2-2 | OSD202938-3-1 | 930 | 0 | 0 | None | Rice |
| OSD202958-2-2 | OSD202958-3-1 | 929 | 1 | 0.11% | None | Rice |
| OSD203003-2-2 | OSD203003-3-1 | 929 | 0 | 0 | None | Rice |
| OSD203044-2-2 | OSD203044-3-1 | 927 | 0 | 0 | None | Rice |
| OSD203080-2-2 | OSD203080-3-1 | 928 | 0 | 0 | None | Rice |
| OSD203081-2-2 | OSD203081-3-1 | 928 | 0 | 0 | None | Rice |
| OSD203095-2-2 | OSD203095-3-1 | 821 | 0 | 0 | None | Rice |
