## Supplementary table 8 for "Multiple nucleotide polymorphism DNA markers for the accurate evaluation of genetic variations"

### Supplementary Table 8. Infringed varieties

| Market varieties compared | | | | | | | Standard varieties | | | | |
|---|---|---|---|---|---|---|---|---|---|---|---|
| Code of market variety 1 | Labbled name of market variety 1 | Code of market variety 2 | Labbled name of market variety 2 | Common genotypes between market variety 1 and market variety 2 | Number of distinct genotypes between market variety 1 and market variety 2 | Ratio of distinct genotypes between market variety 1 and market variety 2 | Code of standard variety | Name of standard variety | Common genotypes between market variety 1 and its standard variety | Number of distinct genotypes between market variety 1 and its standard variety | Ratio of distinct genotypes between market variety 1 and its standard variety |
| market1 | Tianliangyou16 | market8 | Huanghuanzhan | 873 | 3 | 0.34% | OSD9013 | Tianliangyou16 | 815 | 436 | 53.50% |
| market3 | Neixiang2128 | market15 | Wuyoudao4 | 915 | 6 | 0.66% | OSD7363 | Neixiang2128 | 863 | 761 | 88.18% |
| market17 | Cliangyou396 | market16 | Ezhong5 | 890 | 5 | 0.56% | OSD6239 | Cliangyou396 | 857 | 467 | 54.49% |
