## Supplementary table 9 for "Multiple nucleotide polymorphism DNA markers for the accurate evaluation of genetic variations"

**Supplementary Table 9. Multiplex PCR primers used for SSR genotyping in rice varieties**

| Locus code | Forward primer | Reverse primer |
|---|---|---|
| AMPL1563294 | CTTCGTGGCCGAAAAACACT | CCGGCATGACTGAAATCATGC |
| AMPL1563319 | TTTCTCTAAAACACAAGCCGAGAAAAATC | GATTGGTCTAAGTAATTGGTCCACCAAT |
| AMPL1563340 | CAGATTCTGGCCCAAAACATGTG | GAGAGAGAAAGATTGCTCTCTTCTTGG |
| AMPL1563485 | TCAAGCTGATCATATCATCTTTACTCGTTT | TCAAATTTGCCTGTGATGTAATAGCAAATG |
| AMPL1563607 | GTTGTGATTTTCTGTGCACGAACT | TCTTCTTCTTCTCGGGACAAGATGA |
| AMPL1563649 | ACTCCATTGCTTAGCTCTTAACATTATAGC | TCACTGAGATACGCCAACGAGATA |
| AMPL1563754 | TCAGTGAGGCGATCGAGGAT | ATGCCGCTGGCGAGAAGT |
| AMPL1563873 | GCAATACACCTTATGTGTTGTGCA | GGAAGCAAATAGATCTAATTTACCGGTCTA |
| AMPL1563911 | CCGGCTGATTATACAAGTACGAGTAG | GCGAGGAAGAACTTCTTAGATTAGTGC |
| AMPL1563906 | CAAAGCAGTGCAAAGAACATTTCTCT | GTGTTTATCTACCTCCTTTCTAGACCGATA |
| AMPL1563905 | TTCAGTGTCAACTTTTGCAGGGTA | CATCTTCTTCTACTGCACATGAGCA |
| AMPL1563903 | GCATCCATCCATCCATCATAATGC | GTACAAATCACTTCTCCAATTCCTTACCA |
| AMPL1563901 | ACCTCCAAGGTCCTCATCCT | CTACTCTACTCAAGAAACATCAGGCAAT |
| AMPL1563897 | GATTCTCCTCGCTCATCTGAGTG | GGGATTCCAAGTCGATGACCAAT |
| AMPL1563896 | TTGAAGTCGGTGGCGTGT | AGCGGATGCAGGAGCTCTC |
| AMPL1563893 | GAAAATGTATGGCGAGACCCTACA | TGTGTCACAAGGCTAATCCTTGTC |
| AMPL1563891 | AGGCACCACCCTCCTCTTTA | CCCGGAGATGGAGATCCACA |
| AMPL1563890 | TTGGCCAATCAAATCCAAAGAACC | CCAAGAAGAGTGATCAAGAATTGGGTAAAA |
| AMPL1563884 | TAACTATTATGAGCCAAACACAGCATCA | TGGTTTACGCACACAATGGGATAG |
| AMPL1563883 | GGAGTACGTGACATGGAAATAACGA | CTAATCTAATGGTCTAGAATAGTGGGTCCA |
| AMPL1563880 | CCAATGCAGTGGAACACTGAAA | TAGACTTGGAAGTGTCTCCTCCAA |
| AMPL1563879 | CCCGATGAAGTCGAACAGC | GCGTCCTCGTACAAGGAGC |
| AMPL1563877 | TGGCCCACGAAGTTAACTAATTAAACAT | GGAGCTCAGATATTTAGCACCTCATATTTT |
| AMPL1563876 | CTAATAGCAATTGTAGGAGCGCCATA | CCTTGTGGTCATGCTTCCTCATC |
| AMPL1563872 | CTAATTACTAGCTATGCCTATATGGCCAT | TTTGGTAGGTCGCATTGTTTGGATA |
| AMPL1563868 | ATGTTACGTCAACTATAGGTGTGTAGATAAAG | GTAGATTTTGAGTTATCTTCGTAAACTAGTTGG |
| AMPL1563867 | GAGAAGATGGCAACGGAGAATCT | CCGCCATTAATTGGTACTACTAGTCT |
| AMPL1563866 | CTCCGATGCCTTCTTCCTCTTG | CCTGCGACCCTTCCACTACTATA |
| AMPL1563856 | CGTTGGGAGAGGAGATGTTTCC | GCGTCCTACTGCTACTTCTTCCT |
| AMPL1563854 | ACCGACCGATCTGGGATAGAAA | CTCTTCCTCCTCTCCCTAGCAT |
| AMPL1563853 | TTCTTACATCTGACAGCTGTTTTGGT | TGATCATGTTTACCAAAAACAACAACTACA |
| AMPL1563852 | TCCTCGATCGTCAGGAGATTTTTG | TCTGAACAATTAGCTGCTAGTCCATG |
| AMPL1563850 | CACTGTGTGAACTGCTGAACAG | GTGTTGCAGGTAAAGCATGCAA |
| AMPL1563846 | GGGCTTCTTCTCCTTCTCTCCA | CACCAAATATGCAAATACCACCTTTTGTAT |
| AMPL1563844 | CATTGGTTTTGCCTCGCTGAAT | GTGTGTATCCCACGAAGGAACAA |
| AMPL1563834 | CACCGACATGTACTACTGCTCA | GGGAGTAGGGTCCCATTGGATAG |
| AMPL1563830 | GGGATGTGTGCATGCAATTCAT | CACGGCGATTTCTCTGGGAATTA |
| AMPL1563829 | CTGACACACGCGCATGGAGAAT | GAGATACATAGCAGCACCGGAAA |
| AMPL1563826 | GGCGGTTTAGAAGCGTTTATATGAG | GTGGTTACAAGAGTCTAAGTATCACTTGAG |
| AMPL1563825 | TCTTCCGAAGGCTTCGAACTTT | TCGTGGTCTCTTCTTCTTCATCCATA |
| AMPL1563823 | CAGATCCGGCAATGGTGAAGAT | GGTATCCTCTAATCTCACCGATCAAATG |
| AMPL1563822 | CTTATTTATTAGGATGCCATTTGGCAACTT | CATCCTGCTAATCACTCCTGATCAATC |
| AMPL1563821 | ATATACGTCTTGGTCAGCCAAGTTTATC | GATTCCACTGAACATTCAAACACATAACAT |
| AMPL1563820 | AACAGTGGTCGCAATTGTTTCAC | CGGAGGTTGTGGGAGATGATTTC |
| AMPL1563816 | TGTTGCTATTTGCCATACATCTCCT | AGTCAACAAGTAAAAAGAGAAATGGTGAGA |
| AMPL1563813 | ACAAGATAGCATGCAGATAGCATAACAT | GGGATATGGTGTAACGTTGTTTTATTGAAA |
| AMPL1563812 | GGAGCTAAGTCCTTGCCATCG | GAAAATTGCGAAATAGTACTGTACTGGG |
| AMPL1563810 | GTGGGCCAAGCTGCACAAG | TGGTGGTCGCTAGCTGAC |
| AMPL1563809 | GCCATAGCACAACTCTCCCA | AGTATGGTTATTCATAAAGAGGCTACATGAC |
| AMPL1563807 | AAACTGACATACAAGGCGATAAAACAATG | TGAGGTCGGTCTAGGGTTTTAGTATAATT |
| AMPL1563806 | CATGCAATTACTTGTTTTGGTGGTGT | AAAAGGAACATGTCAACAATCTAGCAAG |
| AMPL1563804 | CCGCCTTCTTCTTCTTCTCTATCTT | GGGCTTTACCTGGTTTTGACCA |
| AMPL1563803 | TTAACAAGTAGTAACGGGTGCTTACTTTT | GGAGTAGCAAAATTATCTGGGATCCAT |
| AMPL1563801 | ATGCATTGATGCATGTGGAACAG | TGATGCTCTTTAACTGGAAAGTCGAAA |
| AMPL1563796 | GGCCGTAGACCTTCTTGAAGTA | CACTGCAATTTGGGCCAATTGA |
| AMPL1563795 | GCAACGCACTCTCTATCTCGTA | GATGAGGAATCGAGGGCAGAAAT |
| AMPL1563794 | GGGTATGGTAGGAAAGGAATAATTGAGA | GGGATGCATGACATGTAGCGAT |
| AMPL1563791 | GGCCCTAATAAGCTAAGCTATACCAC | CGCGCTTCAAATATTCTCGTGT |
| AMPL1563789 | CATCAATGGTGGGAGCTCATGA | CTGGTTGAATTTCAAGATCCAAAAATTGTC |
| AMPL1563787 | ATCCTTCCTCTTCTCCTCTATCATAGAAAC | AAATCTGGCAACCACTTGCATG |
| AMPL1563781 | TATGGTAGGTAGTGTACCAATATGACTTGG | CCTCCAACTTGCTTTATTACCAGCTAT |
| AMPL1563780 | CCATTTGCCAACCACCTTCTTAATT | CATTGCACGAGTACTCCGTGTA |
| AMPL1563778 | GTTCGCTGAATATAATAACGATAGCAATGC | GACGATCGAGTGAAGACGATGAG |
| AMPL1563775 | CACGGCGATCTCTGTGTTTATTG | TCATTGGCCCACATTAGTGCTATG |
| AMPL1563774 | ACAGTCTTACATACGTAACAATCCTTTCTC | GATCCATTATCGAGGTCTTTCCATAAGG |
| AMPL1563769 | CCGCTCCTTTCTAGCTCCAT | TGATGGTGGCTCATGGACTTG |
| AMPL1563760 | CCAGTACCTAGCCACAAAAAGGT | CCATTTGAACAGGATGGACTGGTAA |
| AMPL1563750 | CCCTAAGCTTGAGGACAACACA | TCTTACAAACTTTAGACCATTGAATCGCTT |
| AMPL1563740 | AAAGGCATGGCACTTTGTTTTTACTAG | TGGCCATCTCCATTTCGGATTG |

| | | |
|---|---|---|
| AMPL1563739 | ACTATTTGAGGGTGTCCAATTGGAC | GTCAGCCGTGAGTACACCAT |
| AMPL1563735 | CTTGAATTGGCCGATTTTGGCA | AAGAACCAGCAAAGATCTAAACCCAA |
| AMPL1563734 | CGGCAAGGTGGTGATGAAGATC | CATCATATCGCTAATCAATCAGAGCGAT |
| AMPL1563732 | GAAGCACAGTTCTCAGTTGGAATG | AAAACCGTAGCGTAGACGATATTATCTTC |
| AMPL1563731 | TCTTCGTCTGCGCGAATATTTAGA | GCAAAAATTGATCTCACTTGATGTTCAGTT |
| AMPL1563726 | ACTAGGAGGAGAACCATATTTGGCA | GGACCACTTTCAACACTAAGCAAGT |
| AMPL1563719 | GGATCGATTCCCATCCAGTCA | TCAGGCCTTGTAGAAGATCCTAGTG |
| AMPL1563717 | GCGCACATGATCATGGTTCAAC | AGGGATGATTTTACAAGGATTTCATCAGAA |
| AMPL1563715 | CACTTGAGGTGGGATTTGAATTCAAT | TCTCCACTAGTGTATCTACCTATCCATCT |
| AMPL1563699 | CTCTGAAGATCAATTTCAGTTCAGACCT | GCCTACCATACACAAAAATTTTGGACATAA |
| AMPL1563697 | GATAATATCATATGGCGGTTTAGGAGCAT | GCCCAGCAGGCCTAATGTAC |
| AMPL1563688 | CGGCCATGTACAGCTCATAC | GGTACCCGTCGAGCATGAAG |
| AMPL1563671 | CCCTAGTGGCTGTATCAAGTTACG | GGCTCTGCTGCTATCGTCATTT |
| AMPL1563668 | GTAGATAGAGAGACTGTTGGAGTTTGAC | TGTTGGATTTAAACCAAACAAAGAGAATGG |
| AMPL1563665 | TCCTCGAGAGGAAATCGATCCA | GCCTGGTCGTGCATGCTT |
| AMPL1563660 | CCAACAGGACAATGTAAATATTTGACACTG | CAGATTTTACCAGCTTCGCTTTGAG |
| AMPL1563650 | CCTCTCTCACCATTCCTTCAGTT | TTCCAGCCCAACACCTTACAGAAAT |
| AMPL1563644 | CCGGCAAAGAAAGTTGCAAGTC | GAAAGCCTGGTCATGTTGGATTG |
| AMPL1563642 | TGAGCTTCTCCATCACTCCCAT | TCAAATCGTTCGACCACGTGAT |
| AMPL1563640 | TGACCGTACAATCCTACTACTCGT | GGAAAAAATATACTACCTCGAAAAGGTTGGA |
| AMPL1563638 | GCCTTCTTGCACCGTTGA | AAGGCAGTTTCACTGACGTGA |
| AMPL1563624 | CGTCGACTCCTCCAAGGAGT | CCTCCCACGCGTACATCTC |
| AMPL1563618 | CCGTTATAACTGTGAACTGTGATATTTACG | GGCAGAGAGACCTAATTCCTAGTTGAT |
| AMPL1563617 | TGCCAAATATGTTGTCTTCTATGGTGAT | GCCAGTTTGCCTGATATCAATAAAGAAAAA |
| AMPL1563606 | CGCGCATGGATAACTCGT | CAACCTCCTCCTCCCACAA |
| AMPL1563601 | GGTGTATGAGGGTGGTGACAAG | TTGTTAGCTGAAACATTACAAAACAAACCT |
| AMPL1563599 | CAGATGGTCAGATTTGGTGCTGA | GAACCAAAGAGACCAAACTTTATGAGTTTG |
| AMPL1563597 | CCACCAATCTTGTCTTCCGGAT | AATGGTTAGCTAGCTAGTAAGAGAGTGT |
| AMPL1563596 | GCACCCATCAACGTTGGAGT | TGGAGTTCACGATCTTCTCCA |
| AMPL1563594 | AGGCTTAGGCTTAGGCTAGATAGG | GCCAAGCCCATACATGATCCTAG |
| AMPL1563593 | GAAGTTGTACAGATAGAACTTGAGCCTT | CCAGCATACCATACGACTAAATCCC |
| AMPL1563592 | AGGGAAGGGTAGAGAGAGGAAATC | CCTCGAGGAGGATGAGCTCAT |
| AMPL1563590 | CTGCAAGACTCTACTACTCCAACAC | TGTGGAAGCCATCTCACCATTG |
| AMPL1563578 | ATGCAGGAAGGAAGGAGGAGTT | CAAAAGCCCTACTACTCCTATACCTATACT |
| AMPL1563575 | ATTCTTCAAACTTTCCAGATGGGTCAT | CCACTGACGTATTTCTGAATAGCAACT |
| AMPL1563572 | CTCTCGCTATCCGAAAGAACAGA | CTGTAGATCACGTCTCTGTTAAAAGTCT |
| AMPL1563571 | GCCGATCAGGTGACCAAACTTA | CCTGTCTGAACCAGCACAGATTC |
| AMPL1563566 | GTAGCAAGCCACACCCAAGCTA | ACCACACCACGTCGATGATTTT |
| AMPL1563565 | ACACAGAGAAACATTTCAGAAAGCAGA | GTATATGTGATATGCAAGCAGCAGATTTAG |
| AMPL1563556 | CTGATCGCGACGGGAAAACCAA | GAGAAATGAAAGTACTACTGGTACAAGCAA |
| AMPL1563554 | CCCGATCCTCACTACTGTGTCA | GATCGGAGGATCGACGAAACTT |
| AMPL1563551 | AGTTTTGCTTGATCTACTGAATCACAAAAC | GTTTGTTTGTAGCAACGGTAATGGTAAAA |
| AMPL1563546 | GTATGGTATACTAGCAACGGATCTTATGC | GGTTGGATCAGTCATTTCGGCAT |
| AMPL1563542 | ACGATTGTGGTTTGGAAAATCTGC | GCTAAGGTGCCATGTTGCAATTATC |
| AMPL1563540 | GGATCAGGACAGCACAAGGAATG | GCGGATGAGATGAGAATTTTCAACTAAAAT |
| AMPL1563532 | CGCCCTGTTGAAGGAGATACAC | GGATAGGATGTTAAAGCGCCACTT |
| AMPL1563531 | CTCTCCTGGAACTCACTTGGTT | TACTACCGACCTACCGTTCACAA |
| AMPL1563530 | CATCTTCTGTCTAGCCAGGTCAA | ATGAACATTACTTCCTTTCGCTACGT |
| AMPL1563526 | GGATATTGTTGTTCTACTGACATTGCTAGT | CCAGAGAAGCATGCATACCAAATATACAAT |
| AMPL1563523 | CAACCATCACTCTGAAATCTGAACG | AGCTTGAGAGCGATTTTAGAAAAGTTAATG |
| AMPL1563521 | GCTCTCAAGTTGAATGTATAAGCTGGT | GGACAGTGTCATGTGGTGTTATAGAAG |
| AMPL1563514 | AAAACCGGCACATATAGAAAAATGAGC | CCATATGGTGTACCATCGCCAAAT |
| AMPL1563510 | GCATTCGCACGACTCTAACCATA | CGGTGAAAAGCAATGACTCATACACTA |
| AMPL1563508 | CACATCACCAATTGAATAGAGAGAGAAGA | TCACCATGGTTCAAGAGTGAAAACAA |
| AMPL1563506 | CTGCTCATCGATCACAATGTACTGT | GCATGATTGAAACAGGAAAAGTACGTAGA |
| AMPL1563505 | ACAACCACTCCTCTCGTCTCT | CTTCCTCTTCCTCTTGTTCCACT |
| AMPL1563503 | GAGTCTGTCTCAACCTGCATTTG | CAAAGATGAAACCTGGATTGGATCAGA |
| AMPL1563502 | GCTTGGCATTGAGAGTTATATATAGTTGGG | TAAATATTTTGAGCCGGAGGTAGTCTTG |
| AMPL1563496 | GGGTGTTCTTGGTCTACTTTATGGG | CTGTTGCATAGCCTCGCTTTTT |
| AMPL1563494 | ACAAACTATCTGGAAGCTTCTGGAAG | GATATTAGCAGTGGATTCATGGAGCT |
| AMPL1563487 | CCTCCTGGATCTCCTCACGAT | CCTCGGTGAGTGGTGCTTAC |
| AMPL1563486 | CCAACCAAATCTTTGACGATGCT | ATGCTACTGTGCTTTTAATTAACTTGCTG |
| AMPL1563483 | CTCGCAAAACAAATACTGGCCAT | CTGCACGCGAGCTGCAAAC |
| AMPL1563481 | CGTCGATCGAGTTGGTGACAAT | GGAGGAGGAGAGGAGATTGGATC |
| AMPL1563476 | GATTCCTTGGCCCAAAAACCAG | TCTTGTATTAACTGGGAACCAACGATAC |
| AMPL1563475 | GCCATCTAGTCCTCGGAACA | ACCACTTGAGATCAAATTTTCAAAACACTG |
| AMPL1563473 | CACAAAGATCAGCAATGGCTAGTTT | CATCACCTCTTGTACCTTGGTGT |
| AMPL1563471 | CCCTGTTGCAAGGTTCTTTGAG | GAAGCCAAATCACAGCTGCAAAATA |
| AMPL1563459 | GGAGCAGGCTCATACACACATA | GACCAAGTTCTGATCAGTTAGTGTTTTTC |
| AMPL1563452 | CTTTCCTAGTATCGCCTTGTTCCT | CGTACTGCTGCTTCTTTATGCG |
| AMPL1563450 | CCGTAGGCCGTAGCATTAATTACC | CCTTCGATATTGCTGCACCTTTG |
| AMPL1563446 | GGGAGAACTCATTTTTCCCTATAAGATCA | TGATGTTGAAGAAGTCAAACGAAGAGTT |

| | | |
|---|---|---|
| AMPL1563426 | GTAGTGATTGAGAGTAGTACTGTTCACTTG | ATCTCCTGTGGTCTGTTGAATCTATTG |
| AMPL1563421 | GGGAGAAAATTTTCACCTTTGAGACA | GCATTCTTGCATACACATTGATGTGG |
| AMPL1563416 | CGATGCATCATCATCAGACCGAT | CGAGCGAGGAGATAGGTGGATATAG |
| AMPL1563414 | CTACCTAGTACATTGTGCAATTAGTACACA | CAGCACCCTATGGCTAGTTGTAAG |
| AMPL1563413 | CAGAAAAAGCCATGCGTGAAGT | CCGTATCAAAAATTCGATGTGACATATCTC |
| AMPL1563411 | CTGCCCTGCTCTCAAGTTTCTA | GGGAGAAAAACTAACCAAGTGTACTCTTAT |
| AMPL1563408 | CTGGCACAGGAATGGATGAATG | CTCAAACGCACACGATTTAACAATTG |
| AMPL1563407 | GGACGATGTTGCGGAGATAGAG | CCTCAAACATCCTCTTTCCCATCA |
| AMPL1563406 | GTGGCGAAATCCAGGCAGAT | GTCGTCGGTTATAAACTTATAATGGTCTGA |
| AMPL1563405 | GCGTGGAACTCCAAGATCTTCC | AAGCCGATCAGGACCGTAAAG |
| AMPL1563402 | GACCTTTCACCACCAACAACAC | GAGGAGATGAAGAAGAATGCCCAA |
| AMPL1563401 | CCCAAATCACATGAGGCCCATT | GAAATCGCACCATGCATCGATC |
| AMPL1563393 | GTGGAGCTCCGAAGAGAGGAA | GATCCGACGAGCAGTGGATAA |
| AMPL1563391 | AACTTGGAGACACACTAGTCTGGTA | AACTCAAACGTACGACAATCGATTTTTAAA |
| AMPL1563386 | GCGCATATTTCTGATTCCGTCATG | GTATAGCTGGACTAGCATGTCAGTTAC |
| AMPL1563384 | TGTTGTTGTTGCGTCCCTCTA | CCCGAGCTGGAATGGACTAC |
| AMPL1563383 | CCCAGTACGGCAACAAAATCCT | GCCCATTTGGTCACACAGAAAAG |
| AMPL1563379 | CAAATCATTAGGAAGGCAAGCAATGA | CATTCGAGTGTATGGTCAGGTTGAT |
| AMPL1563374 | TCATATGCCACAGCTCCATAGTTTG | GCTCCCTCGATGGTTTCTTACG |
| AMPL1563372 | CCGAGGGTTAGCTTGTTAGTGC | GAACAAGACAAGAGAAGCAATATTTGGTG |
| AMPL1563371 | GCGCATTCATGTCATCGTCTCA | ATCTTTCCAGTTTTTATTGCTCAAAGCTT |
| AMPL1563370 | GACTTCGTTTCAAGTGGTAAACCTG | AGCCTATTCTTGTTTTCCTTCAACATCT |
| AMPL1563368 | CGCCAGGATGATGCTCGA | ATGCCAACACGGTTCATTTCAC |
| AMPL1563367 | TGGAAATTTTGTCCATGTCCTTTTGAAC | ATCAGCGCAATCAAACTTTGGAATATTTAA |
| AMPL1563355 | CACCTCCTCGAAGAACTGCT | CCGCCATTAAAATTGCTCCAAG |
| AMPL1563354 | CCACAGAGTAGCATCAAACGCA | GCAAAGTTTATGCTCACTTTGCCA |
| AMPL1563350 | CATGCCTGCTATACCTTCCTGA | CATCTCTAGCTACTAGTTGTTCTCTTCTCT |
| AMPL1563349 | ATAGACAGACAGAACAGCGGTCAA | GGAGAGATGGATCGGTGCATAG |
| AMPL1563346 | TCTGCTCCACATTTCTCTTGGAAG | GATAATCGGTGTAGCACATGTTTGG |
| AMPL1563342 | CATCCATGTATGGAAGATTACATCTCACA | TCTCTGCTCCATATAAGAATAGACAACTGA |
| AMPL1563341 | CATCTACCAAGCATGCCCAGTTA | GACGACAAGCTCGGAAACAATC |
| AMPL1563339 | TGTGTTCTGTTTGGAATTGTAGGTCA | GTGGCACTATTATTGCGTTGCATT |
| AMPL1563338 | GGGAGATTACATCGAGGGACAAGA | ACGCACAAGAAAAACGCCAAATATTTA |
| AMPL1563335 | ACATATGGAGGCCAGGCTACAT | CGACGGTGAGATTTGATAATACCATGTAG |
| AMPL1563331 | GCAGAGGAAAGATCACCTCCAA | CCTGACAAGGAGAAGGCATGAAG |
| AMPL1563330 | GCTTAGGGAGCGTCTGTAGGAA | CCAACACAAAGGACTAAGATTTGTGACATA |
| AMPL1563328 | CCAACACGATCGATACAGCAAAC | TGGGAGAAATTGCTCTTCATTTTCCTAG |
| AMPL1563323 | TGCATGGCTATCTCTTTTGTTCTTTTC | TCTGTGGCATTTATGCTTTTCCCT |
| AMPL1563321 | CTCCCATCTTTTTGCTGCAATCTTT | GCGATCGATCGACCTAGCTATTG |
| AMPL1563309 | CCACACGACACGAGCTAGTA | GCCATCGCCCTTTTCGTCTT |
| AMPL1563305 | GACGTCAATTCATTTTTCGATCAATTTGC | TGTGAATGAAGGCGAATGGTGT |
| AMPL1563302 | CCCTCCCTCTCGATCTAAAACC | CACAAGCTCTTGAGAGACGGAAA |
| AMPL1563300 | CTATCTTGGCGTCCATCTTGAGT | GTTGAATCAAATACAGTTCGTTTCAGGTT |
| AMPL1563293 | CCATCTCCATCGTTTCGCCAAT | TACTCACACACTGATCACAAAGCAAA |
| AMPL1563287 | AGCATCTCTTTGTACCAAAATTTTTCCTC | TCTGCTTGTCCCAAATCACCA |
| AMPL1563283 | CGGTGGAATTCGTTGAAGGTCA | TTACCCATGCGTTTAACTATTCTAAGCA |
| AMPL1563280 | GCCCGATTTATTCTATTAAAATCTCAAAGG | CGTGGGAATGTGGTTGCAAAAG |
| AMPL1563278 | CTAGTGATGAGAACCGTATCAGAAAAACT | GTGGCGGTGATGTCAGCT |
| AMPL1563277 | TCATGTTCAAACGAGCCCAAGAA | GAGGCTGTGGGTCATTCTTCTC |
| AMPL1563273 | GCCTCATGAACACAATCACAAATCA | CTACCTCTCCCTCTCGCCTT |
| AMPL1563271 | GAAGAAAAATTCGAGCAATTAAAAAGGAGC | CTCTCAAGTTCTCGACGAGGAA |
| AMPL1563270 | GCAGGTGGAAGGAATGTTTAGAGA | GATGAACAGACCAAGCCAAGCTA |
| AMPL1563269 | GTGTTTTCTGGATCCTTTATAGTGAAGTG | AGTTAATGAAAGCAAATAGTTACAAGCACT |
| AMPL1563266 | AATCATAAGAAGCGCATAGGCCTT | GCAGTTTTCTCATATGATTCCGTTCTATGT |
| AMPL1563265 | TGTGGGCACATATGGGTCTCATA | CGCTGGGATGTGTTAGGGTTTT |
| AMPL1563264 | TGTTTGCGCGAAATATAACTTTTCCC | CCTAGTGATGAATACAAGTGTCTATGGGA |
| AMPL1563262 | AGTTGTTTATTAAAGTGACACGTGGC | AAAAGGAACTAAAAGTTCTTTCATAACGTCAAC |
| AMPL1563249 | TAGAGTAACATGTGGCAAGTTAAGAGC | TCCTTTTGTTTCACAGCATTTTCCATG |
| AMPL1563248 | GTGGTTTTAGGCCTGTCAGGTA | TACTGGTATTGGTTCTTGTGGTTTTTGT |
| AMPL1563247 | TTTTTGCCTTGGTTTTTATCGATTTTGC | CCCTCCCACCCTTCTCTTGTTA |
| AMPL1563246 | CCCTGTCTCACGACCATTATTAAACT | ATTTCTGGTGTTTGATGTCTGAAGGA |
| AMPL1563243 | CTCGGTAACTGTAAGCTCCTCAT | GTCTCGTTGATGGGCGTAGATTT |
| AMPL1563241 | AGCAATTTCTCTCTCACACATGCA | GTACTTAACACTGACATGCATGAGTG |
| AMPL1563239 | GATAGCACTAGCCATTATTGAGTTTCTGT | CTACGCTGCTCCAGATTAGGAG |
| AMPL1563237 | TGACGATGCAAGCCGACAAATA | CGATCATATGGAAGCAAAGAAAAGCTTAAT |
| AMPL1563235 | AGATCAAAATGATGAGGATCAACTACTTGG | CAGCACAAGGATCGGAAGTAGAC |
| AMPL1563230 | GCTTCTCCTTGTCTCGAAGCAA | TCCATCCACCCAGAGTAAAAGGA |
| AMPL1563229 | CTCCTCCTCCTCCCAAACAATG | GTTGTCTTCTCCTCCACCCAAAA |
| AMPL1563228 | GAAGAGGTGGAGGCCATGT | GAGACGACGAAGGCAGATAAGTC |
| AMPL1563226 | TGAGCAGAAGCACTCTAGTATATTTAGCT | GCACATGTTCGTCGACCAAGAA |
| AMPL1563224 | AACTCCTCCTCCTCATCACGT | GCCGGTGTGCTGTATGTAGC |
| AMPL1563221 | AACCACTTAGATGATATGCAATAAACAGGT | GCTATCCCGTAGTATTAAACAATCGGT |

| | | |
|---|---|---|
| AMPL1563217 | CATGCACACTATTGTTATAGATGCCATTG | CCGTATCTTCATGCATCAACATATCATGA |
| AMPL1563216 | CAAACTTGCTGCATACACACCTTAAT | ACCCTAACCCAACCTAACCAAC |
| AMPL1563211 | AAGTGAAAAAGAGAAAAGGTAGAAATGTGC | GCAACGTTGAATAATCGTTCAAAATCAGTA |
| AMPL1563210 | CCGAAGTAATAGTCGGTAAACTCATCC | CAGCAGGTAGAGTTTCCACTCATAG |
| AMPL1563209 | AAGAGGCACTGAAAAATACCTCAAGATT | TCAGATTTGACAGTTTCACAAAGAGGT |
| AMPL1563206 | GGTGGGAGGTTTTGGTTTTAAGATAG | GAGTCTATGAGTATAGTTGCCTTGGTTAC |
| AMPL1563204 | GTCACGGTCTAAAGAAAAGAGACGT | GCCCACCAATAGTCAGGTCATG |
| AMPL1563202 | TCTATGTATCGCTCGACAGATTTTTCC | TGATTGGTTGGCTATCGTTTGAGAG |
| AMPL1563201 | GCCTTGACTCTTCATCGCAAAC | CAGATAACGAAATCTGATCAACTCTGCTAA |
| AMPL1563190 | ACCACTACCAGCCAGCCTG | TCGTCGAGTTACAAAACGTCCC |
| AMPL1563183 | CTATATGCTTTCTGTAATGCAATGCCA | ACCTTCTTCATATAATGTCAAAGGTTGGAA |
| AMPL1563181 | AATTGAATAGTTTGCAATGGCATGACA | TGGCTGTTAAAGAAGCAAGACTGAA |
| AMPL1563177 | ATGCATGCAAACCCTAGCTACA | TATAGCTAGCAAGCAGCAAGCGAA |
| AMPL1563171 | GCTACACGCACCCGATACTTTC | CTTCTCGGAGAGCTTCTCCATC |
| AMPL1563170 | ACCTGTCACCTTTACTGTGATCTATTTG | CGGGCAGGTATTGGGATTATTTATTAATCC |
| AMPL1563167 | TGAGTGGATCATAAAAGGTTGAGACATC | CCGTGGCATCTTTACTCTGGAAA |
| AMPL1563165 | TGGTTGTGTGGGTCATGTCAAG | CCATAAACCGGCACTGTGTAATTCT |
| AMPL1563164 | GACCAAACCTGGTTAGCATGATATAACT | TGAACTTCCAGAAGATTTGTAATGTAGCTT |
| AMPL1563160 | GGTGTGGTACCTTTATTGATGCCT | GCATCAACAAATACAAATGCAACAAATGTT |
| AMPL1563155 | TCTCTCCGTCGATGGGAGAT | CTAGTAATCAGTAGCGGAAGAACACTC |
| AMPL1563151 | ACACGTGCAGTAGTGTCTACTG | TGAAAACGACCGGAATATTCTCACC |
| AMPL1563150 | CATCTAACTAGGGTATAGGTGGCAGAT | GACGGAGGGAGTAGTTCATGG |
| AMPL1563149 | CTCTTTACTCACACTCTGCTCGAA | GTCTGTCTTCCTCTCCAAAATAATTCAAGA |
| AMPL1563148 | CATCGAATATCTCTCTTACTCACTACTCGA | ACTTATTTGCTTCCAAGAAAGACTCCTT |
| AMPL1563146 | GTTTATCATCTGCAGAATATGAACGCA | TCCAAATAAATGAGAGGGAGAAATCAATGT |
| AMPL1563145 | GGGAGTAGTTTGTAATCCGTACATGA | GAGAAGGATCCTGATTCCACGAATC |
| AMPL1563144 | CCTGACCTCAGTATCAAGGACGA | GGACAACATAGCGAATTATCTAGTCGATC |
| AMPL1563142 | AAGCATCAATGAATTCGAATGGTCAAG | TCATCTCCCTGTCGGACTTGT |
| AMPL1563141 | GCTGGTCAAGCCACTCAATCTC | AATCTCTATGCCTCCCTCTCACT |
| AMPL1563140 | GGAAGGTGACGTACGTGTGTAT | GGTCGTAATCAGGGAAAAAGAAAAAGC |
| AMPL1563139 | GTCAACACTCTAGCCCTCACAT | TACTGTTCACTCTTTCTGTGGATTGAATC |
| AMPL1563136 | CCACTCTGTCTTTTTCCTTTTTAAAAGAGA | GTTTGGCTCCAATCTAAATTAGGAGC |
| AMPL1563135 | TGATATCATGGATATGCCACCGTTT | ATGAAGAAAAGGCAGGATCATATGCT |
| AMPL1563132 | CTGCGTCCCGACAAGCAAAA | CGCGTTGTAATCGCCTCAAA |
| AMPL1563131 | GAGGAGCTATAGCTACTCATATTATCCTGA | ACTGGTGTGATTACTGTTTCTCTGTTT |
| AMPL1563130 | TCCTCTCCAGAAACCAAGACCT | GGGCGTGTCATGTATAGATAGCG |
| AMPL1563129 | GATAAGGGAAAAATAAACCTTGGATGCA | TCTCTAGCCCTCAGAGCAGTAAG |
| AMPL1563128 | CCAGGTCTCGTGCAATGTTCTT | ATGGACCACAAACGACCTTCTTAG |
| AMPL1563127 | CTGTAAATACATACAGTGGATCATACAGGA | ATTTGTTCATTCGGGTTGCTTTTGT |
| AMPL1563125 | TCGATGTCTGTTCATTGTTCGTTTG | CAATTTGTGTTTTGTTAGTTGGCGAAAG |
| AMPL1563124 | GGTAGTACCAGGAGACCGTGTA | TCACTGTCTTTGTGCTCTCAATTCTC |
| AMPL1563104 | CTCAATGCCAACTTGATATATATCCGGT | GTGACGGAGTAATTGTGTTGCA |
| AMPL1563098 | GGACGATCGCCCTGTACAAG | CTCAAGGCTCACCTTATTATACTAATCTGC |
| AMPL1562999 | GTGAGACCTTGTAGTGATGTGCTATTTA | CGGAGAGGAAGAACAGAGAGTTTC |
| AMPL1562814 | AAATATTGCTCAAATTTCCAATGTTGATCG | CACGTTTGGAATTTCGGTTCGAAAT |
| AMPL1562757 | GCTTCTCCTGCAGGTTCATCTC | CATATGGTCAGCGACAGGTACA |
| AMPL1562697 | GGCCCTCATCACCTTCGTAG | GTCGTCCTCCCTCTCCCTATC |
