## Supplementary figures and images for "Multiple nucleotide polymorphism DNA markers for the accurate evaluation of genetic variations"

### Supplementary Figure 1

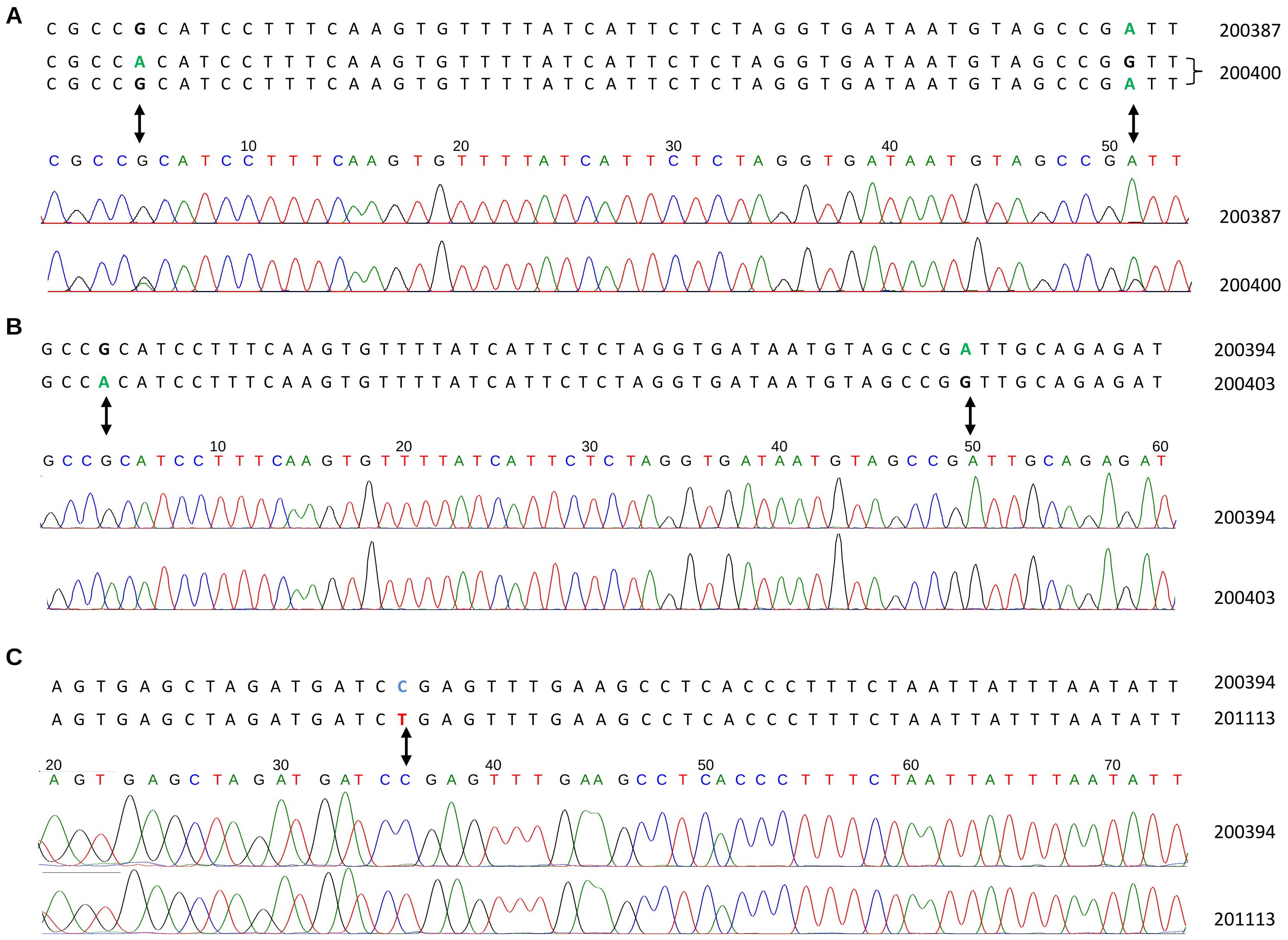


**Supplementary Figure 1.** Validation of distinct MNP genotypes between Nipponbare lines.
